# Transcriptional organization and regulation of the *Pseudomonas putida* flagellar system

**DOI:** 10.1101/2021.08.29.457346

**Authors:** Antonio Leal-Morales, Marta Pulido-Sánchez, Aroa López-Sánchez, Fernando Govantes

**Affiliations:** Centro Andaluz de Biología del Desarrollo, Universidad Pablo de Olavide/Consejo Superior de Investigaciones Científicas/Junta de Andalucía and Departamento de Biología Molecular e Ingeniería Bioquímica, Universidad Pablo de Olavide, Sevilla, Spain.

## Abstract

A single region of the *Pseudomonas putida* genome, designated the flagellar cluster, includes 59 genes potentially involved in the biogenesis and function of the flagellar system. Here we combine bioinformatics and *in vivo* gene expression analyses to clarify the transcriptional organization and regulation of the flagellar genes in the cluster. We have identified eleven flagellar operons and characterized twenty-two primary and internal promoter regions. Our results indicate that synthesis of the flagellar apparatus and core chemotaxis machinery is regulated by a three-tier cascade in which *fleQ* is a Class I gene, standing at the top of the transcriptional hierarchy. FleQ- and *σ*^54^-dependent Class II genes encode most components of the flagellar structure, part of the chemotaxis machinery and multiple regulatory elements, including the flagellar *σ* factor FliA. FliA activation of Class III genes enables synthesis of the filament, one stator complex and completion of the chemotaxis apparatus. Accessory regulatory proteins and an intricate operon architecture add complexity to the regulation by providing feedback and feed-forward loops to the main circuit. Because of the high conservation of the gene arrangement and promoter motifs, we believe that the regulatory circuit presented here may also apply to other environmental pseudomonads.

**ORIGINALITY-SIGNIFICANCE STATEMENT:** This is the first integrative study of the flagellar transcriptional cascade in *Pseudomonas putida*. Our results provide a new transcriptional organization featuring several operons with a nested architecture, detailed regulatory characterization of twenty-two flagellar promoters and a novel hierarchy for the regulatory circuit of the flagellar transcriptional cascade. The results presented represent a significant departure from previous models for other related bacteria. High conservation of flagellar gene organization and promoter sequences suggest that our observations may be relevant to other pseudomonads.

## INTRODUCTION

Motility allows bacterial cells to colonize new niches and escape from unfavourable habitats, minimizing the negative effects of competition and depletion of local resources (McDougald *et al*., 2012). For many bacteria, motility is directed by flagella, complex molecular machines that enable swimming of individual cells in liquid and semi-solid media and collective swarming on humid surfaces by engaging in propeller-like rotation driven by the proton-motive force (Altegoer *et al*., 2014; Kearns, 2010). The flagella-associated chemotaxis machinery, comprised of membrane-bound and cytosolic chemoreceptors and associated signal transduction proteins, promotes directional motion by controlling flagellar rotation in response to chemical gradients of attractants and repellents (Sourjik and Wingreen, 2012).

Bacterial flagella are proteinaceous appendages composed of over 30 different polypeptides (Altegoer *et al*., 2014). Flagella are constituted of three functional units: the basal body, a rotary machine embedded in the cell envelope; the filament, a long and thin helical appendage that protrudes from the cells and acts as a propeller; and the hook, a rigid, curved element that connects basal body and filament. The basal body is the most complex of these units. In Gram-negative bacteria, it is comprised of five separate components: (i) the MS-ring, an annular structure bound to the cell membrane, provides support to the complete flagellar architecture; (ii) attached to the MS-ring, the cytoplasmic C-ring is the flagellar rotor and the target for functional regulation by the chemotaxis machinery; (iii) a hollow rod anchored to the MS-ring spans the cell envelope, connecting the basal body with and transmitting rotation to the outer components of the flagella; (iv) the double LP ring is bound to the peptidoglycan and outer membrane and acts as bushings for smooth rotation of the rod; finally, (v) the in-built flagellar type III secretion system (FT3SS) provides the functions necessary for the export of flagellar proteins through the flagellum itself. Associated to the periphery of the flagellar basal body, the stator complex fuels flagellar rotation by coupling inward proton or sodium transport to the generation of torque (Chevance and Hughes, 2008; Terashima *et al*., 2008). As documented thoroughly in *Escherichia coli* and *Salmonella*, flagellar assembly is sequential, with the inner components of the basal body being assembled first, followed by components of the rod, LP ring and hook, which cannot be exported until the functional FT3SS is completed. Subsequent assembly of the filament proteins is preceded by a shift in FT3SS specificity triggered by completion of the hook structure. Finally, the accessory stator and chemotaxis complexes are assembled. The structure and assembly of the flagella have been reviewed thoroughly elsewhere (Altegoer *et al*., 2014; Chevance and Hughes, 2008; Evans *et al*., 2014).

Gram-negative bacteria differ with respect to the number of flagella, their distribution across the surface and the transcriptional organization and regulation of the flagellar genes. Although flagellar genes are scattered in the genomes of some organisms, most often they are grouped in one or a few genomic regions (Smith and Hoover, 2009). *Pseudomonas putida* is a well-studied soil bacterium endowed with extraordinary metabolic versatility that has become a model organism for biodegradation of organic toxicants and an attractive platform for biofilm-based industrial applications (Gross *et al*., 2007; Li *et al*., 2007; Martins Dos Santos *et al*., 2004). *P. putida* shows lophotrichous flagellation, consisting of a tuft of 5 to 7 flagella at a single pole (Harwood *et al*., 1989). In the reference strain KT2440, most structural components of the flagella, the core chemotaxis signal transduction system and a number of dedicated regulatory proteins are encoded in a ∼70 kbp chromosomal region known as the flagellar cluster (**Fig. 1A**). Additional genes encoding a second stator complex (Toutain *et al*., 2005), several paralogs of the core chemotaxis components (García-Fontana *et al*., 2013) and a set of 27 chemoreceptors (Corral-Lugo *et al*., 2016) are scattered elsewhere in the genome.

**Figure 1.**
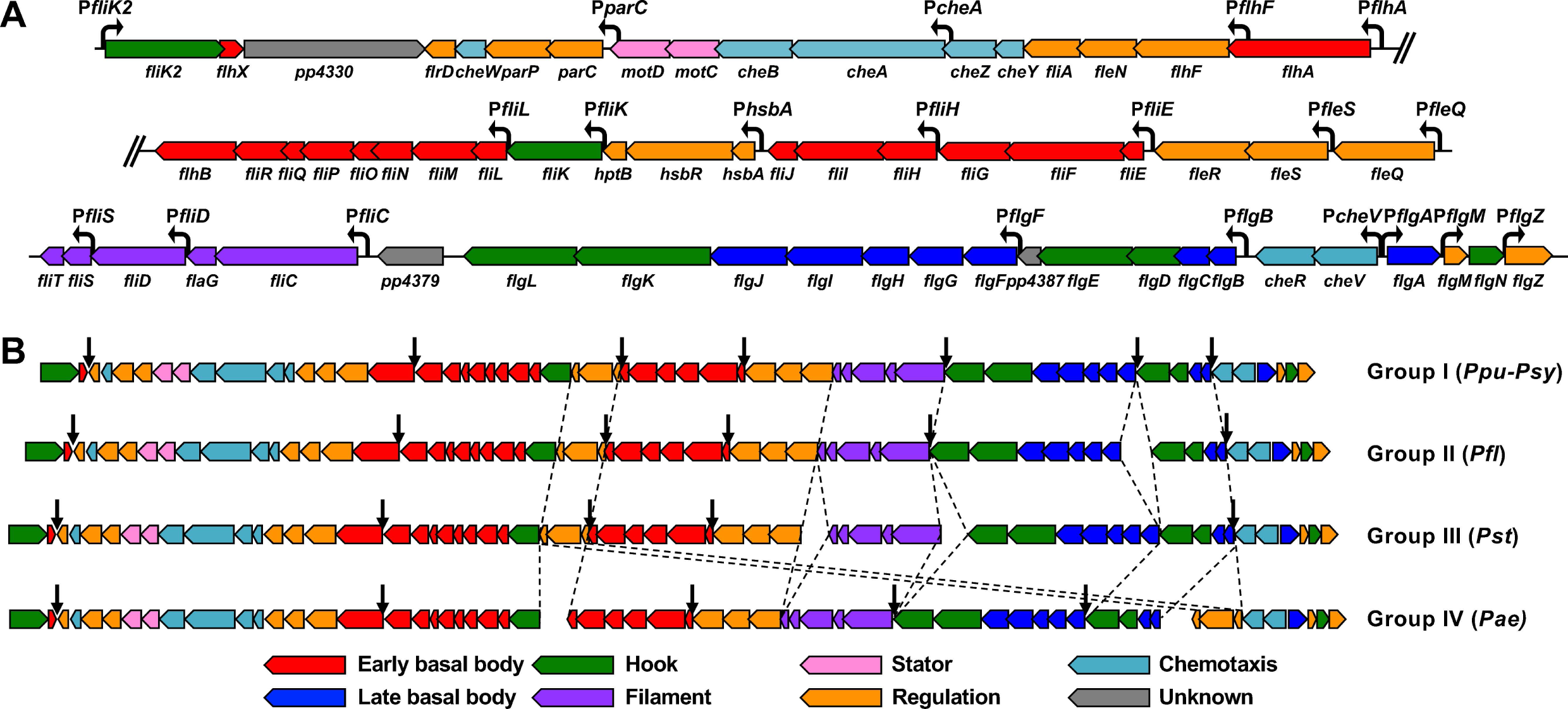
The *Pseudomonas* flagellar cluster. Synteny relationships between the 59 flagella-related genes in the *Pseudomonas* flagellar cluster. Genes are color-coded to denote their putative function as indicated. **A.** Cartoon of the *P. putida* KT2440 flagellar cluster, including designations of all 59 flagella-related genes and the 20 promoters identified in this work. The 7-gene insertion (PP_4345 to PP_4351) upstream from *flhA* is omitted (indicated by two oblique lines). **B.** Simplified cartoons of the *Pseudomonas* Group I, II, III and IV flagellar clusters. Dashed lines indicate points of synteny disruption between groups. Black arrows indicate preferential locations for insertion of small groups of additional genes.

The biosynthesis of bacterial flagella is a complex and ordered process that mirrors the assembly sequence (Smith and Hoover, 2009). In order to achieve the correct temporal regulation of flagellar genes so that the structural protein are produced as they are required for assembly of the nascent flagellum, bacteria use hierarchically organized regulatory circuits in the form of a transcriptional cascades (Smith and Hoover, 2009). Little is known about the transcriptional organization and regulation of the flagellar genes in most members of the genus *Pseudomonas*, and much of it has been extrapolated from the transcriptional cascade of *Pseudomonas aeruginosa* described in the seminal work of Dasgupta *et al*. (2003). In this organism, the flagellar genes are organized into four classes to reflect the timing of their expression and the transcription factors involved in their regulation. Class I genes, encoding the master regulator FleQ and the flagellar *σ* factor FliA, stand at the top of the hierarchy and are not regulated by any of the dedicated flagellar transcription factors. Class II genes encode the cytoplasmic and membrane-bound components of the basal body, the FT3SS, the two-component system FleSR and the auxiliary regulatory proteins FleN and FlhF, involved in control of the number and polar location of the flagella, respectively. Genes in this class are activated by FleQ in a σ^54^-dependent fashion. Class III genes encode the structural components of the rod, the outer LP ring and the hook, and their expression is also σ^54^-dependent and activated by FleR. Finally, Class IV genes, encoding the filament proteins, the core components of the chemotaxis apparatus and the stator complex are dependent on the flagellar *σ* factor FliA. In *P. aeruginosa* FleQ activation of the Class II genes is antagonized by the auxiliary protein FleN and the second messenger c-di-GMP (Dasgupta *et al*., 2000). The histidine kinase FleS likely activates the response regulator FleR by phosphorylation (Dasgupta *et al*., 2003), although the involvement of FleS in flagellar gene regulation has recently been questioned (Zhou *et al*., 2021) and the signals and relevant conditions have not yet been explored. FliA sequestration by the anti-*σ* factor FlgM prevents premature activation of the Class IV promoters (Dasgupta *et al*., 2003; Frisk *et al*., 2002). FliA inhibition is likely released by FlgM secretion through the FT3SS upon completion of the hook, as shown first in *Salmonella* (Karlinsey *et al*., 2000), although experimental evidence of this phenomenon in *P. aeruginosa* is still lacking. In addition, the anti-*σ* factor antagonist HsbA has been shown to prevent FlgM inhibition by a partner-switching mechanism. HsbA activity is in turn regulated by HsbR and HptB in response to as of yet unknown signals (Bhuwan *et al*., 2012). The *P. aeruginosa* flagellar system and its regulation was reviewed recently (Bouteiller *et al*., 2021).

The *P. putida* KT2440 genome encodes all the known elements shown to be involved in flagellar regulation in *P. aeruginosa*, and previous work from our group and others has confirmed the role of FleQ as the master regulator of the flagellar cascade and the role of the *σ* factor FliA activating the expression of some late flagellar genes (Blanco-Romero *et al*., 2018; Jiménez-Fernández *et al*., 2016; Rodríguez-Herva *et al*., 2010; Wang *et al*., 2018). Recently, ChIP-seq studies suggested that FleQ may bind *in vivo* upstream from genes encoding components of the basal body (FlhA, FlhB, FlgA, FlgB, FlgF, FlgG), the hook (FliE, FliK), the motor protein MotB, the anti-*σ* factor FlgM and some chemotaxis elements (CheV, PP2310, PP4888) (Blanco-Romero *et al*., 2018), but the outcome of such interaction in terms of transcriptional regulation has not been explored. Our recent work has clarified the transcriptional organization, promoter arrangement and regulation of the *flgAMNZ* and *flhAF-fleN-fliA* operons (Navarrete *et al*., 2019; Wirebrand *et al*., 2018). Although an integrated model of the *P. putida* flagellar cascade comparable to that of *P. aeruginosa* is as of yet unavailable, the preliminary work suggests significant differences in the transcriptional organization and regulation of the flagellar genes between both species. Significantly, the flagellar/chemotaxis system of *P. putida* and other so-called environmental *Pseudomonas* differs from *P. aeruginosa* in the type of flagellation (lophotrichous *vs.* monotrichous) the identity and functionality of the chemoreceptors. These differences may correlate with the different requirements of the *P. aeruginosa* pathogenic lifestyle *vs.* the free living soil/saprophytic lifestyle of *P. putida* (Lacal *et al*., 2013; Matilla *et al*., 2021; Sampedro *et al*., 2015)

The present study clarifies the transcriptional organization of the flagellar cluster and the regulatory circuit controlling flagellar synthesis in *P. putida*. Our results support a transcriptional circuit in which the temporal regulation of the flagellar components is achieved by the combination of a three-tier transcriptional cascade and several feedback and feed-forward loops to fine-tune gene expression according to the assembly status and relevant physiological and environmental cues.

## RESULTS

### A cluster of 59 flagella-related genes is highly conserved in *Pseudomonas* spp

In order to identify the complete set of genes related to flagellar motility and chemotaxis within the flagellar cluster, we have revised the annotation in the light of recent experimental evidence. ORF PP_4328 encodes a protein of unknown function containing the domain designated “flagellar hook-length control-like, C-terminal” (InterPro accession: IPR021136), found in the hook length control protein FliK. Transcription from the upstream promoter region is dependent on the flagellar *σ* factor FliA in *P. putida* (Jiménez-Fernández *et al*., 2016). Accordingly, we have designated this ORF *fliK2*. The ORF immediately downstream, PP_4329, is an ortholog of *flhX*, a gene identified in several bacterial genomes whose product resembles the C-terminal domain of the FT3SS protein FlhB (Pallen *et al*., 2005). A function of FlhX as a partial replacement for FlhB function has been documented in *Helicobacter pylori* (Smith *et al*., 2009; Wand *et al*., 2006). The *P. putida* PP_4329 product bears 34% identity and 56% similarity to that of HP:1575, its *H. pylori* counterpart. PP_4331 encodes an ortholog of *Vibrio cholerae* FlrD, an auxiliary protein to transcriptional regulation by FlrBC, the *V. cholerae* counterparts of *Pseudomonas* FleSR (Moisi *et al*., 2009). Similarly, PP_4333 and PP_4334 encode orthologs of the *Vibrio* spp. ParP and ParC proteins, required for polar localization of the chemotaxis complexes (Ringgaard *et al*., 2018). Sequence alignments revealed 32%, 44% and 51% identity and 53%, 63% and 70% similarity between the PP_4331, PP_4333 and PP_4334 products and the products of their *V. cholerae* El Tor strain ATCC 39315 orthologs, VC_2058, VC_2060 and VC_2061. Finally, PP_4362, PP_4363 and PP_4364 encode orthologs of *P. aeruginosa* HptB, HsbR and HsbA, involved, along with the anti-sigma factor FlgM, in posttranslational regulation of FliA (Bhuwan *et al*., 2012). Sequence alignments showed 51%, 62% and 64% identity and 66%, 74% and 79% similarity between the products of PP_4362, PP_4363 and PP_4364 and their *P. aeruginosa* PAO1 counterparts PA_3345, PA_3346 and PA_3347 Our results suggest that at least 59 of the genes present in the cluster are potentially related to the flagellar function (**Fig. 1A; Supplementary Table S1**). In contrast, we have been unable to establish such functional relationship for the seven consecutive ORFs (PP_4345 to PP_4350 and PP_5673) with unusually low G+C content located upstream from *flhA*, and the single ORFs PP_4330, PP_4379 and PP_4387 (**Fig. 1A**).

We have examined the conservation of the flagellar cluster in 550 genomes available online at the *Pseudomonas* Genome Database (Winsor *et al*., 2016); available at <u>pseudomonas.com</u>). Precomputed ortholog analyses revealed that 51 of the genes were conserved in all the genomes analyzed, 7 were conserved in over 90% of the genomes, and a single gene, *fliT,* for which members of two separate ortholog groups were found, was conserved in 80% of the genomes. High conservation of the 59-gene set likely suggests that it encodes an integral part of the flagellar apparatus in the vast majority of members of the genus *Pseudomonas*. Synteny analysis performed on 401 of the genomes revealed nearly universal conservation of the gene order and orientation of the flagellar cluster, with very few exceptions (complete analysis in **Supplementary File S1**). Based on the chromosomal location of the flagellar cluster genes, we have defined four groups encompassing most of the genomes analyzed (**Fig. 1B; Supplementary File S1**). In Group I, including multiple *P. putida* and *Pseudomonas syringae* strains among others, all genes are found in a single chromosomal region in the same order and orientation as in *P. putida* KT2440. Group II, including strains of *Pseudomonas fluorescens* and related species, bears the flagellar cluster split in two separate regions, with the split point between *flgF* and *flgE*. Exceptionally, the *fliK2*-*flhX* gene pair was not linked to the cluster in some members of groups I and II (e.g., *P. syringae* and *Pseudomonas savastanoi* strains). In Group III, which includes most *Pseudomonas stutzeri* strains examined, the cluster is disrupted between *fliC* and *flgL*, and in ∼70% of the cases, also between *fleQ* and *fliT*, resulting in the location of their flagellar clusters in two or three separate chromosomal regions. Finally, Group IV, encompassing most *P. aeruginosa* strains, displays the only major disruption in the synteny of the cluster: the genes *hptB*, *hsbR* and *hsbA* are found downstream from *cheR* in one chromosomal location, while segments containing *fliK2* to *fliK* and *fliJ* to *flgB* are located in two additional chromosomal regions. A similar analysis revealed that orthologs of *motA* and *motB*, the only structural flagellar genes not linked to the *P. putida* flagellar cluster, are present in nearly all the *Pseudomonas* genomes analyzed, in a separate chromosomal location, in the order *motAB*, and almost invariably linked to the gene *rsgA* and the orthologs of the KT2440 ORF PP_4906 (**Supplementary File S1**).

Flagellar clusters generally include additional genes; some are related to covalent modification of the flagella or are the result of duplication of specific genes in the cluster ( Dasgupta *et al*., 2003), while others appear unrelated to flagellar function. Synteny analysis also revealed that insertion of additional genes occurs preferentially at some of the split points noted above (*flhB2*-*flrD*, *hsbA*-*fliJ*, *fliC*-*flgL*, *flgF*-*flgE* and *flgB-cheR*), along with the *flhA*-*flhB* and (less commonly) the *fliE*-*fleR* intergenic region (**Fig. 1B; Supplementary File S1**).

Taken together, the results of our analysis suggest that *Pseudomonas* genomes have evolved to preserve the sequence and organization of a set of 59 genes related to the flagellar function. Accordingly, most genes in the cluster are found in the same genomic neighborhood in all the genomes analyzed, and disruption of synteny occurs almost exclusively in a few specific locations.

### Transcriptional organization of the *P. putida* KT2440 flagellar cluster

As a first approach to the identification of the transcriptional units present at the KT2440 flagellar cluster, we analyzed the coverage obtained in a strand-specific RNA-seq experiment (**Fig. 2A, C and E**). We observed continuous, uninterrupted strand-specific transcription, as expected for genes co-transcribed as an operon, in ten regions containing flagellar genes within the cluster: *fliK2*-*flhB2* and *flgAMNZ* at the top strand, and *flhAF-fleN-fliA-cheYZAB-motCD-parCP-cheW-flrD*, *hsbAR-hptB-fliKLMNOPQR-flhB*, *fliHIJ*, *fliEFG*, *fliC-flaG-fliDST-fleQSR*, *flgFGHIJKL*, *flgBCDE-pp4387*, and *cheV-cheR* at the bottom strand. Continuous bottom strand transcription was also observed at a region spanning *motA* and *motB* (**Supplementary Fig. S1**). Since the neighboring genes are transcribed from the top strand, these results suggest that *motA* and *motB* are co-transcribed as a bicistronic operon.

**Figure 2.**
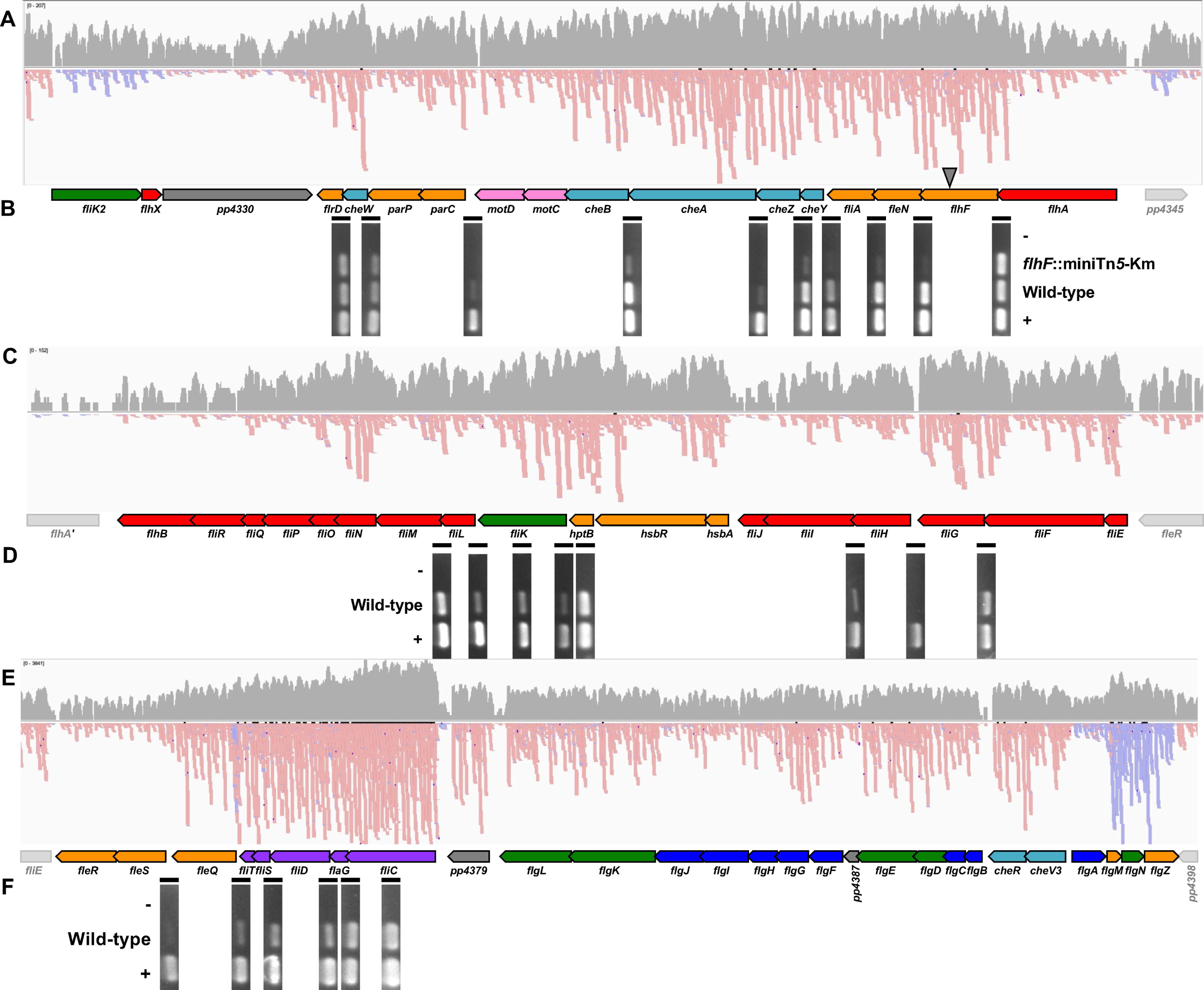
**RNA-seq and RT-PCR analysis of the transcriptional organization of the *P. putida* KT2440 flagellar cluster. A.**, **C.** and **E.** Output of the RNA-seq analysis of the KT2440 flagellar cluster, split in three regions. Coverage is represented in grey on a logarithmic scale. Top strand and bottom strand reads are shown in blue and pink, respectively. **B.**, **D.** and **F.** Agarose gel electrophoresis showing the RT-PCR results for selected regions of the *flhAF-fleN-flhF-fliA-cheYZAB-motCD-parCP-cheW-flrD* (**B**), *hsbAR-hptB-fliKLMNOPQR-flhB*, *fliEFG and fliHIJ* (**C**) and *fliC-flaG-fliDST-fleQSR* (**F**) operons. Plus (+) and minus (-) signs denote PCR reactions performed with genomic DNA and RNA, respectively, as templates. A cartoon of the flagellar genes is aligned with the RNA-seq reads and RT-PCR results. Genes are color-coded as in Fig. 1. An inverted triangle denotes the location of the *flhF*::miniTn*5-*Km insertion. The RNA-seq images represent data from a single representative replicate. Each RT-PCR image was cropped from a single picture and represents consecutive lanes of a representative agarose gel.

In contrast to our RNA-seq results, the fourteen-gene string running from *flhA* to *flrD* is broken into seven separate transcriptional units in *P. aeruginosa* (Dasgupta *et al*., 2003). On the other hand, our recent work showed that at least *flhA*, *flhF*, *fleN* and *fliA* are part of the same operon in *P. putida* (Navarrete *et al*., 2019). To test the validity of the transcriptional organization inferred from the RNA-seq data, we performed RT-PCR with oligonucleotides framing all intergenic regions larger than 3 bp from *flhA* to *flrD* and cDNA from *P. putida* KT2442, a rifampicin-resistant derivative of KT2440, as a template (**Fig. 2B**). Amplification was obtained from all of the intergenic regions tested, strongly suggesting that transcripts initiated from P*flhA* span all fourteen coding regions down to *flrD*. This assay was also performed with cDNA from MRB49, a KT2442 derivative bearing a polar miniTn*5*-Km insertion in *flhF* (Navarrete *et al*., 2019). Transcription levels were diminished for all intergenic regions downstream from the insertion down to *motD-parC*. Polarity was not observed at the *parP*-*cheW* and *cheW-flrD* junctions. The loss of the polar effect in this region and the increased transcription obtained in both the wild-type and mutant strain, suggest the presence of an internal promoter activity. We note that one or more FliA-dependent internal promoters may be present in this region and not be detected in this background, as polarity of the *flhF*::miniTn*5*-Km mutation renders this strain FliA^-^ (Navarrete *et al*., 2019).

Discrepancy with the transcriptional organization proposed for *P. aeruginosa* was also observed at the regions spanning from *hptB* to *fliL*, *fliE* to *fliJ*, *fliC* to *fleR* and *flgA* to *flgZ*. RT-PCR assays using KT2442 cDNA as a template and primers spanning the conflicting intergenic regions confirmed co-transcription of *hptB*, *fliK* and *fliL* (**Fig.2D**) and *fliD*, *flaG*, *fliD*, *fliS*, *fliT*, *fleQ* and *fleS* (**Fig. 2F**), and transcriptional termination at the *fliG-fliH* intergenic region (**Fig. 2D**), consistent with the RNA-seq results. A similar RT-PCR analysis of the *flgAMNZ* operon was previously published (Wirebrand *et al*., 2018). Altogether, the RNA-seq and RT-PCR data and our previously published results (Navarrete *et al*., 2019; Wirebrand *et al*., 2018) strongly suggest that the flagellar genes of the *P. putida* KT2440 flagellar cluster are transcribed as ten separate operons. In addition, at least two of the operons contain internal promoters. Finally, an eleventh flagellar operon not linked to the cluster spans the genes *motA* and *motB*.

### *P. putida* flagellar promoters revealed by sequence alignment

Given the high conservation of the flagellar cluster organization across the genus *Pseudomonas*, we hypothesized that such conservation may also extend to the promoter regions preceding each of the operons. To test this hypothesis, we performed sequence alignments of the intergenic regions upstream from *fliK2*, *flhA*, *hsbA*, *fliH*, *fliE*, *fliC*, *flgF*, *flgB*, *cheV*, *flgA* and *motA*, all of which are expected to bear the primary promoters for the operons defined above. For this purpose, we used the genomic sequences of *P. putida* strains KT2440 and F1, *P. syringae* pv tomato strain DC3000, *P. syringae* pv syringae strain B728a, *P. savastanoi* pv phaseolicola strain 1448a, *P. fluorescens* strain SBW25 and *Pseudomonas protegens* strain Pf5 (**Fig. 3, Supplementary Fig. S2; Supplementary File S2**). While the general conservation of the intergenic sequences was modest, alignments revealed conserved motifs showing high similarity to the *σ*^54^-dependent promoter consensus (Barrios *et al*., 1999) at the intergenic regions upstream from *flhA*, *fliE*, *flgF*, *flgB* and *flgA*. We also found conserved sequences bearing similar to the FliA-dependent promoter consensus sequence (Rodríguez-Herva *et al*., 2010) upstream from *fliK2*, *fliC* and *cheV*, the latter located within the divergent *flgA* coding sequence. A motif resembling the *σ*^70^-dependent promoter consensus featuring an extended -10 and a poor -35 box (Bown *et al*., 1997) was present upstream from *hsbA*. We observed that the intergenic region upstream from *fliH* was too short to accommodate a promoter (<15 bp) in the *P. fluorescens*, *P. protegens*, *P. syringae* and *P. savastanoi* strains used in the alignment. A survey of *P. putida* strains revealed generally longer intergenic regions containing two copies of a repetitive extragenic palindromic sequence previously documented (Aranda-Olmedo *et al*., 2002), but no obvious similarity to known promoter consensus motifs. Yet, as our RNA-seq and RT-PCR results predict that a promoter must be active upstream from *fliH*, we included this region in the analyses below.

**Figure 3.**
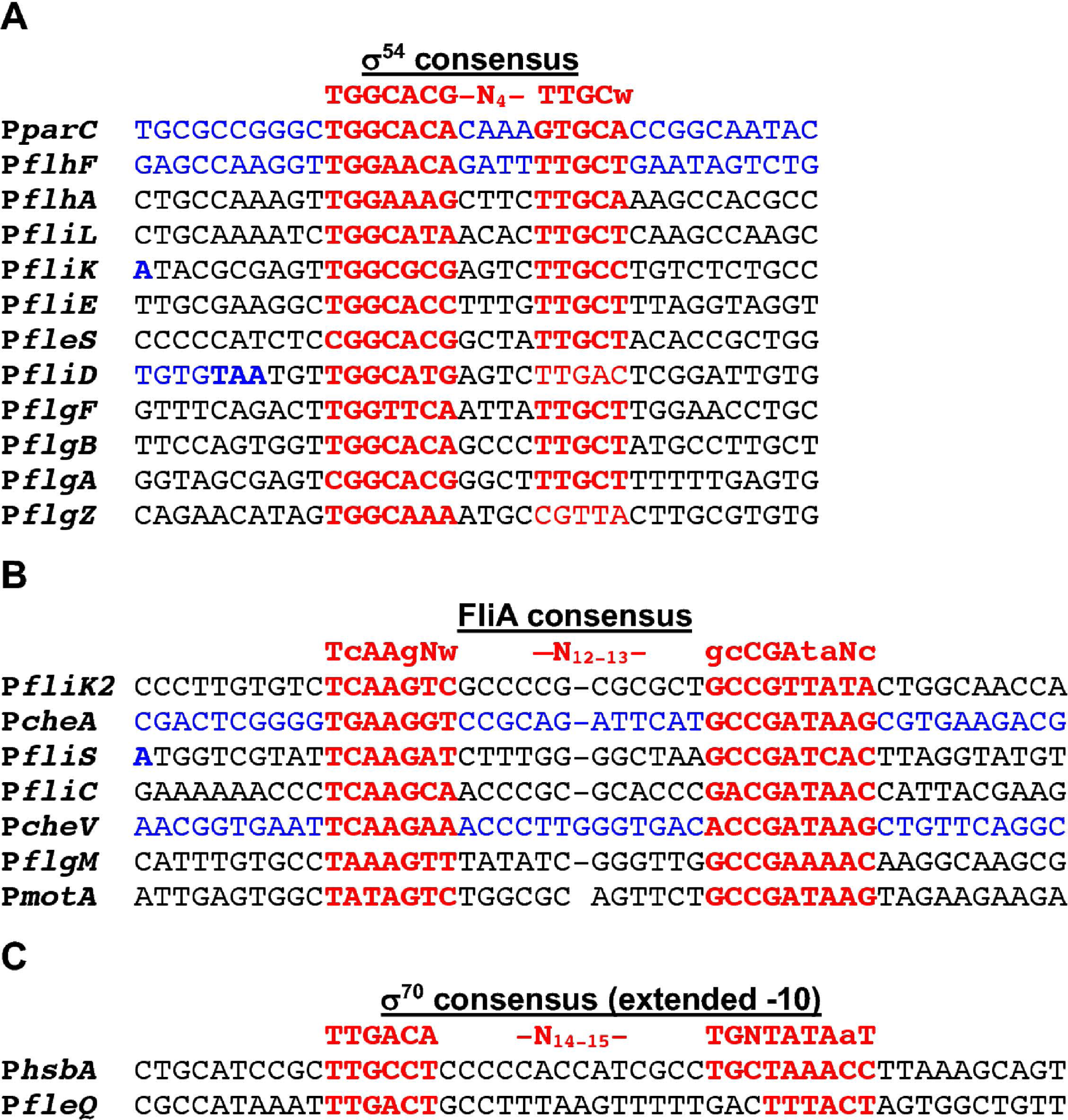
The *P. putida* KT2440 flagellar promoter set. Sequences of the twenty-one putative *P. putida* KT2440 flagellar promoters identified in this work, along with the consensus sequences for the corresponding *σ* factors: *σ*^54^ (**A**), FliA (**B**) and *σ*^70^ (**C**). The promoter motifs are indicated in red, and the coding sequences of the upstream genes are indicated in blue.

Because we have previously shown evidence of the presence of additional promoter activities upstream of *flhF*, *fliS* and *flgM* (Jiménez-Fernández *et al*., 2016; Navarrete *et al*., 2019; Wirebrand *et al*., 2018), we also assessed the presence of putative promoter motifs by means of sequence alignment in multiple additional intergenic regions of the flagellar cluster (**Fig. 3, Supplementary Fig. S2; Supplementary File S2**). Additional matches to the *σ*^54^-dependent promoter consensus were identified at the intergenic regions upstream from *fliL*, *fliK* and *fleS*, while sequences showing conservation at the -24 but not the -12 motif were found upstream from *fliD and flgZ*. We also identified conserved sequences matching the FliA-dependent promoter consensus upstream from *fliS* and *flgM*, and a putative *σ*^70^-dependent promoter motif upstream from *fleQ*. Finally, putative internal promoters located within the coding sequences of the preceding genes were also found upstream from *parC* and *flhF* (*σ*^54^-dependent), and *cheA* (FliA-dependent). In summary, this analysis revealed twenty-one putative flagellar promoters. Twelve such regions bear matches to the *σ*^54^ consensus, seven show similarity with the FliA consensus, and two more harbor putative *σ*^70^ promoter motifs.

### Strategy for analysis of the regulation of the *P. putida* flagellar cluster promoters

In order to characterize the expression from the putative flagellar promoters identified, we expanded our pre-existing collection of *P. putida* promoter fusions (Jiménez-Fernández *et al*., 2016) to include all the potential promoter regions identified. The selected regions were cloned into the Gateway^®^ vectors pMRB2 or pMRB3 to generate transcriptional fusions to the reporters *gfp-*mut3 and *lacZ*, as previously described (Jiménez-Fernández *et al*., 2016). All twenty-two *gfp-lacZ* fusion plasmids and the empty vector control pMRB1 were transferred to selected *P. putida* strains by mating. Expression was subsequently monitored along the growth curve by means of A_600_ and GFP fluorescence measurements in a microtiter plate reader with temperature and shaking control capabilities. GFP fluorescence was plotted *vs* A_600_ and differential rates of GFP accumulation during exponential growth (d*fluorescence*/d*A_600_*) were calculated from the slope of the linear fit of the plotted data (Jacob and Monod, 1961) (**Supplementary Fig. S3**). The results of thiese experiments are summarized in **Supplemental File S3**.

To assess the effect of different regulatory factors on flagellar/chemotaxis promoter transcription we used a subtractive approach in which we compared activity of the gene fusion constructs in the wild-type strain with that in mutants bearing deletions of single regulatory genes in an otherwise wild-type background. To this end, we chose the wild-type *P. putida* strain KT2442 and its mutant derivatives MRB52, MRB149, MRB71 and MRB93, bearing in-frame deletions of the *fleQ*, *rpoN*, *fleN* and *fleR* regulatory genes, respectively. To assess whether the observed regulatory effects are direct or indirect, we used an additive strategy. To this end, we constructed MRB130, a *P. putida* KT2442 derivative lacking the flagellar cluster (Δflagella), including all the regulatory elements therein. In this background, we inserted miniTn*7* derivatives expressing regulatory elements ectopically to test their effect on the expression of the fusion constructs of the flagellar/chemotaxis predicted promoter regions in the absence of other components of the flagellar cascade. In this context, we interpreted restoration of expression in the presence of FleQ or FliA as evidence of direct interaction between each regulatory factor and the cognate promoter region.

### P*fleQ* and P*fliH* transcription is not regulated by FleQ, FleR or FliA

The differential rates of GFP accumulation from the P*fleQ-gfp-lacZ* and P*fliH-gfp-lacZ* fusions were low (4.06±0.30 and 5.45±0.26 arbitrary units, respectively) and indistinguishable from to the empty vector values (3.88±0.42 arbitrary units). As an alternative approach, expression was monitored by end-point *β*-galactosidase assays in exponential and stationary phase. Expression from both promoters was negligible in exponential phase, but the activity levels were significantly increased (4- to 5-fold) in stationary phase (**Fig. 4**). P*fliH* transcription was not significantly altered in strains lacking the transcription factors FleQ and FleR, or the auxiliary regulatory protein FleN. In the case of P*fleQ*, statistically significant changes in GFP accumulation were observed in the Δ*fleQ*, Δ*fleN* and Δ*fleR* mutants in stationary phase, but the extent of the regulation (1.2-fold or less) suggests that these differences may be due to minor changes is cell physiology rather than the result of a true regulatory process. Using the additive approach we noted that the differential rates derived from the P*fleQ* fusion were 2-fold lower in the Δflagella strain in stationary phase. However, the wild-type expression levels were not restored by ectopic production of FleQ or FliA (**Fig. 4A**). P*fliH* expression was not affected by the deletion of the complete flagellar cluster or the ectopic production of FliA (**Fig. 4B**). Taken together, these results suggest that the upstream regions of *fleQ* and *fliH* contain promoters that are poorly expressed in exponential phase, induced in stationary phase, and not substantially regulated by any of the known flagellar transcription factors.

**Figure 4.**
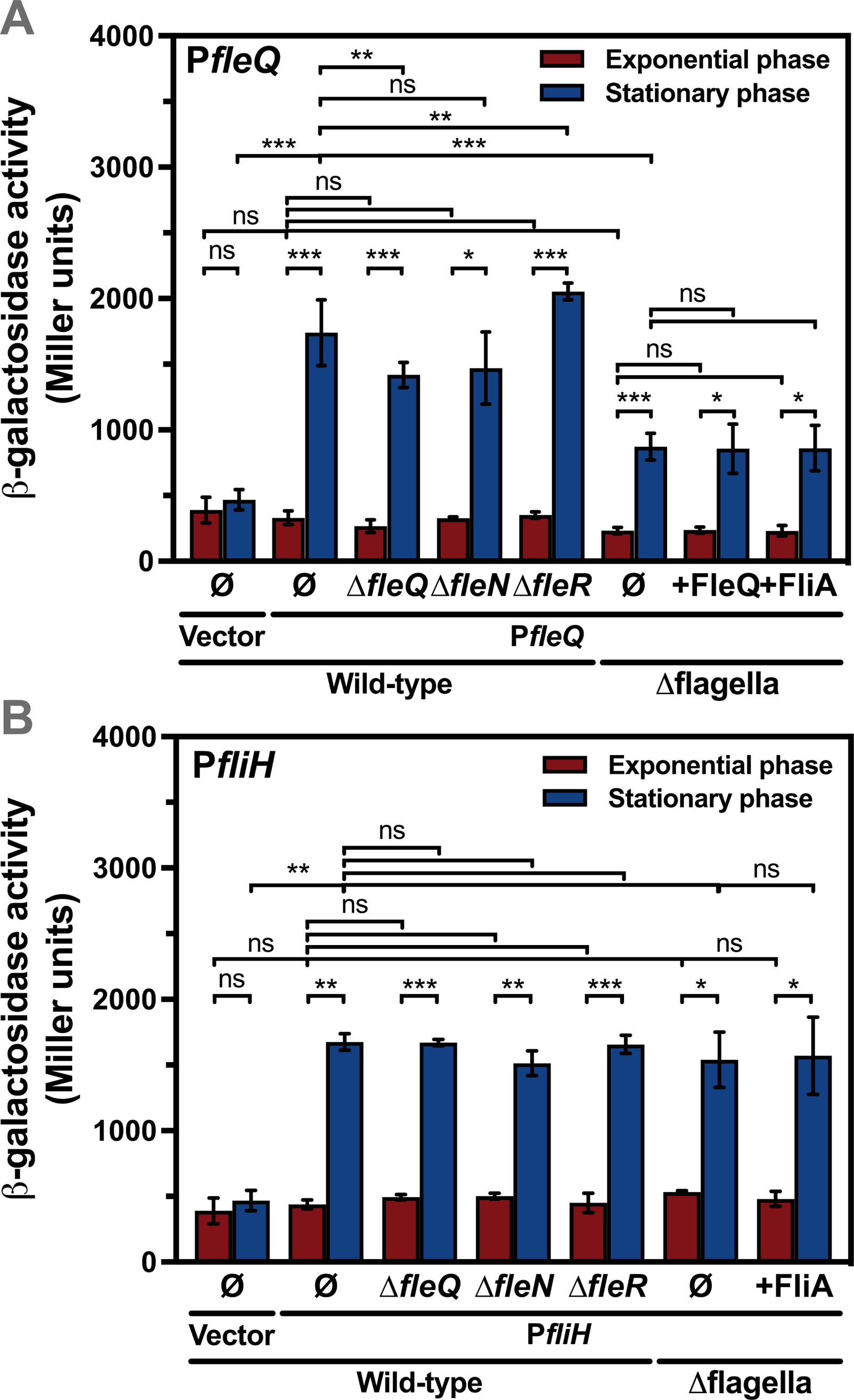
Expression of the P*fleQ* and P*fliH* promoters in *P. putida* KT2442 and mutant derivatives. *β*-galactosidase assays of the P*fleQ* (**B**) and P*fliH* (**B**) promoter fusions in the wild-type strain KT2442 and its Δ*fleQ*, Δ*fleN*, Δ*fleR*, Δflagella and Δflagella ectopically expressing FleQ or FliA derivatives. **Ø** denotes no further modification of the genetic background indicated below. Bars represent the averages and standard deviations of at least three independent assays. Stars designate p-values for the Student’s t-test for unpaired samples not assuming equal variance. ns: *p*≥0.05; *: p<0.05; **: p<0.01; ***:p<0.001. For each promoter fusion, the T-test was performed for (i) empty vector *vs.* promoter fusion in the wild-type strain; (ii) the wild-type *vs* the set of deletion mutants, (iii) the Δflagella mutant *vs.* its derivatives producing FleQ or FliA; and (iv) each fusion-bearing strain in exponential *vs.* stationary phase.

### Flagellar promoters directly regulated by FleQ

Analysis of the differential rates of GFP accumulation showed that all twenty additional flagellar promoters drove significant levels of transcription from the fusion constructs (**Figs. 5 and 6)**. Fifteen of these fusions were clearly downregulated (2- to 41-fold) in the Δ*fleQ* strain, suggesting the direct or indirect involvement of FleQ in their regulation. Statistically significant regulation below the 2-fold threshold was also observed for the weakly active P*fliK*, P*fliD* and P*flgZ* promoter regions, and for P*flhF* and P*hsbA*, which showed relatively high levels of FleQ-independent expression. Taken together, our results are fully consistent with the notion that FleQ is the master activator of the synthesis of the vast majority of flagellar and chemotaxis components.

**Figure 5.**
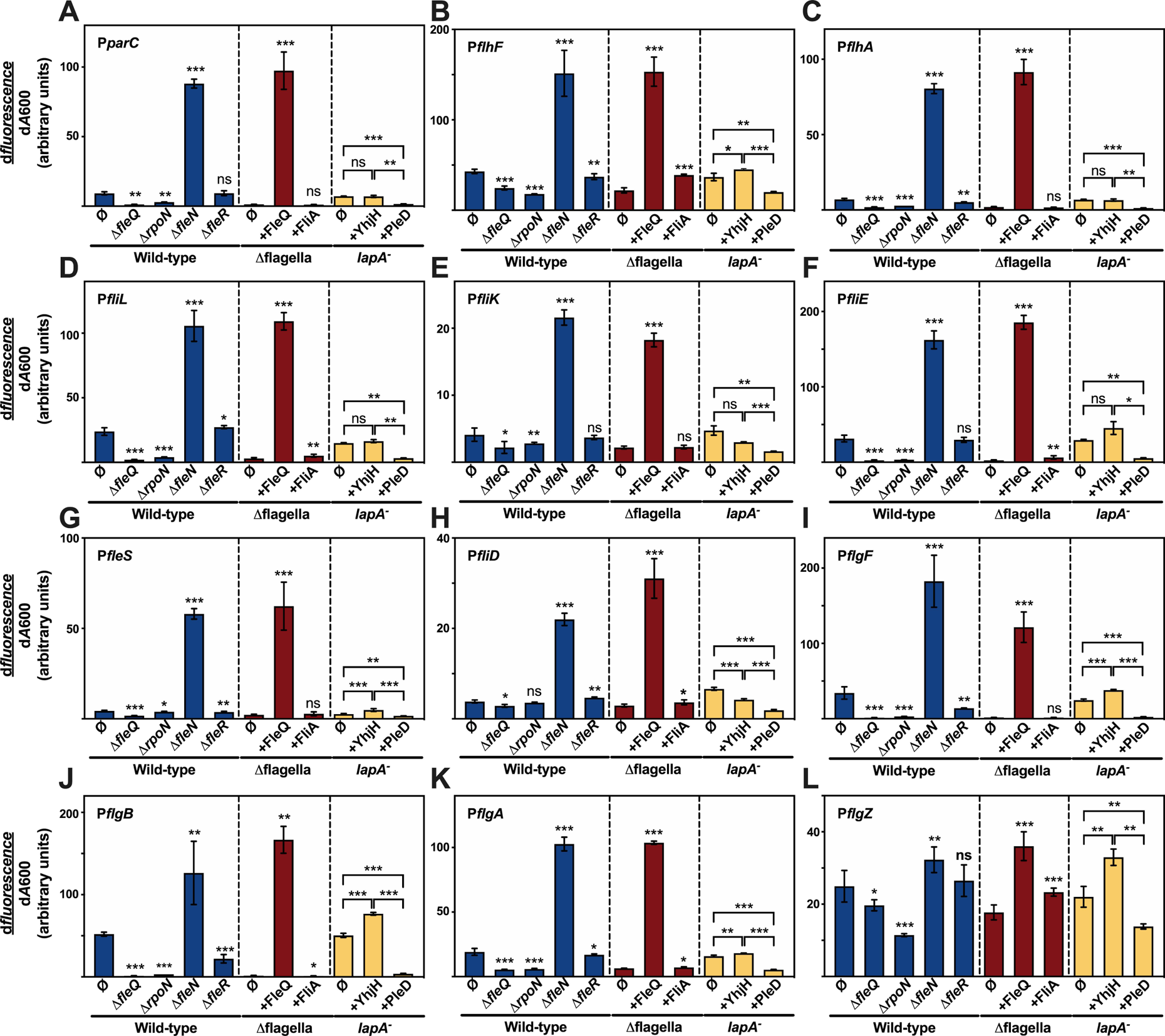
Expression the Class II flagellar promoters in *P. putida* KT2442 and mutant derivatives. Differential rates of GFP fluorescence accumulation of the P*parC* (**A**), P*flhF* (**B**), P*flhA* (**C**), P*fliL* (**D**), P*fliK*(**E**), P*fliE* (**F**), P*fleS* (**G**), P*fliD* (**H**), P*flgF* (**I**), P*flgB* (**J**), P*flgA* (**K**) and P*flgZ* (**L**) promoters in the wild-type strain KT2442 and its Δ*fleQ*, Δ*rpoN*, Δ*fleN*, Δ*fleR*, Δflagella, Δflagella ectopically expressing FleQ or FliA, and *lapA^-^* expressing YhjH or PleD derivatives. **Ø** denotes no further modification of the genetic background indicated below. Bars represent the averages and standard deviations of at least three independent assays. Stars designate p-values for the Student’s t-test for unpaired samples not assuming equal variance. Results are shown for the pairwise comparisons of the Δ*fleQ*, Δ*rpoN*, Δ*fleN* and Δ*fleR* mutants using the wild-type strain as the reference, the Δflagella strain expressing FleQ or FliA using the Δflagella strain as a reference, and all pairwise comparisons involving the *lapA-* mutant and its derivatives expressing YhjH or PleD. ns: *p*≥0.05; *: p<0.05; **: p<0.01; ***:p<0.001. The complete statistical analysis of this dataset is available in **Supplementary File S3**.

**Figure 6.**
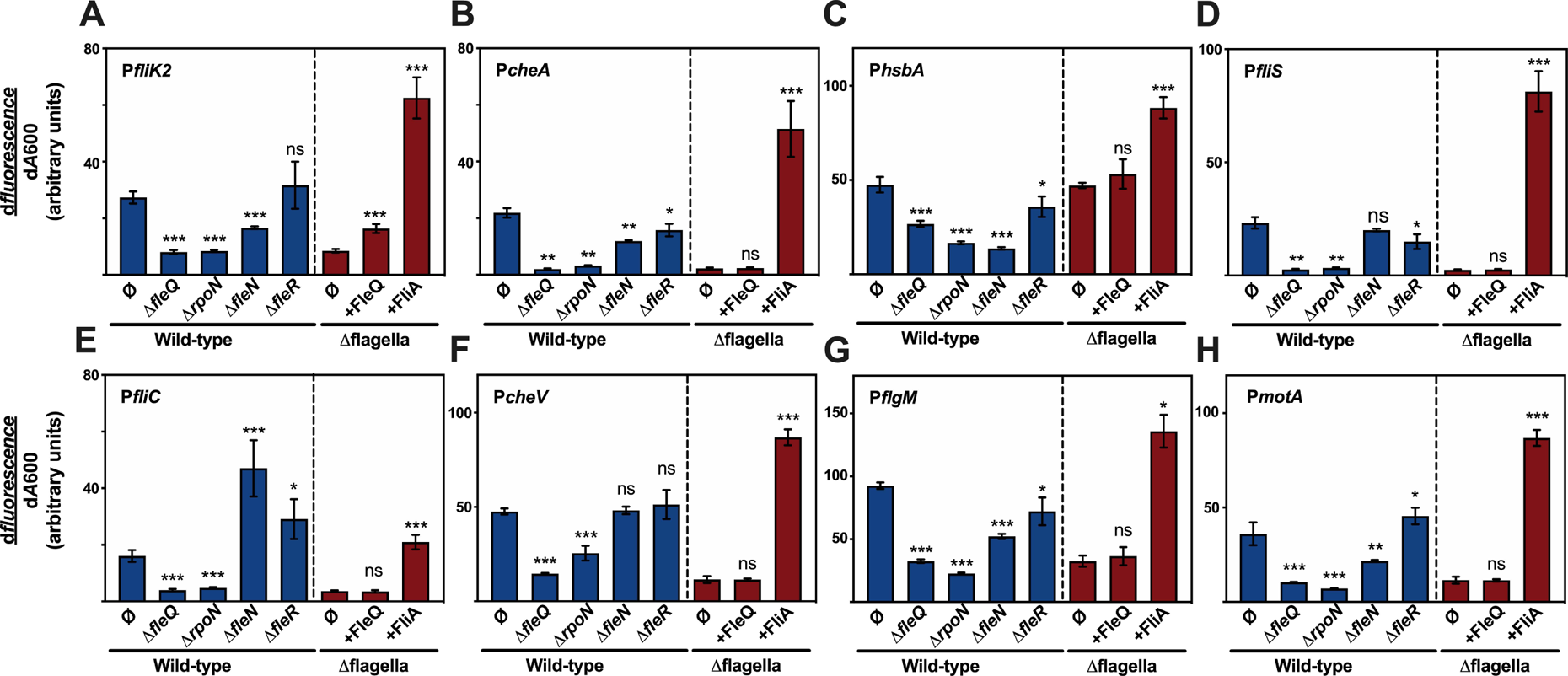
Expression the Class III flagellar promoters in *P. putida* KT2442 and mutant derivatives. Differential rates of GFP fluorescence accumulation of the P*fliK2* (**A**), P*cheA* (**B**), P*hsbA* (**C**), P*fliS* (**D**), P*fliC*(**E**), P*cheV* (**F**), P*flgM* (**G**) and P*motA* (**H**) promoters in the wild-type strain KT2442 and its Δ*fleQ*, Δ*rpoN*, Δ*fleN*, Δ*fleR*, Δflagella and Δflagella ectopically expressing FleQ or FliA. **Ø** denotes no further modification of the genetic background indicated below. Bars represent the averages and standard deviations of at least three independent assays. Stars designate p-values for the Student’s t-test for unpaired samples not assuming equal variance. Results are shown for the pairwise comparisons of the Δ*fleQ*, Δ*rpoN*, Δ*fleN* and Δ*fleR* mutants using the wild-type strain as the reference, and the Δflagella strain expressing FleQ or FliA using the Δflagella strain as a reference. ns: *p*≥0.05; *: p<0.05; **: p<0.01; ***:p<0.001. The complete statistical analysis of this dataset is available in **Supplementary File S3**.

To identify which flagellar promoters are directly regulated by FleQ, we tested the expression of the set of FleQ-activated promoter fusions in the Δflagella strain bearing miniTn*7-nahR-*P*sal-fleQ*, which produces FleQ ectopically from the salicylate-inducible P*sal* promoter (**Figs. 5 and 6**). Preliminary assays showed that salicylate induction greatly decreased the growth rate and was not used in these experiments. Nevertheless, *fleQ* transcription from uninduced P*sal* was still evident in RT-PCR assays (**Supplementary Fig. S4**), and ectopic production of FleQ in these conditions was sufficient to cause a significant increase in the differential expression rates (2- to 110-fold) obtained from the P*parC*, P*flhF*, P*flhA*, P*fliL*, P*fliK*, P*fliE*, P*fleS*, P*fliD*, P*flgF*, P*flgB*, P*flgA* and P*flgZ* promoter fusions. Significantly, the sequence alignment study of these twelve promoter regions revealed the presence of putative *σ*^54^-dependent promoter motifs (**Fig. 3, Supplementary Fig. S2; Supplementary File S2**). In addition, putative FleQ binding sites bearing similarity to the consensus sequences proposed for *P. aeruginosa* (Baraquet and Harwood, 2015) and *P. putida* (Molina-Henares *et al*., 2016) were found either upstream or downstream from the *σ*^54^-binding motifs (**Supplementary Fig. S5**). Still, it may be argued that FleQ regulation may conceivably be indirect in the Δflagella background, as recent ChIP-Seq results suggest that FleQ may bind DNA in the vicinity of some regulatory genes outside of the flagellar cluster (Blanco-Romero *et al*., 2018). We have previously shown that P*flhA*, P*flgA* and P*flgB* are still activated by FleQ when expressed in *E. coli*, an organism that naturally lacks FleQ and uses a completely different strategy to regulate flagellar gene expression (Jiménez-Fernández *et al*., 2016). To clarify this issue, we expanded these results by assessing expression of the twelve promoter fusion set in *E. coli* ET8000 producing *P. putida* FleQ from its own promoter region at the multicopy plasmid pMRB99 (**Supplementary Table S2**). Ten of the twelve promoters were significantly activated in a FleQ-dependent fashion in this background, with induction ratios ranging from 4- to 81-fold. In contrast, P*fleS* and P*flgZ* expression was not significantly altered by the presence of FleQ. These results suggest that FleQ interacts directly at least with the P*parC*, P*flhF*, P*flhA*, P*fliL*, P*fliK*, P*fliE*, P*fliD*, P*flgF*, P*flgB* and P*flgA* promoter regions to activate transcription, while the direct involvement of FleQ in activation of the P*fleS* and the P*flgZ* is at this point unclear (also see below).

Expression from the vast majority fusions in this set showed significantly reduced expression in a Δ*rpoN* background lacking *σ*^54^ (**Fig. 5**). However, the induction ratio was low (less than 2-fold) for P*fliK*, P*fleS* and P*fliD* promoters, and the difference was not statistically significant for P*fliD.* To investigate the involvement of *σ*^54^ further and explore a possible effect of the growth phase on *σ*^54^ regulation of the flagellar promoters, we also assayed expression by means of end-point *β*-galactosidase assays in wild-type and Δ*rpoN P. putida* strains grown to exponential and stationary phase (**Supplementary Table S3**). Statistically significant differences in the measured activity with induction ratios of 2-fold or more were observed with all promoters in stationary phase and all promoters but P*fleS* and P*flgZ* in exponential phase, suggesting that *σ*^54^ contributes to the expression of all twelve promoters and its effect on P*fleS* and P*flgZ* is more prominent in stationary phase.

Expression from the twelve promoter fusion set was also measured in a KT2442 derivative lacking the auxiliary regulatory protein FleN (**Fig. 5**). The differencial fluorescence accumulation rates were clearly increased for eleven of the twelve fusions (2- to 13-fold) in the Δ*fleN* background. Exceptionally, a small (less than 2-fold) but statistically significant increase was observed for the weak P*flgZ* promoter. These results support the notion that FleN antagonizes FleQ activation of the flagellar promoters (Dasgupta and Ramphal, 2001). The second messenger c-di-GMP has been shown to positively and negatively modulate FleQ activity (Baraquet *et al*., 2012; Claudine and Harwood, 2013; Hickman and Harwood, 2008). We also tested expression of the twelve fusions in *P. putida* strains that produce the *Caulobacter crescentus* diguanylate cyclase PleD* or the *E. coli* phosphodiesterase YhjH ectopically, thus increasing or decreasing the intracellular c-di-GMP concentrations. To this end, we chose a KT2442 derivative lacking the high molecular weight adhesin LapA, as c- di-GMP overproduction provokes aggregation of LapA*^+^ P. putida* (Jiménez-Fernández *et al*., 2016) that precludes accurate absorbance and fluorescence measurements. The differential rates of fluorescence accumulation in the unmodified *lapA^-^* strain were generally similar to the strain with artificially decreased c-di-GMP levels, suggesting that c-di-GMP levels are generally low in our experimental conditions. Comparison of the *P. putida* strains bearing modifed c-di-GMP levels showed statistically significant c-di-GMP-dependent downregulation of the twelve fusions, with inhibition ratios of 2- to 21-fold, except for a somewhat smaller (1.8-fold) ratio obtained with the P*fliK* fusion (**Fig. 5**). Taken together, our results indicate that the P*parC*, P*flhF*, *flhA*, P*fliL*, P*fliK*, P*fliE*, P*fliD*, P*flgF*, P*flgB* and P*flgA* promoters are dependent on the alternative *σ* factor *σ*^54^, positively regulated by FleQ, negatively regulated by FleN and inhibited by high c-di-GMP levels. Although we have not been able to demonstrate the direct regulation of P*fleS* by FleQ beyond any doubt, the weight of the bioinformatic and experimental evidence and the coincidence of its behavior with the rest of the set in every other regard strongly suggests that P*fleS* is likely a *bona fide* FleQ-dependent promoter. On the other hand, P*flgZ*, a poorly expressed promoter, exhibited a similar response to FleQ, FleN, *σ*^54^ and c-di-GMP, but the extent of the regulation was modest in all cases.

### FleR is not essential for flagellar function and flagellar gene expression

The FleSR two-component system is responsible for the activation of Class III flagellar genes in *P. aeruginosa* and other bacteria (Dasgupta *et al*., 2003; Echazarreta and Klose, 2019), but is not characterized in *P. putida*. To explore a possible function of FleSR in flagellar motility and the regulation of the flagellar cluster, we constructed MRB92 and MRB93, two KT2442 derivatives bearing in-frame deletions of *fleS* and *fleR*, respectively. Swimming assays were performed with the Δ*fleR* and Δ*fleS* mutant using the wild-type and Δ*fleQ* strains as positive and negative controls, respectively. The sizes of the swimming halos produced by the wild-type, Δ*fleR* and Δ*fleS* strains were indistinguishable, while no motility halo was obtained with the Δ*fleQ* mutant (**Supplementary Fig. S6**). These results indicate that FleR and FleS are not essential for flagellar motility, at least under our experimental conditions.

To determine the contribution of FleR to the regulation of the flagellar promoters, expression from the complete fusion set was also examined in the Δ*fleR* mutant (**Figs. 5 and 6**). Deletion of *fleR* had a statistically significant effect on the differential rates of fluorescence accumulaion by the P*cheA*, P*flhF*, P*flhA*, P*fliL*, P*hsbA*, P*fleS*, P*fliS*, P*fliD*, P*fliC*, P*flgF*, P*flgB*, P*flgA*, P*flgM* and P*motA* promoters. However, the fold changes were modest in all cases, except for 2-fold downregulation of the FleQ-dependent P*flgF* and P*flgB* and 2-fold upregulation of P*fliC*. These results suggest that FleR does not have a major role in the regulation of flagellar gene expression, but may contribute to fine-tune the expression of some flagellar operons. Alternatively, FleR function may involve a signaling event that does not occur under our experimental conditions.

### Flagellar promoters directly regulated by FliA

The additive approach was also used to assess which promoters of the flagellar cluster are directly regulated by the flagellar *σ* factor FliA. To this end, differential rates of fluorescence accumulation were assessed for the complete set of FleQ-regulated flagellar promoters in the Δflagella strain bearing miniTn*7-nahR-*P*sal-fliA*, which produces FliA ectopically from the salicylate-inducible P*sal* promoter. As with the FleQ-producing construct above, salicylate induction was detrimental to growth, but in the absence of salicylate, ectopic production of FliA significantly stimulated expression (4- to 33-fold) from all seven promoter regions (namely P*fliK2*, P*cheA*, P*fliS*, P*fliC*, P*cheV*, P*flgM* and P*motA*) bearing matches to the FliA consensus sequence (**Fig. 6**). In contrast, ectopic production of FleQ failed to significantly alter the expression from any of these fusions, except for a modest (∼2-fold) increase in P*fliK2* (**Fig. 6**). Conversely, ectopic production of FliA had no significant effect on the differential rates of fluorescence accumulation by the FleQ-dependent P*flhA*, P*fliK*, P*fleS*, and P*flgF*, while statistically significant but modest (≤2-fold) stimulation was observed with P*parC*, P*flhF*, P*fliL*, P*fliE*, P*fliD*, P*flgB*, P*flgA* and P*flgZ* (**Fig. 5**). Although most FliA-dependent fusions were downregulated in the Δ*fleQ*, Δ*rpoN* and Δ*fleN* backgrounds, we note that P*fliC* expression was increased in the absence of FleN (**Fig. 6**). Taken together, our results suggest that the promoters contained at the P*fliK2*, P*cheA*, P*fliS*, P*fliC*, P*cheV*, P*flgM* and *motA* promoter regions are dependent on the alternative *σ* factor FliA, and indirectly regulated by *σ*^54^ and FleQ. An additional promoter region not carrying an evident match to the FliA consensus, P*hsbA*, showed a similar response pattern to FliA, FleQ and *σ*^54^, but the extent of the regulation was moderate in all cases and significant basal transcription was observed in all backgrounds. We propose that this promoter region may contain a weak, previously undetected FliA-dependent promoter and an additional FliA-independent promoter activity.

## DISCUSSION

The *P. putida* KT2440 flagellar cluster resides in a ∼70 kbp chromosomal region encoding the vast majority of structural and regulatory flagellar proteins along with the core components of the chemotaxis machinery. Here we expand on our previous work (Jiménez-Fernández *et al*., 2016; Navarrete *et al*., 2019; Wirebrand *et al*., 2018) to identify the transcriptional organization of the *P. putida* KT2440 flagellar cluster and characterize the transcriptional regulation of twenty primary and internal promoters accounting for the expression of all but three relevant. The results presented provide (i) updated annotations and putative functions for seven flagellar genes; (ii) evidence of the transcriptional organization of the flagellar cluster in ten separate operons, while an additional operon, *motAB*, is not linked to the cluster; (iii) an inventory of twenty-two flagellar promoters, including eleven primary and eleven internal promoters; (iv) characterization of the regulation of all twenty-two flagellar promoters regarding their dependence on known flagellar transcription factors (FleQ, FleR, FliA, *σ*^54^), the auxiliary protein FleN and the intracellular signalling molecule c-di-GMP; (v) a novel hierarchy for the transcriptional cascade controlling flagellar gene expression that differs significantly from those previously described; (vi) evidence of high conservation of the flagellar gene complement and organization in the genus *Pseudomonas*; and (vii) evidence of high conservation of the flagellar promoter motifs in a set of environmental *Pseudomonas*, suggesting that many of the results presented here may be applicable to this group of organisms.

We have defined the *P. putida* KT2440 flagellar cluster as the chromosomal region encompassing ORFs PP_4328 (*fliK2*) to PP_4397 (*flgZ*). These boundaries were established based on three lines of evidence. (i) High conservation: 51 of the flagellar genes are present in all the *Pseudomonas* genomes analyzed; the remaining 8 are found in at least 80%; (ii) conservation of gene arrangement: all flagellar genes are clustered in the same order in one to three chromosomal locations in most *Pseudomonas* genomes analyzed (**Supplementary File 1**); and (iii) experimental evidence of involvement in flagellar function: all but one of the 59 flagellar genes in the cluster have been demonstrated to play roles related to assembly and operation of the flagellar and chemotaxis complexes in at least one organism. In addition to the genes previously annotated, we have been able to identify an ortholog to *H. pylori flhX*, encoding a small protein similar to the processed form of FlhB that determines the specificity switch of the FT3SS (Pallen *et al*., 2005). FlhX has been shown to partially complement FlhB function in *H. pylori* (Smith *et al*., 2009), but its actual role in flagellar assembly is not well understood. One more gene in the cluster is an ortholog of *V. cholerae flrD*, encoding a transmembrane signal transduction protein that stimulates transcriptional activation by the FlrBC two-component system (equivalent to FleSR) (Moisi *et al*., 2009). We have also identified orthologs of *Vibrio* spp. *parP* and *parC*, encoding two polar landmark proteins required for polar positioning of the chemotaxis arrays (Ringgaard *et al*., 2018). Finally, we have annotated the orthologs of *P. aeruginosa hptB*, *hsbR* and *hsbA*, encoding a phosphotransferase a response regulator and an anti-*σ* factor antagonist proteins. This system antagonizes FlgM sequestration of FliA by partner-switching mechanism (Bhuwan *et al*., 2012) and indirectly regulates biofilm formation by regulating the activity of the diguanylate cyclase HsbD (Valentini *et al*., 2016). In contrast, the function of *fliK2* is still uncharacterized. However, because *fliK2* is part of an operon with *flhX*, and its transcription is FliA-dependent in *P. putida*, we find it likely that it will eventually demonstrate to be a *bona fide* flagellar gene. With the updated annotation, we conclude that the conserved *Pseudomonas* flagellar cluster gene set encodes all the structural proteins of the flagellum (minus an additional flagellar motor encoded by the *motAB* operon elsewhere), all the elements required for flagellar assembly (including the complete FT3SS), all the core components of the chemotaxis apparatus, three dedicated transcription factors, seven auxiliary proteins to control their activity, three landmark proteins for the polar location of the flagellar and chemotaxis complexes, and one protein involved in functional regulation of flagellar rotation (**Supplementary Table 1**). We note that, in addition to the conserved gene complement described here, some organisms bear species- or strain-specific flagellar genes in their clusters [e.g., genes for flagellin glycosylation (Dasgupta *et al*., 2003)].

The *P. putida* KT2440 flagellar cluster is organized in ten transcriptional units, with primary promoters located upstream from *fliK2*, *flhA*, *hsbA*, *fliH*, *fliE*, *fliC*, *flgF*, *flgB*, *cheV* and *flgA*. In addition, internal promoters are found upstream from *flhF*, *cheA* and *parC* at the *flhAF-fleN-fliA-cheYZAB-motCD-parCP-cheW-flrD* operon, upstream from *fliK* and *fliL* at the *hsbAR-hptB-fliKLMNOPQE-flhB* operon, upstream from *fliD*, *fliS*, *fleQ* and *fleS* at the *fliC-flaG-fliDST-fleQRS* operon and upstream from *flgM* and *flgZ* at the *flgAMNZ* operon (**Fig. 2**). Some of the promoters shown here were identified and partially characterized in our previous work (Jiménez-Fernández *et al*., 2016; Navarrete *et al*., 2019; Wirebrand *et al*., 2018). Our results indicate that much of the *P. putida* KT2440 flagellar cluster is transcribed as part of long operons containing internal promoters. The recent development of high throughput RNA-sequencing techniques with the possibility of genome-wide mapping of transcriptional start and termination sites has revealed that nested operon architectures are much more common in bacterial genomes than expected (Conway *et al*., 2014; Vera *et al*., 2020; Warrier *et al*., 2018; Yan *et al*., 2018).

While the operon arrangement described here is generally reminiscent to that proposed for *P. aeruginosa* (Dasgupta *et al*., 2003), the *P. aeruginosa* cluster does not contain any operons with a nested architecture. Rather, the genes are grouped in shorter transcriptional units. We observe that the *P. aeruginosa* counterparts to the internal promoters shown here are regarded as primary promoters, which leads to the division of these operons into shorter transcriptional units. Whether this is the result of interspecies differences within *Pseudomonas* or a consequence of the different approaches used to characterize the transcriptional organization of these genes remains to be determined. It is noteworthy that long, nested flagellar operons have been described in other polarly-flagellated Gamma-proteobacteria (Y. K. Kim and McCarter, 2000; Prouty *et al*., 2001; Wilhelms *et al*., 2011; Wu *et al*., 2011).

The regulatory architecture of the flagellar transcriptional cascade has been elucidated in the polarly flagellated Gamma-proteobacteria *V. cholerae* (Prouty *et al*., 2001), *Vibrio parahemolyticus* (McCarter, 2001), *Vibrio campbelli* (Petersen *et al*., 2021), *Legionella pneumophila* (Albert-Weissenberger *et al*., 2010), *Aeromonas hydrophila* (Wilhelms *et al*., 2011), *Shewanella oneidensis* (Shi *et al*., 2014), *P. aeruginosa* (Dasgupta *et al*., 2003) and now *P. putida* (**Fig. 7**). Comparative analysis of the models proposed in these organisms suggests a number of common themes along with significant organism-to-organism variability even within the same genus. All the transcriptional cascades described are three- or four-tiered, with the transcription factor FleQ (also known as FlrA) at the top of the regulatory hierarchy. Accordingly, *fleQ/flrA* is invariably a Class I gene (i.e., its transcription does not require any of the dedicated flagellar transcription factors). Synthesis of FleQ/FlrA is likely activated in response to environmental and physiological cues that favor a motile lifestyle, but the conditions and regulatory mechanisms involved are generally not well understood. Regulation of *fleQ* expression in diverse *Pseudomonas* species may involve repression by the CRP ortholog Vfr (Dasgupta *et al*., 2002), AmrZ (Tart *et al*., 2006), OsaR (Ma *et al*., 2021) and activation by the AlgU-regulated antisense RNA *fleQas* (Markel *et al*., 2018). We have found that P*fleQ* is induced in stationary phase and not regulated by FleQ, FleR or FliA, suggesting that it belongs to Class I (**Fig. 4A**). Since transcription from P*fleQ* is not interrupted at the intergenic region between *fleQ* and *fleS* (**Fig. 2E and F**), *fleQ*, *fleS* and *fleR* must be considered Class I flagellar genes in *P. putida*, even though *fleS* and *fleR* are also transcribed from their own FleQ-dependent promoter. Although we have not explored regulation of the P*fleQ* promoter further, we note that the AlgU-dependent promoter motifs for *fleQas* are conserved in a wide range of *Pseudomonas* (Markel *et al*., 2018), and we observe significant levels of antisense transcription around the 5’ end of *fleQ* coding sequence (**Fig. 2E**), suggesting that *fleQas* may also be produced and operate in *P. putida*. A second promoter, P*fliH* driving the synthesis of the secretion ATPase components FliH,

**Figure 7.**
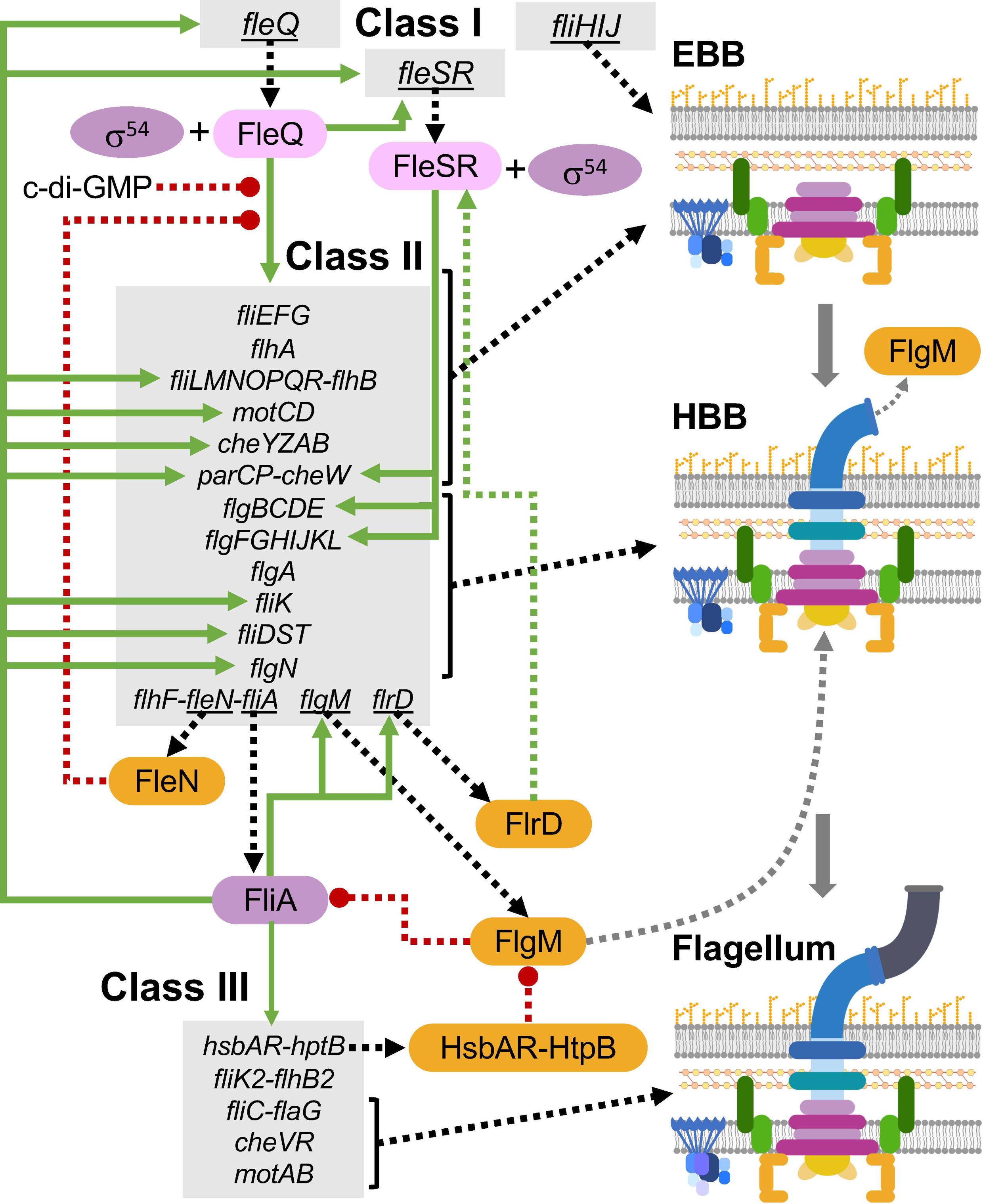
Hypothetical model of the flagellar transcriptional cascade in *P. putida* KT2440. Cartoon representing the transcriptional hierarchy, feed-forward and feedback loops of the *P. putida* KT2440 flagellar cascade. Genes are shaded in grey, transcription factors are shaded in pink, *σ* factors are shaded in purple. and auxiliary regulatory proteins are shaded in orange. Transcription is shown as green arrows and translation is shown as black dashed arrows. Green dashed arrows and red dashed lines with a round end denote positive and regulatory events based on protein-protein interaction, respectively. FlgM secretion by way of the FT3SS is denoted by a dashed grey arrow. Three stages of flagellar assembly are represented: early basal body (EBB), the hook-basal body complex (HBB), and the complete flagellum.

FliI and FliJ, showed a similar behavior to P*fleQ*: its expression was low in exponential phase, induced in stationary phase and unresponsive to any of the dedicated flagellar transcriptions factors (**Fig. 4B**). Accordingly, we also classify FliH, FliI and FliJ as Class I genes. We note that the presence of additional Class I genes other than the master regulation is not uncommon in polarly flagellated Gamma-proteobacteria (Dasgupta *et al*., 2003; Petersen *et al*., 2021; Wilhelms *et al*., 2011), and *flrB* and *flrC* are actually Class I genes in *V. campbellii* (Petersen *et al*., 2021).

FleQ activation of Class II genes sets in motion the assembly of the basal body (**Fig. 7**). Polar localization of the flagella is directed by the signal recognition particle-like GTPase FlhF (Schuhmacher *et al*., 2015). FlhF is likely the first flagellar gene product to locate to the cell pole and has been shown to recruit the first assembly building block, the MS-ring structural protein FliF, in *V. cholerae* (Green *et al*., 2009). As basal body assembly progresses, the MinD-like ATPase FleN (also known as FlhG) provides a negative regulatory loop by inhibiting the ATPase activity of FleQ, required for *σ*^54^ promoter activation (**Fig. 7**) (Baraquet and Harwood, 2013). Deletion of *fleN* provokes hyperflagellation and *fleN* overexpression prevents the synthesis of flagella (Schuhmacher, *et al*., 2015), suggesting that it may become active in response to an checkpoint that indicates that assembly of the correct number of flagella has been initiated. Recently, Blagotinsek *et al*., (2020) showed that FlhG is recruited to the flagellar pole as an inactive monomer by the C-ring protein FliM in *S. oneidensis*. Membrane-bound FlhG may then undergo ATP-induced dimerization to become active in a process that is stimulated by interaction with membrane lipids (Schuhmacher *et al*., 2015). The ATP-bound dimer is the form that interacts with FleQ to prevent Class II promoter activation (Baraquet and Harwood, 2013). Thus, assembly of the C-ring(s) may be the signal that triggers FleN-dependent inhibition of Class II promoter transcription. The extent of flagellar assembly enabled by FleQ-activated transcription is most often restricted to all or part of the early basal body (EBB) structures, namely the MS-ring, the C-ring and the FT3SS (Dasgupta *et al*., 2003; Khan *et al*., 2020). In contrast, we have shown that *P. putida* Class II genes encode all structural flagellar components except for the filament. This arrangement is reminiscent of that proposed for *V. campbellii* (Petersen *et al*., 2021) and *S. oneidensis* (Shi *et al*., 2014).

An additional tier, Class III, activated by the FleSR (also known as FlrBC) two-component system is required in *P. aeruginosa*, *V. cholerae*, *V. parahemolyticus* and *A. hydrolitica* to complete the assembly of the components of the hook-basal body (HBB) complex (Dasgupta *et al*., 2003; Khan *et al*., 2020; Wilhelms *et al*., 2011). Activation of FleSR/FlrBC was recently shown to respond to the assembly of the EBB in *V. cholerae* and *P. aeruginosa* (Burnham *et al*., 2020), as previously observed with the Epsilon-proteobacteria *Campylobacter jejuni* and *Helicobacter pylori* (Boll and Hendrixson, 2013; Tsang and Hoover, 2015). Sensing of the assembly status may be mediated by FlrD, a membrane protein containing a HAMP domain that stimulates FlrBC-dependent activation in *V. cholerae* (Moisi *et al*., 2009). *P. putida* encodes orthologs of FlrD and FleSR (**Supplementary Table S1**). However, because transcription of all the HBB genes is directly dependent on FleQ, FleR is not essential for the assembly of functional flagella (**Supplementary Fig. S6**). Yet, we have shown that FleR shows statistically significant effects on the transcription of multiple flagellar promoters. Particularly, transcription from the P*flgB* and P*flgF* promoters, driving the synthesis of the structural rod, LP ring and hook proteins, is somewhat diminished in the Δ*fleR* strain (**Fig. 5**), and sequence alignment revealed multiple conserved sequences with high similarity to the proposed FlrC binding motif (CGGCAA) in our set of seven environmental *Pseudomonas* strains in both promoter regions (**Supplementary File 2**). These observations suggest that FleR may play an unrecognized role in the synthesis of the late basal body and hook components. Interestingly, replacement of the *V. cholerae* P*flgB*, P*flgF* and P*flgK* FlrC-dependent promoters for the FlrA-dependent P*fliE* resulted in normal flagellation and motility in the wild-type strain, but provoked the loss of the flagellum and motility in a Δ*flhG* background, suggesting that the FlrBC may provide robustness to the regulatory circuit in the event of faulty FlhG function (Burnham *et al*., 2020). Accordingly, we hypothesize that FleSR activation of the P*flgB* and P*flgF* promoters in *P. putida* may act as a safeguard against a shortage of rod and hook subunits if FleQ activation is compromised by altered FleN signaling. While this evidently does not happen in our experimental setting, there may be conditions in which such backup activation by FleR may relevant. Overlapping functions of FleQ and FleR have been documented in a few other systems. FlrC, the FleR ortholog of *S. oneidensis*, is not required for motility and plays a minor role in the regulation of the flagellar promoters (Shi *et al*., 2014). In this organism ectopic production of FlrC can suppress the nonmotile phenotype of a Δ*flrA* mutant (lacking the cognate FleQ ortholog) by replacing FlrA as the activator of the flagellar *σ*^54^-dependent promoters (Gao *et al*., 2018). In a similar vein, the functions of FlrA and FlrC are partially redundant in the flagellar cascade of *V. campbelli*; however, in this case, it is FlrA, and not FlrC, that is dispensable for flagellar motility (Petersen *et al*., 2021). Finally, in *Rhodobacter sphaeroides*, FleQ and the FleR ortholog FleT cooperate in the regulation of Class III flagellar promoters by forming a heterooligomeric protein (Poggio *et al*., 2005). These and other possibilities of interplay between the two *P. putida σ*^54^-dependent activators are worth exploring in the future.

The final tier in the flagellar cascades is triggered by the release of FliA from inactivation by the anti-*σ* factor FlgM. As first demonstrated in *Salmonella*, a shift in secretion specificity upon completion of the hook structure leads to FlgM secretion by way of the FT3SS (Karlinsey *et al*., 2000). This is generally considered to be a conserved mechanism to activate FliA-dependent transcription in an assembly checkpoint-dependent fashion, and has been directly demonstrated in *V. cholerae* (Correa *et al*., 2004). In addition, the *P. aeruginosa* anti-sigma factor antagonist HsbA has been show to prevent FlgM-dependent inactivation of FliA by a partner-switching mechanism in response to a phosphorylation cascade involving several membrane hybrid sensor kinases, the histidine phosphotransferase HtpB and the response regulator HsbR (Bhuwan *et al*., 2012), but the signals triggering this system are currently not known and its connection with flagellar assembly is therefore unclear. As the *P. putida* flagellar cluster encodes FlgM, HsbA, HsbR and HtpB orthologs (**Supplementary Table S1**), any of these mechanisms may be functional, but we have not explored them in this work. The set of FliA-dependent Class IV genes in polarly flagellated Gamma-proteobacteria normally includes flagellin and other filament components, along with chemotaxis and flagellar stator genes. In *P. putida* FliA is essential for expression of *fliC* and *flaG*, encoding flagellin and a regulator of filament length, the chemotaxis operon *cheVR*, the stator genes *motAB* and the uncharacterized *fliK2-flhX* operon (**Figs. 2, 5, 6 and 7**), and boosts transcription of the distal chemotaxis and stator genes of the Class II *flhAF-fleN-fliA-cheYZAB-motCD-parCP-cheW-flrD* operon. Accordingly, FliA is essential for assembly of the filament and completion of the chemotaxis signal transduction system and one of the stator complexes (**Fig. 7**). We note that, although the core chemotaxis and the *motCD* stator genes were previously assigned to Class IV in *P. aeruginosa* (Dasgupta *et al*., 2003), re-examination of the evidence revealed that none of the orthologs of the *flhAF-fleN-fliAcheYZAB-motCD-parCP-cheW-flrD* operon genes were significantly downregulated in a Δ*fliA* mutant, and should therefore be re-classified as Class II [(see Supplementary Tables 2-6 in Dasgupta *et al*. (2003)]

FliA also participates in feed-forward and feedback loops on several regulatory elements (**Fig. 7**). On the one hand, it boosts the synthesis of the anti-sigma FlgM (**Fig. 6**), an antagonist of FliA itself. Most intriguingly, FliA-dependent transcription of *fleQ* and *fleSR* as part of the *fliC-flaG-fliDST-fleQSR* operon (**Fig. 2**) provides positive feedback to the synthesis of the transcription factors involved in early flagellar gene expression (**Fig. 7**). Since FleQ function and Class II transcription may be transiently impaired by the formation of an inactive complex with FleN (Baraquet and Harwood, 2013), FleQ newly synthesized by way of readthrough transcription from the strong Class III P*fliC* promoter may titrate out FleN, restoring expression of the Class II genes. Unlike peritrichous flagellates, in which new flagella can be continuously added during cell elongation, polar flagellates such as *P. putida* must assemble their flagella in a discontinuous fashion, as a new pole is generated only once per cell cycle during cell division. New flagella must then be assembled at the new pole in time for their completion at the following cell division. In this scenario, the reactivation of early flagellar gene expression by this FliA-mediated positive feedback mechanism appears to be a suitable mechanism to enable *de novo* flagella synthesis at the new pole while assembly at the old pole is completed.

## EXPERIMENTAL PROCEDURES

### Bacterial strains and growth conditions

The bacterial strains used in this work are summarized in **Supplementary Table S4**. Liquid cultures of *E. coli* and *P. putida* strains were routinely grown in Luria-Bertani (LB) broth (Sambrook and Russell, 2000) at 37°C and 30 °C, respectively, with 180 rpm shaking. For fluorescence monitoring, 1/10 strength LB was used to minimize background fluorescence. For solid media, Bacto-Agar (Difco) was added to a final concentration of 18 g l ^−1^. Antibiotics and other additions were used, when required, at the following concentrations: ampicillin (100 mg l^−1^), carbenicillin (0.5 g l^−1^), kanamycin (25 mg l^−1^), rifampicin (10 mg l^−1^), chloramphenicol (15 mg l ^−1^), gentamycin (10 mg l^−1^), tetracycline (5 mg l^−1^), 5-bromo-4-chloro-3-indoyl-β-D-galactopyranoside (X-gal) (25 mg l^−1^) and sodium salicylate (from 0.1 mM to 2 mM) All reagents were purchased from Sigma Aldrich, except for X-Gal, which was purchased from Fermentas.

### Plasmid and strain construction

Plasmids and oligonucleotides used in this work are summarized in **Supplementary Table S4**. All DNA manipulations were performed following standard procedures (Sambrook and Russell, 2000). Restriction and modification enzymes were used according to the manufacturers’ instructions (Fermentas, Roche and NEB) When required, blunt ends were generated using the Klenow fragment or T4 DNA polymerase. *E. coli* DH5*α* was used as a host in cloning procedures. All cloning steps involving PCR were verified by commercial sequencing (Secugen or Stab Vida) Plasmid DNA was transferred to *E. coli* and *P. putida* strains by transformation (Inoue *et al*., 1990), triparental mating (Espinosa-Urgel *et al*., 2000) or electroporation (Choi *et al*., 2006). Site-specific integration of miniTn7 derivatives in *P. putida* strains was performed essentially as described (Choi *et al*., 2005). The flagellar cluster and *rpoN* gene were deleted by plasmid-based gene replacement as previously described (Martínez-García and de Lorenzo, 2021). For *rpoN* deletion, 500-bp fragments flanking the respective gene were PCR-amplified with oligonucleotide pairs rpoN-fwd-EcoRI/rpoN-overlap-Rev (upstream region) and rpoN-overlap-fwd/rpoN-rev-BamHI (downstream region), subsequently combined by overlap extension PCR and cloned via *Eco*RI and *Bam*HI restriction sites into plasmid pEMG, a Km-resistant plasmid that incorporates two I-SceI sites flanking a *lacZ*α polylinker, to yield pMRB290. For flagellar cluster deletion, the plasmid pEMG-flagella (Martínez-García *et al*., 2014) containing 750-bp upstream and 816-bp downstream of PP_4297 and PP_4329 genes respectively was used. These plasmids were mobilised by triparental mating to *P. putida* KT2442 harbouring the I-SceI expressing plasmid pSW-I (Martínez-García and de Lorenzo, 2011) The positive co-integrates were selected through PCR amplification of flanking regions and resolved through induction of the I-SceI enzyme, derived from the pSW-I plasmid using 15 mM 3-methyl benzoate (3-MB) Then, the induced culture was plated onto LB agar plates and the positive colonies were assessed for the loss of the Km resistance marker. Deletions were verified by PCR. The pSW-I plasmid was cured after successive passes of the deleted strains in LB without antibiotics.

To construct *P. putida* KT2442 derivatives with in-frame deletions of *fleN*, *fleQ* and *fleR*, upstream and downstream chromosomal regions flanking these genes were PCR-amplified with an specific oligonucleotide pairs. For *fleN* deletion, fleN-up-fwd/fleN-up-rev (upstream region) and fleN-down-fwd/fleN-down-rev (downstream region) For *fleQ* deletion, fleQ-up-fwd/fleQ-up-rev (upstream region) and fleQ-down-fwd/fleQ-down-rev (downstream region) For *fleR* deletion, fleR-up-fwd-EcoRI/fleR-up-rev-BamHI (upstream region) and fleR-down-fwd-BamHI/fleR-down-rev-HindIII (downstream region) The PCR products were cleaved with EcoRI and BamHI or BamHI and HindIII, respectively, and three-way ligated into EcoRI- and HindIII-digested pEX18Tc, a gene replacement vector. A BamHI-excised FRT-flanked kanamycin resistance cassette from pMPO284 was then cloned into the BamHI site, yielding pMRB96 (*fleQ*), pMRB104 (*fleN*) and pMRB192 (*fleR*) These plasmids were transferred to *P. putida* KT2442 by electroporation. Selection of integration, allelic replacement and FLP-mediated excision of the kanamycin resistance marker was performed essentially as described (Hoang *et al*., 1998; Llamas *et al*., 2000) to generate Δ*fleQ* (MRB52), Δ*fleN* (MRB71) and Δ*fleR* (MRB93) The structure of the deleted loci was verified by PCR.

For the construction of pMRB178 and pMRB286, plasmids producing FleQ and FliA, respectively, the *fleQ* and *fliA* open reading frames were PCR-amplified with the oligonucleotide pairs FleQ-XbaI-SD-fwd/ FleQ-SpeI-rev and FliA-fwd-XbaI-SD/FliA-rev-SpeI-Stop, digested with appropriated enzymes and cloned into XbaI- and SpeI-cleaved pMRB172, a salicylate-inducible expression plasmid bearing Tn7 transposon for site-specific genome integration.

A 750-bp fragment containing the putative P*fleS* promoter region (positions -750 to - 1 relative to *fleS* start codon) was amplified using oligonucleotides PfleS-fwd/PfleS-rev as primers and cloned via SpeI and PstI restriction sites into previously cleaved pMRB196, a pMRB3-derived broad host-range *gfp*mut3::*lac*Z transcriptional fusion vector, yielding pMRB291. P*fliE* and P*flgF* 500-bp promoter regions were amplified by using oligonucleotide pairs PfliE-fwd-SpeI/PfliE-rev-PstI and PflgF-fwd-SpeI/PflgF-rev-PstI and cloned as described above to yield pMRB250 (P*flgF*) and pMRB259 (P*fliE*)

### Flagellar motility (swimming) assays

Swimming assays were adapted from Parkinson (1976). Tryptone motility plates containing 0.3% Bacto-agar (Difco) were toothpick-inoculated with fresh colonies and incubated for 14 h at 30°C. At least 3 biological replicates were assayed for each strain.

### *In vivo* gene expression assays

Differential rates of GFP accumulation during exponential growth were used to measure gene expression in cultures bearing *gfp-lacZ* transcriptional fusions to different promoters. Overnight LB cultures with the corresponding antibiotics of the fusion-bearing strains were diluted in 1/10 strength LB broth with the same additions, and dispensed into the wells of a Costar 96 microtiter polystyrene plate (Corning) (150 µl per well). The plate was incubated in a Tecan Spark microliter plate reader/incubator at 30 °C with 510 rpm shaking to mid-exponential phase. Cultures were then diluted in the same medium and incubated in the same conditions for 23 hours. A_600_ and GFP fluorescence (485 nm excitation, 535 nm emission) were monitored in 15-minutes intervals during incubation with the gain value set to 55. Differential rates of GFP fluorescence accumulation were calculated as the slopes of the linear regression (R^2^>0.98) performed on the exponential growth fluorescence vs. absorbance plots (Supplementary Fig. 3) (Jacob and Monod, 1961). The results (arbitrary units) were divided by 1000 to avoid excessive trailing zeros in the chart axis legends.

For end-point fluorescence assays cells bearing the corresponding fusion plasmids grown on LB with the appropriate additions were diluted in 3 ml of the same medium to an A_600_ of 0.01 and incubated at 30°C with shaking (180 rpm) for 24 hours (stationary phase). 0.5 ml samples were withdrawn, cells were washed three times in 1xNaCl-phosphate buffer (M9 salts lacking NH4Cl) (Sambrook and Russell, 2000) suspended in 5 ml 1xNaCl-phosphate buffer. Samples (150 µl) were transferred to Costar 96 plates and end-point absorbance and fluorescence measurements were performed as above.

For β-galactosidase assays, the fusion strains were grown and diluted as above and incubated for 4.5 or 24 hours (for exponential or stationary phase assays, respectively). Growth was then stopped, and β-galactosidase activity was determined from sodium dodecyl sulphate- and chloroform-permeabilized cells as described (Miller, 1993). Results are reported as Miller units (Miller, 1993).

For all expression assays, data are reported as the average and standard deviation of at least three biological replicates. Significance of the differences reported was assessed by means of the Student’s t-test for unpaired samples not assuming equal variance. Differences were considered significant when the *p-*value was below 0.05.

### RNA isolation and RT-PCR

Total RNA from stationary phase cultures was extracted as described (García-González *et al*., 2005) Reverse transcription (RT) of total RNA (2 µg) was carried out using the High-Capacity cDNA Achieve kit (Applied Biosystems) following the manufacturer instructions. Amplification reactions were performed with 50 ng of cDNA as template in a 30-cycle PCR program, using appropriate oligonucleotides detailed in **Supplementary Table S2** and Illustra^TM^ PuReTaq^TM^ Ready-To-Go^TM^ PCR Beads (GE Healthcare) Negative and positive controls were performed with total RNA or genomic DNA as templates, respectively. The RT-PCR products obtained were resolved by 1% agarose gel electrophoresis and visualized by ethidium bromide staining.

### RNA-seq assay

5 µg of total RNA were sent to the CABIMER Genomics facility (Seville, Spain) Depletion of rRNA and preparation of cDNA libraries were carried out with the Illumina Stranded TOTAL RNA preparation RIBO-ZERO PLUS kit. Single-end sequencing was performed with the Illumina NextSeq 500 HIGH-Output flow cell and 1 x75 bp length parameters at the CABIMER Genomics unit (Centro Andaluz de Biología molecular y Medicina regenerativa, Sevilla, Spain)..

The RNA-seq raw data was pre-filtered using FASTQ Toolkit v.1.0.0. Quality of the reads was checked with FastQC (https://www.bioinformatics.babraham.ac.uk/projects/fastqc/) and then mapped to the *P. putida* KT2440 reference genome (GenBank: NC_002947.4) using Bowtie2 v.2.3.5.1 (Langmead and Salzberg (2012). Mapped data were converted to BAM files using Samtools v1.11 (Li *et al*., 2009) and reads were visualized with the Integrative Genomics Viewer (IGV) (Robinson *et al*., 2011). Raw dataset is available at the Gene Expression Omnibus (GEO) database (http://www.ncbi.nlm.nih.gov/geo) under accession number GEO: GSE173832. The genome-aligned data were displayed using the Integrated Genomics Viewer (IGV) software.

### Bioinformatics methods

We used the pre-computed *Pseudomonas* ortholog groups available at the *Pseudomonas* genome database (Winsor *et al*., 2016) (<u>pseudomonas.com</u>) to assess the conservation of the flagellar gene set in the genus *Pseudomonas*. Synteny was analyzed by computing the number of intervening genes between the orthologs of all consecutive genes in the *P. putida* flagellar cluster in all 550 genomes in the *Pseudomonas* ortholog database using a simple Microsoft Excel script and the numerical part of the Locus Tag for each pair of genes in each genome as a positional reference. The resulting dataset was manually curated and genomes yielding inconsistent or ambiguous results were discarded. A total of 401 genomes provided consistent results for the complete cluster (**Supplementary File 1**) Sequence alignment of intergenic sequences used 500 bp fragments upstream from the start codon for each gene. Occasional inconsistencies in the annotation of the start codons were corrected when required. Multiple sequence alignments were performed using MUSCLE (Edgar, 2004), available at https://www.ebi.ac.uk/Tools/msa/muscle/.

## ACKNOWLEDGMENTS

We wish to thank María G. Velasco, Laura Claret and Álvaro Escobar for their help in some of the constructions used in this work, the CABIMER Genomics unit for their technical help with the RNA-seq assays, Guadalupe Martín for technical assistance and all members of the Govantes and Santero laboratories at CABD and the Andalusian Network of Bacterial Flagella Researchers (RedFlag) for providing materials and critical discussions. This work was funded by the Spanish Ministerio de Ciencia, Innovación y Universidades and European Regional Development fund, grants BIO2013-420173-P and PGC2018-097151-B-I00 awarded to FG. M P-S is the recipient of a PhD training grant (FPU) of the Spanish Ministerio de Ciencia, Innovación y Universidades. The authors declare no conflict of interest.

## SUPPORTING MATERIALS LEGENDS

**Supplementary File S1. Synteny analysis of the flagellar cluster in 401 *Pseudomonas* genomes.** Each of the four groups is shown in a separate tabs. The values indicate the number of intervening genes between each of the gene pairs above in each genome. Yellow frames indicate the flagellar operons of *P. putida* KT2440. Absent genes are denoted by an X.

**Supplementary File S2. Sequence alignments of the flagellar promoter regions in seven environmental *Pseudomonas* strains.** Output of the MUSCLE alignment in CLUSTALW format. PpuKT2440: *P. putida* KT2440; PpuF1: *P. putida* F1; PsyDC3000: *P. syringae* pv tomato DC3000; PsyB728a: *P. syringae* pv syringae B728a; Psa1448a: *P. savatanoi* 1448a; PflSBW25: *P. fluorescens* SBW25; PprPF5: *P. protegens* PF5. Asterisks indicate positions conserved in all seven strains. Coding sequences are shown in blue, and putative promoter motifs are shown in red.

**Supplementary File S3. Statistical analysis of the differential rate of fluorescence accumulation dataset represented in** Figs. 5 and 6. The average, standard deviation, the numerical and symbol representation of the T-test results and the fold difference in pairwise comparisons are shown for each combination of promoter fusion and genetic background. Fold differences are positive when expression is greater in the reference strain and negative when expression is smaller in the reference strain. Lines are color-coded as follows. Green: statistically significant difference, fold change greater than 2; yellow: statistically significant difference, fold change smaller than 2; red: difference not statistically significant.

**Supplementary Table S1. Inventory of flagellar genes *P. putida* KT2440 flagellar cluster.** *P. putida* locus tags are according to the TIGR sequencing project annotation available at the *Pseudomonas* Genome Database (<u>pseudomonas.com</u>). Putative functions are based on experimental evidence obtained in *P. putida* or other organisms.

**Supplementary Table S2. Class II flagellar gene expression in *Escherichia coli*.** The values show *β*-galactosidase activity (Miller units) stationary phase cultures of *E. coli* ET8000 bearing the empty vector pSB1K3, or pMRB99, expressing *P. putida* KT2440 FleQ from its own promoter grown in LB. Also shown are the number of biological replicates (n) for each fusion and strain, the averages and standard deviations, the fold differences (ET8000/pMRB99 / ET8000/pSB1K3 ratio) in the *β*-galactosidase activity and the T-test *p-* values are shown. ns: *p*≥0.05*; = *p*<0.05; ** = *p*<0.01; *** = p <0.001.

**Supplementary Table S3. Effect of *rpoN* deletion on Class II flagellar gene expression.** The values show *β*-galactosidase activity (Miller units) from exponential and stationary phase cultures of *P. putida* KT2442 (wild-type) and MRB149 (Δ*rpoN*) grown in LB containing 10 mM glutamine. The number of biological replicates (n) for each fusion and strain, the averages and standard deviations, the fold differences (wild-type/Δ*rpoN* ratio) in the *β*-galactosidase activity and the T-test *p-*values are shown. ns: *p*≥0.05*; = *p*<0.05; ** = *p*<0.01; *** = p <0.001.

**Supplementary Table S4. Bacterial strains, plasmids and oligonucleotides used in this work.** Underlined bases indicate oligonucleotide positions that differ from the corresponding templates. ^1^Reference list for this Table shown as Supplementary References below.

**Supplementary Figure S1. RNA-seq analysis of the *motAB* operon.** Output of the RNA-seq analysis of the genomic region containing the *motAB* operon. Coverage is represented in grey on a logarithmic scale. Top strand and bottom strand reads are shown in blue and pink, respectively. A cartoon of the flagellar genes is aligned with the RNA-seq reads. Genes are color-coded as in Fig. 1.

**Supplementary Figure S2. Identification of putative flagellar promoters.** Sequences from the upstream regions of selected flagellar genes from *P. putida* KT2440 and F1, *P. syringae* DC3000 and B728a, *P. savastanoi* 1448a, *P. fluorescens* SBW25 and *P. protegens* Pf-5 with the putative promoters aligned along with the established consensus for *σ*^54^, FliA and *σ*^70^-dependent promoters. No gaps are introduced, except when required for alignment of the promoter motifs. Promoter sequences are indicated in red and coding sequences of the upstream genes are indicated in blue.

**Supplementary Figure S3. An example of the GFP fluorescence vs. absorbance plot.** Fluorescence data collected along the growth curves of the wild-type (red) and Δ*fleQ* (blue) strains bearing the P*flgB-gfp-lacZ* fusion plasmid pMRB272 were plotted against the A_600_ data. Circles, squares and triangles denote three independent biological replicates, with the darker colors denoting the data used for the linear regression and the lighter colors denoting those not used. Dotted lines denote the linear fit obtained for the selected data of each replicate.

**Supplementary Figure S4. RT-PCR of *fleQ* expressed from its natural genomic context and ectopically.** Ethidium bromide-stained 1% agarose electrophoresis gel showing the amplification reaction products obtained with *fleQ*-specific (lanes 1-5) and *fliT*-specific (lanes 6-10) oligonucleotides. The template DNA was KT2442 genomic DNA (lanes 1 and 6), KT2442 cDNA (lanes 2 and 7), MRB130(Δflagella) cDNA (lanes 3 and 8), or MRB130/miniTn*7-nahR-*P*sal-fleQ* cDNA (lanes 4 and 9). Lanes 5 and 10 contain reactions performed with no template DNA. The picture shows one representative result of three biological replicates.

**Supplementary Figure S5. Putative FleQ binding sites at the Class II promoter regions. A.** Cartoon of each Class II promoter region showing the putative s54 promoter in red and the putative FleQ binding sites in green, along with the distances between the different *cis-*acting elements. **B. and C.** The putative FleQ binding sites, aligned, and the logo derived from the sequences of these sites using WebLogo 3.7.4. (Crooks *et al*., 2004) (+) and (-) indicate the orientation of the sequences as shown in **B.** relative to the promoter (coding and non-coding strand, respectively).

**Supplementary Figure S6. Swimming assay of the Δ*fleR* and Δ*fleS* mutants.** Swimming halos obtained in tryptone-soft agar plates with the wild-type strain KT2442 and its Δ*fleQ*, Δ*fleR* and Δ*fleS* derivatives. The images show representative halos from identical plates grown in parallel in the same conditions. Cropping was performed maintaining the scale of the respective images.

