## Supplemental File 2 for "Transcriptional organization and regulation of the *Pseudomonas putida* flagellar system": Supplementary file 2-All alignments.pdf

**Supplementary File S2. Sequence alignments of the flagellar promoter regions in seven environmental *Pseudomonas* strains.** Output of the MUSCLE alignment in CLUSTALW format. PpuKT2440: *P. putida* KT2440; PpuF1: *P. putida* F1; PsyDC3000: *P. syringae* pv tomato DC3000; PsyB728a: *P. syringae* pv syringae B728a; Psa1448a: *P. savatanoi* 1448a; PflSBW25: *P. fluorescens* SBW25; PprPF5: *P. protegens* PF5. Asterisks indicate positions conserved in all seven strains. Coding sequences are shown in blue, and putative promoter motifs are shown in red.

**P*fliK2***

**MUSCLE alignment (https://www.ebi.ac.uk/Tools/msa/muscle/)**

**CLUSTAL multiple sequence alignment by MUSCLE (3.8)**

```
PpuKT2440_PfliK2_PP_4328      GCGGCTGGCCGGCTGGTGCAGGGCGCACAGCCAGGTGAGATTCTCCTCGGCGGTGAGCAA
PpuF1_PfliK2_Pput_1539        ------------------------------------------------------------
PsyDC3000_PfliK2_PSPTO_3636   --------------------------AGAGCCAGGCCAGGTTCTCAAGCGGTGTGAGCAA
PsyB728a_PfliK2_Psyr_3394     ----------------------------AGCCAGGACAGGTTTTCCAGTGGCGTCAGCAC
Psa1448a_PfliK2_PSPPH_3315    -----------------------------GCCAGGAAAGATTCTCCAGCGGTGTCAGCAG
PflSBW25_PfliK2_PFLU_1757     ---------------------------------GCTGAGGTTTTCTTCAGCGGTGAGCAC
PprPF5_PfliK2_PFL_1679        -----------------------------------------TCTCCTCGGGGCTCAACAG

PpuKT2440_PfliK2_PP_4328      GTCCTTGA-TGCCGGCGGCGTGGCCAATCCACAGCAGGGCGCTGGCCAAGGCATGACGCT
PpuF1_PfliK2_Pput_1539        ---------TGCCGGCGGCGTGGCCAATCCACAGCAGGGCGCTGGCCAGGGCATGACGCT
PsyDC3000_PfliK2_PSPTO_3636   GTCCTTGAGCGCTGGC-GCATGGCCAAGCCACAACAGATCGCAGCCCGGCGCT---CGCT
PsyB728a_PfliK2_Psyr_3394     ATCCTTGATCGCTGGC-GCATGACCGATCCACAACGGTTTGCGGCCCGGTGCGGCGGGCT
Psa1448a_PfliK2_PSPPH_3315    GTCCTTGAGCGCCGAC-GCATGGCCAACCCACAACAGATTGAGCCCCGGCGCTGTCGGCT
PflSBW25_PfliK2_PFLU_1757     GTCCTTGA-TGCCGGCGGCGTGGCCGATCCAGACCAGGTTGCGCGCCAGTTCAGTGCGTT
PprPF5_PfliK2_PFL_1679        CTCCTTGA-TGCCGGCGGCATGGCCGATCCACAACAGGTTGCGGGCCAGTTCCGCGCGCT
                                        ** * * ** ** ** * *** * * *   *    **    *     * *

PpuKT2440_PfliK2_PP_4328      GTTCTGCCAGGGGCTTGCCACCCAGCAGAATCTGCCCGGCCGTCGGCTGCATCAGCCCGG
PpuF1_PfliK2_Pput_1539        GTTCTGCCAGTGGCTTGCCACCCAGCAGAATCTGCCCGGCCGTCGGCTGCATCAGCCCGG
PsyDC3000_PfliK2_PSPTO_3636   GGCGGCCCAGCACTTGCCCGTTGAGCAGCACCTGCCCGGCGGTCGGCTGCATCAGCCCGC
PsyB728a_PfliK2_Psyr_3394     GCCGGCCCAGTGGCTGCGCGTCGAGCAGGATTTGCCCGGCCGTTGGCTGCATCAGCCCGC
Psa1448a_PfliK2_PSPPH_3315    GGCGGCCCGCTGGCTGGCCGTTGAGCAGCACGTGCCCGGCCGTGGGTTGCATCAGCCCGC
PflSBW25_PfliK2_PFLU_1757     GTGCGTTGAGGGGCTTGCCGTTTAACCGAACTTCACCCGCCGTCGGCTGCATCAACCCGG
PprPF5_PfliK2_PFL_1679        GGGCGTGCAATGGCTGGCCATTGAGCAGCACCTGGCCGCTGGTGGGCTGCATCAACCCGG
                              *             *   *    * * * *  *  **    ** ** ******* ****

PpuKT2440_PfliK2_PP_4328      CCAGCAGGCGCAACAGGCTGGTCTTGCCGCTGCCGTTGGGGCCACTGATCTGCAGCATGT
PpuF1_PfliK2_Pput_1539        CCAACAGGCGCAACAGGCTGGTCTTGCCACTGCCGTTGGGGCCACTGATCTGCAGCATGT
PsyDC3000_PfliK2_PSPTO_3636   ACAGCAGGCGCAGCAGGCTGGTCTTGCCGCTGCCGTTGGGGCCGCTGACCTGCAACATGT
PsyB728a_PfliK2_Psyr_3394     ACAGCAGGCGCAGCAGGCTGGTCTTGCCGCAGCCGTTGGGACCGCTGATCTGCAGCATGT
Psa1448a_PfliK2_PSPPH_3315    ACAACAGGCGCAGCAGGCTGGTCTTGCCACTGCCATTGGGGCCGCTGACCTGCAGCATGT
PflSBW25_PfliK2_PFLU_1757     CCAGCAGGCGCAGCAAACTGGTCTTGCCGCTGCCGTTGGGGCCGCTGACCTGCACCATGT
PprPF5_PfliK2_PFL_1679        CCAGCAGGCGTAGCAGGCTGGTCTTGCCGCTGCCGTTGGGGCCGCTGATCTGCACCATGT
                               ** ****** * **  *********** * *** ***** ** **** ***** *****

PpuKT2440_PfliK2_PP_4328      CGCCGGCGCCCAGCTCGAAATCGAGCTGTTCGAACAGCAGGCGCCAGTCGCGCTCGCAGG
PpuF1_PfliK2_Pput_1539        CACCGGCGCCCAGCTCGAAATCGAGCTGTTCGAACAGCAGGCGCCAGTCGCGCTCGCAGG
PsyDC3000_PfliK2_PSPTO_3636   CCCCGGCGCACAGGTGCAGATCGAGGTTTTCGAACAGCACCCGCCAGTCCCGCTCGCATG
PsyB728a_PfliK2_Psyr_3394     CGCCTGTGTGCAGGTGAAGATCGAGGTTTTCGAACAGCATCCGCCCGTCCCGCTCACACG
Psa1448a_PfliK2_PSPPH_3315    CCCCCGTGCGCACCTGCAGATCAAGGTTTTCGAACAGCATCCGCCCGTCCCGCTCACACG
PflSBW25_PfliK2_PFLU_1757     CGCTGCCGGCCAGGCGCAGTTCGAGGTGCTCGAACAGCAGGCGCAGGTCGCGTTCACAGG
PprPF5_PfliK2_PFL_1679        CGCCGCCCGTCAGACGCAACTCGAGGTGTTCAAATAGCATCCGCCAATCGCGCTCACAGG
                              * *       **     *  ** ** *  ** ** ****  ***   ** ** ** ** *

PpuKT2440_PfliK2_PP_4328      CCAGGCCCGCGGCTTGGAGGTGAAG---GGTCACGGTTTCGCCTTATCTTCAGTAGGGCT
PpuF1_PfliK2_Pput_1539        CCAGGCCTGCGGCTTGGAGGTGAAG---GGTCACGGTTTCGCCTTATCTTCAGTAGGGCT
PsyDC3000_PfliK2_PSPTO_3636   CCAGAGCCGTCGCTTGCAAAAATGGGGGGGCT--GGGGTCATC------------AGGCG
PsyB728a_PfliK2_Psyr_3394     CCAGAGCCGTCGCTTGCAAAAATGGGGTGGCT--GGGATCATC------------GGGCG
Psa1448a_PfliK2_PSPPH_3315    TCAGAGCCGTCGCTTGCAAAAATGGGGGGGCT--GGGATCATC------------GGGCG
PflSBW25_PfliK2_PFLU_1757     CAAGCGCTATGGCTTCAAGAAGAGGGCTGGTCAAAAGATCG-C------------GGGCC
PprPF5_PfliK2_PFL_1679        CGAGTGCTACGGCTTCGAGAAGAGGGCTGGTCAAGGGATAG-C------------GGGCC
                                **  *    ****  *      *   **        *   *             ***

PpuKT2440_PfliK2_PP_4328      TT----------------------------------------------------------
PpuF1_PfliK2_Pput_1539        TTGGAGCGGCGTCGCCTGTTTCGCGGGTGAACCCGCTCCCACAGAACCCTGTAGGAGCGG
PsyDC3000_PfliK2_PSPTO_3636   TT---------------------------------------TCCAGTGCTGTTGCGTCGC
PsyB728a_PfliK2_Psyr_3394     TT---------------------------------------TCCAGTGTT-TTGCGTCGC
Psa1448a_PfliK2_PSPPH_3315    TT---------------------------------------TCCAGTGCTGGAGCGTCGC
PflSBW25_PfliK2_PFLU_1757     TT---------------------------------------TAC----------------
PprPF5_PfliK2_PFL_1679        TT---------------------------------------TACA---------------
                              **

PpuKT2440_PfliK2_PP_4328      ----------GAAGCAGCCAACACCGCTCCAGAACCCTTGTGTCTCAAGTCGCCCCGCGC
PpuF1_PfliK2_Pput_1539        GTTCACCCGCGAAGCAGCCAACATCGCTCCAGAACCCTTGTGTCTCAAGTCGCCCCGCGC
PsyDC3000_PfliK2_PSPTO_3636   GAAATGTGTCAAAACAGACACATACGGAGTAAAAG----ACGGTTCAAGTCAGGATTGAT
PsyB728a_PfliK2_Psyr_3394     GAAATGTGCCAAAACAGACACATACGGAGTAAAAC----GCGGTTCAAGTCGGGGTTGAT
Psa1448a_PfliK2_PSPPH_3315    GAAATGTACCAAAACAGACACATACGCAGTAAAAG----GCGGTTCAAGTCGGGGGTGAT
PflSBW25_PfliK2_PFLU_1757     -----------------------------------------GGTTCAAGTCGGCGGTGCA
PprPF5_PfliK2_PFL_1679        -----------------------------------------GGTTCAAGTCGGCAACGGA
                                                                       *  *******

PpuKT2440_PfliK2_PP_4328      GCTGCCGTTA-------------------TAC------TGGCAACC--------------
PpuF1_PfliK2_Pput_1539        GCTGCCGTTA-------------------TAC------TGGCAACC--------------
PsyDC3000_PfliK2_PSPTO_3636   GCAGCCGTTACATGAAGAGAACT------TAA------TGGCGGCCAGCCC---------
PsyB728a_PfliK2_Psyr_3394     ACGGCCGTTACATGAAGAGAAGT------TAA------TGGCGGCTGGTCC---------
Psa1448a_PfliK2_PSPPH_3315    ATGGCCGTTACATGAAGAGAAGT------TAA------TGGCGGCTGGTCC---------
PflSBW25_PfliK2_PFLU_1757     GCGGCCGTTA----AAGAGAAATGCACAATAACGGCTTTGGCGACCGATTTTAGAGAGCT
PprPF5_PfliK2_PFL_1679        GCGGCCGTTA----AAGAG--ATGCAGGATAAATGCATTGGCGGCCTGTTCTAGAGAGCT
                                 *******                   **       ****  *
```

```
PpuKT2440_PfliK2_PP_4328        --------------------------------------ACAGGCGCGGCATTATTACACG
PpuF1_PfliK2_Pput_1539          --------------------------------------ACGGGCGCTCGGCATTATTACACG
PsyDC3000_PfliK2_PSPTO_3636     -----------------GGTTTC-------------------GGCCGGTATTA-TACATA
PsyB728a_PfliK2_Psyr_3394       -----------------GGTTTT-------------------GGCCGGTATTA-TACATG
Psa1448a_PfliK2_PSPPH_3315      -----------------GGTTTT-------------------GGCCGGTATTA-TACATG
PflSBW25_PfliK2_PFLU_1757       GGATCAAATAGTTGTGAGGTTTTTAGCGCTCTCTTTGCAGACGGGCGGCATTA-TACATG
PprPF5_PfliK2_PFL_1679          GCGTCAAACAATTGCAATGTTTTCACCACTC-CCCCAAAGACGGGCGGCATTA-TACATG
                                                                          *  *** **** ****

PpuKT2440_PfliK2_PP_4328        CGCCGCCGCGCTGCCAAGAGGCC-GGGCTCGACAGGTTGT----ATACAATC------
PpuF1_PfliK2_Pput_1539          CGCCGCCGCGCTGCCAAGAGGCC-GGGCTCGACAGGTTGT----ATACAATC------
PsyDC3000_PfliK2_PSPTO_3636     CGA-GTCCTGTTATCAAAGGGGC-AATTTCCTCAGGTCG-----ACATTTCA------
PsyB728a_PfliK2_Psyr_3394       CGA-GTCCTGTTATCAAAGGGGC-AATTTCCTCGGGTCG-----ATAGTTCA------
Psa1448a_PfliK2_PSPPH_3315      CGA-GTCCTATTATCAAAGGGGC-AATTTCCTCAGGTCG-----ATATTTCA------
PflSBW25_PfliK2_PFLU_1757       TGATGCCCTACTCTAAAGAGGGC-AATTTCCCCAG---------AGACGACGCGCGCG
PprPF5_PfliK2_PFL_1679          CGAAGCCCTACGCTTATGAGGGCTAATTTCCACAGGTTGTGTCCGAATGACAGGCGAA
                                 *  * *        *   ** *     **  * *           *
```

**P*parC***

**MUSCLE alignment (https://www.ebi.ac.uk/Tools/msa/muscle/)**

**CLUSTAL multiple sequence alignment by MUSCLE (3.8)**

```
Psa1448A_PparC      GCCATACCCAGCGACAAGGCCTTCACGATCATTGAAAAGGTTTCCGGAATCGTCAAACGC
PsyDC3000_PparC     -CCATGCCCAGCGACAAGGCCTTCACCATCATCGAAAAGGTCTCGGGCATCGTCAAACGG
PsyB728A_PparC      -CGATGCCCAGCGACAAGGCCTTCACCATCATTGAAAAGGTCGCCGGGATCGTCAAACGC
PpuKT2440_PparC     ------------------------------------------------------------
PpuF1_PparC         ------------------------------------------------------------
PflSBW25_PparC      ------CCCAGCGACCAGGCGTTTACCATCATCGACAAAGTGGCGTCGATCCTGCGGCCG
PprPF5_PparC        ------CCCAGCGACCAGGCGTTCACCATCATCGACAAGGTGGCGAAGATCCTCAAACCC

Psa1448A_PparC      TTCGACAATCCCATACACGTTGAAGGCTTCACTGACGACCAGCCGATCAGCACTGCGCAG
PsyDC3000_PparC     TTCGATAACCCGATTCATGTTGAAGGCTTTACCGACGACCAGCCGATCAGCACTGCACAG
PsyB728A_PparC      TTTGACAACCCGATCCATGTCGAGGGTTTTACCGACGACCAGCCGATCAGTACTGCACAG
PpuKT2440_PparC     ---------------------------------------------------CACCGCGCAG
PpuF1_PparC         ------------------------------------------------------------
PflSBW25_PparC      TTTGAAAACCCGATCCACGTTGAAGGCTTTACCGACAACTTTCCGATCAACACGGCGCAG
PprPF5_PparC        TTCGACAACCCGATCCATGTCGAGGGTTTCACCGACAACCTGCCGATCCAGACCGCGCAG

Psa1448A_PparC      TTTCCGACCAACTGGGAACTGTCTTCGGCTCGATCTGCCAGCATTGTGCGCATGCTGGCG
PsyDC3000_PparC     TTTCCGACCAACTGGGAGCTGTCGTCGGCGCGCTCTGCCAGCATTGTGCGCATGTTGGCC
PsyB728A_PparC      TTTCCGACCAACTGGGAACTGTCTTCGGCGCGTTCTGCCAGCATTGTGCGCATGCTGGCC
PpuKT2440_PparC     TACCCTACCAACTGGGAGTTGTCCTCGGCGCGCGCGGCCAGCATCGTGCGGCTGCTGGCC
PpuF1_PparC         -----------------------------GCGCGCGGCCAGCATCGTGCGCCTGCTGGCC
PflSBW25_PparC      TACCCGACCAACTGGGAACTGTCTTCGGCGCGTGCCGCCAGCATCGTTCGCATGCTCGCG
PprPF5_PparC        TACCCGACCAACTGGGAGCTGTCCTCGGCCCGTTCTGCGAGCATCGTGCGCATGCTGGCG
                                                  **  * ** ***** ** **  ** * **
Psa1448A_PparC      ATGGACGGTGTCAATCCAGCGCGACTGGCCTCTGTGGGCTACGGTGAGTTTCAACCGATC
PsyDC3000_PparC     ATGGACGGTATCAATCCGGCGCGGCTTGCGTCGGTAGGCTACGGTGAGTTTCAGCCCATC
PsyB728A_PparC      ATGGACGGCGTCAATCCGGCGCGACTCGCCTCGGTGGGCTACGGTGAGTTTCAGCCGATC
PpuKT2440_PparC     ATGGAGGGCGTCAACCCGGCACGCATGGCTTCGGTGGGCTATGGCGAGTACCAGCCGGTG
PpuF1_PparC         ATGGAGGGCGTCAACCCGGCACGCATGGCTTCGGTGGGCTACGGCGAGTACCAGCCCGTG
PflSBW25_PparC      ATGCAGGGCGTGAACCCCGGACGGCTGGCCTCGGTGGGGTACGGCGAGTTCCAGCCGGTG
PprPF5_PparC        ATGCAGGGGGTGAACCCGGCGCGCATGGCCTCGGTGGGCTACGGTGAGTTCCAGCCAGTG
                    *** * **  * ** ** *  **  * ** ** ** ** ** ** ****  ** **  *
Psa1448A_PparC      GCGCCCAATACCACTGCCGCCGGACGCGCCAAGAACCGCCGCGTAGTGCTGGTTATCTCG
PsyDC3000_PparC     GCCCCCAACACCACGGCAGCCGGACGCGCCAAGAACCGTCGTGTGGTGCTGGTGATCTCG
PsyB728A_PparC      GTTCCCAACACCAGCACCGCCGGACGCGCCAAGAACCGCCGTGTAGTGCTGGTTATCTCC
PpuKT2440_PparC     GCCAGCAACGATACCGCCGAAGGGCGCGCACGCAACCGCCGGGTGGTGCTGGTGATTTCC
PpuF1_PparC         GCCAGCAACGATACCGCCGAAGGGCGCGCACGCAACCGCCGGGTGGTGCTGGTGATTTCC
PflSBW25_PparC      GCCAATAACGCTACGGTGGAAGGCCGCGCGCGCAACCGTCGCGTGGTGCTGGTGGTGTCC
PprPF5_PparC        GCCAACAACGCCACCGCCGAAGGCCGGGCGCGCAATCGCCGGGTGGTGCTGGTGGTGTCG
                    *     **    *     *  ** ** **    ** ** ** ** ********  * **
Psa1448A_PparC      CGAAATCTGGATGTGCGTCGCAGCCTGACCAGTTCAGGGTCGGCCAATGTGCAACCGGAT
PsyDC3000_PparC     CGAAATCTGGACGTTCGCCGCAGCCTGACCGGTTCCGGCACGGCCAATGCGCAGCCCGAC
PsyB728A_PparC      CGGAATCTGGATATTCGCCGCAGCCTGACCAGCGCCGGATCAGCCAATGTGCAGCCGGAT
PpuKT2440_PparC     CGTAACCTGGAAGTGCGCCGCAGCCTGACCGGCTCGGGCAGCGCCAATGCCACACCGGAT
PpuF1_PparC         CGTAACCTGGAAGTGCGCCGCAGCCTGACAGGCTCGGGCAGCGCCAATGCCACACCGGAT
PflSBW25_PparC      CGCAACCTGGATGTACGCCGCAGCCTGACCGGCACGGGCACCGCCAATGCAACACCGGAT
PprPF5_PparC        CGCAACCTGGATGTGCGCCGCAGCCTGACCGGTACCGGCACCGCCCATGCAACACCGGAT
                    ** ** *****  * ** ***********  *  * **    *** ***     ** **
Psa1448A_PparC      GCTGCACTGAGGCGTGCTGGCACACAAACTGCACCGGTGCCAGCAAAGCCGCCGGTACGT
PsyDC3000_PparC     GCTGCATTACGCCGTGCTGGCACACAAACTGCACCGGTGCCAGCAAAGCCGCCGGTACGC
PsyB728A_PparC      GCTGCATTGCGGCGTGCTGGCACACAAACTGCACCGGTGCCAGCAAAGCCGCCGGTACGC
PpuKT2440_PparC     GCCGCATTGCGCCGGGCTGGCACACAAAGTGCACCGG---CAATACAAGCAACGGTA-GC
PpuF1_PparC         GCCGCATTGCGCCGTGCTGGCACACAAAGTGCACCGG---CAATACAAGCAACGGTA-GC
PflSBW25_PparC      GCGGCCTTGAAGCGGGCTGGCACACAAACTGCACCGGCCCCTGCAAAGCCGCCGGTTCGT
PprPF5_PparC        GCGGCACTCAAGCGTGCTGGCACACAAACTGCACCAACCCCGGTCAAAACGCCGGTACAG
                    ** **  *    ** ************* ******     *     *  *  ****
Psa1448A_PparC      GCGAACGC----CGTCAATTCTCCGTCAC----------------------------CTG
PsyDC3000_PparC     GCGAACGC----CGTCAATTCTCCGTCAC----------------------------CCG
PsyB728A_PparC      GCGAACGC----CGTCAATTCTCCGTCAC----------------------------CCG
PpuKT2440_PparC     GAGGCCGTGATACGTCAATTCCCCGTCGC--------------------------CGCTG
PpuF1_PparC         GAGGCCGTGATACGTCAATTCCCCGTCGCTGATACGAACACGATTTTGAATGTGCCGCCG
PflSBW25_PparC      CAGAGTGC----CGTCAATTCTCCGTCGC----------------------------CGG
PprPF5_PparC        GGAAGCGC----CGTCAATTCTCCGTCAC----------------------------CTG
                          *     ********* ***** *                            * *
Psa1448A_PparC      CCCCCT------------------------------AAC-------ATCGTATCGGTATGG
PsyDC3000_PparC     CCCCCT------------------------------AAT------TATCGTATCGGTATGG
PsyB728A_PparC      CCCCCT------------------------------AAA------CATCGTATCGGTATGG
PpuKT2440_PparC     TCCCCTGCAGGCGCGGGTTTACCCGCGAGCCGGGCGACGCGGTGTATGGCACCGGTTGCA
PpuF1_PparC         CCCCCTGTGGGAGCGGGCATGCCCGCGAAGCCAGCGACTCGGTGAATGGCACCGGCTACG
PflSBW25_PparC      CTCAAT------------------------------AACCTA-TGCAATGTCTCGGTCGCG
PprPF5_PparC        CTTTAT------------------------------AACTTAGCGCTATGTCTC-GCCCCG
                         *                              *           *   * *
```

```
Psa1448A_PparC      TCGTTGCTGTACGCAA--------------------------------------------
PsyDC3000_PparC     TCGTCGCTGTACGGAA--------------------------------------------
PsyB728A_PparC      TCGTCGCTGTACGGAA--------------------------------------------
PpuKT2440_PparC     CCGGTGTTCGCGGGTAAACCCGCTCCTACATTGATTGCGCGAGCTTTCGAATTTTGAGCA
PpuF1_PparC         CCGGTGTTCGCGGGCTTGCCCGCTCCCACAGAGGGGCGTGCGAGCTTTGAAGTTTTGCGC
PflSBW25_PparC      CCAATGCCTGCCGG----------------------------------------------
PprPF5_PparC        CTGTGGCCGGGCGG----------------------------------------------
                         *      *

Psa1448A_PparC      --------------------------GGAAAAACATCA---------------
PsyDC3000_PparC     --------------------------GGAAAAACATCA---------------
PsyB728A_PparC      --------------------------GGAAAACAATTA---------------
PpuKT2440_PparC     AGG-------------CAGTGTCCCTGCCAAATCAATAGATAGAGTTTGCTTG
PpuF1_PparC         ATGACGGTTGTTTCTACAGTGTCCCTGCCAAATCAATAGATAGAGTTTGCTTG
PflSBW25_PparC      --------------------------GAGGAACCAACCGA-------------
PprPF5_PparC        --------------------------GAGGAACCAACCGA-------------
                              *   **  *
```

### P*cheA*

**MUSCLE alignment (https://www.ebi.ac.uk/Tools/msa/muscle/)**

**CLUSTAL multiple sequence alignment by MUSCLE (3.8)**

```
PpuKT2440_PcheA     -ACCGCACCATGGACCTGGTCGAGGAAAGCACACCGGTGCTCAATGAGCTGGCCAACGAG
PpuF1_PcheA         -ACCGCACCATGGACCTGGTCGAGGAAAGCACACCGGTGCTCAATGAGCTGGCCAGCGAG
Psa1448A_PcheA      ----------------------AGGAAAGCACACCGCTCATGAGCGGTCTGAGCTCCGAT
PsyDC3000_PcheA     ----------------------AGGAAAGCACGCCGATCATGAATGGTCTGAGTTCCGAC
PsyB728A_PcheA      ----------------------AGGAGAGTACGCCGGTCATGAACGGCCTGAGTTCCGAC
PflSBW25_PcheA      AACCGCACCATGGACCTGGTGGAAAACGCCACGCCCCTGGTCAACGGCATGGCCACCGAG
PprPF5_PcheA        -ACCGCACCATGGACCTGGTGGAGAGCAGCACCCCGCTGGTCAACGGCCTGAGCACTGAA
                                          *       ** **  *  * *  *   **      **

PpuKT2440_PcheA     GCCAAGGCCCTGAGCACCGACTGGCAGCGCTTCATGCGCCGCGAAGTGGCTGCGCCGGAA
PpuF1_PcheA         GCCAAGGCCCTGAGCACCGACTGGCAGCGCTTTATGCGCCGCGAAGTGGCTGCGCCGGAA
Psa1448A_PcheA      GCCAAGGCATTGAGTGACGACTGGGGGCGTTTCATGCGCCGCGAAATCGGTGCTGAAGAA
PsyDC3000_PcheA     GCCAAGGCGTTGAGCGAAGACTGGGGACGCTTCATGCGTCGCGAAATCGGTGCCGAAGAG
PsyB728A_PcheA      GCCAAGGCGTTGAGCGACGACTGGGGACGTTTCATGCGCCGCGAAATCGGTGCCGAAGAG
PflSBW25_PcheA      GCCCAGGCCTTGAGCCACGACTGGGGCCGTTTCATGCGCCGCGAAGTCGGGGCTGAAGAG
PprPF5_PcheA        GCCCAGGCCTTGAGCGCAGACTGGGGGCGCTTCATGCGGCGCGAGGTCGGCGCTGAAGAG
                    *** ****  ****    ******   ** ** ***** *****  * *  **    **

PpuKT2440_PcheA     TTTCGTGATCTGGTCAAGCGAGTCGACAGTTTCCTGACGCACAGCGCCGAGGGTAACCGC
PpuF1_PcheA         TTCCGTGATCTGGTCAAGCGCGTCGACAGTTTCCTGACGCACAGCGCCGAGGGTAACCGC
Psa1448A_PcheA      TTTCGTGAGCTGGCCAAGCGGGTCGATGGTTTTCTGACGCGCACCGAGCAGGAGGCCGAT
PsyDC3000_PcheA     TTTCGTGAACTGGCCAAGCGCGTCGATGGTTTTCTGACGCGTACCGAGAAGGAAAGCCAT
PsyB728A_PcheA      TTCCGCGAGCTGGCCAAGCGGGTCGATGGTTTTCTGACGCGTACCGAGAAGGAAACCCAT
PflSBW25_PcheA      TTTCGTGAGTTGGCGCGTCGGGTCGACGGGTTCTTGTCACGCAGCGAGCAGGAAAACCGC
PprPF5_PcheA        TTTCGCGAGCTGGCGCGCCGGGTCGACAGTTTCCTGACGCGCAGCGAACAGGAAACCCGT
                    ** ** **  ***     ** *****  * **  ** * *  * **   ***    *

PpuKT2440_PcheA     AAGGTTTCCGGGCACCTCAACGACATTTTGCTGGCCCAGGACTATCAGGACCTGACCGGC
PpuF1_PcheA         AAGGTTTCCGGGCACCTCAATGACATTTTGCTGGCCCAGGACTATCAGGACCTGACCGGC
Psa1448A_PcheA      AGGGTTTCCGGGCATCTCAACGATATTTTGCTTGCCCAGGATTACCAGGACCTTACGGGG
PsyDC3000_PcheA     ACGGTTTCCGCGCACCTCAACGATATTCTGCTTGCTCAGGATTACCAGGACCTCACCGGT
PsyB728A_PcheA      CAGGTTTCCGCACATCTGAATGACATTCTGCTCGCCCAGGATTACCAGGACCTTACCGGT
PflSBW25_PcheA      ACGGTTTCCAGCAACCTCAACGACATTCTGCTGGCCCAGGATTACCAGGACCTCACCGGT
PprPF5_PcheA        ACCGTTTCCGGCCACCTCAACGACATTCTGCTGGCGCAGGATTACCAGGACCTGACCGGT
                       ******    * ** ** ** *** **** ** ***** ** ******** ** **

PpuKT2440_PcheA     CAGGTGATCAAGCGCGTCACCACACTGGTGACCGAGGTGGAAAGCAACCTGCTCAAGCTG
PpuF1_PcheA         CAGGTGATCAAGCGCGTCACCGCACTGGTGACCGAGGTGGAAAGCAACCTGCTCAAGCTG
Psa1448A_PcheA      CAGGTTATCAAGCGCGTGACGCAATTGGTTACGGAAGTCGAGAGCAATCTGCTCAAACTG
PsyDC3000_PcheA     CAGGTGATCAAGCGCGTCACGCAACTGGTGACCGAAGTAGAGGGCAATTTGCTCAAGCTG
PsyB728A_PcheA      CAGGTCATCAAGCGTGTGACGCAACTGGTCACTGAAGTCGAGAGCAATCTGCTCAAACTG
PflSBW25_PcheA      CAGGTGATCAAGCGTGTGACCCAATTGGTCACCGAAGTGGAAAGCAACTTGCTCAAATTG
PprPF5_PcheA        CAGGTGATCAAGCGTGTGACCCAGTTGGTCACGGAAGTCGAAAGCAATCTGCTCAAACTC
                    ***** ******** ** **     **** ** ** ** **  ****  *******  *

PpuKT2440_PcheA     GTGCTGATGGCCAGCCAGGTCGACCGCTTTGCCGGTATAAAACATGACCACGATCAATTG
PpuF1_PcheA         GTGCTGATGGCCAGCCAGGTCGACCGCTTTGCCGGTATAAAACATGACCACGATCAATTG
Psa1448A_PcheA      GTGCTCATGGCCAGCCATGTCGATCGCTTCGCAGGCATCGAACATGACGAAGAATCCATC
PsyDC3000_PcheA     GTGCTCATGGCCAGCCATGTCGATCGCTTCGCTGGCATCGAACATGACGAAGAATCCATC
PsyB728A_PcheA      GTGCTCATGGCCAGTCATGTCGATCGCTTCGCAGGCATCGAACATGACGAACAATCCATC
PflSBW25_PcheA      GTGCTTATGGCAGGCCAGGTTGATCGTTTCGCCGGCATTGAACATGACCGCGAAGCGATC
PprPF5_PcheA        GTCTTGATGGCCAGCCAGGTCGATCGTTTCGCCGGTATCGAACATGACCGTGAATCGATC
                    **  * *****  * ** ** ** ** ** ** ** **  ********    *     *

PpuKT2440_PcheA     CGTGCAGAAAAAGATCGAGAAAAACATCCGACTCGGGGTGAAGGTCCGCAGATTCATGCC
PpuF1_PcheA         CGTGCAGAAAAAGATCGAGAAAAACATCCGACTCGGGGTGAAGGTCCGCAGATTCATGCC
Psa1448A_PcheA      CTTGCTGAAAAAGATCCTAAAAAACATCTCGCCAAGGGTGAAGGTCCGCAGATTCATGCC
PsyDC3000_PcheA     CTTGCTGAAAAAGATCCTAAAAAACATCTCGACAAGGGTGAAGGTCCGCAGATTCATGCC
PsyB728A_PcheA      CTCAATGAAAAAGATCCTAAAAAACATCTCGCTCAGGGTGAAGGTCCGCAGATTCATGCC
PflSBW25_PcheA      CTCTCGGAAAAAGATCCACAAAAACATCTCGCCAAGGGTGAAGGTCCGCAGATTCATGCC
PprPF5_PcheA        CTCGCTGAAAAAGATCCGCAAAAACATCTCGCCAAGGGTGAAGGTCCGCAGATTCATGCC
                    *     **********   *********       *************************

PpuKT2440_PcheA     GATAAGCGTGAAGACGTCGTGTCCGGTCAGGACGATGTCGATGATCTGCTGTCCAGCCTT
PpuF1_PcheA         GATAAGCGTGAAGACGTCGTGTCCGGTCAGGACGATGTCGATGATCTGCTGTCCAGCCTT
Psa1448A_PcheA      GATAAACGTGAAGACGTCGTATCCGGACAGGATGATGTAGACGATCTGCTATCGAGCCTC
PsyDC3000_PcheA     GATAAACGTGAAGACGTCGTATCCGGACAGGATGATGTAGACGATCTGCTATCGAGCCTC
PsyB728A_PcheA      GATAAACGTGAAGACGTCGTATCCGGACAGGATGATGTAGACGATCTGCTATCGAGCCTC
PflSBW25_PcheA      GATAAACGTGAAGACGTTATGTCAGGTCAGGATGACGTAGATGACCTGTTATCCAGTTTA
PprPF5_PcheA        GATAAACGGGAAGACGTTATGTCCGGTCAGGACGATGTAGACGATTTGCTTTCCAGCCTT
                    ***** ** ********  * ** ** ***** ** ** ** **  ** * ** **  *

PpuKT2440_PcheA     GGTTTTTAA-----------------------GGAGCACGTTTG-
PpuF1_PcheA         GGTTTTTAA-----------------------GGAGCACGTTTG-
Psa1448A_PcheA      GGCTTCTAAGATCTGTTGACGGTTAATTTTAGGGAGCACCCC---
PsyDC3000_PcheA     GGCTTCTAAGGTCTGTTGACGGTTAATTTTAGGGAGCACCCC---
PsyB728A_PcheA      GGCTTCTAAGGTCTGTTGACGGTTAATTTTAGGGAGCACCCC---
PflSBW25_PcheA      GGCTTCTAA-----------------------GGAGCACACCC--
PprPF5_PcheA        GGATTCTA------------------------GGAGCACCCATTA
                    ** ** **                        *******
```

**P*flhF***

**MUSCLE alignment (https://www.ebi.ac.uk/Tools/msa/muscle/)**

**CLUSTAL multiple sequence alignment by MUSCLE (3.8)**

```
PpuKT2440_PflhF      AAGGTGCTGCAAGCCCTGTTGTCGGAACAGGTGCCAGTGCGCGATATTCGCAGTATTGCC
PpuF1_PflhF          AAGGTGCTGCAAGCCCTGTTGTCGGAACAGGTGCCAGTGCGCGATATTCGCAGTATTGCC
PsyDC3000_PflhF      AACGTGCTGCAGGCGTTGCTGGCCGAGCACGTACCTGTGCGGGATATTCGCAGCATTGCA
PsyB728A_PflhF       AACGTGCTGCAGGCGCTGCTGGCTGAACACGTACCTGTGCGGGATATTCGCAGCATTGCA
Psa1448A_PflhF       AACGTGCTGCAAGCGCTTCTGGCTGAGCATGTGCCTGTGCGTGATATTCGTAGCATTGCA
PflSBW25_PflhF       -AAGTGCTGCAGGCGCTGCTGGCCGAACAAGTGCCGGTACGCGACATTCGCAGCATCGCC
PprPF5_PflhF         AAGGTCCTGCAAGCGCTGCTGGCCGAGCAGGTGCCGGTACGCGACATCCGCAGCATTGCC
                      * ** ***** **  *  ** * ** ** ** ** ** ** ** ** ** ** ** **

PpuKT2440_PflhF      GAGGCCATCGCGAACAACGCGGGCAAGAGTCAAGATACCGCCGCACTGGTGGCGGCGGTG
PpuF1_PflhF          GAGGCCATCGCGAACAACGCCGGCAAGAGTCAAGATACCGCCGCACTGGTGGCGGCGGTG
PsyDC3000_PflhF      GAGGCTATCGCCAACAACGCCGGGAAGAGTCAAGATACCGCCGCTCTGGTGGCCGCCGTG
PsyB728A_PflhF       GAGGCTATCGCCAACAACGCCGGGAAGAGTCAAGATACCGCCGCTTTGGTGGCCGCAGTG
Psa1448A_PflhF       GAAGCTATCGCCAACAACGCCGGCAAGAGTCAAGATACCGCCGCTTTGGTGGCTGCGGTG
PflSBW25_PflhF       GAGGCCATCGCCAACAACGCCGCCAAGAGTCAAGATACTGCCGCCTTGGTGGCTGCGGTG
PprPF5_PflhF         GAAGCCATCGCCAACAATGCCGCCAAGAGTCAAGATACTGCCGCTTTGGTTGCCGCGGTA
                     ** ** ***** ***** ** *  ************** *****  **** ** ** **

PpuKT2440_PflhF      CGCGTCGGATTGTGTCGCGCCATCGTGCAAAGCATTGTCGGCGTTGAGTCGGAGCTACCA
PpuF1_PflhF          CGCGTCGGATTGTGTCGCGCCATCGTGCAAAGCATTGTCGGCGTTGAGTCGGAGCTGCCA
PsyDC3000_PflhF      CGTGTCGGTCTGAGTCGCGCAATCGTCCAAAGCATTGTAGGCGTTGAGCCGGAGCTGCCT
PsyB728A_PflhF       CGCGTGGGTCTGAGCCGCGCAATCGTCCAAAGCATTGTAGGCGTTGAGCCGGAGCTGCCT
Psa1448A_PflhF       CGAGTGGGTCTGAGCCGCGCAATCGTCCAAAGCATTGTAGGTGTTGAGCCGGAGCTGCCT
PflSBW25_PflhF       CGCGTCGGATTGTCGCGTGCAATCGTGCAAAGCATTGTAGGGCTTGACTCCGAGCTGCCT
PprPF5_PflhF         CGCGTCGGATTGTCTCGTGCCATCGTCCAAAGCATTGTAGGCACTGAGTCTGAGCTGCCT
                     ** ** **  **   ** ** ***** *********** **   ***  * ***** **

PpuKT2440_PflhF      GTGATTACCCTGGAGCCAAGGTTGGAACAGATTTTGCTGAATAGTCTGCAAAGGGCCGGG
PpuF1_PflhF          GTGATTACCCTGGAGCCAAGGTTGGAACAGATTTTGCTGAATAGTCTGCAAAGGGCCGGG
PsyDC3000_PflhF      GTTATCACTCTGGAACCAAGGTTGGAACAGATATTGCTCAATAGTTTGCAGAAGGCTGGT
PsyB728A_PflhF       GTTATCACTCTGGAGCCAAGGTTGGAACAGATATTGCTCAATAGTTTGCAGAAGGCTGGT
Psa1448A_PflhF       GTTATCACTCTGGAACCAAGGTTGGAACAGATATTGCTCAATAGTTTGCAGAAGGCTGGT
PflSBW25_PflhF       GTGATCACCTTGGAACCAAGGTTGGAACAAATATTGCTCAATAGTATTCAGAAGGCAGGA
PprPF5_PflhF         GTGATCACCTTGGAGCCAAGGTTGGAACAAATTTTGCTCAGCAGTCTGCAGAAGGCAGGA
                     ** ** **  **** ************** ** ***** *  *** * ** * *** **

PpuKT2440_PflhF      CAGGGTCAGGAAGACGGTGTTCTTCTGGAACCTAGCATGGCCGAGAAGCTGCAACGTTCG
PpuF1_PflhF          CAAGGTCAGGAAGACGGTGTTCTTCTGGAACCTAGCATGGCCGAGAAGCTGCAACGTTCG
PsyDC3000_PflhF      CAGGGTCAGGAAGAAGGCGTTCTGCTTGAACCGAGCATGGCTGAAAAGCTCCAGCGTTCG
PsyB728A_PflhF       CAGGGTCAGGAAGAAGGCGTTCTTCTTGAGCCAAGCATGGCTGAGAAGCTCCAGCGTTCG
Psa1448A_PflhF       CAGGGTCAGGAAGAAGGCGTTCTTCTTGAGCCAAGCATGGCTGAGAAGCTCCAGCGTTCG
PflSBW25_PflhF       CAAGGCCAGGAAGAGGGCGTTCTGCTGGAGCCAAGCATGGCTGAGAAGCTGCAGCGTTCG
PprPF5_PflhF         CAAGGCCAGGAAGAAGGCGTTCTGCTGGAGCCAAGCATGGCAGAGAAGCTGCAGCGTTCG
                     ** ** ******** ** ***** ** ** ** ******** ** ***** ** ******

PpuKT2440_PflhF      CTGATCGAGGCGGCCCAGCGCCAGGAAATGCAGGGTCAGCCGGCCATTCTGCTGGTCGCC
PpuF1_PflhF          CTGATCGAGGCGGCCCAGCGCCAGGAAATGCAGGGTCAGCCGGCCATCCTGCTGGTCGCC
PsyDC3000_PflhF      TTGATTGAAGCGGCCCAGCGTCAGGAGATGCAAGGGTTGCCGGTCATTCTTCTGGTAGCA
PsyB728A_PflhF       TTGATCGAGGCCGCCCAGCGTCAGGAGATGCAGGGGTTGCCGGTAATTCTTCTGGTAGCA
Psa1448A_PflhF       TTGATTGAGGCCGCCCAGCGTCAGGAGATGCAAGGGTTGCCGGTAATTCTTCTGGTAGCA
PflSBW25_PflhF       TTGATCGACGCCGCACAGCGCCAGGAAATGCAGGGCCAACCGGTGATCCTGCTGGTGGCA
PprPF5_PflhF         CTGATCGATGCGGCGCAGCGTCAAGAGATGCAAGGTCAACCGGTGATTCTGTTGGTAGCG
                      **** ** ** ** ***** ** ** ***** **    ****  ** **  **** **

PpuKT2440_PflhF      GGCCCGATCCGCGCCATGTTGTCGCGTTTCGGTCGCCTGGCTGTACCGAATTTGCATGTT
PpuF1_PflhF          GGCCCGATCCGCGCCATGTTGTCGCGTTTCGGTCGCCTGGCTGTACCGAATTTGCATGTT
PsyDC3000_PflhF      GGTCCTGTCCGGGCGATGCTGTCACGATTTGGACGATTGGCTGTACCGAACATGCATGTT
PsyB728A_PflhF       GGGCCTGTCCGGGCGATGCTGTCACGATTTGGACGATTGGCTGTACCGAACATGCATGTT
Psa1448A_PflhF       GGGCCTGTCCGGGCGATGCTGTCACGATTTGGACGATTGGCTGTACCGAACATGCATGTT
PflSBW25_PflhF       GGCCCCGTCCGGGCGATGTTGTCGCGGTTTGGGCGCCTGGCAGTACCGAATTTGCACGTT
PprPF5_PflhF         GGGCCGGTCCGCGCCATGCTCTCGCGGTTCGGCCGTCTCGCCGTCCCGGGTTTGCATGTG
                     ** **  **** ** *** * ** ** ** ** **  * ** ** ***    **** **

PpuKT2440_PflhF      CTGGCGTATCAGGAAATACCTGACAACAAGCAAGTCACCATCGTTGCCACCGTGGGCCCT
PpuF1_PflhF          CTGGCGTATCAGGAAATACCTGACAACAAGCAAGTCACCATCGTTGCCACCGTGGGCCCT
PsyDC3000_PflhF      CTGGCGTACCAGGAAATACCGGACAACAAGCAAGTAACCATCGTAGCGACTGTCGGGCCG
PsyB728A_PflhF       CTGGCGTATCAGGAAATTCCGGACAACAAGCAAGTAACCATCGTAGCGACTGTCGGGCCA
Psa1448A_PflhF       CTGGCGTATCAGGAAATTCCGGACAACAAGCAAGTAACCATCGTAGCGACTGTCGGGCCG
PflSBW25_PflhF       TTGGCTTATCAGGAAATTCCTGACAACAAGCAAGTGACTATCGTCGCGACAGTAGGGCCC
PprPF5_PflhF         CTGGCGTACCAGGAAATTCCGGACAACAAGCAAGTGACCATCGTTGCGACAGTAGGGCCC
                      **** ** ******** ** ************** ** ***** ** ** ** ** **

PpuKT2440_PflhF      AACGGCTGA-GGTAGGGGATA
PpuF1_PflhF          AACGGCTGA-GGTAGGGGATA
PsyDC3000_PflhF      AACGGCTGA-GGTAGTGAGTC
PsyB728A_PflhF       AACGGCTGA-GGTAGTGAGTC
Psa1448A_PflhF       AACGGCTGA-GGTAGTGAGTC
PflSBW25_PflhF       AACGGCTGAGGGTAGTGGGTT
PprPF5_PflhF         AACGGCTGA-GGTAGTGGGTT
                     ********* ***** *  *
```

**P*flhA***

**MUSCLE alignment (https://www.ebi.ac.uk/Tools/msa/muscle/)**

**CLUSTAL multiple sequence alignment by MUSCLE (3.8)**

```
PpuKT2440_PflhA                       ----------------------------CGTAGCGACTCGACCAC------ATAATCCA-
PpuF1_PflhA_start_codon_correcte      -----------------------------GTAGCGACTCGACTAC------ATAATCCA-
PflSBW25_PflhA                        ------------------------GCAGGTTCGCGGACCGGTTACCAATGGACCCGGCAT
PprPF5_PflhA                          ------------------------GTGGTGTCGAGATCAGCCTGT------ATCCGGCG-
Psa1448A_PflhA                        ----------------------------CGTCCGCTCTTGGGCGT------GTGGTGCG-
PsyDC3000_PflhA                       ------------------------------------------------------------
PsyB728A_PflhA                        TCTACTACAGCACCGAGCTTGACCAGGAGATCCCTGCCGGGCTGT------ATCTGGCAG

PpuKT2440_PflhA                       --TCGCGGTTTTATGGCG-------------------------------------ATGTT
PpuF1_PflhA_start_codon_correcte      --TCGCGGTTTTATGGCG-------------------------------------ATGTT
PflSBW25_PflhA                        ACCCGAGGGTATCCCGCGCA---CAGCCACCATAAGGCCGCACAGTAGACGATAAAAAGG
PprPF5_PflhA                          --GCGCTGGCC--GGGCTGA-------ATCT------------------------GGATC
Psa1448A_PflhA                        --GCGCAGATTTGTGACGCAGAGCGTCACCC----------------------------A
PsyDC3000_PflhA                       --GCCCAGGTAT-TGGCTTA---CGTCTACC------------------------AGATC
PsyB728A_PflhA                        TCGCCCAGGTGC-TGGCCTA---CGTCTACC------------------------AGATC
                                         *   *        *

PpuKT2440_PflhA                       TGTCAGGAACGATAAAGCTATCGAGAGAG----TTCGCCACCAT----------------
PpuF1_PflhA_start_codon_correcte      TGTCAGGAACGATAAAGCTATCGAGAGAG----TTTGCCACCAT----------------
PflSBW25_PflhA                        CGTCAAC----TGAAAGCGGACAATGGCGCTTCTGCGCATTGGTAAGCCGGGCCGCGACG
PprPF5_PflhA                          TGGCAGTAGCGCCCATGCAG--CAGAGTC----TGCTCAACGAGTGCAAGAGC-------
Psa1448A_PflhA                        CGGCATT----CCCACGCTG----GAGCG----TGAGGAACGATC---------------
PsyDC3000_PflhA                       CGCCAAT----TCCACTCAGGCCAGGGCA----AGCGCCCCGACCCCCTTGGCGACGTCA
PsyB728A_PflhA                        CGCCAGT----TCCGCGCCGGTCAGGGCA----AGCGCCCCGACCCTCTGGGCGACATCA
                                       * **            *        *

PpuKT2440_PflhA                       --GTTCTCCGGGTCTAGCGATGTAGTCAAAAGCGC----------TGGCGAAGGTATCAC
PpuF1_PflhA_start_codon_correcte      --GTTCTCCGGGTCAAGCGATGTAGTCAAAAGCGC----------TGGCGAAGGTATCAC
PflSBW25_PflhA                        CCCGACCACTGCGTTTGCAGCGGTGTACGGAGACT-------------------------
PprPF5_PflhA                          --AACCTGCGTCTCCTGGAGTATGGCGCCTGTGGT---------------------TATC
Psa1448A_PflhA                        --ATCTCAACTATCGTGCGACGCTCCGCATTGCAT---GCCGTTCCGGGCGCTCTGCGTC
PsyDC3000_PflhA                       AAATCCCTCCGGATTTGCAGCGCGATGCTTAGCCTTCAGTGGCTGTCATAAAGTTGCCTT
PsyB728A_PflhA                        AGATCCCGCCCGACCTGCAGCGCGACGCGTAGTCT-------------------------
                                                      *

PpuKT2440_PflhA                       AGACTCACTAGAGCTACC-ATCGAGCA------------------GTTGGCATGATGTTC
PpuF1_PflhA_start_codon_correcte      AGACTCACTAGAGCTACCAATCGAGTA------------------ATTGGCATGATGTTC
PflSBW25_PflhA                        -------------------AGAGAGCG-------------GCTTTCCTAGGGGCAACCGC
PprPF5_PflhA                          CGGTCATCTGCAGTGATCTGGCGTGTT------------------TCCGTGGTGATTTGC
Psa1448A_PflhA                        CGCTCTTTGCGATTCCCACGTAGAGCGTGAGGAGCGATCATCTCAACTATCGTGCGACGC
PsyDC3000_PflhA                       AGCTCTCTTCCAACGCTCTGGCGTGCG----------CCTTTAAATGTACGGCGCGGCGC
PsyB728A_PflhA                        ------------GCTCCTGGACGCGTG------------------------GTGCGGCGC
                                                            * *                                  *

PpuKT2440_PflhA                       ATGCCTTTGA-----GTAGTCATTGGCTCTCTACATCCGCATAGA---AACCGTGAGCTT
PpuF1_PflhA_start_codon_correcte      ATGCCTTTGA-----GCAGTCATTGGCTCCCTACATCCGCATAGA---AACCATGAGCTT
PflSBW25_PflhA                        AAAACCTTGTCAGAAATT-TCCTCATCTCT------------------ACCCGCGAACCC
PprPF5_PflhA                          ----CGGTGACTCGGGTTGCCAATCAC----------------------TTCCAGGACTG
Psa1448A_PflhA                        ---TCCGCGTCGCATGCCGTTCCGGGCGCTCTGCGTCCGCTCTGTGCGATTCCCACGC-A
PsyDC3000_PflhA                       AGATCTGTGACGCGCAGCGTCACGACAGGC------------------ATTCCCATGCTG
PsyB728A_PflhA                        AGATTTGTGACGCAGAGCGTCACCCACGGC------------------ATTCCCGCGCTG
                                              *                                          *     *

PpuKT2440_PflhA                       TTGCATGATCA-----AGTCGCTCAGATTTTTTTGTCGTGGCGCCACCCTCTCTGTGG-
PpuF1_PflhA_start_codon_correcte      TTGCATGATCA-----AGTCGCTCAGAATTTTTTTGCAGTGGG-CCTCCCTCTCTTTGG-
PflSBW25_PflhA                        C--TGTGGTGAGGGAGCTTGCTCCCGCTGGGGCGCGTAGCGCCCCCAAATCCTATGTATG
PprPF5_PflhA                          GAGGGCGGCCATCCGCATGCATCTGGATGATCCACA----GGC-CAATGCCCGGGCCGGG
Psa1448A_PflhA                        GAGCGTGAGGAACGATCATCTC---------AACTATCGTG---CGACGCTCCGCATTG-
PsyDC3000_PflhA                       GAGTGGTAGGAACGATAGGTGTCCGGAAGAACACTATGGTGGC-CCATGCTCCGCGTGGG
PsyB728A_PflhA                        AAGCGTGAGGAGCGAGATTTTTCCGGCAGACACCTGC-------CCATGCTCCGCGCGGG
                                                *                                 *      *

PpuKT2440_PflhA                       ----CTTTCCAAGAGCTTAAGCCTT------------TGGCCATTGCCGTAGAGGCGATT
PpuF1_PflhA_start_codon_correcte      ----CTTTTCAAGAGGTTCAGCCTT------------GGGCCTTTGCCGTTGAGGCGAGT
PflSBW25_PflhA                        CGATCGGTCAGGTAAATC--GCATC-----------------------------------
PprPF5_PflhA                          GATGCCCTGCGTGAAGTCGTCCGGCGCGACTGGATGCTGGACGCTAGCGCATTGATGGCG
Psa1448A_PflhA                        CATGCCGTTCCGGACGCTCTGCGTC----------------CGCTCTT------------
PsyDC3000_PflhA                       CATGCCGTTCTGGACGCTTTGCGTC----------------CGCTCTTTAGTGTGCGCAG
PsyB728A_PflhA                        CATGCCGTTCCGGACGCTCCGCGTC----------------CGTTCTTGCACAGGTAAAC
                                          *  *     *       *

PpuKT2440_PflhA                       GGGCGGG-AGGTCTCGTTTTTCTTGGTGATTGCTTGCTCATGAGCCACCCGTCAACGCGG
PpuF1_PflhA_start_codon_correcte      GGGGCGGGAGGTCTCGTTTCTCTCGGTGATTGCCTGCTGATGAGCCATCAGCCAACGCGA
PflSBW25_PflhA                        -GGCTGG----------TTTTG----GGGCTGCTACGCAGCCCAGCGGGAGCAAGCTCCC
PprPF5_PflhA                          TGGCGTG----------CTGCC----TGGCTGCCGGAT-------------TAATCACGG
Psa1448A_PflhA                        --GAAGA----------TTTTC----TGACTGCGC----------------AAGGTATGC
PsyDC3000_PflhA                       CGCGAGG----------TTCTC----TGGCTTCAC----------------TGATCACCT
PsyB728A_PflhA                        GAGAAGA----------TTTTC----TGACTGCGC----------------TCAGCGCAC
                                                        *        *  * *
```

```
PpuKT2440_PflhA                       T------GGTGCTCGCCCGGTTCATCTTTTCG------TGTGGGTGTCT-----------
PpuF1_PflhA_start_codon_correcte      T------GGAGTTCGCCCGGTTCATCTTTTCG------TGTGGGTGTCA-----------
PflSBW25_PflhA                        TCGCCACAGGTGCGACATGATCTG-----------------CGGCATGGG----------
PprPF5_PflhA                          CCACTCTGAGGTTTCCGGCATTTTCGGCTCAA------TGCCGGCGCCATATTTCCTTTC
Psa1448A_PflhA                        C-------GGTTTCACATTGCTTCTGCCTTTAAAGCCTTACCAGCACTGGCTTT------
PsyDC3000_PflhA                       C--------GTCTTGCGCAATTTCTACCTCTAAAGCCTTGCCGGCACTGGCTTT------
PsyB728A_PflhA                        C-------GGATTTACGTGACTTCTGTGCCTAAAGCCTTACCGCCACTGGCTTT------
                                                     *

PpuKT2440_PflhA                       -GCCAAAGTTGGAAAGCTTCTTGCA--AAGCCACGCCCAGGCGCCCCCTGGGCGTCAAAG
PpuF1_PflhA_start_codon_correcte      -GCCAAAGTTGGAAAGCTTCTTGCA--AAGCCACGCCCAGGCGCCCCCTGGGCGTCAAAG
PflSBW25_PflhA                        -GCAAAAGTTGGAAGGCTTCTTGCAATACCGCC--TGCAGGACGCCTCCCGGCGTCAAAA
PprPF5_PflhA                          GGTAAAAGTTGGAAGGTTTCTTGCAGTA--GCAGCCCCGTGCCGCTT-CAGGCGTCAAAA
Psa1448A_PflhA                        -GTAAAAGTTGGACAGCTTTTTGCAACAGGGTCGATACAGGTCGCCA-TCGGCGTCAATG
PsyDC3000_PflhA                       -GTAAAAGTTGGACAGCTTTTTGCAATAAGGTCGATACAGGTCGCCA-TCGGCGTCAATG
PsyB728A_PflhA                        -GTAAAAGTTGGACAGATTTTTGCAACAGGGTCGATACAGGTCGCCA-TCGGCGTCAATG
                                       *  *********  * ** *****  *         *  *   *     ********

PpuKT2440_PflhA                       TTTTGCATCAA---------GAGGACTCGCG
PpuF1_PflhA_start_codon_correcte      TTTTGCATCAA---------GAGGACTCGCG
PflSBW25_PflhA                        GTTTGCTTCAAGGAAACGAGGAATACCGGTG
PprPF5_PflhA                          GTTTGCTTTACGGAACGGGGTA--AATCG--
Psa1448A_PflhA                        TTTTGATCTAGTGCGCAGGTAAACAATCA--
PsyDC3000_PflhA                       TTTTGATCTAGTGCGCAGGTAACGAATCA--
PsyB728A_PflhA                        TTTTGATCTAGTGCGCAGGTAACGAATCA--
                                       ****    *           *  *
```

**P*fliL***

**MUSCLE alignment (https://www.ebi.ac.uk/Tools/msa/muscle/)**

**CLUSTAL multiple sequence alignment by MUSCLE (3.8)**

```
PsyDC3000_PfliL      ------------------------------------------------------------
PsyB728A_PfliL       ------------------------------------------------------------
Psa1448A_PfliL       ------------------------------------------------------------
PpuKT2440_PfliL      AGCCAGCAGAACTCGGTCGCCTCGACATTCGCGTCAATGTTGCGGCGGACCAGGCCACCC
PpuF1_PfliL          ------------------------------------------------------------
PflSBW25_PfliL       ------------------------------------------------------------
PprPF5_PfliL         -------------------------------------------------CCAGCAGACCC

PsyDC3000_PfliL      ----------------ACTCCGCGTTGGCAAGCAGTTCGTGATGC--TCCGCGTCGCATC
PsyB728A_PfliL       ----------------------------------TGGCAGGACGC-----------CAGG
Psa1448A_PfliL       -------------------------TATT-----ACGCATGATGCTCCTTGCCTCACATG
PpuKT2440_PfliL      AGGTCACCTTCATCAGCGGCCACGCCGGC-----GTACGTGACGC------CCTCGACAG
PpuF1_PfliL          ---TCACCTTCATCAGCGGCCACGCCGGC-----GTACGTGATGC------CCTCGACAG
PflSBW25_PfliL       --------------AGCGCCCACCCGGTG-----GTGCGTGAAGC------GCTGGAAAG
PprPF5_PfliL         AGGTCACCTTCATGAGTGCCCATGCTGGC-----GTTCGCGAAGC------CCTGGAAGG
                                                          *  ** **

PsyDC3000_PfliL      TGTGTTGCAACTCA---ACGTTATGTCTTACTCA-TAGTCACT-----GCACATAAACGA
PsyB728A_PfliL       CTAGCCGCACCGTG-----------------CCA-------------------TGGATGG
Psa1448A_PfliL       GCAGCCGCGACGCG---AAGTCATGCCTTGTCTGCTAGTTGTTAAGGCAGAAATGCTTTT
PpuKT2440_PfliL      CCAGGTGCACCGCCTGCGCGAACTGT-TTGCCCAGCAGGGGTT--GGCGCAGCCGGATGT
PpuF1_PfliL          CCAGGTGCACCGCCTGCGCGAACTGT-TTGCCCAGCAGGGGTT--GGCGCAGCCGGATGT
PflSBW25_PfliL       CCAGTCCGGCCGTCTGCGGGAGATGT-TTGCCCAGCAGGGAAT--GGGGCAGGTGGACGT
PprPF5_PfliL         GCAGATGCACCGCATGCGCGACATGT-TCAACCAGCAGGGCCT--GGGGCAGGTGGACGT
                        *      *

PsyDC3000_PfliL      AAGCGCTTTTTACCGCGCTGGCACTGACGACGCAGGGTGCGACGTAAGTGACAGG-----
PsyB728A_PfliL       CG-------------CGTTG----CGGCGACCC------CCGGATCAGTGTCAGG-----
Psa1448A_PfliL       CGATCC---------CGATGGTACAGACGACGCAAATTGCGACCTTAGTGACAGG-----
PpuKT2440_PfliL      CAACGT---------GGCTG--ACCAGTCGCGCGGG---CAGCAACAACAGCAGG---GG
PpuF1_PfliL          CAACGT---------GGCTG--ACCAGTCGCGCGGG---CAGCAACAACAGCAGG---GG
PflSBW25_PfliL       CAACGT---------CTCCG--ACCAGTCCCGTGGC---CAGGAGCAGCAACAAC---AA
PprPF5_PfliL         CAACGT---------CTCCG--ATCAGTCCCGTGGCTGGCAGGGCCAGGAGCAGGCGCAG
                                        *          *        *      *    **

PsyDC3000_PfliL      ---GCGAAGGAACCCGACGA----------------AGTCGAGCCGAAATCT--------
PsyB728A_PfliL       ---GCGAAGGAACCCGACGA----------------AGTCGGGCCGGAAACCGGAGCGAG
Psa1448A_PfliL       ---GCGAAGGAACCCGACGA----------------AGTCGGGCCGGAAACAGGAGCGAG
PpuKT2440_PfliL      CAGGCGCAGG---GCAGCAACCTTTCCGG-------GGTGGCGGCG--------------
PpuF1_PfliL          CAGGCGCAGG---GCAGCAACTTTTCCGG-------GGTGGCGGCG--------------
PflSBW25_PfliL       CAGGCCCAGA---CCCGTGGCGTGAGCAGCAGCGGCGGTCGGGGTGGCGATAACGGCGAC
PprPF5_PfliL         CAGGAGCAGAATCGCCGCAGCGGTTCCAGCACCAGCGGCGGGCGCCTGGACAATGGCGAT
                        *   **     *                      *  *

PsyDC3000_PfliL      ------------CATTGATTC--------------GCAGATCTGT---------------
PsyB728A_PfliL       GGGTTTTGCCTACTTTGGCCCCATCAAAGTAGGTCGCCGAGAGGCGAAAAGGTGCCCTGA
Psa1448A_PfliL       GGGATTTGCCTACTTTGGCCCCATCAAAGTAGGTCGCCGAGGGGCGAAAAGGTAACCTGA
PpuKT2440_PfliL      ------------CGCCGGGCCGAGCAGGGTGGG--GCCGAAGCGGTCGATAGCGCCCGGC
PpuF1_PfliL          ------------CGCCGCTCCGAGCAGGGTGGG--GTCGAGGCGGTCGATAGCGCCCGGC
PflSBW25_PfliL       AG----------CGGTAGCCCTGCTGATGTG----GCTGAAGCGG----TAGCGCCG---
PprPF5_PfliL         GA--------TGAGTCGTCCCCGGCCGTG------GCAGAAGTGC----AGGCTCC----
                                         *              *  **   *

PsyDC3000_PfliL      --------------------------------------GTGACGCAGAGCGTCACGAAAT
PsyB728A_PfliL       GTCGAACCTCAATCCACCTGACAAAGAAAAACCGCCGTGTGACGCAGAGCGTCACGAACC
Psa1448A_PfliL       GCCTGACACAAACATAACTGACATAGCCGAACCGCTGTGTGACGCAGAGCGTCACGAAAG
PpuKT2440_PfliL      CTTTG---------------------------------GAGCAGCAGGTGGTTGTCGGCG
PpuF1_PfliL          CTTTG---------------------------------GAGCAGCAGGTGGTTGTCGGCG
PflSBW25_PfliL       --------------------------------------GTAACGTCGACGGTGATCGGCT
PprPF5_PfliL         ----------------------------------------GGTACAGACCGTGATCGGCA
                                                                  *   **

PsyDC3000_PfliL      GC----------ATTTCCACGCAGAGCTTGGGAACGATA----ATCAACGGCCTTCGGCC
PsyB728A_PfliL       GC----------ATTCCCACGCTGGAGCGTGAGGAACGATAGTCGTGGCAGCCGCGACGC
Psa1448A_PfliL       GC----------ATTACCACGC--------------------------------------
PpuKT2440_PfliL      ACAGTGCGGTCGATTACTACGC-----------------------------CTGATAACT
PpuF1_PfliL          ACAGTGCGGTCGATTATTACGC-----------------------------CTGATAACG
PflSBW25_PfliL       CCAGCGCCGTCGACTACTACGC--------------------------------------
PprPF5_PfliL         GCAGCGCCGTCGACTACTACGC--------------------------------------
                      *          * *   ****

PsyDC3000_PfliL      AGAATCAAAAGCTGACTTGTGTATGACGCTGAGCGCGGAGAGTGGGAGCGG--------T
PsyB728A_PfliL       GACGTCACGAACGGCATTCCCCCTG----------GGGAGCGTGGGGACGA--------T
Psa1448A_PfliL       ------------------------------------GGAGCGTGGGGACGA--------T
PpuKT2440_PfliL      GTTGGCCTGGGCTGGCTTCTTCGCG-------GGCGAGCCCGCGAAGAAGC---------
PpuF1_PfliL          GTTGGCCTGGGCTGGCTTCTTCGCG-------GGCGAGCCCGCTCCAACAGGCGTTTCGT
PflSBW25_PfliL       ------CTGATCTGATTTTTGCGCT-------GGATGAAATGTGGGAGGGGGCTTGCTCC
PprPF5_PfliL         -----------CTGACCT-----------------------GTGGCGGGGG--------C
                                                              *
```

```
PsyDC3000_PfliL      AATCATCATGGTCCAAGCCCGGTT--TTCAGGGTTCATCAGGACTTATGTCATGTCCAGC
PsyB728A_PfliL       AGTCATCAACTCCCGGGCCGGGCC--CTCAG----CGCTGGCACAAGAATCACGATCCCT
Psa1448A_PfliL       AGTCATCAGCTCCCGGGTCGGGCTGATTCGG----CACTGACACAAGCATCACCATCCCT
PpuKT2440_PfliL      ------------------------------------------------------------
PpuF1_PfliL          TGATTTCGAACCCTGTGC------------GGTAGCTGTGGGAGCGGGCGTGGCCGCGAA
PflSBW25_PfliL       CGATGAGGGGTTGTCAACTGAAAC---ATGGGTGACTGATACACCGCTATCGGGGGCAAG
PprPF5_PfliL         CGCTACCGGCCCCTGCACTGATCC---TCGCGGTGTATCTGCAACAGCTTGCACCCCAGG

PsyDC3000_PfliL      GCCTGTCTCCGCGCGTAACCGTTTGTGTCAGATACATGCCCCTCTCGTACAACTCTGGCA
PsyB728A_PfliL       GCGCGCCCTGGCGCGTAACCGGTTGTGTCCGATACATCCCCGTTTCGTACAACTCTGGCA
Psa1448A_PfliL       CCGCGCCCAGCCGCGTAACCGGTTGTGTCCAATACATCTCCATCTCGTACAACTCTGGCA
PpuKT2440_PfliL      -----CAGCACCGCAAAACCTGCTG----GAATCCATCCC-----CTGCAAAATCTGGCA
PpuF1_PfliL          GCAGCCAGCACCGCAAAACCTGCTG----GAATCCATCCC-----CTGCAAAATCTGGCA
PflSBW25_PfliL       CCCCCTCCCACAATTTGACTTGCCGCGTCCGTACGTCCCCCACTTCAGACAACTCTGGCA
PprPF5_PfliL         TTTCCCTTCGCAGCGTTGCCCTTCGCTGTTGATCCCTGCC--TTTCTGACAAATCTGGCA
                                       *     *              *     *    ** *******

PsyDC3000_PfliL      TAACACTTGCTCTTCCTCTGTCACGTATTGAAGAACCTCTTGAATAGTGACGGATTATTG
PsyB728A_PfliL       TAACACTTGCTCTGCCTCTGTCACGTATTGAAGAACCTCTTGAATAGTGACGGATTATTG
Psa1448A_PfliL       TAACACTTGCTCTGCCTCTGTCACGTATTGAAGAACCTCTTGAATAGTGACGGATTATTG
PpuKT2440_PfliL      TAACACTTGCTCAAGCCAAGCCATTCTCCTTTGAA-CCCCCGATGGACGACGGATTATTG
PpuF1_PfliL          TAACACTTGCTCAAGCCAAGCCATCCTCCCTTGAACCCCCCGATGGACGACGGATTATTG
PflSBW25_PfliL       TAACACTTGCTCTTGCCTTTCCGTAGATCTATGAA-TCCCCGAATAGTGACGGATTATTG
PprPF5_PfliL         TAACACTTGCTCTTGCCTTGCCGTGCGACTGTGAA-CCCCCGATTAGTGACGGATTATTG
                     ************   *     *          ***   *  **     ************

PsyDC3000_PfliL      GC
PsyB728A_PfliL       GC
Psa1448A_PfliL       GC
PpuKT2440_PfliL      GC
PpuF1_PfliL          GC
PflSBW25_PfliL       GC
PprPF5_PfliL         GC
                     **
```

**P*fliK***

**MUSCLE alignment (https://www.ebi.ac.uk/Tools/msa/muscle/)**

**CLUSTAL multiple sequence alignment by MUSCLE (3.8)**

```
PflSBW25_PfliK       ------------------------------------------TT----------------
PpuKT2440_PfliK      -GGGCGTTGCGCGCATGTGGAGTTCGTATGGTGAGCCACGGGTTGGCTCGGGCATAATCT
PpuF1_PfliK          CGGGCGTTGCGCGCATGTGGAGTTCGTATGGTGAGCCACGGGTTGGCTCGGGCATAATCT
PprPF5_PfliK         -------GAGTTTTCTTGGGAGGCTCTGGCATAATTCCCGGTT----------------
PsyDC3000_PfliK      -------------------GGAGTGGCATAATCCGTTCCGGGTT----------------
PsyB728A_PfliK       ---------------------------------CCGCCTGGAGT----------------
Psa1448A_PfliK       ------------------------------------------------------------

PflSBW25_PfliK       --AGTGATTTGGGG--GAGTTGGTGGACCAGATTGATCGGGAGTTTGCGTTGGTGAAGCC
PpuKT2440_PfliK      GGCTTGATCAAGGAGTGAACAAGTGACTGACATGCATATTGACCACAAGGTACTCAGTGA
PpuF1_PfliK          GGCTTGATCAAGGAGTGAACAAGTGACTGACATGCATATTGACCACAAGGTACTCAGTGA
PprPF5_PfliK         --CTTGATCAAGGAGTGAACAAGTGGCTGAGATTCATCTGGACCACACGGTGCTCAATGC
PsyDC3000_PfliK      --CTTGATCAAGGAGTGAACAAGTGTC---GATTCATCTGGATTACAGCGTACTGAGTGC
PsyB728A_PfliK       --CTTGATCAAGGAGTGAACAAGTGTC---GATTCATCTGGATTACAGCGTGTTGAATAC
Psa1448A_PfliK       ----TGATCAAGGAGTGAACAAGTGTC---GATTCATCTGGATTACAGCGTGCTGAATGC
                         ****   **   **    ***      **  **   **        *  * *

PflSBW25_PfliK       GTTGTATGAGACGGAGCGGCAGCGCTTTCAGATTTGAGTTGAGGTTGTGTACATATCCGT
PpuKT2440_PfliK      CCTGCGTGAGG----------------TCA--------TGGAAGATG----GCTACCTGC
PpuF1_PfliK          CCTGCGTGAGG----------------TCA--------TGGAAGATG----GCTACCTGC
PprPF5_PfliK         TTTGCAAGAGG----------------TCA--------TGGAGGATG----AATATCCCG
PsyDC3000_PfliK      CCTGCAGGAAG----------------TCA--------TGGAGGACG----AATATCCGA
PsyB728A_PfliK       CTTGCAGGAGG----------------TCA--------TGGAAGACG----AATATCCAA
Psa1448A_PfliK       CTTGCAGGAAG----------------TCA--------TGGAGGACG----AATATCCGA
                       **   **                  ***        * ** *  *      ** *

PflSBW25_PfliK       TGCTGCGGTAACGGCCACTTAGGG-----GGGCCGCGCTTACAGCGGCTAACCCGTCGTC
PpuKT2440_PfliK      AGTTGGTGCAGACTTTTCTGGACGACTCGGAGCGGCGTCTTGGGCAGTTG---CAC---G
PpuF1_PfliK          AGTTGGTGCAGACTTTTCTCGACGACTCAGAGCGGCGTCTTGGCCAGCTG---CAC---G
PprPF5_PfliK         TCCTGCTGGATACCTTCGTCTGCGATTCCGAGGAGCGTGTGCGGTTGCTG---CAT----
PsyDC3000_PfliK      CTTTGCTGGATGTATTCCTCAAGGATTCCGAGCAGCGTCTCGCCCAACTG---CGCCTTG
PsyB728A_PfliK       CCTTGCTGGATGTCTTTCTCAAGGATTCCGAGCAGCGTATCGCGCAGCTG---CGTCTGG
Psa1448A_PfliK       CCTTGCTGGATGTGTTCCTCAAGGATTCCGAGCAGCGTATCGCGCAACTG---CGTCTTG
                        **  * *        *    *     * *  ***  *        *    *

PflSBW25_PfliK       CTGCCGGGTTCTTACTGTCGATCTCGGTTAAATGTGGGAGGGGGCTTGCTCCCGATAGCA
PpuKT2440_PfliK      CGGCCAAAGC-----GCCGA----------------GGAG-----CTTGGCGCGGCAGC-
PpuF1_PfliK          CGGCCAAAGC-----GCCGA----------------GGAG-----CTTGGCGCGGCAGC-
PprPF5_PfliK         -----AAGTCCGAGGATACGATTTG--------------------CTGGTAGCCACCGC-
PsyDC3000_PfliK      CGGTTGAAACC-----GGCAACCTTGATC---TTCAGGAA-----CTGAGCCTCACGGC-
PsyB728A_PfliK       CTGCCGGGTCC-----CGCAATCCTGACT---TTCAGGAG-----TTGAGCCTTACCGC-
Psa1448A_PfliK       CTGTTCAAGTC-----AGCCATCCTGACT---TTCAGGAA-----TTGAGCCTCACCGC-
                               *       * *                         *          **

PflSBW25_PfliK       GTGGGGCAGTTGCAGGTGTATTAGCTGACACACTGCAATAAGCCCCCTCCCACCGTTTGA
PpuKT2440_PfliK      ---TCACAGTTTCAAGGGCAGCAGCAGCAAC------ATGGGCGCGGTGGCCTTGGCCAG
PpuF1_PfliK          ---CCACAGTTTCAAGGGCAGCAGCAGCAAC------ATGGGCGCGGTGGCCTTGGCCAG
PprPF5_PfliK         ---CCACAGCCTCAAGGGCAGCAGCAGCAAC------ATGGGCGCCACCCGCCTGGCCGA
PsyDC3000_PfliK      ---GCACAGCTTCAAGGGCAGCAGCAGCAAC------ATGGGTGCCCTGCAGCTTTCGCA
PsyB728A_PfliK       ---TCACAGTTTCAAGGGCAGCAGCAGCAAC------ATGGGGGCCTTGCAGCTTTCCGA
Psa1448A_PfliK       ---GCACAGTTTCAAGGGCAGCAGCGCCAAT------ATGGGTGCCTTGCAACTTTCCGA
                           ***   ** * * *  ***    *       **  *  *

PflSBW25_PfliK       TCTCCATCCATCAGGTAGA----------TATCAGCCTGCTTATGATCGTGATCTGGGGC
PpuKT2440_PfliK      CCTGTGT-CAGCAGCTCGAAGAGCGCGCGGCGGCCCCCGTTGTACGGTATTGAAGACT
PpuF1_PfliK          CCTGTGC-CAGCAGCTCGAAGAGCGCGCACGGCGGCCCCCGTTATATGGAATTGAAGACT
PprPF5_PfliK         ACTCTGT-CATCAGCTGGAACTGCGGGCCGGGCAGTCGGCCGCGGATGGCATTCGACAAT
PsyDC3000_PfliK      ATTGTGT-CACCAGCTTGAGGAGCGTGCTCGTCAGAACGATTCCTCGGATCTGCGAGACC
PsyB728A_PfliK       GCTGTGC-CGTCAGCTCGAGGAGCAGGCCCGTCAGGAGCAGTTCGCAGGTCTGGCAGAAT
Psa1448A_PfliK       GTTGTGT-CGTCAGTTGGAAGAGCAGTCCCGTCAGCAGTTGTTCACAGGTCTGACAGAGT
                       *     *  *** * **             * *

PflSBW25_PfliK       ACATCGAGCCCAGGCGAGGTGCCGAGTGCTGGGGCG--------------GA--------
PpuKT2440_PfliK      TGATCAG---CCGCATCGACATGGAATTCGTCGTTGTGCAGCACTTTTACCG--------
PpuF1_PfliK          TGATCAG---CCGCATCGACATGGAATTCGTCGTGGTGCAGCACTTTTACCG--------
PprPF5_PfliK         TGGTGGG---GGAGATCGACGGCGAGTTCGCTATCGTTCGACGCTTGTGCGAGCAGGAGC
PsyDC3000_PfliK      TGATCGG---CAGGATAGGCAGCGAATACCTGACCGTCCGTGGTTTGTTCAA--------
PsyB728A_PfliK       TGATTGG---CACGATCGATAGCGAATACATGACGATTCGCGGGCTGTTCAA--------
Psa1448A_PfliK       TGATCGG---TAAAATCGACAGCGAATACATGACCATTCGCGGTTTGTTTAA--------
                        *             *     ** * *

PflSBW25_PfliK       --------------CCCCGATCAGCCGTTACCCACACAACGGATAGGTACCCGTAAAA--
PpuKT2440_PfliK      --------------TGGTGAGCAGCAAC-GTATTT-CCGCAGGCTGAT------------
PpuF1_PfliK          --------------CGGTGAGCAGCAAC-GTATTT-CCGCAGGCTGAT------------
PprPF5_PfliK         TGCAGCGCTTTCATTGTTGAGAAGCAGCTTTTTAT-CTGCGGATTGAT------------
PsyDC3000_PfliK      --------------TGCTGAGCGCCAGCTCTTCAT-CAGTTGATCGAT------------
PsyB728A_PfliK       --------------TTCCGAGCGGCAGCTGTTCGT-CATCTGAGCCCTTAGAGTCATACC
Psa1448A_PfliK       --------------CGCTGAGCGGCAGTTGTTTGT-CAGCTGATCACTGATTTCCAGATC
                                       **    *           *    *     *
```

```
PflSBW25_PfliK       --------------------------------------TCAAAACCTGGCCCAACTCTTG
PpuKT2440_PfliK      --------------------------------------ACGCGAGTTGGCGCGAGTCTTG
PpuF1_PfliK          --------------------------------------ACGCGAGTTGGCGCGAGTCTTG
PprPF5_PfliK         -----------------------------------------AAAGCTGGCCCGGCCTTTG
PsyDC3000_PfliK      ---AATGCCCTGTCGAAGGGCTGTAT------------TTTAAAGTTGGCCCACATATTG
PsyB728A_PfliK       CTCGATCC-CCGCTATGGGGAATCAT------------TAGATAACTGGCCCGTATATTG
Psa1448A_PfliK       CCCGATCCTCCTTTATTCGGCAACCTTTGAAGTGGGTGTGGAAGGTTGGCCCGCATATTG
                                                                   **** *     ***

PflSBW25_PfliK       CTCT-ACCTCTAGATACTGCATCCCCGATCCCCA--TGCAGTGGAGACCTTTACCT
PpuKT2440_PfliK      C------CTGTCTCTGCCTGAATGATTTTTTGTTTCTGGCGCGGAGACCGCTCG--
PpuF1_PfliK          C------CTGTCTCTGCCTCGATGATT-TTTGTTTCTGGCGCGGAGACCGCTCG--
PprPF5_PfliK         CAGTCAAGTCTTCCAACTGTACCTATGATCCCCG--TGCAGCGGAGACTGTCCC--
PsyDC3000_PfliK      C--T-ATGTCTTGTTG-TATGGCAACAACCAGCAGCTTATGCGGAGACCGAA----
PsyB728A_PfliK       C--T-AACTATTTCGG-TAAGGCGATAACCAGCAGCTGATGCGGAGACCGAA----
Psa1448A_PfliK       C--T-CAGTGTTCTGG-TAAGGCGATAACCAGCAGCTTATGCGGAGACCGAA----
                     *       * *                         *   * ******
```

**P*hsbA***

**MUSCLE alignment (https://www.ebi.ac.uk/Tools/msa/muscle/)**

**CLUSTAL multiple sequence alignment by MUSCLE (3.8)**

```
PsyDC3000_PhsbA     TGTGGTCGAGATGGCAGAAGCGGCCGAGCGTACGGCGGCTCAGCGGCTCGGGCATTTTCA
PsyB728A_PhsbA      --TGGTCGAGATGGCAGAGGCTGCCGAGCGAACTGCCGCCCAGCGCCTTGGGCACTTCCA
Psa1448A_PhsbA      ---GGTCGAGATGGCGGAAGCCGCCGAGCGAACGGCGGCTCAGCGGCTTGGGCATTTTCA
PpuKT2440_PhsbA     ------------------------------------------------------------
PpuF1_PhsbA         ------------------------------------------------------------
PflSBW25_PhsbA      ------------------------------------------------------------
PprPF5_PhsbA        -----TGGACATGGCAGAGAGTGCCGAGCGCACGGCCGCCCAGCGCCTGGAGTACTTTCA

PsyDC3000_PhsbA     GGGGCAGGTCAATCTGGCCAACAACAAGCTGCAGGAGCTGGACCAGTTTCGTCAGGACTA
PsyB728A_PhsbA      GGGGCAGGTCAATCTTGCCAACAACAAGCTGCAGGAACTTGACCAGTTTCGTCAGGATTA
Psa1448A_PhsbA      GGGGCAGGTCAATCTGGCCAACAACAAGCTGCAGGAGCTTGACCAGTTTCGTCAGGATTA
PpuKT2440_PhsbA     ----------------------------------GAGCTGGAGCGCTTTCGCGAGGATTA
PpuF1_PhsbA         ------------------------------------------------------------
PflSBW25_PhsbA      ------------------------------------------------------------
PprPF5_PhsbA        CGGCCAGGTGCGTGTGGCCGAGAGCAAGCTGGGGGACCTGGAGCGTTTTCGTGGCGACTA

PsyDC3000_PhsbA     TCAGCAGCAATGGCTGCAGCGCGGCAGTGCCGGGGTGTCCGGACAGTGGTTGCTGGGCTA
PsyB728A_PhsbA      TCAGCAACAATGGTTGCAGCGCGGCAGTGCCGGGGTTTCCGGGCAGTGGCTGCTGGGTTA
Psa1448A_PhsbA      TCAGCAGCAATGGTTGCAACGCGGCAGTGCCGGGGTTTCCGGGCAGTGGTTACTGGGTTA
PpuKT2440_PhsbA     CCAGTTGCAGTGGATCAACCGCGGCGGGCAAGGGGTCAATGGCAGCTGGCTGGTCAATTA
PpuF1_PhsbA         ------------------------------------------------------------
PflSBW25_PhsbA      -----------------------------------------------------------A
PprPF5_PhsbA        CCAGGAACAGTGGGTCGAGCGCGGCAAGTTGGGGGTCAGTGGCCACTGGCTGATGAACTA

PsyDC3000_PhsbA     CCAGCGCTTCCTCAGCCAGCTTGATGTCGCGGTTGCCCAGCAATACAAAAGTCTCGAATG
PsyB728A_PhsbA      TCAACGCTTTCTCAGTCAGCTCGATGTCGCCGTTGCCCAGCAATACAAAAGTCTTGAATG
Psa1448A_PhsbA      TCAGCGCTTTCTCAGCCAGCTCGATGTCGCCGTTGCCCAGCAATACAAAAGCCTTGAGTG
PpuKT2440_PhsbA     CCAGCGCTTTCTGGGGCAGCTGGAAACGGCCATGACCCAGCAGCGCCAGAGCCTGGTCTG
PpuF1_PhsbA         ---------------------------------------------GCAAAGCCTGGTCTG
PflSBW25_PhsbA      CCAGGGCTTTCTCAACCAATTGGAAACGGCCGTCGGCCAGCAGCGCCAGAGCCTGGCGTG
PprPF5_PhsbA        CCAGTACTTTCTCAGCCAGCTGGATACCGCCGTCGGTCAGCAGCGACAGAGCCTGGCCTG
                                                                   * ** ** *  **
PsyDC3000_PhsbA     GCACAAGGCCAACCTCGATAGAGCGCGCAGTGCCTGGCAGGATTGCTATGCGCGGGTCGA
PsyB728A_PhsbA      GCACAAAGCCAATCTGGATCGGGCGCGCAGCGCCTGGCAGGATTGCTACGCGCGGGTAGA
Psa1448A_PhsbA      GCACAAGGCCAATCTCGATAGGGCGCGGAGCGCCTGGCAGGATTGTTACGCGCGGGTCGA
PpuKT2440_PhsbA     GCACCAGAACAACCTCAATAACGCCCGTGGTACCTGGCAGCAGGCCTATGCCCGGGTGGA
PpuF1_PhsbA         GCACCAGAACAATCTCAATAATGCCCGTGGTGCCTGGCAGCAGGCCTATGCCCGGGTGGA
PflSBW25_PhsbA      GCACCAGAACAACCTCGATAAAGCGCGCGAGAGCTGGCAAGCGGCGTTTGCCCGGGTCGA
PprPF5_PhsbA        GCACCAGAACAACCTCGATCGCGCCCGGGAGGCCTGGCAGCAGGCCTATGCCCGGGTCGA
                    **** *   *** **  **   ** **      ******       *  ** ***** **
PsyDC3000_PhsbA     GGGCTTGCGCAAGCTGGTCCAGCGCTACATGGATGAAGCCCGCAGGCTTGAAGACAAGCG
PsyB728A_PhsbA      GGGCTTGCGCAAGCTGGTGCAGCGGTACATGGACGAGGCGCGCAGGCTGGAAGACAAGCG
Psa1448A_PhsbA      AGGCTTGCGTAAGCTGGTGCAGCGTTATATGGATGAAGCACGCAGGCTGGAGGATAAGCG
PpuKT2440_PhsbA     AGGTTTGCGCAAGCTGGTACAGCGCTACCAGGACGAGGCCCGGCGCGCTGAAGACAAGCG
PpuF1_PhsbA         AGGTTTGCGCAAGCTGGTACAGCGCTACCAGGACGAGGCCCGGCGCGCTGAAGACAAGCG
PflSBW25_PhsbA      AGGCCTGCGCAAGCTGGTGCAGCGCTATATCGACGAAGCGCGGGCACTTGAAGACAAACG
PprPF5_PhsbA        GGGGCTGCGCAAGCTGGTGCAGCGCTATATCGACGAGGCGCGGGCGCTTGAGGACAAGCG
                     **  **** ******** ***** **    ** ** ** **       ** ** ** **
PsyDC3000_PhsbA     CGAGCAGAAGTTGCTCGACGAGTTGTCGCAACGTTTGCCTCGTCACGAACAATTCTGACG
PsyB728A_PhsbA      CGAGCAGAAATTGCTCGACGAGTTGTCGCAGCGCCTGCCTCGTCATGAACAATTCTGACG
Psa1448A_PhsbA      GGAGCAGAAATTGCTCGACGAGTTGTCGCAGCGTCTGCCTCGTCACGAACAATTCTGACG
PpuKT2440_PhsbA     CGAGCAGCGCTTGCTGGATGAGCTGTCGCAGCGTCTGCCGCGGCAAAACCCTCTATAG--
PpuF1_PhsbA         CGAGCAGCGCTTGCTGGATGAGCTGTCGCAGCGCCTACCGCGGCAAAACCCTCTATAG--
PflSBW25_PhsbA      CGAGCAAAAGCTGCTCGACGAACTCTCTCAGCGCCTTCCACGCCAAGAACAGTATTAAGG
PprPF5_PhsbA        TGAGCAGAAGTTGCTGGATGAACTTTCCCAACGCTTGCCTCGTCAGAGCCCATACTGATG
                     *****     **** ** **  * ** ** **  * ** ** **    *     *
PsyDC3000_PhsbA     -----------------------------------------TTTCATTTAATCAGTCCC-
PsyB728A_PhsbA      -----------------------------------------TTTGACTCAATCAGACCCG
Psa1448A_PhsbA      -----------------------------------------TTTGACCCAATCAGACCCG
PpuKT2440_PhsbA     ---GTGTTCGCCGCAGGC--------CTTGCAGCGGAATAATTGGCCCGAAACAAATG--
PpuF1_PhsbA         ---GTGTTCGCCGCAGGC--------CTTGCAGCGGAATAATTGGCCCGAAACAAATGCG
PflSBW25_PhsbA      CGCGGGCTTACCTGCGATGACACACGCGTGTAGTGAGCGGGCTTGCCC--------CGCG
PprPF5_PhsbA        ---GTGCTCA-------------------------------TTTGCCCCG----------
                                                          *
PsyDC3000_PhsbA     ------------------------------------------------------------
PsyB728A_PhsbA      C-----------------------------------------------------------
Psa1448A_PhsbA      C-----------------------------------------------------------
PpuKT2440_PhsbA     ------------------------------------------------------------
PpuF1_PhsbA         ATGTCTTGCCCGCGAATGCGGTGGCGGCTGCGACGGTGTATGACCGGGTATATTGGCTGG
PflSBW25_PhsbA      CTGGGCTGC-----------GAAGCGGCCCTAAAACCTGACACCGCGTTTTACCTGAATA
PprPF5_PhsbA        ------------------------------------------------------------
```

```
PsyDC3000_PhsbA     -------------------GGAGATCTTCCGG----------------------------
PsyB728A_PhsbA      -------------------AGTGATCTCTCTG----------------------------
Psa1448A_PhsbA      -------------------GGTGATTTCTCTG----------------------------
PpuKT2440_PhsbA     -------------------AGAGGTCTTGCCCGCGAAGAG--------------------
PpuF1_PhsbA         CAGGTCCGGCCTATTCGCGGGTGAACCCGCTCCTACCACGGTATTGGGCGACACCACCGA
PflSBW25_PhsbA      ACCCTATGGCCTTTTTGGGGGAGGCTTCGCCCCCAGCGCGG-------------------
PprPF5_PhsbA        ---------------TGCGGGTTGTCCCCCCG----------------------------
                                        *        *

PsyDC3000_PhsbA     ------------------------------TCTGGTCTACGCTATTAATGTCTCGG----
PsyB728A_PhsbA      ------------------------------GCTGGTCTACGCTATCAATGTCTCGG----
Psa1448A_PhsbA      ------------------------------GCTGGTCTACGCTATCAATGTCTCGG----
PpuKT2440_PhsbA     ------------------------------GCCGGTACAGGCCATCAAAGCCATGAGCC-
PpuF1_PhsbA         TTGATGTAGGAGCGGGTTTACCCGCGAAGAGGCCGTACAGGCCATCAAAGCCACGAGCC-
PflSBW25_PhsbA      -------------------------GGCAAGCCCGCTCACTACAAATAGTGCATAGCCCA
PprPF5_PhsbA        ------------------------------GCGAGTGGG---------------------
                                                  *

PsyDC3000_PhsbA     ----------------------------------------TGATGAAACACCTCGAGGTT
PsyB728A_PhsbA      ----------------------------------------TGATGAAACACCTCGAGGTT
Psa1448A_PhsbA      ----------------------------------------TGATGAAACACCTCGAGGTT
PpuKT2440_PhsbA     -----------TGCATCCGCTTGCCTCCCCCACCATCGCCTGCTAAACCTTAAAGCAGTT
PpuF1_PhsbA         -----------TGCATCCGCTTGCCACCCCCACCATCGCCTGCTAAACCTTAGAGCAGTT
PflSBW25_PhsbA      CACACATTTGATGTGTTGTGTTGCCCATAGCCGGTTCACCTGCTAAACCTTGT-GTCATT
PprPF5_PhsbA        ----------------------------------------TGCTAAACCTTTTAGACGTC
                                                            ** * ** *     *   *

PsyDC3000_PhsbA     CCC---------------------------
PsyB728A_PhsbA      CCC---------------------------
Psa1448A_PhsbA      CCCC--------------------------
PpuKT2440_PhsbA     GCATCCGCCATCATCGAAGGAATCGCTAGC
PpuF1_PhsbA         GCATCCGCCATCATCGAAGGAATCGCTAGC
PflSBW25_PhsbA      GCCAATGAC-------AAGGAAGCAGTCGC
PprPF5_PhsbA        GTCAATGAC-------AAGGAAGCAGTCAC
```

**P*fliH***

**MUSCLE alignment (https://www.ebi.ac.uk/Tools/msa/muscle/)**

**CLUSTAL multiple sequence alignment by MUSCLE (3.8)**

```
PsyB728A_PfliH       ------AGGAATTGAACCAGATTCTCGAGAAGCAGTTCTCCGGTAACGCCAACACGTCGC
PsyDC3000_PfliH      ------AAGAGCTGAACCAGATTCTCGAGAAGCAGTTCTCCGGCAACGCCAACACGTCGC
Psa1448A_PfliH       ------AAGAGCTTAACCAGATTCTCGAGAAGCAGTTCTCCGGTAACGCCAACACGTCGC
PpuKT2440_PfliH      ------------------------------------------------------------
PpuF1_PfliH          ------------------------------------------------------------
PflSBW25_PfliH       C-CTCAAAGAGCTCAACACGATTCTGGAGAAGCAGTTCTCCGGCAACTCGAATGCTTCGC
PprPF5_PfliH         CGCTCAAGGAACTCAACCAGATTCTCGAGAAGCAGTTCTCCGGCAACTCCAACGCATCGC

PsyB728A_PfliH       GTACCACACTGGGTGGTATCAAGCGTGCGGCCGATATCATGAACTTTCTCGACAGCTCGA
PsyDC3000_PfliH      GTACTACCCTGGGTGGTATCAAGCGTGCGGCCGACATCATGAACTTCCTCGACAGCTCCA
Psa1448A_PfliH       GTACTACCCTGGGTGGTATCAAGCGTGCAGCCGACATCATGAACTTCCTCGACAGCTCTA
PpuKT2440_PfliH      ------------------------------------------------TCGACAGC----
PpuF1_PfliH          ------------------------------------------------TCGACAGC----
PflSBW25_PfliH       GTACCACCTTGGGTGGCATCAAGCGCGCGGCCGATATCATGAACTTCCTCGACAGCTCGG
PprPF5_PfliH         GGACCACCCTGGGTGGTATCAAGCGTGCAGCCGACATCATGAACTTCCTCGACAGCTCGA
                                                                     ********

PsyB728A_PfliH       TCGAAGGTTCGCTCATGGACTCGATCCGCGAGGTCGACGAAGACCTCTCGGTGCAGATCG
PsyDC3000_PfliH      TCGAAGGCGCGCTCATGGATTCGATTCGCGAGGTCGACGAAGACCTTTCGGTGCAGATCG
Psa1448A_PfliH       TCGAAGGCGCGCTCATGGATTCGATCCGCGAGGTCGACGAAGACCTTTCGGTACAGATCG
PpuKT2440_PfliH      -----------------------------------------GACCTGTCGGAGCAGATCG
PpuF1_PfliH          -----------------------------------------GACCTGTCGGAGCAGATCG
PflSBW25_PfliH       TCGAAGGCCAGTTGATGGACTCGATCCGCGAGATCGACGACACCCTGTCCGGCCAGATCG
PprPF5_PfliH         TCGAAGGCCAGCTGATGGATTCGATCCGCGAAGTGGACGAAGACCTGTCGGGCCAGATCG
                                                                *** ** *  *******

PsyB728A_PfliH       AGGACCTCATGTTCGTCTTCAACAATCTTTCCGATGTCGACGACCGTGGTATCCAGGCGC
PsyDC3000_PfliH      AAGACCTCATGTTCGTCTTCAACAACCTCTCCGACGTGGACGACCGCGGTATACAGGCGC
Psa1448A_PfliH       AAGACCTCATGTTTGTCTTCAACAACCTCTCCGACGTGGACGACCGCGGTATTCAGGCGC
PpuKT2440_PfliH      AAGACCTGATGTTTGTCTTCAACAACCTGGCCGACGTCGACGACCGGGGCATCCAGGCGC
PpuF1_PfliH          AGGACCTGATGTTTGTCTTCAACAACCTGGCCGACGTCGACGACCGGGGCATCCAGGCGC
PflSBW25_PfliH       AAGACCTCATGTTCGTGTTCAACAACCTCTCCGATGTCGACGACCGTGGCATCCAGGCGT
PprPF5_PfliH         AAGACCTGATGTTCGTCTTCAACAACCTCGCCGATGTCGACGATCGCGGCATCCAGGCGC
                     * ***** ***** ** ******** **  **** ** ***** ** ** ** ******

PsyB728A_PfliH       TGCTGCGCGAAGTATCCTCGGACGTGCTGGTACTCGCGCTCAAGGGCTCGGACGAAGGCA
PsyDC3000_PfliH      TGTTGCGCGAAGTGTCTTCGGACGTGCTGGTACTGGCGCTCAAAGGCTCGGACGAGGCAA
Psa1448A_PfliH       TGTTGCGCGAAGTATCCTCGGACGTGCTGGTACTGGCGCTGAAGGGCTCGGACGAGGCGA
PpuKT2440_PfliH      TGTTGCGCGAAGTGTCCTCCGACGTGTTGGTGGTGTCGCTCAAGGGCGCCGACGAGCGGG
PpuF1_PfliH          TGTTGCGCGAAGTGTCCTCCGACGTGTTGGTGGTGTCGCTCAAGGGCGCCGACGAGCGGG
PflSBW25_PfliH       TGCTGCGCGAAGTGTCCTCCGACGTGCTGGTGCTGGCGCTCAAGGGCTCCGACGAAGGCG
PprPF5_PfliH         TGCTGCGCGAAGTCTCTTCCGATGTACTGGTGCTGGCCCTCAAGGGCTCGGACGAAGGGG
                     ** ********** ** ** ** **  ****  *  * ** ** *** * *****

PsyB728A_PfliH       TCAAGGAAAAAATCTTCAAGAACATGTCCAAGCGTGCCTCTGAGCTGTTGCGCGACGATC
PsyDC3000_PfliH      TCAAGGAAAAAATCTTCAAGAACATGTCCAAGCGTGCCGCCGAACTGTTGCGCGACGACC
Psa1448A_PfliH       TCAAGGAAAAAATCTTCAAGAACATGTCCAAGCGTGCCGCCGAACTGTTGCGCGACGACC
PpuKT2440_PfliH      TCAAGGACAAGATCTTCAAGAACATGTCCAAACGTGCTTCGGAACTGCTGCGTGACGATC
PpuF1_PfliH          TCAAGGACAAGATCTTCAAGAACATGTCCAAACGTGCTTCGGAGCTGCTGCGCGACGATC
PflSBW25_PfliH       TCAAGGAAAAGATCTTCAAGAACATGTCCAAGCGTGCCTCGGAACTGTTGCGCGACGACC
PprPF5_PfliH         TCAAGGAAAAGATCTTCAAGAACATGTCCAAGCGTGCGGCGGAACTGCTGCGCGACGACC
                     ******* ** ******************** *****  * ** *** **** ***** *

PsyB728A_PfliH       TGGAAGCAAAAGGTCCGGTGCGTGTCAGCGATGTGGAAACCGCCCAGAAAGAAATCCTCA
PsyDC3000_PfliH      TGGAAGCAAAAGGTCCGGTGCGTGTCAGCGATGTGGAAACCGCCCAGAAAGAAATCCTCA
Psa1448A_PfliH       TGGAAGCAAAAGGTCCAGTGCGTGTCAGCGACGTGGAAACCGCCCAGAAAGAAATCCTCA
PpuKT2440_PfliH      TGGAGGCCAAAGGGCCGGTACGGGTCAGCGACGTGGAAACCGCGCAGAAAGAAATCCTCA
PpuF1_PfliH          TGGAGGCCAAAGGGCCGGTACGGGTCAGTGACGTGGAAACCGCGCAGAAAGAAATCCTCA
PflSBW25_PfliH       TGGAAGCCAAAGGGCCGGTGCGCGTCAGCGACGTCGAAACCGCGCAGAAAGAGATCCTCA
PprPF5_PfliH         TGGAAGCCAAGGGCCCGGTTCGCGTCAGCGATGTGGAAACCGCGCAGAAGGAAATCCTCA
                     **** ** ** ** ** ** ** ** ***** ** ** ******** ***** ** *******

PsyB728A_PfliH       CCATTGCGCGTCGTATGGCCGAGGCCGGCGAGATCGTACTCGGTGGCAAGGGTGGCGAGG
PsyDC3000_PfliH      CCATTGCGCGTCGTATGGCCGAGGCCGGCGAGATCGTACTCGGTGGCAAGGGTGGCGAGG
Psa1448A_PfliH       CCATTGCGCGTCGTATGGCCGAGGCCGGCGAGATCGTACTCGGTGGCAAGGGTGGCGAGG
PpuKT2440_PfliH      CCATTGCCCGGCGCATGGCCGAGGCCGGCGAGATCGTGCTCGGCGGCAAGGGCGCCGAGG
PpuF1_PfliH          CCATTGCCCGGCGTATGGCCGAGGCCGGCGAGATCGTCCTCGGCGGCAAGGGCGCCGAGG
PflSBW25_PfliH       CCATTGCCCGCCGTATGGCCGAAGCCGGAGAAATCGTTCTCGGCGGGAAGGGCGGCGAAG
PprPF5_PfliH         CCATTGCCCGCCGTATGGCCGAAGCCGGAGAAATCGTTCTCGGCGGGAAGGGCGGCGAAG
                     ******* ** ** ******** ***** ** ***** ***** ** ***** * *** *

PsyB728A_PfliH       AAATGATCTAAGGATTCAGC-----------------CGACTC-----------------
PsyDC3000_PfliH      AAATGATCTAAGGATTCAGC-----------------CGACTC-----------------
Psa1448A_PfliH       AAATGATCTAAGGATTCAGC-----------------CGACTC-----------------
PpuKT2440_PfliH      AAATGATCTAAGGTATTCGCGATCCCATAGCCGACGCCGCCTTTTGTAGGAGCGGGTTTA
PpuF1_PfliH          AAATGATCTAAGGTATTCGCGATCCCATAGCCGACGCCGCCTTTTGTAGGCGCGGGTTTG
PflSBW25_PfliH       AAATGATTTAAGG------------------------CGCTTTCC---------------
PprPF5_PfliH         AAATGATCTAAGGT-----------------------CGCTTT-----------------
                     ******* *****                        **  *

PsyB728A_PfliH       ------------------------------------------------------------
PsyDC3000_PfliH      ------------------------------------------------------------
```

```
Psa1448A_PfliH          ------------------------------------------------------------
PpuKT2440_PfliH         CCCGCGAAGGCGTCGGTCCTGGCAATCGAGGTTGTCGCTGCCGACGCATTCGCGGGTGAA
PpuF1_PfliH             CCCGCGAAGGCGTCGGCCCTGGCAATCGAGGTTGTCGCTGCCGACGCATTCGCGGGTGAA
PflSBW25_PfliH          ------------------------------------------------------------
PprPF5_PfliH            ------------------------------------------------------------

PsyB728A_PfliH          -----------------------------------------------------
PsyDC3000_PfliH         -----------------------------------------------------
Psa1448A_PfliH          -----------------------------------------------------
PpuKT2440_PfliH         CCCGCTCCTACAAGGGACCGTGTCGGCCGAACATGGTTTGAGAACATGACAGC
PpuF1_PfliH             CCCGCTCCTACAAGGGACCGTGTTGGCCGAACATGGTTTGAGAACATGACAGC
PflSBW25_PfliH          -----------------------------------------------------
PprPF5_PfliH            -----------------------------------------------------
```

**P*fleS***

**MUSCLE alignment (https://www.ebi.ac.uk/Tools/msa/muscle/)**

**CLUSTAL multiple sequence alignment by MUSCLE (3.8)**

```
PpuKT2440_PfleS                       ------------------------------------------------------------
PpuF1_PfleS                           ------------------------------GCGATCATGCACCCGTACGGGGTCATTGGC
PsyB728A_PfleS                        ------------AACCTGGTCGAGCGCATGGCGATCATGCACCCGTACGGTGTGATTGGC
PsyDC3000_PfleS_start_codon_corr      ------------AACCTGGTCGAGCGCATGGCGATCATGCACCCGTACGGTGTGATCGGC
Psa1448A_PfleS                        ------------AACCTGGTCGAGCGCATGGCGATCATGCACCCGTACGGCGTGATCGGC
PflSBW25_PfleS                        ------------------------------------------------------------
PprPF5_PfleS                          CGTGAGCTGGCCAACCTGGTGGAGCGCATGGCGATCATGCATCCCTATGGTGTGATTGGC

PpuKT2440_PfleS                       ---------------------------------------------------------GAC
PpuF1_PfleS                           GTGTCGGAACTGCCGAAGAAATTCCGCTATGTCGATGACGAAGACGAGCAGATGGTCGAC
PsyB728A_PfleS                        GTGGCTGAGCTGCCAAAGAAGTTTCGCTACGTCGACGACGAAGACGAGCAGATGGTCGAC
PsyDC3000_PfleS_start_codon_corr      GTGGCCGAGCTGCCGAAAAAATTCCGCTATGTCGATGATGAAGACGAGCAGATGGTCGAC
Psa1448A_PfleS                        GTGGCCGAGCTGCCGAAGAAATTCCGCTATGTCGATGACGAGGACGAGCAGATGGTCGAC
PflSBW25_PfleS                        ------------------------------------------------------------
PprPF5_PfleS                          GTCGTCGAGTTGCCGAAGAAGTTCCGCTATGTGGATGACGAAGACGAGCAGTTGGTGGAC

PpuKT2440_PfleS                       AGCCTGCGCAGCGACCTGGAAGAGCGCGTGGCGATCAATGGCCACACACCAAACTTCAGC
PpuF1_PfleS                           AGCCTGCGCAGCGACCTGGAAGAGCGCGTGGCGATCAATGGCCACACACCCAATTTCAGC
PsyB728A_PfleS                        AGCATGCGCAGCGAGATCGAAGAGCGTGTGGCGATCAACAGCAATACGCCGAACTTCGCT
PsyDC3000_PfleS_start_codon_corr      AGCATGCGCAGCGATATCGAAGAGCGCGTGGCCATCAACAGCAACACACCGAACTTCGCA
Psa1448A_PfleS                        AGCATGCGCAGCGATATCGAAGAGCGCGTCGCCATCAACAGCAACACGCCGAATTTCGCG
PflSBW25_PfleS                        ---------------------------------------GGTCATACCCCGGATTTCGGC
PprPF5_PfleS                          AGCCTGCGCAGCGATCTGGAAGAGCGCGTAGCGATCAACGGTCACGCGCCAGACTTCGCC
                                                                              *  *  * **  * ***
PpuKT2440_PfleS                       AACCACGCCATGCTGCCGCCCGAAGGCCTGGACCTCAAGGACTACCTGGGTAGCCTGGAG
PpuF1_PfleS                           AACCACGCCATGCTGCCGCCCGAAGGCCTGGACCTCAAGGACTACCTGGGTAGCCTGGAG
PsyB728A_PfleS                        TCCGGCGCGATGCTGCCGCCTGAAGGTCTTGATCTGAAAGACTACCTCGGTGGTCTGGAG
PsyDC3000_PfleS_start_codon_corr      TCCGGTGCCATGCTGCCGCCTGAAGGGCTTGACCTGAAAGACTACCTGGGTGGTCTGGAA
Psa1448A_PfleS                        TCGGGCGCAATGCTGCCGCCTGAAGGCCTGGACCTGAAAGACTATCTGGGTGGTCTGGAG
PflSBW25_PfleS                        TCCACTGCCATGCTGCCGCCCGAAGGCCTGGACCTGAAGGACTACCTCGGCAACCTGGAA
PprPF5_PfleS                          GCCACGGCCATGCTGCCGCCGGAAGGCCTGGACCTGAAGGACTACCTCGGCAGCCTGGAG
                                            ** *********** ***** ** ** ** ** ***** ** **    *****
PpuKT2440_PfleS                       CAGGGCCTGATCCAGCAGGCGCTGGACGATGCCAACGGTATCGTTGCGCGCGCGGCTGAA
PpuF1_PfleS                           CAGGGCCTGATCCAGCAGGCGCTGGACGATGCCAACGGTATCGTTGCGCGCGCGGCTGAA
PsyB728A_PfleS                        CAGGGCCTGATTCAGCAGGCGCTCGACGATGCCAACGGTATCGTTGCCCGTGCTGCGGAG
PsyDC3000_PfleS_start_codon_corr      CAAGGTCTGATTCAGCAGGCGCTGGACGATGCCAACGGCATTGTGGCTCGCGCAGCGGAG
Psa1448A_PfleS                        CAGGGTCTGATTCAGCAGGCGCTCGACGACGCGAACGGTATTGTTGCCCGTGCTGCAGAG
PflSBW25_PfleS                        CAAGGCCTGATCCAACAGGCCCTGGACGACGCCAACGGCATCGTCGCCCGCGCCGCCGAG
PprPF5_PfleS                          CAAGGGCTGATCCAGCAGGCGCTGGACGACGCCAATGGCATCGTGGCGCGGGCCGCAGAG
                                      ** ** ***** ** ***** ** ***** ** ** ** ** ** ** ** ** ** **
PpuKT2440_PfleS                       CGTTTGCGTATTCGCCGTACCACCCTGGTCGAGAAAATGCGCAAGTACGGCATGAGCCGC
PpuF1_PfleS                           CGTTTGCGTATTCGCCGTACCACCCTGGTCGAGAAAATGCGCAAGTACGGCATGAGCCGC
PsyB728A_PfleS                        CGCTTGCGTATTCGTCGTACCACGCTGGTCGAGAAGATGCGCAAGTACGGTATGAGCCGC
PsyDC3000_PfleS_start_codon_corr      CGCTTGCGGATTCGCCGGACCACCCTGGTCGAGAAGATGCGCAAGTACGGTATGAGCCGT
Psa1448A_PfleS                        CGTTTACGGATTCGTCGCACCACGCTGGTCGAGAAGATGCGCAAGTACGGTATGAGCCGT
PflSBW25_PfleS                        CGCCTGCGTATCCGTCGAACCACGCTGGTGGAGAAGATGCGCAAGTACGGTATGAGCCGC
PprPF5_PfleS                          CGCTTGCGCATCCGTCGCACCACCCTGGTGGAGAAGATGCGCAAGTACGGTATGAGCCGT
                                      **  * ** ** ** ** ***** ***** ***** ************** ********
PpuKT2440_PfleS                       CAGGGCGGTGAGGACCAGGCGGAGGATTGATGTTTGGGCTGGGTGTTCGCCTTGGGGGTT
PpuF1_PfleS                           CAGGGTGGCGAGGAACAGGCCGAGGATTGA------------------------------
PsyB728A_PfleS                        CGTGAGGGCGATGAACAGGCAGATGATTGA------------------------------
PsyDC3000_PfleS_start_codon_corr      CGTGAGGGTGATGAACAGGCGGAGGATTGA------------------------------
Psa1448A_PfleS                        CGCGAAGGTGATGAACAGGCAGATGATTGA------------------------------
PflSBW25_PfleS                        AAAGAAGGTGATGAACAGGCGGATGATTGA------------------------------
PprPF5_PfleS                          CGTGAAGGTGATGAACAGGCGGATGATTGA------------------------------
                                         *  ** ** ** ***** ** ******
PpuKT2440_PfleS                       GCGTTTGGGCGGGGTGTTCGCCTTGAGGTTTTTGTTTGGGCGGCGCTCGATCTCCGCGGC
PpuF1_PfleS                           ------------------CGCCTGGCGGTT------------------------------
PsyB728A_PfleS                        ------------------CGCCG----GTTTTTT--------------------------
PsyDC3000_PfleS_start_codon_corr      ------------------CGCCT----GTTTCAA--------------------------
Psa1448A_PfleS                        ------------------CGCCT----GTCTGTA--------------------------
PflSBW25_PfleS                        ------------------CGCCT----GTTTT----------------------------
PprPF5_PfleS                          ------------------CGCCG----GTTTT----------------------------
                                                        ****     **
PpuKT2440_PfleS                       ACAACAAAACTCACGGCGAACCCTTCTCGGTCTTAACG----------------------
PpuF1_PfleS                           ----------GCCTG-------TGCCGGCTTCTTCGCG----------------------
PsyB728A_PfleS                        GTGCCAAAAACCAAG-------CCCCTGATTTTTCAGG----------------------
PsyDC3000_PfleS_start_codon_corr      ACGCCAAAAATCAAG-------CCCTTGAATTTTCAAG----------------------
Psa1448A_PfleS                        GCGCCAAAAATCAAG-------CCCCTGAATTTTCAGG----------------------
PflSBW25_PfleS                        ----------TCAAG-------CCGCTGATTTTGGGTGTCCGGCCTCGCCAATACTGTTC
PprPF5_PfleS                          ----------TCAAG-------TCTTTGAAAACTGGGT----------------------
                                                 *  *
```

```
PpuKT2440_PfleS                       ------------------------------------------------------------
PpuF1_PfleS                           ------------------------------------------------------------
PsyB728A_PfleS                        ------------------------------------------------------------
PsyDC3000_PfleS_start_codon_corr      ------------------------------------------------------------
Psa1448A_PfleS                        ------------------------------------------------------------
PflSBW25_PfleS                        CGTTTTTGACGGCGTCTGTTGCCGTCATACTGCGTTGCAGCTCCTCCCCATAGCCAGCTA
PprPF5_PfleS                          ------------------------------------------------------------

PpuKT2440_PfleS                       --------------------------------------------------------CAAT
PpuF1_PfleS                           -------------------------------------------------GGCGTGCCCGC
PsyB728A_PfleS                        ------------------------------------------------------------
PsyDC3000_PfleS_start_codon_corr      ------------------------------------------------------------
Psa1448A_PfleS                        ------------------------------------------------------------
PflSBW25_PfleS                        TGAGTCGTCGCTGCGCCTTGTCTGACGGCAACAGCCACTCGTCAAAACGAAACTGTATTG
PprPF5_PfleS                          ------------------------------------------------------------

PpuKT2440_PfleS                       CCTACGTCAAACCCAAACCCCCATCTCCGGCACGGCTATTGCTACACCGCTGGCAACACA
PpuF1_PfleS                           GAAGAGGCCGGCACAAATCCCTATCTCCGGCACGGCTATTGCTACATCGCTGGCAACACA
PsyB728A_PfleS                        -----------------GGTTTTTTTTAGGCACGGGGATTGCTAAGTCTCTCTTAACACA
PsyDC3000_PfleS_start_codon_corr      -----------------GGTTTTTTTTCGGCACGAGGATTGCTAAGTCTCTCTTAACACA
Psa1448A_PfleS                        -----------------GGTTTTTTTTCGGCACGAGGATTGCAAAGTCCCTCTCAACACA
PflSBW25_PfleS                        GCGAGGCCGGGCACCTTGGTTTTTTTTCGGCACAAGTATTGCTACAGCCCTCGCAACGTT
PprPF5_PfleS                          -----------------GGTTTTTTTTAGGCACGGGTATTGCTAAGTCTCTTGCAACGTT
                                                             * *  *****    ***** *   * **   ***
PpuKT2440_PfleS                       CCGTTT-TATGACGGTCAGCCACGCGAGAGA-GCACG
PpuF1_PfleS                           CCGTTTTTATGACGGTCAGCCACGCGAGAGA-GCACG
PsyB728A_PfleS                        CCGTTT-ACTGACGGTTCGCCACGAGAGAGATGAATC
PsyDC3000_PfleS_start_codon_corr      CCGTTT-ACTGACGGTTCGCCACGAGAGAGATGAACT
Psa1448A_PfleS                        CCGTTT-ACTGACGGTTCGCCACGAGAGAGATGAACT
PflSBW25_PfleS                        CCGTTTAACTGACGGTCAGCCAAGCGAGAGA-GCATG
PprPF5_PfleS                          CCGTTTAACTGACGGTCAGCCACGCGAGAGT-GCACG
                                      ******   *******  **** * *****   * *
```

**P*fleQ***

**MUSCLE alignment (https://www.ebi.ac.uk/Tools/msa/muscle/)**

**CLUSTAL multiple sequence alignment by MUSCLE (3.8)**

```
PpuKT2440_PfleQ                       -----GCCTCCGCCGAGCAATAATACCGTAAGGAGCCCCCATGAGC---GCGCTGCAGC
PpuF1_PfleQ                           ACCAGTCGTCGGCCTCCTGATA-------AAGGAGCAAGACATGAGCGAAGTAATCACCC
PsyDC3000_PfleQ_start_codon_corr      ------AACCTGCTGCTCTAG--------GAGAACAATAAAATGAGTGCTGCGCTCAAGC
PsyB728A_PfleQ                        ------AATCTGCTGGCCTG---------AGGAGAACAATAATGAGTGCTGCGCTCAAGC
Psa1448A_PfleQ                        --------GTTGCTGCCCTGAG-------GAGAACAACAAGATGAGTGCTGCGCTCAAGC
PflSBW25_PfleQ                        GAGAAAGGCCTGAGGGCCTT---------GAGGAACACATCATGAGCCAAGCACTGCAAC
PprPF5_PfleQ                          GCACCGGGTCCGCAGTTTTA---------AGGAGGAACACCATGAGTGCTGTCCTGCAAC
                                                 *                   *         *****    *   *    *

PpuKT2440_PfleQ                       AAATCGAAGACACCCGCCTTGCCCTGAGCGAGGCCCTGCAAGTGCGCGACTGGGCTGCCA
PpuF1_PfleQ                           GAATCGAACAGACCCGCGACATGCTGTTGGCGGCGTTGGCCGACCGCGACTGGGAGGCGG
PsyDC3000_PfleQ_start_codon_corr      GTATCGAAGAAACCCGGGAAGCCTTGGTCGGGGCGTTGGCCGAGCGCGACTGGGAGGCTA
PsyB728A_PfleQ                        GTATCGAGGAAACCCGCGAAGCCTTGGTAGGCGCGCTGGCAGATCGTGACTGGGAGGCTA
Psa1448A_PfleQ                        GTATCGAAGAAACCCGGGAAGCCCTGGTGGGGGCGCTTGCCGAACGCGACTGGGAGGCTA
PflSBW25_PfleQ                        GCATTGATGAAACCCGCGAGGCGCTGATGGGCGCGCTGGCGGACCGCAACTGGGACGCCA
PprPF5_PfleQ                          GCATTGAAGAAACCCGTGAGGCGCTGGTCGGCGCCCTGGCCGAACGCAACTGGCAAGCCA
                                        ** **  * *****        **   *  **  *    *  **  *****   **

PpuKT2440_PfleQ                       TTGGTGAGTTGGACTTGGCCTGTCGTGCGCTGATGGATGAATTACTGCGAGAGAATCAGG
PpuF1_PfleQ                           TGGGGGAGTTGGACCTGGAATGCCGTTTGCGCATCGGTGACATGGTGTCCGAAGCCAAAG
PsyDC3000_PfleQ_start_codon_corr      TCGTCAAGCTTGACCTGGCGTGTCGCGAGTGTGTGGATGCGGTGGTCAGCGAAGCGCCTG
PsyB728A_PfleQ                        TCGTCAAGCTGGACCTGGCGTGTCGCGAGTGTGTCGACGCGGTGGTCAGCGAAGCGCCCG
Psa1448A_PfleQ                        TCGTCAAACTTGACCTGGCGTGTCGCGAATGTGTCGATGCGGTGGTCAGCGAAGCGCCTG
PflSBW25_PfleQ                        TCGGCGAACTGGACCTGGGCTGCCGTCACGTGATCGACGAAGTACTCAGCGAAGCACCGG
PprPF5_PfleQ                          TTGGCGAGCTGGACCTGGCGTGCCGTTCATGCATGGAGGACGTGCTCAGCGAGGCTGAGG
                                      * *   *  * *** ***  ** **        * *  *   *  *    **       *

PpuKT2440_PfleQ                       ATGATGAGGCCGAAATCCGGGCCAATCTGGAAGCGCTGATGGCGGTTTATCGTCAGCTGA
PpuF1_PfleQ                           GAAATGAGGCGGACGTGAGTGCCAGTCTCGAGGAATTGCTGGTCGTTTACCGCCAACTAA
PsyDC3000_PfleQ_start_codon_corr      CCGACGAACCTGCGCTGCGTAGCAACCTTGAAGAGCTGCTGGGTGTCTATCGGCAACTGA
PsyB728A_PfleQ                        ACGACGAACCTGCACTGCGCAGCAACCTTGAGGAGTTATTGGGTGTCTACAGGCAACTGA
Psa1448A_PfleQ                        CTGACGAACCGGCTCTGCGCAGCAATCTTGAGGAGTTGTTGGGGGTCTACAGGCAACTGA
PflSBW25_PfleQ                        TAGATGAGGATGCGTTGCGGGAGAAGCTTGAAAGCTTGCTGGCGGTATACCAGCAACTAC
PprPF5_PfleQ                          TCGACGAGGCGGCACTGCGCGACAATCTGGAGGAGTTGTTGGGGGTCTACCGGCAACTGC
                                         * **    *   *  *    *  ** **     *  ***  ** **    ** **

PpuKT2440_PfleQ                       TCGACGTTGCAAGTGACGAGCGTCAATCCTTGGTCGCACGAATGTCGAATATCAACCAAG
PpuF1_PfleQ                           TCGAGGTTGCAAGTGGCGAGCGTCAATCCTTGGTCAATGAAATGACGAAAATCCGTCAGG
PsyDC3000_PfleQ_start_codon_corr      TTGATGTTGCAACCGGCGAGCGTCAAGCCGTCGTCGATGAAATGACGCAAATCCAGAATG
PsyB728A_PfleQ                        TTGATGTTGCAACCGGTGAGCGTCAAGCGGTTGTCGAAGAAATGACTCAAATCCAGAATG
Psa1448A_PfleQ                        TTGATGTTGCAACCGGGGAGCGTCAAGCAGTCGTCGATGAAATGACTCAAATCCAGAATG
PflSBW25_PfleQ                        TTGAGGTGACGACTGGCGAGCGCCAGGCGATTTTCGAAGAGATGTCGCAGATCAACCAAG
PprPF5_PfleQ                          TTGAGGTGGCAACAGGGGAGCGCCAGGCGATAGTCGACGAGATGTCGCAGATCCAGCAAG
                                      * ** **  * *  * ***** **  *  *  **      *** *  * ***    * *

PpuKT2440_PfleQ                       CGAAACACGCCACAAAGGTATACCATCTGTTCAGCTGAAGCTGGGCAGAGAG--CAACGC
PpuF1_PfleQ                           CGAAGAACGCGGCAAAGGTATACCATTTGTTCACCTGACC----ACCGATGTTGCAAAAA
PsyDC3000_PfleQ_start_codon_corr      CCAAAAATGCGACGAAGGTATACCATCTGTTCGGTTGACTGATAATTGTCATT-TAATCC
PsyB728A_PfleQ                        CGAAGAGCGCGACGAAGGTATACCATCTGTTTGGTTAATGTTTAATTGTCATTTTTATTG
Psa1448A_PfleQ                        CGAAGAACGCGACGAAGGTATACCATCTGTTCGGTTAATGTTTAATTGTCATTTTTATTG
PflSBW25_PfleQ                        CGAAAAACGCGTCAAAGGTTTACCATCTGTTCGGCTGAACAGTCGCAGTTAATCCTACCG
PprPF5_PfleQ                          CACAGAACGCGGCAAAGGTTTACCATTTGTTCGGTTGATGTTGA---GTTAATCCCGAAC
                                      *  *    **  * ***** ****** ****    * *         *

PpuKT2440_PfleQ                       TTCG---------GCGCCATAAATTTGACTGCCTTTAAGTTTTTGACTTTACTAGTGGCT
PpuF1_PfleQ                           GCCG--------TGCGCCATTAATTTGACTGATCGGTGTTTTTTGACTTTACTAGTGGCT
PsyDC3000_PfleQ_start_codon_corr      GTCGACAAGAGTTTCGCCATAAATTTGACTGTGTTCGTTTTTTTGACTTAACTAGT-GAT
PsyB728A_PfleQ                        TTCGACAAGAATTTCGCCATAAATTTGACTGTGTCCGTTTTTTTGACTTAACTAGT-GAT
Psa1448A_PfleQ                        TTCGACAAGAATTTCGCCATAAATTTGACTGTGTTCGTTTTTTTGACTTAACTAGT-GAT
PflSBW25_PfleQ                        G-----------TGCGTCATAAATTTGACCGTGCCAGCTTTTTTGACTTAACTAGT-GCT
PprPF5_PfleQ                          ACTG--------CGCGCCATAAATTTGACTGTGCACGCTTTTTTGACTTAACTAGTGGCT
                                                    ** *** ******** *        ********** ****** * *

PpuKT2440_PfleQ                       GTTTTCGAATTTCAGACGTCCGAACACTGACCATTCAGTCGGAATGA-TGTCGAGCACGC
PpuF1_PfleQ                           GTTTTCGAATTTCAGACGTCCGAACACTGACCATTCAGTCGGAATGA-TGTCGAGCACGC
PsyDC3000_PfleQ_start_codon_corr      GTTTACTATTTTCTGGCGTCT---CATGGTTATACCGGCTGAGATGC----CAAGCTGCC
PsyB728A_PfleQ                        GTTTACTATTTTCTGGCGTCT---CAAGGTTTATACAGCTGAGAAGC----CAAGCTGCC
Psa1448A_PfleQ                        GTTTACTATTTTCTGGCGTCT---CAAGGTTTATACAGCTGAGATGC----CAAGCTGCC
PflSBW25_PfleQ                        GTTTTCAGATTTCAGACGTCT--AGTAAGGTAAGAAGTCTTACAGGCGCCTCGA--TTGC
PprPF5_PfleQ                          GTTTTCAGATTTCAGGCGTCT--ATAAGGCAGAAAATGCCTACAAGCGCGTCTAGCTTGC
                                      **** *   **** * ****        *              * *     * *     *

PpuKT2440_PfleQ                       CCCTG----CAGGGCGCCGAT-ATGACTAGGGAAGTTGCTATTGC
PpuF1_PfleQ                           CCCTG----CAGGGCGCCGAT-ATGACTAGGGAAGTTGCTATTGC
PsyDC3000_PfleQ_start_codon_corr      CCCCAATCAT-GGGCATTGAG-TTGACTAGGGAAGTTGCTATTGC
PsyB728A_PfleQ                        CCCCAATCAT-GGGCATTGAG-TTGACTAGGGAAGTTGCTATTGC
Psa1448A_PfleQ                        CCCCAATCAT-GGGCATTGAG-TTGACTAGGGAAGTTGCTATTGC
PflSBW25_PfleQ                        CCTCAATCTCGGGGCAGGAAGTTTTACTAGGGAAGTTGCTATTGC
PprPF5_PfleQ                          CCCTTATCT--GGGCATTGAG-TTGACTAGGGAAGTTGCTATTGC
                                      **         ****    *   * ********************
```

**P*fliS***

**MUSCLE alignment (https://www.ebi.ac.uk/Tools/msa/muscle/)**

**CLUSTAL multiple sequence alignment by MUSCLE (3.8)**

```
PpuF1_PfliS      -----------------------------AAGATGGCACCCTCGAGTTCAACTCGACCAA
PprPF5_PfliS     ----------------------------------------CTGGATTTCGACGACAAGAA
PsyB728A_PfliS   --------------------------------------AATTGACGTTAAGCACGACCGA
PsyDC3000_PfliS  ----------------------------------------------CTCGACACCGCTAC
Psa1448A_PfliS   ----------------------------------------------CTGGACACTGCTAC
PpuKT2440_PfliS  GAAGAGCACCGGGGCGGGGAGCATCTCCATGCTTTCGCAGATGGGGATCAATACCGATCA
PflSBW25_PfliS   ------------------------------------------------------------

PpuF1_PfliS      GTTCACCG------CGGCGATGAACGACAAGAAGCTGGGCTC------------------
PprPF5_PfliS     ATTCACGG------CTGCCATGACCGACAAGAAGCTCGGTGG------------------
PsyB728A_PfliS   ACTCACAA------CAGCGCTGAGCGACAAGAAGCTGGGCAG------------------
PsyDC3000_PfliS  ATTCACCG------CAGCGCTGAACGACAAAAAGCTGGGCAG------------------
Psa1448A_PfliS   ATTCACAG------CAGCGCTGAACGATAAAAAACTGGGCAG------------------
PpuKT2440_PfliS  GCAGACGGGCCTGTTGGTGCTGGACGATACCAAGTGGGACAAGGCTGTTGCCAAGGGTGC
PflSBW25_PfliS   -ACGACTGGGCTGCTGACAATGGATGACAAAAAGTGGGATGCGGCGGCTAAAACCAATGC
                     **              **   ** *  **    *

PpuF1_PfliS      ---CGAAGTTCAGGAGTTGTTCACCGGT------------------------------TC
PprPF5_PfliS     ---TGAAGTGCAGAAGCTGTTTACCGGT------------------GATGGTACC---AA
PsyB728A_PfliS   ---CCAGATTCAGGCGATGTTCACCGGT------GTGACCAATGCTGACGGCACCAGCTC
PsyDC3000_PfliS  ---CCAGATCCAGACCATGTTCACCGGTACGGGCGCAACAAATGCTGACGGTACGGTGGA
Psa1448A_PfliS   ---TCAGATTCAGACGATGTTCACTGGTACGGGTGCAACAAATGCCGACGGTACGGTTGA
PpuKT2440_PfliS  AGCCGATATCGCGGCGGTGTTCACCGGT------------------------------GA
PflSBW25_PfliS   GGCTGACATCAATAGTATGTTCACCGGT------------------------------AA
                      *  *        **** ** ***

PpuF1_PfliS      CAATGGGCTGATCGAGCGCATGAACAAGGCCATCGATCCGTACAAC------------TC
PprPF5_PfliS     CCCGGGTCTGCTGGAACGCATGACTAACGCGATCAAGCCGTTTACCCAGGGCGTGGGTAG
PsyB728A_PfliS   GGGTGGCTTGATCGCACGCATGAACACTGCGCTTGAGCCGTATACG------------AA
PsyDC3000_PfliS  TGGCGGCCTGGTTGCTCGGATGAGCAAAGCGCTTTTACCCTACACG------------AA
Psa1448A_PfliS   TGGCGGCCTGGTTTCGCGTATGACCAAGGCGCTTCTGCCCTACACA------------AA
PpuKT2440_PfliS  CAAGGGCCTGATCAAGCGCATGACTGCGGCCACGGATGCTTACACT------------GG
PflSBW25_PfliS   AAACGGCATGCTGGCGCGCATGAAAAGCGCCACGGACGACTTTGCC------------AA
                     **  ** *    ** ****     **          *

PpuF1_PfliS      CA---CGGACGGCAGCTTGGCCACACGCAAGAGCAACCTGGATAAGGTGGCGAAGAACCT
PprPF5_PfliS     CATGACCGAGGGGATCCTGACCACTCGTACCAAGACCCTGGATATCACCAAGAAAAAGCT
PsyB728A_PfliS   AG---CCGACGGCGTTCTGGCTGGCAAGACCACCAGTCTTAATAAAGTGCAGACGCGACT
PsyDC3000_PfliS  GG---CCGACGGCGTTCTGGCCAGCAAGACAAGCAGTCTGACTAAAGTCCAGACCCGAAT
Psa1448A_PfliS   GT---CTGACGGTGTTCTGGCCAGCAAGACGAGCAGTCTTAATAAAATACAGACTCGTAT
PpuKT2440_PfliS  CA---CGACCGGAATCTTGGCAACGCGGACAAAGAACCTCAACGACAACCTAACTGAGCT
PflSBW25_PfliS   GGCGTCCACGGGCGTTCTAGCCACGCGTTCAACCTCACTGTCCGACAGTCTGAAAGACCT
                      *    **     *  *          *     **             *      *

PpuF1_PfliS      GGCAGACCAGCAGACTGCGCTGGATCGTCGCACCGAGGCTTTGACCGAGTCGCTGACCAA
PprPF5_PfliS     CAGTGATCAACAGGATGCCTTGGATCGCCGGGTGGCTACCCTGACAGCGGTACTGACCAA
PsyB728A_PfliS   GGCCAGTGATCAGGAGGCGCTCAACCGTCGTATTACGACATTGACGGCAACCCTCACCAA
PsyDC3000_PfliS  CGCCAGTGATCAGGATGCCCTTGATCGTCGTATCGAAAGTCTGACGGCCAGCCTGACCAA
Psa1448A_PfliS   TGCCAGTGATCAGGACGCACTGGATCGTCGTATTACAAGCTTGACGGCGAGTTTGACCAA
PpuKT2440_PfliS  CACCAAGCAGCAAGAGGCGCTTGATCGTCGTATTGAGACATTGACGGCCACGCTGACAGC
PflSBW25_PfliS   GTCTAAACAGCAGGATACCTTGAACGAACGCATGGACCTGCTGACCAAGAGCCTCTCCGA
                         * **     *  *  *    **           ****        *  *

PpuF1_PfliS      GAAGTGGGTGGCAATGGACACCGCCGTGGCCAAGCTGAATGCCCAGGCCGACCAGATCAA
PprPF5_PfliS     GAAGTACAACGACATGGACACCCTGGTGGGCAAGCTGAAGGCCACCGCCAGTAACATCAC
PsyB728A_PfliS   GAAATATAACGCAATGGACCTGGTCGTCGGGCAGTTAAAGGCAACGGCTTCAAGCATTAC
PsyDC3000_PfliS  GAAGTACAACGCGATGGACCTGGTGGTCGGGCAGCTCAAAGCGACCGCGACCAGTATCAC
Psa1448A_PfliS   GAAATACAACGCGATGGACCTGGTTGTAGGGCAGCTCAAGGCGACCGCTACCAGCATCAC
PpuKT2440_PfliS  CAAATACAACGCGATGGATACCCTCGTCGCCAAGCTCAATGCCACGAGTTCGAGCGTGAT
PflSBW25_PfliS   CAAATACAACGCCATGGACACCCTGGTCGCTCAATTGCGTCAGCAAAGCACCAGTGTCAT
                  ** *     *  *****       ** *   *  *                    * *

PpuF1_PfliS      ATCGGTTTTCGACGCTATCAACGCTCAGGCCAAGAACTCCTGATCGTTCTCGCCTTTTGC
PprPF5_PfliS     GTCGATGTTTGAAGCCTTGACCGCACAGCAAA---AAGGCTGAT--------ATTCACTC
PsyB728A_PfliS   TTCGATTTTTGAAGCGATGAACGCGCAGAAAA---ACGCCTCGTAAAAAACCAGTGCAGA
PsyDC3000_PfliS  TTCGATATTCGAAGCGATGAACGCGCAGAAAA---ACGCTTCGTAAATTTAATTGACAGC
Psa1448A_PfliS   GTCGATCTTCGAAGCGATGAACGCGCAGAAAA---ACGCTTCGTAAATTCGTTTTACAGA
PpuKT2440_PfliS  GACGACCCTGAACGCGATGAACAAGGCCAATAACGACGACTGAT----------------
PflSBW25_PfliS   GACCACGCTCAATGCCTTGAACAACCCTAAAACCAATACTTGAT----------TGCGGC
                   *     *  * **  * * *         *   *    *  *

PpuF1_PfliS      CATGT-AAAAGCCCGGCAGCGACCCTTGTCGCTCCGGGCTTTTTGCGT------------
PprPF5_PfliS     AGCGCCAAAAGCCCGGCAGCGTTTTTCAGCGCTCC-GGGCTTTTGTAT------------
PsyB728A_PfliS   AGTACCAGGAACCCGACGCTGTGTC-----------GGGTTCTTGCTT------------
PsyDC3000_PfliS  AGTATCAAGAACCCGACTTCGTGTC-----------GGGTTCTTGCTT------------
Psa1448A_PfliS   AGTATCAAGAACCCGACTCCGTGTC-----------GGGTTTCTGCTT------------
PpuKT2440_PfliS  ------------------------------------GG----------------------
PflSBW25_PfliS   CACGAAAAAAGGCCAGACGGTTGTCT----------GGCCTTTTTTTTCATCTGGAGACG
                                                     **
```

```
PpuF1_PfliS         --TGGCACTAAAGTTTTTTGACGCTCAGGCGATA-CACAGGTACAAGCAACCTTTT----
PprPF5_PfliS        --TTGGCCTAAAGTTTTTTGACGTTGCGTCGATATATTGCTTATACGAACCCTTG-----
PsyB728A_PfliS      --TTAGGCTAAAGTTTCCGCGCACTGCGTCGATAGCTTGAGTATCAGTATTCTGG-----
PsyDC3000_PfliS     --TTGGGCTAAAGTTTCTGCGCGATGCGTCGATAGCTTGAATATCAGTATTCTGG-----
Psa1448A_PfliS      --TTGGGCTAAAGTTTCTGCGCCATGCGTCGATAGCTTGAGTATCAGTATTATGG-----
PpuKT2440_PfliS     --TCGTATTCAAGATCTTTGGGGCTAAGCCGATCACTTAGGTAT--GTAGCCCCTGTGTA
PflSBW25_PfliS      ACTCGCGCTAAAGCTTTTTGCGAGCCAGTCGACATATTGGTTACTGAAATCCTTTTCGCT
                      *     * *** *            * ***         **     *

PpuF1_PfliS         --TCGCTGCAACGAGGTAGATCC-
PprPF5_PfliS        ---AGTTTTGAAGAGGTATTAC--
PsyB728A_PfliS      ---ATTTGTTACGAGGCCCATT--
PsyDC3000_PfliS     ---ATTTGTGACGAGGCCCCCCAT
Psa1448A_PfliS      ---ATTTGCGACGAGGTCCCCCTT
PpuKT2440_PfliS     GCCCCGTTCAAAGAGGATCCCCC-
PflSBW25_PfliS      GTCGCCTGTCACGAGGTAGAAAC-
                          *   * ****
```

**P*fliD***

**MUSCLE alignment (https://www.ebi.ac.uk/Tools/msa/muscle/)**

**CLUSTAL multiple sequence alignment by MUSCLE (3.8)**

```
PpuF1_PfliD          -----------------------------TCTAAGGCACTTCCCCGCCTTTTCGTTAT--
PprPF5_PfliD         ---------------------------GCGCTGGTCGTGCTCTCTCGCTTTTTACTTTAA
PsyB728A_PfliD       -------------------------------------------------TTTTTGCTTTCA
PsyDC3000_PfliD      -------------------------------------------------TTTTTACTTTCG
Psa1448A_PfliD       ------------------------------------------------TTTTTTACTTTCG
PpuKT2440_PfliD      CGATTCTGTGATGAATCGCGAAAGGGTGCGGCAACGTACCCTTTCGTTTTCTGATTCTAC
PflSBW25_PfliD       ---GCGCTTGGTCACTAACGGAAGGGCGCGCTGA-GTGCCCTTCCGTTTTTTCGGTTTAG
                                                                     ** *   * *

PpuF1_PfliD          GAGGTGGTAGACATGGACATGAGCGTCAAGCTGAAT-CTGTCCTACCCG-----GCATCC
PprPF5_PfliD         GAGGTGATGGACATGGATATGAGCGTGAAGCTGAAC-TTGTCTTATCCA-----GCGGCC
PsyB728A_PfliD       GAGGTGATGGTCATGGATATGAGTGTGAAGCTGAGC-TTGTCTTATCCG-----GCCGTT
PsyDC3000_PfliD      GAGGTGATGGTCATGGATATGAGTGTAAAGCTGAAC-GTGTCTTATCCG-----GCCGTT
Psa1448A_PfliD       GAGGTGATGGTCATGGATATGAGTGTAAAGCTGAAC-GTGTCTTATCCG-----GCCGCT
PpuKT2440_PfliD      GAGGCAAGCATCATGGATATGAGTGTCAAGCTGAACTCTGTTCTCTCTGCGCCCGCCACT
PflSBW25_PfliD       AGGAGATTGGACATGGACATGAGTGTAAAGCTGAAC-CAGTCTTATCC------GCCGGT
                       *        ****** ***** ** *******     **  *  *       **

PpuF1_PfliD          CGCCCCGCCGAACAGGC-----TGGCGAA-AAACCGGTTGCTCGCGCTGAGGAGGCTTCG
PprPF5_PfliD         AAGCCGGT------GACGCCTGTCGCTGACAAGCCCGTGGAGAAGCCTCGAGAGGT----
PsyB728A_PfliD       CAACCGGCAGGCCAGGCCGTTGTGGCCGCCCCGCCGGTTGACAAGCCTGCCGACGCGAAG
PsyDC3000_PfliD      CAACCGGCAAGTCAGGCCCCCGTGTCGGACAAGTCTGTTGACAAGCCAGCCGACGTCAAA
Psa1448A_PfliD       CAACCGGCGAGTCAGGTGCCAGTGCCAGACAAGTCTGTTGACAAGCCTGCCGATACGCCG
PpuKT2440_PfliD      CAAGAAGCCAGGAGTGCACAGCCATCGAGCAGATCGAATGCTGAGCTACCAGACGTGCCG
PflSBW25_PfliD       TGCCCCTCAAGGCGCAC-----------------------------CTGTGCAGGTGTCC
                                                                         *

PpuF1_PfliD          CGCGGCGGCAGTGACGCGCTGGTACCCAATGCCATCAGTGCTGCCGAGCACAGCAATGAT
PprPF5_PfliD         -----------CGTTCCTGCAGTTGCCAAAACCGCATCTCAGGACAGTAAGAAAGAAGCT
PsyB728A_PfliD       CCCGTGGAGCGGGTTGCGGCCCCGGCCGAAAGTAAAGGTTCCGATCTGCATAAGGATAAT
PsyDC3000_PfliD      TCGGTGGAGCGTGTCGCTGCTCCTGAAGAAAGTAAAGGCTCGGATTTGCACAAGGATGAT
Psa1448A_PfliD       TCAGTCGAGCGTGTTGCCGCCACGGCGGAAAGTAAAGGCTCAGATCTGCACAAGGATGAC
PpuKT2440_PfliD      AC-----------------------------------------------CAAGGAAGTC
PflSBW25_PfliD       GCCGATAAGAGCGTCACTAAAGTCCAGGAAAC------CCCGGCTACAGGCAAAGA----
                                                                        *   *

PpuF1_PfliD          GCC---------GAAAAAGTGAAAAGCGCTGTTTCGGAAATTGAAAAGTTTCTCAGCGCC
PprPF5_PfliD         CGGACCGAGCAGGAAAAGTTGAAAATGGCCGTTCAGGAAATTGAAAAGTTCGTCCAGTCG
PsyB728A_PfliD       CCCAAGGACGATGCTAAAGTCAAAGCAGCGGCCGAGGATATTCAAAAATTCTTTCACGCG
PsyDC3000_PfliD      TCTCAGGACGAGGCGAAAGTCAAAGCCGCTGCAGAAGACATCCAGAAATTTTTTCACTCG
Psa1448A_PfliD       TCGCACGACGAGGCGAAAGTCAAAGCCGCGGCAGAAGACATCCAGAAATTTTTTCACTCG
PpuKT2440_PfliD      C------AGCGGCCGGAACTCGAGAAAGCGGTCAGCGACATACAAGAATACGTAAAAGCT
PflSBW25_PfliD       --GCCTGAGCGTGCGGACCTGGAAAAAGCAGTCACCGATATTCGCGAATTTGTACAAGCG
                                     *  *  *    ** *     ** **     * *   *     *

PpuF1_PfliD          ACCCGACGCAACCTGGAATTCTCCACGGACGAAGAATCCGGAAAGATCGTAGTTAAGGTC
PprPF5_PfliD         GTCAAGCGCAATCTAGAGTTCTCGATCGACGAGCCTTCCGGAAAGGTCGTGGTCAAAGTC
PsyB728A_PfliD       GTGAAGCGCAATCTTGAGTTCTCGATTGACGAAGATTCTGGCAAAGTCATTGTCAAAGTG
PsyDC3000_PfliD      GTGAAGCGCAATCTCGAGTTTTCGATTGATGAAGCTTCTGGCAAAGTGATCGTCAAGGTC
Psa1448A_PfliD       GTCAAACGCAATCTCGAGTTTTCGATTGATGAAGCTTCTGGCAAAGTGATCGTCAAGGTC
PpuKT2440_PfliD      TCGCAGCGGCAGTTGGATTTTTCCGTGGACGATTCCACGGGGACCCTGGTGGTCAAGGTT
PflSBW25_PfliD       TCCGAGCGTAAGCTGGACTTTTCCATTGATGACTCCACCCATCGCGTGGTGGTCAAGGTT
                           **  *  * ** ** **    ** **     *        *  * ** ** **

PpuF1_PfliD          ATCGCCAGTGAAACCGGCGAGTTGATCCGCCAGTTGCCTTCTGAAGAGGCATTGCGGATT
PprPF5_PfliD         ATTGCCAGTGATTCGGGTGAAGTGGTTCGCCAGATCCCCAATGAAGAAGTGCTTAAACTG
PsyB728A_PfliD       ATTGCAAGCGACTCCGGTGAAGTAGTAAGACAGATTCCCAATGCAGAAATCCTGAAACTG
PsyDC3000_PfliD      ATTGCCAGTGACTCGGGTGAAGTGGTAAGACAAATTCCCAATGCAGAAATCCTGAAACTG
Psa1448A_PfliD       ATTGCCAGTGACTCGGGTGAAGTGGTAAGACAAATTCCCAATGCAGAAATCCTGAAACTG
PpuKT2440_PfliD      ATTGCCACCGGTAGTGGCGAAGTCATTCGACAGCTTCCAAGTGAGGCAGCCCTGAGCCTG
PflSBW25_PfliD       ATCGCCACCGACAGCGGTGAAGTGATTCGCCAGATTCCGTCGGAAACGGCACTGAAGCTG
                     ** ** *  *     ** **   *  *  * **  * **    *         *     *

PpuF1_PfliD          GCGCATAACCTGAGCGATGTAAACAGTATTTTGTTCGACGCCAAGGTTTGATCGGCGG--
PprPF5_PfliD         GCCAATAGCTTGAATGATGCGAGTAGTCTGTTGTTCAGTGCCAAGGCCTGACAGCTGGCA
PsyB728A_PfliD       GCCGATAGCTTGAGTGATGCGAACAGTTTGTTGTTCCGCGCCAAGGCCTGATCGCCGGTA
PsyDC3000_PfliD      GCCGATAGCCTGAGTGATGCGAACAGCTTGTTGTTCCGCGCCAGAGCGTGATCGCCGGTA
Psa1448A_PfliD       GCCGATAGCCTGAGTGATGCGAACAGCTTGTTGTTCCGCGCCAAGGCCTGATTGTCAGTA
PpuKT2440_PfliD      GCTCGCAGTCTGGCGGAGGGTGATGGTTTTTTACTTGATGACAATGTGTAAT-GTTGGCA
PflSBW25_PfliD       GCACAGAACCTTTCCAGCGCAAGCCACCTGTTGTTTGACGACAAGGTCTGA--GCTGGCA
                     **    *   *       *         * **   *    * **  *  * *  *   *

PpuF1_PfliD          -AGAATGTGCTAAAGCAGGAATCACGTCGGGCTTGCAGGAGCAGGCCC--TGACGGTC--
PprPF5_PfliD         TGAAAATTGTTGCTGGTTACGTTTGAGCGCTGTTGTGGTCAAAAGACTGGCGGCACAC-T
PsyB728A_PfliD       CCGCATGTGTTGTCGGGTCATTCGCC-TTTTAAAAGGGCGGTTAACCCTGCGGCATTCTT
PsyDC3000_PfliD      CGGTACTTGTTGCAGGTTGCCTGATG-CGTTGACGCGACAGAAAAACCGGCGACATTCTT
Psa1448A_PfliD       CGGTATTTGTTGCCGTTTTGCGAGAC-CTGTGAAAAGGCGGTGAAACCGGCGACATTCTT
PpuKT2440_PfliD      TGAGTCTTGACTCGGATTGTGTCT---CGCTCATTCATC-GTCAAGCAATTGGCATTCGC
PflSBW25_PfliD       TGAAAATTGTTGC----TGTCATGTCTTGAGGCTTCGTCAGTCAAGATAACGGCAGCCAC
                            **                                          * *   *
```

```
PpuF1_PfliD           -AACAAAGAGGGATAACTGT
PprPF5_PfliD          GAA--GGGAGAGACAC----
PsyB728A_PfliD        GAA--GGGAGAAGCACC---
PsyDC3000_PfliD       GAA--GGGAGAAGCAAA---
Psa1448A_PfliD        GAA--GGGAGAAGCAA----
PpuKT2440_PfliD       TGATCAGG-GATATAAAGCA
PflSBW25_PfliD        CGAACAGGAGAAACGAAG--
                        *    * *
```

**P*fliC***

**MUSCLE alignment (https://www.ebi.ac.uk/Tools/msa/muscle/)**

**CLUSTAL multiple sequence alignment by MUSCLE (3.8)**

```
PprPF5_PfliC        ---------------------------------CAGCTAGAGCCTATGTTTT---CCCGA
PsyB728A_PfliC      ------------------------------------------------------------
PsyDC3000_PfliC     ------------------------------------------------------------
Psa1448A_PfliC      ------------------------------------------------------------
PpuF1_PfliC         ACATCGACCTGTACTGCCTGCATCAGGGCAGCGCGGCGATCGTGGATGCCGTTTCGTCGC
PpuKT2440_PfliC     ------------------------------------------------------------
PflSBW25_PfliC      -------TGCCTTCTGCATTCACCAGGGTAGTGCGGCCATTGTCGACGCCGT---GGCGC

PprPF5_PfliC        GCC-------AGAGAAGTCGCCGATCTGTTTCGTCTGGGGCGTGATGTCGAGGCGGTGGG
PsyB728A_PfliC      ----------AGACAAACCCGAG---AAGTTTCTCAAGGACATG--GTCGAAACCGGCAA
PsyDC3000_PfliC     ----------AGACAAGCCCGAG---AAGTTTCTCAAGGACATG--GTCGAAACGGGTAA
Psa1448A_PfliC      ----------AGACAAACCCGAG---AAATTTCTCAAGGACATG--GTTGAAACGGGTAA
PpuF1_PfliC         GCTTCGATGAAGGCGAGCACAAGCAGAGGTTCGTCAAGGATATG--GTCGAAACGGGCAA
PpuKT2440_PfliC     ------------------------------------------------------------
PflSBW25_PfliC      GTCGGTTTGAAGACGCTCCGGTGGACAAGTTCATCAAGGACATG--GTCGAGACCGGCAA

PprPF5_PfliC        TGTCATG--GTTGAGCTGTTCGATCCCATCCAGTCGTTGGTGGATGGAGCTGCGCTGCCT
PsyB728A_PfliC      CACCGTGTCATCCAG-----------CATTCCGCTGCTGATCGAGAAGCATGTGCTCGGT
PsyDC3000_PfliC     CACGGTGTCTTCCAG-----------TATTCCATTGCTGATTGAAAAGCATGTGCTTAAC
Psa1448A_PfliC      CACCGTGTCGTCGAG-----------CATTCCATTGCTGATTGAAAAGCACGTACTCAAC
PpuF1_PfliC         TACCGTGTCCTCGAG-----------CATACCGCTGCTGCTGGAAAAGCACGTGCTTGGC
PpuKT2440_PfliC     -------------------------------CGCTGCTGCTGCAGAAGCACATGCTCGAT
PflSBW25_PfliC      CACCGTTTCATCGAG-----------CATTCCGTTGCTGCTGGAAAAACACGTGATGGAC
                                                       * ** *  *           *

PprPF5_PfliC        GTCCAGCAGGATTGGGC-GCAATTGCTGACGCTGATGCTGGCGTGCCAGGAAGGGCAGAA
PsyB728A_PfliC      TCTTCATGGAAGCGGGTCGCCTTGAGCGGTTTTGGTGTGGGGTTGTCC------------
PsyDC3000_PfliC     TCTGACTGGAGTCGGGTTGCGCTGAGCGGTTTCGGTGTAGGCCTTTCC------------
Psa1448A_PfliC      TCTTCGTGGCATCGAGTAGCACTGAGCGGTTTTGGCGTGGGGCTTTCC------------
PpuF1_PfliC         TCCGGCTGCAAGCGCGTGGCGCTGAGTGGCTTCGGGGTGGGCTTGTCG------------
PpuKT2440_PfliC     GGCAGCTGGAAGCGCGTGGCGATCAGTGGCTTTGGCGTGGGCCTGTCG------------
PflSBW25_PfliC      GCCACCTGGAAGCGCGTGGCCATCAGTGGTTTTGGTGTGGGCCTGTCG------------
                                 * *  **  *    *     *  *  **  *  *

PprPF5_PfliC        CTGGTTGGGTTTGGCCGACTATCTGGAATACGAGCTGCTGGACTGGTTGCAGGCCGAGTT
PsyB728A_PfliC      -----TGGGGCTCTGCAATTATCTA---TAAAGACTGAG---TTCACCTCGCCTTACAAA
PsyDC3000_PfliC     -----TGGGGTTCTGCAATTATCTA---TAAAGCCTGAA----TCACCGCAGGCTAAAAA
Psa1448A_PfliC      -----TGGGGTTCTGCGATTATCTA---TAAAGCCTGAA----TCGCCGCAGGCTAAAAA
PpuF1_PfliC         -----TGGGGCTCGGCCATTATCGAACGCTGCGACTGAC-CGTTCGTCGCCTTC----AA
PpuKT2440_PfliC     -----TGGGGCTCGGCCATCCTGTA---CCGCGACTGA-----TCCCAGTCAGCCCCTAA
PflSBW25_PfliC      -----TGGGGTTCGGCGATTCTTTA---TCGCCCTTGATGCTTTAATCGCTTCAATAAAA
                         ****  *   * *   *             **      *

PprPF5_PfliC        TGGTGGATGAGCGGTGCGGCAGGCTGGGCGAGGGTTTTTGGT------------------
PsyB728A_PfliC      GAACGCCTG---CGGATAAAACCCCA---GGCGTTCTTAAAT------------------
PsyDC3000_PfliC     GAGCGCCCG---TGGCTAGTTCATCG---GGCGCTTTTTTTT------------------
Psa1448A_PfliC      CAGCGCCCG---TGACTAGTTCGCCG---GGCGCTTTTTTTT------------------
PpuF1_PfliC         CAACGCCGT---------CATCGTCTGACGGCGTTTTACTTT------------------
PpuKT2440_PfliC     AAGCGCCCGGC-------GGCCACCGCCCGGCGCTTTTTTTTCTGCCGAAAATAAGCTTC
PflSBW25_PfliC      AAACGCCGATGATCGAAAGATCATC----GGCGTTTTTTATT------------------
                        *                        *  * * *    *

PprPF5_PfliC        ------------------------------------------------------------
PsyB728A_PfliC      ----------------------------------------------TAAATAGTGATCTA
PsyDC3000_PfliC     ----------------------------------------------GACTAATCGATTCA
Psa1448A_PfliC      ----------------------------------------------GAATAATCGATTCT
PpuF1_PfliC         ------------------------------------------------------------
PpuKT2440_PfliC     TGCCTAACGCAGCCCATTGTAGGAGCGGGTTCACCCGCGAACACCGGCAAAGCCGGTGCC
PflSBW25_PfliC      ------------------------------------------------------------

PprPF5_PfliC        ------------------------------------------------------------
PsyB728A_PfliC      CGTCACA-CTTTGAACTTC--GTGGCGCAATGCGCTC---------------GCCTGCCG
PsyDC3000_PfliC     TTGCACATTTTTAAACTTGGACGTGTGTCATCTGCGCGTGGCAAATGA------------
Psa1448A_PfliC      GGTCACATTTTTGAACTTTTACAGTGGCCTGCTCTTTTTGGGACGTGA------------
PpuF1_PfliC         ------------------------------------------------------------
PpuKT2440_PfliC     ACCCATCGCGTCTCCTTCTTCGCAGCCTCACCCGCTCCACCGACGGGATGAGGCATTCAG
PflSBW25_PfliC      ------------------------------------------------------------

PprPF5_PfliC        -----------------GGGTATTTTCTGGCGTCAG-------------TTTTTCGTTCG
PsyB728A_PfliC      CC-ACTTCTGACTGCCGTGAATTCCCCTCTTTCCCG-------------CCGTGGGTTTA
PsyDC3000_PfliC     ---AGTCCAGGCAGC----AGTTCTCCTCGTGTCAC---------------ATGGGTTCG
Psa1448A_PfliC      ---AGTTCAGACAAC----GGGTTTTCTCGCGTCAT---------------GTTGGTTTG
PpuF1_PfliC         ----------------------CTATCCTGCATCAA-----------AATGGCGCCTGTA
PpuKT2440_PfliC     CCCAGTTCAAACAGT---------TTTTGACGCCAA----------------------TA
PflSBW25_PfliC      ---------------------------TGGCGCCAGAAAACCCAATGAAATCGGGGTTTC
                                                     *
```

```
PprPF5_PfliC          GTGTCGGCTTAACCCCTTGTTTTCCAAGGGATGGCCGGGTGATGGCA-AATATTTTTGAA
PsyB728A_PfliC        ----CGCCCCAAACCCTTGTTTTATAAGGGGTGTCACGATGATGGCA-AAATATTAAAAA
PsyDC3000_PfliC       GTTTCGCCTTAACCCCTTGTTTTATAAGGGGTGGAGGAATGATGGCA-AAATATTCAAAA
Psa1448A_PfliC        GTTTCGTCTCAAACCCTTGTTTTATAAGGGGTACAGGCATGATGGCA-AAATATTCAAAA
PpuF1_PfliC           AATTGGCGTCAACTCCTTGATTTCATTGGACTGGCACGGTGCCAGCA-ATTTTTTTTGAA
PpuKT2440_PfliC       AACCGCCCTTAACCCCTTGATTTCTCAGGGCTGGCCCCATGCCAGTATTTTTTTTTTGAA
PflSBW25_PfliC        GAGCCTGCGTAAACCCTTGAATTACAAGGGGTGGCACGACGCCAGTA-AATATTTTCAAA
                                **  *****  **    **  *        *   * *      **   **

PprPF5_PfliC          AAAGCCCTAAAGCAACTTGCATTACCGACGATAACTATTACGAAGGTTCTCTAGGCCATT
PsyB728A_PfliC        AAAACACTCAAGCAACCTGCCATCGCGACGATAACTATTACGAAGGTTCTCTAGG-CA-T
PsyDC3000_PfliC       AAAACGCTCAAGCAACCTGCCATCGCGACGATAACTATTACGAAGGTTCTCTAGG-CA-C
Psa1448A_PfliC        AAAACACTCAAGCAACCTGCCATCGCGACGATAACTATTACGAAGGTTCTCTAGG-CA-C
PpuF1_PfliC           ATTGCGCTAAAGCAAGACGCCCTGACGACGATAACTATTACGAAGGTTCTCTGTGCCA-C
PpuKT2440_PfliC       AAAACCCTCAAGCAACCCGCGCACCCGACGATAACCATTACGAAGGTTCTCTAGGCCA-C
PflSBW25_PfliC        AAAATCCTCAAGCAACCGGCTACCACGACGATAACTATTACGTAGGTTCTC-AGGCCA-C
                      *     ** ******   **     ********** ****** ********   * **

PprPF5_PfliC          ACCCGGCG-GTTGCCAGGGCCGGAAGC--CGCAGTACCC-AACCAACGAGGAATTCGTC
PsyB728A_PfliC        ACCCGGCG-CTTGCAGGGGCCGGAAGCCACGTAGTACCAAAACCACCGAGGAATTCATC
PsyDC3000_PfliC       ACCCGGCG-CTTGCAGGGGCCGGAAGCCACGTAGTACCA-AACCAACGAGGAATTCATC
Psa1448A_PfliC        ACCCGGCG-CTTGCAGGGGCCGGAAGCCACGTAGTACCA-AACCAACGAGGAATTCATC
PpuF1_PfliC           CTCCGGCGAATTGCCAAGGCCGGAAGC--CGTAGTACCCAATCCAACGAGGAATTCGTC
PpuKT2440_PfliC       ACCCGGCT-GTCGCCAGGGCCGGAAGC--CGCAGTATCC-ATACAACGAGGAATTCGTC
PflSBW25_PfliC        ACCCGGCG-GTTGCCAGGGCCGGAAGC--CGCAGTATCC-ATCCAACGAGGATTTCGTC
                        *****   * **   **********  ** **** *  *  ** ****** *** **
```

**P*flgF***

**MUSCLE alignment (https://www.ebi.ac.uk/Tools/msa/muscle/)**

**CLUSTAL multiple sequence alignment by MUSCLE (3.8)**

```
PpuKT2440_PflgF       ----------------------GCGCAGAGCAACTACCAGGCCAATGCCAAGACCATTTC
PpuF1_PflgF           ------------------------------------------------------------
PsyB728A_PflgF        ------------------------------------------------------------
PsyDC3000_PflgF       ------------------------------------------------------------
Psa1448A_PflgF        ------------------------------------------------------------
PflSBW25_PflgF        ------------------------------------------------------------
PprPF5_PflgF          GCGGCAACTGGAGAACGTGCTGTTTCAGGCGGTTTCCCTGTGCGATGGCGGTACGGTCAA

PpuKT2440_PflgF       CACTGAAAGCACCATCATG-CAGACCATCATTCA-GATGACCTGATGATTGTATTG----
PpuF1_PflgF           ---------CACCGTGATG-CAGACCATCATTCA-GATGACTTGATCGTCAT--------
PsyB728A_PflgF        ------------------G-CAGACGCTGATTCA-GTCGAC---CTGATCAGATCGAGCA
PsyDC3000_PflgF       ------------------G-CAGACGCTGATTCA-GTCAAC---CTGATCAGATTGAGCA
Psa1448A_PflgF        ------------------G-CAGACGCTGATTCA-GTCGAC---CTGATCAGATTGAGCA
PflSBW25_PflgF        --------------------GAAGCGAT--------------------------------
PprPF5_PflgF          GACCGAGCACATCCGGCTGCCGGACTATGGTGCGCGCCAGCCGCTTGGCGATTTCTCCCT
                                              *  *

PpuKT2440_PflgF       ---------------------GCTTTGGTGTTTGTAATTGGGGCCGCAAAGCGCCAAGAA
PpuF1_PflgF           --------------------CGTTCGAGTGAGTGCAGGCGAACCCCTTTCTCGTGAAGGG
PsyB728A_PflgF        GTCGGCTTTGATGAAGTGGCCTTTCAGGCGTTTCCAGCCTGGGCCC-----------GGG
PsyDC3000_PflgF       GTCGGATTTGACGAAGTGACCTTTCAGGCGGGTCCAGCCCGGCCCC-----------GGG
Psa1448A_PflgF        GTCGGATTTGATGAAGTGACCTTTCAGCCGTGTCCAGCCAGGCCCG-----------GGG
PflSBW25_PflgF        ---------------------TGTCGGGCGGTTCGAGAAAGCG--------------GTG
PprPF5_PflgF          GGAGGGTGGCCTGGAACAGATTGTCGGGCGCTTTGAAAAGGCC--------------GTG
                                             *     *  *  *                     *

PpuKT2440_PflgF       TGTGAAGGTTCAC--GCGAATGAAGCGGCTGTCGTT--------------ACTGGCCTTG
PpuF1_PflgF           GTTCGTTGTTTTTGCAGGAGGGGTGATGAAAAGGCT--GTTTGTCGGAATGCTGCTCGCC
PsyB728A_PflgF        CT----GGCCTGGGCGCAGACGATCCGGCTGTCGCCCTGCCATTTGTAATAGTGGGCGGG
PsyDC3000_PflgF       CT----GGTCTGGGCGCAGACAATCCGGCTGTCGCCCTGCCAGCGGTAATAGTGCGCAGG
Psa1448A_PflgF        CT----GGTCTGGGCGCAGACAATCCGGCTGTCGCCCTGCCATTTGTAATAGTGGGCGGG
PflSBW25_PflgF        CTCGAAGCCTTATATGCCGAGCATCC--CAGCAGCC--GGCAGATGGGCAAGCGTCTGGG
PprPF5_PflgF          CTCGAGAGTCTGTACGCCGAGCACCC--CAGTAGCC--GTCAGCTGGGCAAGCGGCTGGG
                                                       *                   *

PpuKT2440_PflgF       TTCCCCTTGCTGGTGCTGGTTCTGCCTGCCCCCGCTGCCCCCGCCCCGTTCTATCAATG-
PpuF1_PflgF           TGCTGTCT---TGGGCAGGT---------GCAGGCGGCGCCAGCGCCATACTGGCGATG-
PsyB728A_PflgF        ATCGGCCTGAGAGAGCAGGGCAAGGGTCATCAGGCT-CGCGCTCGCGATACCGATGATGC
PsyDC3000_PflgF       GGCGGCCTGGCAGAGCAGGGCGAAGGTCATCAGGCT-CGCGGTCGCGACGCCGACAACGC
Psa1448A_PflgF        GGCGGCCTGGGCGAGGAGGGCGAGGGTCATCAGGCT-CGCGCTGGCGATGCCGGCAATGC
PflSBW25_PflgF        GGTGTCCC------ACACCACCATCGCTAACAAGCTGCGCGACTACGAAATTCTCAAGGC
PprPF5_PflgF          GGTTTCCC------ACACCACCATCGCCAACAAACTGCGCGAATACGAGGTCGGCAA---
                                                    *   *  * *     *          *

PpuKT2440_PflgF       -------GCAAAGCCTGGTCACCG--------------------------GCCGCTACCT
PpuF1_PflgF           -------GGAAAGCCTGGTAGATG--------------------------GGCGTCTGGT
PsyB728A_PflgF        ACGCACGAAAAGGCAT----GACG--------------------------GTTCCCGTTC
PsyDC3000_PflgF       ACACACGAAAAAGCAT----GACG--------------------------GTTCCCGTTC
Psa1448A_PflgF        ATGCACGAAAAAGCAT----GAGG--------------------------GTTCCCGTTC
PflSBW25_PflgF        TAATAAATAAATCCTTTGGGAAGGGGCTTGCTGTGGTGAGTGGGGCTTGTGCCCCCATTC
PprPF5_PflgF          --------GAATGCC--GGCGACA--------------------------ATCCCTAGGC
                               **  *

PpuKT2440_PflgF       GTG--------TTCGGCCCACAACCCGGGCAAAGGCTG--GGTGC----GCCACTCTGGC
PpuF1_PflgF           GTG--------TTCGCAGTGGTCGCCAGGTGAAGGCTGGAGAC------GCTTTGCCGGG
PsyB728A_PflgF        AGG--------GTTGCGGTGTCGGCAAGGTTATCACTGTCGATCC----GGTTCTCGGCG
PsyDC3000_PflgF       AGG--------GTTGCGGTGCCTGCAAGATTATCACTGTCGATTA----GCCTGTTAGCG
Psa1448A_PflgF        AGG--------GCTGTGGTGTCGGCAAGATTATCACTGTCGATCC----GCTTGTCAGCC
PflSBW25_PflgF        AAGGGGGCTGCTTCGCAGCCCGGCGCAGGGCAAGCCTGCTCACCACGGGGCAAGCCTGCT
PprPF5_PflgF          TTG------------------------------------------------TATCCGGGC
                        *                                                      *

PpuKT2440_PflgF       CCCTACAACAACGCCGC--------------CTGCCGCGCCCATTGAGCAAGC-------
PpuF1_PflgF           CCCTACAACAATGCGCGTTGCCGTGAC----CTGCCGTAGGCACAGGTTTGCCTGCAAAT
PsyB728A_PflgF        CAGTTCGATGAAGTTTTT-------------TTGGCGGTGTTTCCGGTCTG---------
PsyDC3000_PflgF       CGTATCGATGAAGTTTTT-------------TTGGCGGTGTGTCCTGCTCG---------
Psa1448A_PflgF        CATACCAATGAAGTTTTT-------------TTGGCGGTGCTCGCTGCTTG---------
PflSBW25_PflgF        CACCACAGTAAACCCCTCCCACATTCGATCTTTGTCGCTTTCAAGGACGGG---------
PprPF5_PflgF          CGTTAC-------------------------TTGCCGGT---------------------
                      *    *                          ** **

PpuKT2440_PflgF       -------CCTTGCCGCAGGCGGCAAACATCCGCCGACAAACTGAC---------------
PpuF1_PflgF           GAAACCTTTTTGCCGCCAACGGCAAAACAGCGCCGGATAACCGGCGGCCGT---------
PsyB728A_PflgF        -------GCAAGCGGCAGGCGGCAGTCCACCGCCGCTATT--GGCTGACGC--------G
PsyDC3000_PflgF       -------ATGATCAGCAGGCGGCAGTCCGCCGCCGCTATT--GGATGGCGC--------T
Psa1448A_PflgF        -------ATGATCGGCAGGCGGCAGTCCTCCGCCGCTATT--GGGTGGCGC--------T
PflSBW25_PflgF        -------GCTTGCCGCAGGCGGCAGTAGTCCGCCGTTTTTTCGACTCTCAC--------T
PprPF5_PflgF          -----------------AGCGGCATAAGACCGCCGTTTTTCCGTCTTCTGCCAACTCTGT
                                         *****      *****       *
```

```
PpuKT2440_PflgF      ---------GGTTATTGCCCCTTGCCGCGCTGAAAGCCCCGTAAACCGGGTGTTT-----
PpuF1_PflgF          -CGCTGCGCGGGCTTTCTCAACTGCCCGTATTCA-------------TGGGCTTTTTCCT
PsyB728A_PflgF       CGCTCCTCAGGCCGTCGGCGAATGCCGGTTTTTA---CTGGCTAATCAGGCATTTGTATC
PsyDC3000_PflgF      TTGCCTGAAGGTGATCGTCGCGTGCCGGTTTTTG---CTGGTTAATCAGGCATTTGTATC
Psa1448A_PflgF       TTGCCCGAAGGGCATCATCGAATGCCGGTTTTTG---CTGGTTAATCAGGCATTTGTATC
PflSBW25_PflgF       CAATCCTGAAATCTTCATACCTCGCCGCAAACCC---CCGCCCCGCCTGCACTTCATGCT
PprPF5_PflgF         TTTTTCCTCTGTAATCAGGCAAAGCC-----------TTGCCCCGTCTGGGGATCACCAC
                                   *        ***                      *    *

PpuKT2440_PflgF      -------CAGACTTGGTTCAATTATTGCTTGGAACCTGCC----CAGTGACAGCGCGGCA
PpuF1_PflgF          GCTTTTGGCAAGTTGGTTCGGAACTTGCTTTAGCCCCTGTAGAGCAACGGCGA-ACGGCA
PsyB728A_PflgF       GAGCCTGACAAGTTGGTTCAAAAATTGCTTTAACCCTTTCAGAACAGCTACAG-GCGGCA
PsyDC3000_PflgF      GAGGCTGACAAGTTGGTTCAAAAATTGCTTTAACCCTTTCAGAACAGCTACAG-GCGGCA
Psa1448A_PflgF       GAGCCCGACAAGTTGGTTCAAAAATTGCTTTAACCCTTTCAGAACAGCTACAG-GCGGCA
PflSBW25_PflgF       CTTTTT-AAAAGTTGGTTCGCAAATTGCTTAAGCCTCATCAGTACAGCGGTGG-GCGGCA
PprPF5_PflgF         TGCGTGAAAAAGTTGGTGCGCTTATTGCTTATGGCTTCGCAGTACAGCGGTGG-GCGGCA
                               * ***** *     ******    *         **         *****

PpuKT2440_PflgF      GCTACGCG--------CAGAGGAGAAACT
PpuF1_PflgF          AACGGCCGCCAGTGCAATGGAGGATGATT
PsyB728A_PflgF       AGCGTCTGTCG-----GAGGAGGAAGACT
PsyDC3000_PflgF      AGCGTCTGTCG-----GAGGAGGAAGACT
Psa1448A_PflgF       AGCGTCTGTCG-----GAGGAGGAAGACT
PflSBW25_PflgF       AGCGTCCGTCAG---AAAGAGGAAAGAGC
PprPF5_PflgF         AACGTCCGGCAG---ACAGAGGAAAGACT
                            *          *  *    *
```

**P*flgB***

**MUSCLE alignment (https://www.ebi.ac.uk/Tools/msa/muscle/)**

**CLUSTAL multiple sequence alignment by MUSCLE (3.8)**

```
PprPF5_PflgB         ACTTCGACATGGACGGCGAGATCGACGGCAAGCCGTTCAGCGACTCCTTCGAGCTGCCGC
PsyDC3000_PflgB      ------------------------------------------------------------
PsyB728A_PflgB       ------------------------------------------------------------
Psa1448A_PflgB       ------------------------------------------------------------
PflSBW25_PflgB       ACATGGTGTTTTGCCGCAACGT---------GCTGATCTACTTCTCGGCCGAAGTGAAGA
PpuKT2440_PflgB      ------------GTCGCAATGT---------GCTGATCTATTTCTCGGCGCAGGTGAAGA
PpuF1_PflgB          ------------------------------------------------------------

PprPF5_PflgB         GCGATACCGCCTTCAACTTCGCCAGCGATGCCACGCGGGTCGCCCA--------------
PsyDC3000_PflgB      ----------------CGATAACCGCATTGCCGTGCGGTATGCCTA--------------
PsyB728A_PflgB       --------------------AACCGCATCGCCGTGCGCTATGCCTA--------------
Psa1448A_PflgB       ---------------------ACCGCATTGCCGTGCGCTATGCCTA--------------
PflSBW25_PflgB       AAGACATCCTGCTGCGCATTCACGGCACGCTCAAGCGCGGCGGCTATCTGTTCCTCGGAG
PpuKT2440_PflgB      AGGACATCCTGCTGCGCATTCACAGTACCCTCAAGCCCGGTGGCTACCTGTTCCTCGGCG
PpuF1_PflgB          ------------------------------------------------------------

PprPF5_PflgB         ----------GAAGCACGGCCTG-----------------------------CACCCGAA
PsyDC3000_PflgB      ------------TGAATGGCACGATGA-------------------------CTCGGGCA
PsyB728A_PflgB       ------------CGAATGGCATGACGA-------------------------CTCCGGCA
Psa1448A_PflgB       ------------TGAATGGCATGACGA-------------------------CTCTGGCA
PflSBW25_PflgB       CTTCCGAAGCGTTGAATGGCCTGCCGGACCATTACCAGATGGTGCAGTGCAGCCCTGGGA
PpuKT2440_PflgB      CCTCGGAAGCGCTGAACGGCTTGCCGGACCATTACCAGATGGTGCAGTGCAGCCCGGGGA
PpuF1_PflgB          ----------------------------------------------------CCCGGGGA
                                                                         * *  * *

PprPF5_PflgB         GTTCGGCGCGATCACCCGG----GTGCACAAGGAATACGATGC--------GATGTTCGA
PsyDC3000_PflgB      ACTGGTTCCGTTCCTATGGTAATGAAAACTGGGAGTTCGCGGCG-------AACGGCCTC
PsyB728A_PflgB       ACTGGTTCCGTTCGTATGGCAATGAAAACTGGGAGTTCGCTGCA-------AACGGCCTC
Psa1448A_PflgB       ACTGGTTCCGTTCGTATGGCAACGAAAACTGGGAGTTCGCTGCA-------AACGGCCTC
PflSBW25_PflgB       -------TCATTTACCAGGCCAAGTAA------------------------GACGGTCTC
PpuKT2440_PflgB      -------TCATCTACCAGGCCAAGTAAGCACGAAGGCCGGTATCGAGAGATGCCGGCCTT
PpuF1_PflgB          -------TCATCTACCAGGCCAAGTAAGCACGAAGGCCGGTATCGAGAGATGCCGGTCTT
                             *        **    *                              *  *

PprPF5_PflgB         GGACATT----------CGCGCCAAGC----------TGCATGCCCATCCCGGCGAACCA
PsyDC3000_PflgB      ATGCAAC----------GGCGCTTTGCCTGCATCAATGATTTGCCGATCGCCGAGTCCGA
PsyB728A_PflgB       ATGCAAC----------GGCGCTTTGCCTGCATCAATGATTTGCCGATCGCCGAGTCCGA
Psa1448A_PflgB       ATGCAGC----------GGCGCTTTGCCTGCATCAACGATTTGCCGATCGCCGAGTCCGA
PflSBW25_PflgB       ACTCGATCAATGTGGGAGGGGGCTTGC--------------TCCCGATGGCGGTGGTTCA
PpuKT2440_PflgB      TT------------------TGTATGCCTATCCGCTGTTCCTGCCGAGCTCCCCGTAGGA
PpuF1_PflgB          TT------------------TATTTGCCTATCCGCTGTTCCTGCCGAGCTCCCCGTAGGA
                                              **              * ** *   *   *    *

PprPF5_PflgB         GTGGATCTCGAGCGGATCATTCGCCACGAGTGAGCCGTGTCCCCGGCGGCC---------
PsyDC3000_PflgB      GCGCAAAT-------ACCGCTGGCCACTGGGGCGGC---GACCAGATGATCATCCTG---
PsyB728A_PflgB       GCGCAAAT-------ACCATTGGCCGCTGGGGCGCC---GACCGGATGATCACCCGG---
Psa1448A_PflgB       GCGCAAAT-------ACCATTGGCCACTGGGGCGCC---GACCTGATGACCACCCTG---
PflSBW25_PflgB       GTTGATAT----------AGCTATCACTGACACACC------------------------
PpuKT2440_PflgB      GCGGGTTT-------AC------CCGCGAAGAGGCC--GGCCCAGG--------------
PpuF1_PflgB          GCGGGTTT-------AC------CCGCGAAGAGGCC--GGCCCTGCTGGTCAATCTCCAC
                     *      *                * *        *

PprPF5_PflgB         ---------------------------GGAAGCATGGGAAAGGCCC--------------
PsyDC3000_PflgB      ---------------------------GCTTGAGTGAGCTGGGGCTCT------------
PsyB728A_PflgB       ---------------------------GGTTGAGCGAGCTTGGCCTCT------------
Psa1448A_PflgB       ---------------------------GGTTGAGTGAGCTTGGCCTCT------------
PflSBW25_PflgB       ---------------------------GTCATCGGGAGCAAGCCCCCT------------
PpuKT2440_PflgB      ------------------------------------------------------------
PpuF1_PflgB          CCGCCGTTCACCGTTGCTGCCGCTACCGTATTCGCGGGTAAACCCGCTCCTACGACGGAT

PprPF5_PflgB         ------------------------------------------------------------
PsyDC3000_PflgB      -------------------------------------------------GA-TCA-TCGC
PsyB728A_PflgB       -------------------------------------------------GA-TCAATCGT
Psa1448A_PflgB       -------------------------------------------------GATTCAGTCGT
PflSBW25_PflgB       ------------------------------------------------------------
PpuKT2440_PflgB      ------------------------------------------------------------
PpuF1_PflgB          CGGGGGTGGGCATCGGTCGGTGGTTGGGCACAGGCCGCGTGTAGGAGCGGGTTTACCCGC

PprPF5_PflgB         -------------GATATCGACGATATCGGGCCTTTTTTCGTTTGC--------------
PsyDC3000_PflgB      TACTCGCCTTTTATATGAAGAGACCTCAGGGTCTCTTTTTTATTGG--GTTCGACTCA--
PsyB728A_PflgB       CA---------------AAGAGACCTTCGGGTCTCTTTTTTATTGGGTGTCCGGTTGAGG
Psa1448A_PflgB       CA---------------AAGAGACCTTCGGGTCTCTTTTTTATTGAGTGTCCGAATGAGC
PflSBW25_PflgB       ---------------------------CCCACATTTTTTTTGCGGC--------------
PpuKT2440_PflgB      -----------------AAAACACTGACGTCAATTTTTTGTTTTGC--------------
PpuF1_PflgB          GAAGAGGCCCGCACAGGAAAACACTGACGTCAATTTTTTGTTTTGC--------------
                                                      * ****     *
```

```
PprPF5_PflgB         --------------------------------------------CGGAGAAGCCGTTCCA
PsyDC3000_PflgB      --------AGGCGC------TGCCACACGGCAAACCCATACTGGCAAAAAAACGGCACCC
PsyB728A_PflgB       CAATGATGAAGAGCCACGATTGCCACGCGGCAAGTCGATACTGGCAAAAAAACGGCACCC
Psa1448A_PflgB       CATTAATGAAGTGCCGCCATTGTCAGGCGGCAAGCCGATATTGGCAAAAAAACGGCACCC
PflSBW25_PflgB       --------------------------------------AAATTTCAAGAGAGCGGCAACT
PpuKT2440_PflgB      --------------------------------------CGCTTT--------TGCCGCCC
PpuF1_PflgB          --------------------------------------CGCTTT--------TGCCGCCC
                                                                               * 

PprPF5_PflgB         GGGGCTTGCCACCCTTGCCGGC--AAGCGGAAGCGGCTTGCCGCTTGCCGCTTCCGGGTT
PsyDC3000_PflgB      ACC--TTGCCGCTTTGCCTCTCCACAGCGGAAATGCCTTGCCGCTT-----TTCTGGCAT
PsyB728A_PflgB       GCC--TTGCCGCTTTACCCCTCTGCAGCGGAAATGCCTTGCCGCTT-----TTCTGGCAT
Psa1448A_PflgB       CTC--TTGCCGCTTTCTTCTCCCACAGCGGAAATGCCTTGCCGCTT-----TTCTGGCAT
PflSBW25_PflgB       CCCCATTGCCGCTTTACTGGCGCCACGCGGAAATCGCTTGCCGCTT-----TTCTGGCAT
PpuKT2440_PflgB      ATCAATTGCCGCTTTTGTCACCTCAGGCGGAAGGCCTTTGCCGCTT-----TTCTGGCAT
PpuF1_PflgB          ATCAATTGCCGCTTTTGTCACCTCAGGCGGAAGGCCTTTGCCGCTT-----TTCTGGCAT
                          ***** *  *           ******     *********     *** **  *

PprPF5_PflgB         ------------ACCCTTAAATAAACTCAAGTTATTGAATATAAAGGTTTTTAATAATTT
PsyDC3000_PflgB      TGCC--------GCCTCTGCGCTGTCTGTAACCCTCGCAATCATTGGCTTTTGA-AACAT
PsyB728A_PflgB       TGCC--------GCCTCGCCGCCGCCTGTGGACCCTGCAAACATTGGCTTTTAC-AACAT
Psa1448A_PflgB       CGCT--------GCCTCGCCGCTGGCTGCGGCCCCCGCAATCATTGGCTTTTGC-AACAT
PflSBW25_PflgB       CGTC--------GCCGGCAACCGCCTCGCAAGCCCTTGATATACGGGCCTCCACCAGATT
PpuKT2440_PflgB      CAATCTGACAGTACCGCGCCGC------AAAACCCCATAAATACGGGCTTTCCAGTGGTT
PpuF1_PflgB          CAACCTGACAGTACTGCGCCGC------AATCCCCCCTAAACACGGGCTTTTCAGTGGTT
                                  *                        *   *  **  *         *

PprPF5_PflgB         GGCACGGGCCTTGCTGAAGTGTTGGCGACGAAAGCCA---CACCGGCAAACAGGTTCGCA
PsyDC3000_PflgB      GGCACGCCGATTGCTTTGTCAGTAACAACAGAA-TCAGGTCAACCGGCAAAGGTTTCCC-
PsyB728A_PflgB       GGCACGCCGATTGCTTTGTCAGTAACAACAGAA-TCAGGTCAACCGGCAAAGGTTTCCC-
Psa1448A_PflgB       GGCACGCTGATTGCTTTGTCAGTAACAACAGAA-TCAGGTCAACCGGCAAAGGTTTCCC-
PflSBW25_PflgB       GGCATGCACCTTGCTATAACCCTGTTAACGAAAAGCAGGTCAGCC--TAAAGGTTTCC--
PpuKT2440_PflgB      GGCACAGCCCTTGCTATGCCTTGCTCAACGAAATTCCGGTCAACCTCTGAAGGTTTCCCT
PpuF1_PflgB          GGCACAGCCCTTGCTATACCTTGCTCAACGAAATTCCGGTCAACCTCTGAAGGTTTCCCC
                     ****      *****            **  **  *    ** *     *  * ***

PprPF5_PflgB         GCC
PsyDC3000_PflgB      GAC
PsyB728A_PflgB       GAA
Psa1448A_PflgB       GAA
PflSBW25_PflgB       GCC
PpuKT2440_PflgB      GAC
PpuF1_PflgB          GAC
                     *
```

**P*flgA***

**MUSCLE alignment (https://www.ebi.ac.uk/Tools/msa/muscle/)**

**CLUSTAL multiple sequence alignment by MUSCLE (3.8)**

```
PsyDC3000_PflgA_start_codon_corr     ---TTATCGACCCGGGTTACTGCGGTCAGGTAATGATCGCGCCCGGTGCCCTTGGGTGGT
PsyB728A_PflgA                       --GTTATCAACCCGGGTAACGGCGGTCAGGTAATGATCGCGCCCGGTGCCCTTGGGTGGT
Psa1448A_PflgA                       ---TTATCAACCCGGGTGACGGCGGTCAGGTAATGATCGCGCCCGGTCCCCTTGGGTGGT
PpuKT2440_PflgA                      -----GTCCACCCGGGTGACAGCGGTCAGGTAGTGATCACGGCCGGTACCCTTGGGTGGC
PpuF1_PflgA                          -----GTCAACCCGGGTGACCGCGGTGAGGTAGTGATCACGGCCGGTACCCTTGGGTGGC
PprPF5_PflgA_start_codon_corr        TGGTTGTCCACCCGAGTGACCGCCGTCAGGTAGTGATCGCGTCCGGTCCCCTTGGCGGC
PflSBW25_PflgA                       -------CGACCCGCGTCACCGCCGTGAGGTAGTGATCGCGACCGGTGCCCTTGGGTGGC
                                            * ***** ** ** ** ** ***** ***** ** ***** ******** **

PsyDC3000_PflgA_start_codon_corr     GGATGAATCTCTTCCCAGTTCATGTTGACGATGCGTTCGACAGAATGCACCAGAAACCCC
PsyB728A_PflgA                       GGATGAATCTCTTCCCAGTTCATGTTGACGATGCGTTCGACAGAATGCACCAGAAAGCCC
Psa1448A_PflgA                       GGATGAATCTCTTCCCAGTTCATGTTGACGATGCGTTCTACAGAATGCACCAGAAACCCC
PpuKT2440_PflgA                      GGATGGATCTCTTCCCAGTTCATGTTGACGATGCGCTCCACCGAATGCACCAGGAAGCCT
PpuF1_PflgA                          GGATGGATCTCTTCCCAGTTCATGTTGACGATGCGCTCCACCGAATGCACCAGGAAACCT
PprPF5_PflgA_start_codon_corr        GGATGGATCTCTTCCCAGTTCATGTTGACGATGCGCTCCACCGAGCGCACCAGGAAACCC
PflSBW25_PflgA                       GGATGGATCTCCTCCCAGTTCATGTTGACGATGCGCTCCACCGAGCGCACCAGGAAACCC
                                     ***** ***** *********************** ** ** **  ******* ** **

PsyDC3000_PflgA_start_codon_corr     TGAACCTTGGTGTTGTACTCGGTAATGATGACGAAGGAATTGA------CCAGCGAAATC
PsyB728A_PflgA                       TGAACCTTGGTGTTGTACTCGGTAATGATGACGAAGGAATTAA------CCAGCGAAATC
Psa1448A_PflgA                       TGAACCTTGGTGTTGTACTCAGTAATGATGACGAAGGAATTGA------CCAACGAAATC
PpuKT2440_PflgA                      TGGGTCTTGGTGTTGTACTCGGTGATGATCACGAAGCTGTTGCGTGTGTCTTCCTGCAAC
PpuF1_PflgA                          TGGGTCTTGGTGTTGTACTCGGTGATGATCACGAAGCTGTTGCGTGTGTCTTCCTGCAGC
PprPF5_PflgA_start_codon_corr        TGGGTCTTGGTGTTGTACTCAGTGATGATGACGAACGGATTGCTCTGGTCCTG------C
PflSBW25_PflgA                       TGGGTCTTGGTGTTGTATTCGGTAATGATCACGAACGGGTTCTCGCGGTCCTG------C
                                     **   ************ ** ** ***** *****    **        *         *

PsyDC3000_PflgA_start_codon_corr     ATCCCTGTGGAGCCCGTCGCCATTGCCAGATCCATAATCGGAATGGTACCGCCACGAATA
PsyB728A_PflgA                       ATCCCTGTGGAACCTGTCGCCATTGCCAGATCCATGATCGGGATCGTGCCGCCACGAATA
Psa1448A_PflgA                       ATGCCCGTGGAACCCGTCGCCATCGCCAGATCCATGATCGGGATCGTACCGCCACGAATA
PpuKT2440_PflgA                      GGCCGC---AACCCGGTGGCCATCGACAAGTCGAGGATCGGGATGGTCGCCCCGCGAATA
PpuF1_PflgA                          GGCCGC---AACCCGGTGGCCATCGACAAGTCGAGGATCGGGATGGTCGCCCCGCGAATA
PprPF5_PflgA_start_codon_corr        AGGCCACCAGAACCCGTAGCCATTGCCAGATCCAGGATCGGAATGGTTGCCCCTCGTATA
PflSBW25_PflgA                       AACCCAGCAGAACCGGTGGCCAGGGCCAAATCAAGAATCGGAATCGTCGCGCCACGGATG
                                        *      * ** ** ****  * **  ** *  ***** ** **  * ** ** **

PsyDC3000_PflgA_start_codon_corr     TTGGCCACGCCACGAACGATCTTGCTCGATTTGGGCATGATGGTGAGTTTCGGGCATTGC
PsyB728A_PflgA                       TTGGCCACGCCCCGAACGATCTTGCTCGATTTGGGCATGATGGTGAGTTTCGGGCATTGC
Psa1448A_PflgA                       TTGGCCACACCCCGGACGATCTTGCTCGATTTGGGCATGATGGTGAGTTTCGGGCATTGC
PpuKT2440_PflgA                      TTGGCAACGCCACGCACCACAGGGTGCGCCTTGGGCAGCAGGGTCAGCGCAGGGCATTGC
PpuF1_PflgA                          TTGGCAACGCCACGCACCACAGGGTGCGCCTTGGGCAGCAGGGTCAGCGCAGGGCATTGC
PprPF5_PflgA_start_codon_corr        TTCGCGACACCGCACACCACAGGACTGGATTTGGGCATCAGGGTCAGCTTAGGGCATTGC
PflSBW25_PflgA                       TTAGCGACGCCACAGACCACAGGGCTGGACTTGGGCATGATGGTCAACTTGGGGCACTGC
                                     ** ** ** ** *  ** *        *  *******  * *** *     ***** ***

PsyDC3000_PflgA_start_codon_corr     AGTACTTCTTTTACCTTGAAGACGTTGATCCCGTAGAGCTGCTTGCCGTCGAGACGGAAC
PsyB728A_PflgA                       AGTACTTCTTTTACCTTGAAGACGTTGATCCCATAGAGCTGCTTGCCGTCGAGACGGAAC
Psa1448A_PflgA                       AGTACTTCTTTTACCTTGAAGACGTTGATCCCGTAGAGCTGCTTGCCGTCGAGACGGAAC
PpuKT2440_PflgA                      AGTACTTCCCTGACCTTGAACACATTGATGCCGTAAAGCTGTTCGCCATTGAGGCGGAAC
PpuF1_PflgA                          AGTACTTCCCTGACCTTGAACACATTGATGCCGTAAAGCTGTTCGCCATTGAGGCGGAAC
PprPF5_PflgA_start_codon_corr        AGCACCTCGCGAACCTTGAATACGTTGATCCCGTAGAGCTGCCCACCGTCCAGACGGAAC
PflSBW25_PflgA                       AGCACTTCCCGCACCTTGAACACATTGATGCCATACAGCTGCTTGCCATCCAGGCGAAAA
                                     ** ** **    ******** ** ***** ** ** *****    ** *  ** ** **

PsyDC3000_PflgA_start_codon_corr     AAGAGCAGCTCCAGGCGATTCTGACCCACCAGCTGAGTTCGCTGGTTTACCGAATCCATT
PsyB728A_PflgA                       AAGAGCAGCTCCAGGCGATTCTGACCCACCAGCTGAGTTCGCTGGTTTACCGAATCCATT
Psa1448A_PflgA                       AAGAGCAGCTCCAGGCGATTCTGACCCACCAGCTGAGTTCGCTGGTTTACCGAATCCATT
PpuKT2440_PflgA                      AGCAGCAATTCCAGGCGATTCTGCCCCACCAGCTGCGTGCGCTGGTTGACCGAATCCATA
PpuF1_PflgA                          AGCAGCAATTCCAGGCGATTCTGCCCCACCAGCTGCGTGCGCTGGTTGACCGAATCCATA
PprPF5_PflgA_start_codon_corr        AGCAACAACTCTAGTCGATTCTGCCCAACCAATTGCGTGCGCTGGTTCACCGAATCCATT
PflSBW25_PflgA                       AGCAACAACTCCAGGCGATTCTGCCCTACCAGCTGTGTGCGCTGGTTTACTGAATCCATT
                                     *  * **  ** ** ******** ** ****  ** ** ******** ** ********

PsyDC3000_PflgA_start_codon_corr     ACACCTGCCATGCCGAACCCCCTTACCAAAATGACC-------GCCACCCAGGCACCAGA
PsyB728A_PflgA                       ACACCTGCCATGCCGAACCCCCTTA-CAAAATGACC-------GCCGCCCAGACACCAGA
Psa1448A_PflgA                       ACACCTGCCATGCCGAACCCCCTTACCAAAATGACC-------GCCGCCCAGACACCAGA
PpuKT2440_PflgA                      ACACCCGCCATGCC-AGACTCCTTAACGTTTCAGCCTTTCGTTGCAACTCAGGTAGCGAG
PpuF1_PflgA                          ACACCCGCCATGCC-AGACTCCTTAACGTTTCAGCCTTTCGTTGCAACTCAGGTAGCGAG
PprPF5_PflgA_start_codon_corr        ACACCAGCCATGCCCAGACTCCCCAA------AGCTTTAAGATCCGACGCACGCTC---A
PflSBW25_PflgA                       ACTCCTGCCATGCCTTGACTCCTACGCTAAAACCTTTGGGGGTGCGATGCGCACCC---A
                                     ** ** ********    * **                      *    *

PsyDC3000_PflgA_start_codon_corr     CCAACAAGCGGCACGGGCTTTGCTATTTGTTACGTC
PsyB728A_PflgA                       TCAACAAGCGGCACGGGCTTTGCTATTTATTACTTC
Psa1448A_PflgA                       TCAACAAGCGGCACGGGCTTTGCTATTTATTACCTC
PpuKT2440_PflgA                      T-------CGGCACGGGCTTTGCTTTTTTGAGTGTC
PpuF1_PflgA                          T-------CGGCACGGGCTTTGCTTTTTTGAGTGTC
PprPF5_PflgA_start_codon_corr        TTGCTAAACGGCACGGCGCTTGCTTTTTAACTGTT-
PflSBW25_PflgA                       ACACTAAACGGCACGAGCTTTGCTATTTATTGGTTT
                                             *******    ***** ***      *
```

**P*flgM***

**MUSCLE alignment (https://www.ebi.ac.uk/Tools/msa/muscle/)**

**CLUSTAL multiple sequence alignment by MUSCLE (3.8)**

```
PflSBW25_PflgM_start_codon_corre      GGCCAATGAAACGCACCGGCATCATTGGTTTCGAAGATGTGGTACTGCGCGAACGCGACA
PprPF5_PflgM                          ----------GCGCGCCGGCATCATCGAACCGGAAGACGTGGTGCTGCGCGAACGGGACA
PpuKT2440_PflgM                       -----------------------GTCGGCGAAGGTGACGTGGCCCTGCGCGAGCGCGATG
PpuF1_PflgM                           -----------------------GTCGGCGAAGGTGACGTGGCCCTGCGCGAGCGCGATG
PsyB728A_PflgM                        --------------------ATCGTCACCGAAGAAGACGTTGCGATGCGCGAGCGGGATG
PsyDC3000_PflgM_start_codon_corr      -------------------CATCGTGACTGAAGAAGATATCGCGATGCGTGAGCGGGATG
Psa1448A_PflgM                        --------------------ATCGTTACTGAGCAAGACGTTGCGCTGCGTGAGCGCGACG
                                                              *          **  * *   **** ** ** **

PflSBW25_PflgM_start_codon_corre      TCAGCATGATCAGCCAGGGCTACCTGACCTCCCTCGACCAAGCGGTCGGACAGAAATTAA
PprPF5_PflgM                          TCAGCCAGAACAATCAGGGCTACCTCACCTCAGTGGACCAAGCCATTGGTCAGAAACTTG
PpuKT2440_PflgM                       TCGGCACCCTGGGCCAGGGCTTTCTGACCGAGCTGGACCAGGCGGTGGGCATGAAGATGC
PpuF1_PflgM                           TCGGCACCTTGGGCCAGGGCTTTCTGACCGAGCTGGACCAGGCGGTGGGCATGAAGATGC
PsyB728A_PflgM                        TCAGCAGCCTGGGTCAGGGTTTCCTGGCATCGCTGGATCAGGCAGTCGGTCAGAAGGTTG
PsyDC3000_PflgM_start_codon_corr      TCAGCAGCCTGGGCCAGGGTTTCCTGGCATCGCTTGATCAGGCAGTCGGTCAGAAGGTTG
Psa1448A_PflgM                        TGAGCAGCCTGGGCCAGGGATTCCTGGCCTCCCTTGATCAGGCAGTCGGTCAGACGGTTG
                                      *  **         ***** *  **  *     * ** ** **  * **   **   *

PflSBW25_PflgM_start_codon_corre      CCCGACCAGTGGTAACAGACCAAGTCATCACCCTGGTGCATCTTGAACAGGCAGAGGTGA
PprPF5_PflgM                          TCCGACCAATGGTCGCCGACCAGCTCGTTACCCTGGCCCATCTGGAGCAGGCGGAAGTCG
PpuKT2440_PflgM                       TGCGCCCCACGGTGATCGACCAGGTGCTCACCCCGCAACATCTGGAACAGGCCGAGGTGG
PpuF1_PflgM                           TGCGCCCCACGGTGATCGACCAGGTGCTCACCCCGCAACACCTGGAACAGGCCGAGGTGG
PsyB728A_PflgM                        TCCGACAGATGGTCATCGACCAGGTCATCACCCCGGTGGCGCTTGAGCAGCCACAAATGA
PsyDC3000_PflgM_start_codon_corr      TCCGACAAATGGTCATCGATCAGGTCATCACCCCCGTGGCGCTTGAACAGCCCCAGATGA
Psa1448A_PflgM                        TCCGACAAATGGTCATCGACCAGGTCGTTACCCCGGTGGCGCTGGAACAGCCACAAATGA
                                        ** *    ***    ** **  *  * ****        ** ** *** *  *  *

PflSBW25_PflgM_start_codon_corre      TTCGCAAGGGCGATCAGGTGGTGATTTCCGCCAGCAGTGGCGGGCTGAACGTAAAAATGC
PprPF5_PflgM                          TGCGCAAGGGCGATCAAGTGGTGATATCGGCACGCAGCGGCACGTTGAATGTACGGATGC
PpuKT2440_PflgM                       TGCGCAAGGGTGACCAGGTGGTGATCATTGCCCGCAGTGGCAGCCTGAGTGTGCGCATGC
PpuF1_PflgM                           TGCGCAAGGGTGACCAGGTGGTGATCATCGCCCGCAGTGGCAGCCTGAGTGTGCGCATGC
PsyB728A_PflgM                        TTCACAAGGGCGATCAGGTGGTGATCATCGCCCGCAGCGGCTCGCTGGCGGTGCGCATGC
PsyDC3000_PflgM_start_codon_corr      TTCACAAGGGCGATCAGGTGGTGATCATCGCCCGCAGCGGCTCGCTGGCGGTACGGATGC
Psa1448A_PflgM                        TTCACAAGGGCGACCAGGTGGTCATCATCGCCCGCAGCGGTTCGCTGGCGGTGCGCATGC
                                      * * ****** ** ** ***** **    **  **** **     **   **    ****

PflSBW25_PflgM_start_codon_corre      CGGGGGAGGCACTGTCCAACGGTGGCATGAGTGAGCAGATACGGGTCAAGAACCTCAACT
PprPF5_PflgM                          CGGGCGAGGCGCTGTCCAACGGCGGCATGAGCGAACAGATCCGGGTCAAGAATCTCAATT
PpuKT2440_PflgM                       CGGGCGAAGCCTTGAGCAAGGGCGGCCTGAGCGAGCAGATTCGGGTACGCAACCTCAATT
PpuF1_PflgM                           CGGGCGAAGCCCTGAGCAAGGGCGGCCTGAGCGAGCAGATCCGGGTACGCAACCTCAATT
PsyB728A_PflgM                        CCGGCGAAGCGATGTCCGATGGAGGCTTCAACGAACAGATACGGGTGAAAAACCTCAATT
PsyDC3000_PflgM_start_codon_corr      CCGGCGAAGCGATGTCCGACGGCGGTTTCAACGAACAGATACGGGTGAAAAACCTTAACT
Psa1448A_PflgM                        CCGGCGAAGCGATGTCCGACGGCGGTTTCAACGAACAGATACGGGTGAAAAACCTTAATT
                                      * ** ** **  **  * * ** **  * *  ** ***** *****    ** ** ** *

PflSBW25_PflgM_start_codon_corre      CCAACCGCGTCATCAAGGCGCGAGTGACGGCCCCCGGGCAAGTCGAGGTTGCTTTATAGA
PprPF5_PflgM                          CGCAACGAGTGATCAAGGCCCGCATCACGGCGCCAGGGCAAGTGGAAGTGGCCATGTAGA
PpuKT2440_PflgM                       CCAAACGCGTGGTCAAGGCCAGGGTGACCGGCCCGGGCCAGGTCGAGGTCAGTATGTAGG
PpuF1_PflgM                           CAAAACGCGTGGTCAAGGCCAGGGTGACCGGCCCGGGCCAGGTCGAGGTCAGTATGTAGG
PsyB728A_PflgM                        CCCAGCGGGTGATCAAGGCAAGTGTCACCGGCCCAGGGCAGGTGGAAGTCGCTATGTAGT
PsyDC3000_PflgM_start_codon_corr      CCCAACGGGTGATCAAAGCGAATGTCACCGGTCCAGGACAGGTAGAAGTGGCCATGTAAT
Psa1448A_PflgM                        CCCAGCGGGTGATCAAAGCGAATGTCACCGGCCCAGGGCAGGTGGAAGTGGCCATGTAAT
                                      *  * ** **  **** **     * ** *  ** ** ** ** ** **     * **

PflSBW25_PflgM_start_codon_corre      GT--------------GCTGGCAGAGGACGCTGGGTTTTTCTACACTGTGCACGAGATAT
PprPF5_PflgM                          TA--------------GCTGGCGCTCAAGCCGTGCGTTCCCTAAACTGTG-G-CCGATA-
PpuKT2440_PflgM                       CTAGCCACCAGATTGCGCTGGCAGTGAGGAACCGCTTTTCCTAAACTGTG-TCAGAACA-
PpuF1_PflgM                           CTGGCCACCAGATTGCGCTGGCAGTGAGGAACCGCTTTTCCTAAACTGTG-TCAGAACA-
PsyB728A_PflgM                        GCAT--------TGATGGTGGCGCGAAGGGTGTGTTTTTTTTAGACTGCG-G-GGAATG-
PsyDC3000_PflgM_start_codon_corr      GCAT--------TGAGGCTGGCGCAGAGGGTGTGTTTTTTTTAGACTGCG-G-GGAATG-
Psa1448A_PflgM                        GCAT--------TGATGCTGGCGCAGAGGGTGTGTTTTTTTTAGACTGCG-G-GGAATA-
                                                      * ****           *  **   ** **** *      *

PflSBW25_PflgM_start_codon_corre      TCCGTGCGCAG------------ATTTATTGT------------CATTCGCGCCTAAAGT
PprPF5_PflgM                          AAGGGTCGCAGATACCTGACCTGGTTTGTCGA------------CAATCGAGCCTAAAGT
PpuKT2440_PflgM                       CGGGTTCGCGG------------GCTTGCCGACAGGCGGCAATGCATTTGTGCCTAAAGT
PpuF1_PflgM                           CGGGTTCGCGG------------GCTTGCCGACAGGCGGCAATGCATTTGTGCCTAAAGT
PsyB728A_PflgM                        CCCGACGGCAAATGTC-------GTCTATTGGGAAGCTTTCTCACAATCA-GCCTAAAGT
PsyDC3000_PflgM_start_codon_corr      CCCGACGGTGAAAAGC-------ATCTATTGAGAAGCTTTATCACAATCG-GCCTAAAGT
Psa1448A_PflgM                        TCCGACGGTGAAAAGC-------GTCTATTGGGAAGCTTTCTCACAATCG-GCCTAAAGT
                                         *   *                  *   *             ** *   *********

PflSBW25_PflgM_start_codon_corre      TCGTCCGGGTATTGCCGAAAACATGGCAAGCGTCCAAATACCCAGAGGTTTTT-TGATC
PprPF5_PflgM                          TTTTCAGGGTATGGCCGAAAACATGGCAAGCGTCCAAATACCCAGAGGTTTACTTCATC
PpuKT2440_PflgM                       TTATATCGGGTTGGCCGAAAACAAGGCAAGCGTCCAAATACCCAGAGGTTTCT--GATC
PpuF1_PflgM                           TTATATCGGGTTGGCCGAAAACAAGGCAAGCGTCCAAATACCCAGAGGTTTCT--GATC
PsyB728A_PflgM                        TTGATGGGGGTTGGCCGAAAACAAGGCAAGCGTCCAAACACCCAGAGGTTTTTTCAATC
PsyDC3000_PflgM_start_codon_corr      TTAATTGGGGTTGGCCGAAAACAAGGCAAGCGTCCAAACACCCAGAGGTTTTT-CAATC
Psa1448A_PflgM                        TTGATAGGGGTTGGCCGAAAACAAGGCAAGCGTCCAAACACCCAGAGGTTTTTTCAATC
                                      *      **  * ********** ************** ************     ***
```

**P*flgZ***

**MUSCLE alignment (https://www.ebi.ac.uk/Tools/msa/muscle/)**

**CLUSTAL multiple sequence alignment by MUSCLE (3.8)**

```
PpuKT2440_PflgZ                       GATGCAGGAATTGCTCGACCTTCTGCACAAGGAAGCAATCGCCCTGCATGGCCGCGACAT
PpuF1_PflgZ                           GATGCAGGAACTGCTCGACCTTCTGCAGAAGGAGTCTGTAGCCCTGCATGGCCGCGATAT
PsyDC3000_PflgZ_start_codon_corr      ---CCGGCAGCTTCTTGAGCTGTTGCAGGCTGAATCACTGATTCTTCACGGCCGCGACAT
PsyB728A_PflgZ                        ---CCGACAGCTTCTTGAGCTGTTGCAGGCCGAATCACTGATTCTTCACGGCCGCGACAT
Psa1448A_PflgZ                        ---CCGACAGCTTCTAGAGCTGCTGCAGGCCGAATCCCTGATTCTTCACGGCCGCGACAT
PflSBW25_PflgZ_start_codon_corre      ----------------GAGCTACTCAGGGAAGAATCCCTGGCCCTCTACGGCCGCGACAT
PprPF5_PflgZ                          -----------------AGCTGCTACGCGCCGAGTCTATCGCCCTCCACGGCCGCGACAT
                                                       * **  *       **  *  *    **  * ******** **

PpuKT2440_PflgZ                       GCCGCTGCTGGAGCAAATCCTGGCGCGCAAGCAGTCGTTGATCATCTTGCTCGAGCAGCA
PpuF1_PflgZ                           GGCGCCACTGGAAAATATCCTGGCCCGCAAGCAGTCTTTGATCGTGTTGCTGGAGCAGCA
PsyDC3000_PflgZ_start_codon_corr      GAGCGAGATGGAACAGGTGCTGGCGCAGAAACAGGCGTTGGTCATCCTGCTGGATCAGCA
PsyB728A_PflgZ                        GGGCGAGATGGAGCAGGTGCTGGCGCAGAAACAGGCGCTGGTCATTCTTCTCGACCAGCA
Psa1448A_PflgZ                        GGGCGAGATGGAGCAGGTGCTGGCGCAGAAACAGGCGCTGGTCATTCTTCTCGATCAACA
PflSBW25_PflgZ_start_codon_corre      GCCGCTGCTGGAAGAAATTCTGGCGCGCAAGCAATCGCTGATTGTCCTGCTGGAACAGCA
PprPF5_PflgZ                          GCCGCTGCTGGAAGATATCCTGGCGCAGAAGCAAGCCTTGGTCATCTTGCTGGAACAGCA
                                      *       ****  *  * ***** *  ** **  *  ** *  *  * ** ** ** **

PpuKT2440_PflgZ                       TGGCCAGCGCCGCAGCCAATTGCTCGGCAGCCTGGGCCTGAGTGCCGACCGCACCGGTGT
PpuF1_PflgZ                           AGGCCTGCGGCGCAACAACCTGCTCACCAGCCTTGGCCTGAGCGCAGATCGCGCCGGCGT
PsyDC3000_PflgZ_start_codon_corr      CGGCCGCAAACGTAGCCAGGTGCTCGCCGGTATGGGGCTGCCCGCCAATCGCAGTGGTCT
PsyB728A_PflgZ                        TGGTCGCAAACGTAGCCAGGTGCTTGCGGGCATGGGGCTGCCTGCCAATCGCAGTGGTCT
Psa1448A_PflgZ                        TGGGCGCAAACGTAGCCAGGTGCTTGCGGGCATGGGGCTGCCAGCCAATCGCAGTGGTCT
PflSBW25_PflgZ_start_codon_corre      TGGCAAAAAACGCAGCCAGATCCTCGTCAGCCTTGGCCTGCCCGCAGACCATGACGGCCT
PprPF5_PflgZ                          TGGCCGTAAGCGCACGCAAATTCTTGCCAGCCTGAACCTGCCGACCAACCGCGAGGGGCT
                                       **       ** *   *  * **     *  *    ***    *  * *     **  *

PpuKT2440_PflgZ                       GCAGGCTGTAGCTGCGCAGTCACCCCATGGTGATGTGATGCTGCAGCGGCTCGACATGTT
PpuF1_PflgZ                           GGAGGCTGTGGCGGCCCAATCGCCTAACGGTGAATTGATCCTGCAACAACTGGACGTGAT
PsyDC3000_PflgZ_start_codon_corr      GCAGCAACTGGCCAGCCAGTCCGAGGTCGGCGAGCAATTGCTGACGAGCAGCGATGAGCT
PsyB728A_PflgZ                        CCAGCAACTGGCCAGCCAGTCGGATATCGGTGAGCAACTGCTGCTGGCCGCCGACGAACT
Psa1448A_PflgZ                        CCAGCAACTGGCCAGCCAGTCGGATGTCGGCGAGCAATTGCTGATGGCCGGTGACGAACT
PflSBW25_PflgZ_start_codon_corre      GGCGCAGCTGGCCAGCCACTCTTCGGTCGGCGATCAATTACTGGCCCAGAGCAAAGAACT
PprPF5_PflgZ                          CGAGCAACTGGCCAGCCACTCCAGCCTGGGCGCGCAATTGCTCGAACAGAGCAATGTCCT
                                         *    * **    ** **       ** *      * **           *     *

PpuKT2440_PflgZ                       GTCGCAACTGATGGACGACTGCCAGCAGGTCAACCAGACCAATGGCCGGATCATCCAGGT
PpuF1_PflgZ                           CAGCCAGTTGATGCAAGCCTGCCAACAGCTTAACGAAACCAATGGCCGGATCATCCAGGT
PsyDC3000_PflgZ_start_codon_corr      GAATGCGCTGATCAATGAATGCCAGACGCTGAACGATCAGAATGGTAGTCTGATCCAGTT
PsyB728A_PflgZ                        GAACACGTTGATCAACGAATGCCAGACGCTCAACGAGCGCAATGGTAGTCTGATCCAGTT
Psa1448A_PflgZ                        GAATGCGTTGATCAATGAATGCCAGACGCTGAACGAGCAGAATGGTAGTCTGATCCAGCT
PflSBW25_PflgZ_start_codon_corre      CAATCAATTGCTCGCCCAGTGCCAGGAAGCCAACCTGCTCAACGGTCAGTCGATCCAGCT
PprPF5_PflgZ                          GACCCAATTGCTTGCCGAGTGCCAAGCTGCCAACGAGCTCAATGGTCGCTCGATCCAGAT
                                              ** *       *****       ***      ** **       ****** *

PpuKT2440_PflgZ                       GCAGCAACACGTCACCAACAACCAGATCCGCATCCTTCAGGGCGGCGATTCGCCTTCGCT
PpuF1_PflgZ                           GCAGCAACACGTCACCAACAACCAGATCCGGATCCTCATGGGCGGCGACTCGCCTTCGCT
PsyDC3000_PflgZ_start_codon_corr      GCAACAGATCAGCACCGCTCACCAGCTTCGCATTCTCAATGGCGGTGAAACGCCAACGCT
PsyB728A_PflgZ                        GCAACAGGTCAGTACCGCTCACCAGCTTCGCATCCTGAACGGTGGCGATACGCCTACACT
Psa1448A_PflgZ                        GCAACAGATCAGCACCGCTCATCAACTTCGCATACTGAACGGCGGTGAGACGCCCACGCT
PflSBW25_PflgZ_start_codon_corre      TCAGCAAGCTACGACCGCCAACCAGTTGCGTATACTTCACGGCGGAGAGCCTCCAGCGCT
PprPF5_PflgZ                          CCAACAGGCCACCACCGCCAACCAGCTGAAAATCCTCAATGGTGGCGAACCTCCTGCCCT
                                       ** **       ***    * **  *    ** **    ** ** **  * **  * **

PpuKT2440_PflgZ                       CTACGATAGCCGTGGCAGCACCTCGCCGCTGGCCAAGCCGCGCGCGCTCAGCCAAGTGTG
PpuF1_PflgZ                           CTACGACAGCCGTGGCAGCACCTCGCCGCTGGCCAAGCCGCGCGCGCTCAGCCAAGTGTG
PsyDC3000_PflgZ_start_codon_corr      CTATGATGCTCGCGGCTCGACTGCGCTTAGAGCAAAGCCACGCCCCTTGAGTCAGGCGTA
PsyB728A_PflgZ                        CTATGACAGCCGCGGATCGACTGCGCTTAGAGCAAAACCACGCCCTCTGAGCCAGGCGTA
Psa1448A_PflgZ                        CTATGACAGTCGCGGGTCAACTGCGCTTAGAGCAAAACCACGCCCTCTGAGTCAGGCGTA
PflSBW25_PflgZ_start_codon_corre      TTACAACGCTCAAGGTTCCACCTCGCGCCTGGTCAAGCCAAGCACTCGCAGCCAAGCCTG
PprPF5_PflgZ                          ATACGACGCCCGCGGCTCTACCTCAATGCTTGCGAAGCCACGCCCACTCAGTCAGGCTTA
                                       **  *    *  **    **  *       ** ** **  ** *    ** ** *  *

PpuKT2440_PflgZ                       ATTTT---------------TCTA-TCAAGGCACAGAACATAGTGGCAAAATGCCGTTAC
PpuF1_PflgZ                           ATTTT---------------TCTA-TCAAGGCACAGAACATAGTGGCAAAATGCCGTTAC
PsyDC3000_PflgZ_start_codon_corr      ACTTCC--------------TGTA-TCAAGGTTTCGTGCGTAGTGGCAGAATGCCTGTTG
PsyB728A_PflgZ                        ACTTCC--------------TGTA-TCAAGGTTTCGTGCATAGTGGCAGAATGCCTGCTG
Psa1448A_PflgZ                        ACTTCC--------------TGTA-TCAAGGTTTCGTGCGTAGTGGCAGAATGCCTGTTG
PflSBW25_PflgZ_start_codon_corre      ACGCCGGTTTAC-AGCGCG-GCTATCCAAGACGCGCCACATACTGGCAGAATGCCTGCTC
PprPF5_PflgZ                          AACCCAAAAAATGAGCGCGCGCTA-CCAAGCCGTGAAACATGCTGGCAAAATGCTGGCTC
                                      *                     **  ****        * *  ***** *****

PpuKT2440_PflgZ                       TTGCGTGTGTCGC----TTTTGCCTGGAGAACAATAAGCT
PpuF1_PflgZ                           TTGCGTGTGTCGC----TTTTGCCTGGAGAACGATAAGCT
PsyDC3000_PflgZ_start_codon_corr      TTGCGT-TGCCGTAGTATTTTGCCTGGAGATTCAAGATC-
PsyB728A_PflgZ                        TTGCGT-TGCCGTAGTATTTTGCCTGGAGATTCAAGAAC-
Psa1448A_PflgZ                        TTGCGT-TGCCGTAGTATTTTGCCTGGAGATTCAAGAAC-
PflSBW25_PflgZ_start_codon_corre      TTGCG--TGTAGTCGTATTTTGTCTGGAGATGGAAGAACC
PprPF5_PflgZ                          CTGCG--TGTAGTTGTATTTTGTCTGGAGATTGATGAACC
                                       ****  **  *     ***** *******   *  * *
```

**P*motA***

**MUSCLE alignment (https://www.ebi.ac.uk/Tools/msa/muscle/)**

**CLUSTAL multiple sequence alignment by MUSCLE (3.8)**

```
PsyDC3000_PmotA      ------------------------------------------------------------
PsyB728A_PmotA       ------------------------------------------------------------
Psa1448A_PmotA       ------------------------------------------------------------
PpuKT2440_PmotA      GCTGGCTGATCATCTGGAACTGGCACAGTACCGGCGGAATTTCCTCCACGGGCAGCGCCG
PpuF1_PmotA          GCTGGCTGATCATCTGGAACTGGCACAGTACCGGCGGAATTTCCTCCACGGGCAGCGCCG
PflSBW25_PmotA       ------------------------------------GAATCTCGTTTTCGGGCACTGCGG
PprPF5_PmotA         ------------------------------------GGATCTGCTCTTCAGGCAGCGTCG

PsyDC3000_PmotA      -------------AAGCGTTTCGGTACGCGCCAGCCCCAGACGGTTTATCGCGATTTCGA
PsyB728A_PmotA       --------------AGGACTTCGGTACGCGCCAGACCCAGGCGGTTGATGGCAATTTCAA
Psa1448A_PmotA       --------------AGGATTTCAGTACGCGCCAGCCCCAGGCGGTTGATCGCAATCTCGA
PpuKT2440_PmotA      GCAGGCGTTTGAGAAGCTGCTCGCTGCGCTCCAGGCCCAGGCGGTTGAGAGCAACCTCCA
PpuF1_PmotA          GCAGGCGTTTGAGAAGCTGCTCGCTGCGCTCCAGGCCCAGGCGGTTGAGGGCGACTTCCA
PflSBW25_PmotA       GCAAGCGCGCCAGCAGTTCTTCGGCGCGCTTGAGGCCCAGGCGGTTAAGCGCCACTTCGA
PprPF5_PmotA         GCAGTTGCTCGAGCAACTCCTCGGTACGCTTGAGGCCCAGGCGGTTGACCGCGATCTCCA
                                   *     **    ***   ** ***** ***** *  ** *  ** *

PsyDC3000_PmotA      GGCTCTCGGCCTGTTCGGTCAAGC---CATGAGTGTGATGATTGGCCTCGCGCATCACGC
PsyB728A_PmotA       GGCTCTCGGCCTGCTCGGTCAAAC---CGTGGGTGTGATGGTTGGCCTCGCGCATCACGC
Psa1448A_PmotA       GGCTCTCGGCCTGCTCGGTCAAAC---CGTGGGTGTGATGGTTGGCCTCGCGCATCACAC
PpuKT2440_PmotA      GGCTTTCGGCGGGTTCGGCGAGGCTGGCATTGGTGGGGTGATTGGCTTCGCGCATCACTG
PpuF1_PmotA          GGCTTTCGGCGGGTTCGGCAAGGCTGGCATTGGTGGGGTGATTGGCTTCGCGCATCACTG
PflSBW25_PmotA       GATTTTCTGCCGGCTCAGCCAGGCTGCCATGGGTGTGGCTGTTGGCCTCGCGCATCACGC
PprPF5_PmotA         GGTTTTCCGCCGGTTCGCTCATGCTGCCGTGGGAATGCCGGTTGGCTTCGCGGATCACGC
                     *  * ** **  * **    *  *   * *  *   *    ***** ***** *****

PsyDC3000_PmotA      TGAGCACCAGCGCAGGGCTTTCCTGCATCATTTCGGCGATTTCACGCAGCGAGCGGCGGC
PsyB728A_PmotA       TCAGCACCAGCGCAGGACTTTCCTGCATCATTTCGGCAATTTCACGCAGCGACCGACGAC
Psa1448A_PmotA       TCAATACCAGTGCGGGGCTTTCCTGCATCATTTCAGCGATTTCACGCAGCGAGCGGCGAC
PpuKT2440_PmotA      ACAGCACCAGTGCCGGGCTATCCTGCATCAGTTCGGCGATATCGCGCAGCGAGCGGCGGC
PpuF1_PmotA          ACAGCACCAGTGCCGGGCTATCCTGCATCAGTTCGGCGATATCGCGCAGCGAGCGACGGC
PflSBW25_PmotA       TGAGCACCAGCGCCGGGCTGTCTTGCATCAGCTCGGCAATATCGCGCAATGAACTGCGGC
PprPF5_PmotA         TCAAGGCCAGTGCCGGGCTGTTTTGCATCAGATCGACAATCTCCCGCAGCGAGCTGCGGT
                       *   **** ** ** ** *  *******  **  * ** ** ****  ** *  **

PsyDC3000_PmotA      TGTCGTTCAACGCTGAACGAACGTGAGCATGATTGACGGCCGGCACCGGCAGCGCGATGC
PsyB728A_PmotA       TGTCGTTCAGTGCGGCGCGGACGTGTCCATGATTGACGGCAGGGACCGGCAACGCTATGC
Psa1448A_PmotA       TGTCGTTCAATGCTGCGCGAACGTGCCCATGATTGACCGCCGGGACCGGCAACGCGATGC
PpuKT2440_PmotA      TATCGTGGATGGCAGCCATGACCCGGTCGTAGCTGTGCTTCGGCACGGGGATGCGAACAC
PpuF1_PmotA          TATCGTGGATGGCAGCCATGACCCGGTCGTAGCTGTGCTTCGGCACGGGGATGCGAACAC
PflSBW25_PmotA       TGTCGCGGATGCCCTTGCACACACGGTCGTGGCTGGCCTGTGGAACAGGCAGCAGCACAT
PprPF5_PmotA         TGTTGGCGATGGCGCGGCAGACCCGGTCATGGCTGGCCTGGGGCACCGGCAGGCGCACGC
                     * * *   *   *       **  *  * *   **      ** ** ** *     *

PsyDC3000_PmotA      CGTCAAGCTGTTTGACCCAGGCATCAAGCGTCTTGGGAACCAGATGATGTGCTACATGTG
PsyB728A_PmotA       CGTCAAGCAGTTTGACCCAGGCATCCAGCGTCTTCGGAACGAGGTGATGTGCTGCGTGTA
Psa1448A_PmotA       CGTCAAGCAATTTGACCCAGGCATCCAGCGTTTTCGGAACGAGATGATGTGCTGCGTGTG
PpuKT2440_PmotA      TCTCGAGCAGCTTTACCCAGGCCTCTAGCGTACGCGGGGC--------------------
PpuF1_PmotA          TCTCGAGCAGCTTTACCCAGGCCTCTAGCGTACGCGGGGC--------------------
PflSBW25_PmotA       CGTCCAGGCGCTTGATCCAAGCGGCGAGAG-AGGTAGGTTTAGCAGTTGGGACTGTCGTT
PprPF5_PmotA         CATTGAGGAGCTTTACCCAGGCGTCGAGAG-AGGCGGGCGGAGAATGTGGGACGCTGGTT
                       *  **    ** * *** **  * ** *      *

PsyDC3000_PmotA      CTGTCGACATTCTATGGCCTGTGCGGCTGATTATCCGGGCTGGAAGACTGCCAGTGTGCG
PsyB728A_PmotA       CTGTCGGCATTCGAGGGCCTGTGCGGCTGATTATCTGAGCTGGGAAACCGCCAGCGTACG
Psa1448A_PmotA       CTGTCGACATTCGATGGCCTGTGCGGCTGATTATCTGAGCTGGGAAACCGCCAGCGTACG
PpuKT2440_PmotA      ----------------------------------CTGAGTGGGCACCTTGGTTTCAATTG
PpuF1_PmotA          ----------------------------------CTGAGTGGGCACCTTGGTTTCAATTG
PflSBW25_PmotA       TCATTAGCC--------------------ATGATTGGGGCGCGATCATCACTTGCGGTCA
PprPF5_PmotA         TCATTAGAC--------------------ATGGTC-GGGCGCAATCATCGTCTGCTGTGA
                                                         * *

PsyDC3000_PmotA      GGGTTGGACAGCCAGTTTCCCCTTTTTATCGGACCGGGAAAAT-GGTTTTTCACAAGAAC
PsyB728A_PmotA       GGGTGCTGAGGCCATTTTCCTCCTTTTATGGAGGCGCATAAATGGGTTTTTCAAAAGAAC
Psa1448A_PmotA       GGACGCTGAGGCCATTTTCCTCCTTTTATGGAGGCGGATAAATGGGTTTTTCAAAAGAAC
PpuKT2440_PmotA      GCATTTTGACATCA---------------------GTCCGAACTGGCTTTTCGCATTGAG
PpuF1_PmotA          GCATTTTGACATCA---------------------GTCCGAACTGGCTTTTCGCATTGAG
PflSBW25_PmotA       ACACACCCGGAGCG---------------------GGCCGAACTGGCTTTTCGCCTTAAC
PprPF5_PmotA         ACACGCCCAGAGCG---------------------GGCCAAACTGGCTTTTCGCCTTAAC
                                 *                      *    **  ** *****      *

PsyDC3000_PmotA      CGGCTATAGTCTGGCCCAGTTAGGCCGATAAGGAGAAGAAGAGATTCAAGAACTTCCGGA
PsyB728A_PmotA       TGGCTATAGTCTGGCCCAGTTAGGCCGATAAGGAGAAGAAGAGATTCAACAACTTCCGGA
Psa1448A_PmotA       CGGCTATAGTCTGGCCCAGTTAGGCCGATAAGGAGAAGAAGAGATTCAAGAACTTCCGGA
PpuKT2440_PmotA      TGGCTATAGTCTGGCGCAGTTCTGCCGATAAGTAGAAGAAGAGTTTTAA-AGCTCCCGTC
PpuF1_PmotA          TGGCTATAGTCTGGCGCAGTTCTGCCGATAAGTAGAAGAAGAGTTTTAA-AGCTCCCGTC
PflSBW25_PmotA       GGGCTATAGTCTGGCGCAGTTTTGCCGATAAGAGGAACAAGAGTTTTCAGTTCCCCCCAA
PprPF5_PmotA         TGGCTATAGTCTGGCGCAGTTTTGCCGATAAGTAGAAGAAGAGTTTTAAGAACTTCCGAA
                      ************** *****  *********  *** ***** **  *   *  **
```

```
PsyDC3000_PmotA       -CTGTCCCTGAACCCGACTCAGTAAAGTATTTTTT-CCT
PsyB728A_PmotA        -CTGTCCCTGAACCCGACTCAGTAAAGTATTTTTT-CCT
Psa1448A_PmotA        -CTGTCCCTGAACCCGACTCAGTAAAGTATTTTTT-CCT
PpuKT2440_PmotA       TCTTACCCTGACCACGAC--AGCAAGTGCTTTGAA-CCT
PpuF1_PmotA           TCTTCCCCTGACCACGAC--AGCAAGTGCTTTGAA-CCT
PflSBW25_PmotA        TATGTCCATGAACCCGACGGCGCAAGTACTCATCT-CCT
PprPF5_PmotA          TATGACCTTGAACCCGACTCAATAAGTACTCCCCTACCT
                        *  ** *** * ****     **    *       ***
```
