## Supplemental Tables and Figures for "Transcriptional organization and regulation of the *Pseudomonas putida* flagellar system": Supplementary Tables and Figures_REVISED.pdf

**Supplementary Table S1. Inventory of flagellar genes *P. putida* KT2440 flagellar cluster.** *P. putida* locus tags are according to the TIGR sequencing project annotation available at the *Pseudomonas* Genome Database (pseudomonas.com). Putative functions are based on experimental evidence obtained in *P. putida* or other organisms.

| Locus tag | Gene name | Function |
|---|---|---|
| PP_4328 | *fliK2* | Protein containing FliK domain. Function unknown |
| PP_4329 | *flhX* | Protein containing FlhB domain. Auxiliary to export substrate specificity switch |
| PP_4331 | *flrD* | Ortholog of *Vibrio flrD*. Auxiliary protein to FleSR-dependent regulation |
| PP_4332 | *cheW* | Chemotaxis adaptor protein |
| PP_4333 | *parP* | Ortholog of *Vibrio parP*. Chemotaxis apparatus polar landmark protein |
| PP_4334 | *parC* | Ortholog of *Vibrio parC*. Chemotaxis apparatus polar landmark protein |
| PP_4335 | *motD* | Flagellar motor/stator protein |
| PP_4336 | *motC* | Flagellar motor/stator protein |
| PP_4337 | *cheB* | Chemotaxis protein-glutamate methylesterase |
| PP_4338 | *cheA* | Chemotaxis histidine kinase |
| PP_4339 | *cheZ* | Chemotaxis protein phosphatase |
| PP_4340 | *cheY* | Chemotaxis response regulator |
| PP_4341 | *fliA* | RNA polymerase alternative $\sigma$ factor ($\sigma^F$ or $\sigma^{28}$) |
| PP_4342 | *fleN* | Flagellar number regulator and FleQ modulator protein |
| PP_4343 | *flhF* | Flagellar polar landmark protein |
| PP_4344 | *flhA* | Flagellar secretion system export gate protein |
| PP_4352 | *flhB* | Flagellar secretion system export substrate specificity protein |
| PP_4353 | *fliR* | Flagellar secretion system export gate protein |
| PP_4354 | *fliQ* | Flagellar secretion system export gate protein |
| PP_4355 | *fliP* | Flagellar secretion system export gate protein |
| PP_4356 | *fliO* | Flagellar secretion system export gate protein |
| PP_4357 | *fliN* | Flagellar C ring/switch protein |
| PP_4358 | *fliM* | Flagellar C ring/switch protein |
| PP_4359 | *fliL* | Flagellar motor/stator-switch connector protein |
| PP_4361 | *fliK* | Flagellar hook length control protein |
| PP_4362 | *hptB* | Phosphorelay histidine phosphotransferase. Regulator of FliA activity |
| PP_4363 | *hsbR* | Histidine kinase/reponse regulator. Regulator of FliA activity |
| PP_4364 | *hsbA* | Anti-$\sigma^F$ factor antagonist |
| PP_4365 | *fliJ* | Flagellar secretion system ATPase subunit |
| PP_4366 | *fliI* | Flagellar secretion system ATPase subunit |
| PP_4367 | *fliH* | Flagellar secretion system ATPase subunit |
| PP_4368 | *fliG* | Flagellar C ring/switch protein |
| PP_4369 | *fliF* | Flagellar MS ring structural protein |
| PP_4370 | *fliE* | Flagellar hook-basal body adaptor protein |
| PP_4371 | *fleR* | Two-component system transcription activator |
| PP_4372 | *fleS* | Two-component system sensor histidine kinase |
| PP_4373 | *fleQ* | $\sigma^{54}$-dependent transcription activator |
| PP_4374 | *fliT* | Flagellar filament cap protein secretionchaperone |
| PP_4375 | *fliS* | Flagellin secretion chaperone |
| PP_4376 | *fliD* | Flagellar filament cap protein |
| PP_4377 | *flaG* | Flagellar filament length control protein |
| PP_4378 | *fliC* | Flagellin |
| PP_4380 | *flgL* | Flagellar hook-associated protein |
| PP_4381 | *flgK* | Flagellar hook-associated protein |
| PP_4382 | *flgJ* | Peptidoglycan hydrolase |
| PP_4383 | *flgI* | Flagellar P ring structural protein |
| PP_4384 | *flgH* | Flagellar L ring structural protein |
| PP_4385 | *flgG* | Flagellar rod structural protein |
| PP_4386 | *flgF* | Flagellar rod structural protein |
| PP_4388 | *flgE* | Flagellar hook structural protein |
| PP_4389 | *flgD* | Flagellar hook cap protein |
| PP_4390 | *flgC* | Flagellar rod structural protein |
| PP_4391 | *flgB* | Flagellar rod structural protein |
| PP_4392 | *cheR* | Chemotaxis methyltransferase protein |
| PP_4393 | *cheV* | Chemotaxis adaptor protein |
| PP_4394 | *flgA* | Flagellar P ring secretion chaperone |
| PP_4395 | *flgM* | Anti-$\sigma^F$ factor |
| PP_4396 | *flgN* | Flagellar hook-ssociated proteins secretion chaperone |
| PP_4397 | *flgZ* | Flagellar c-di-GMP effector and brake protein |

**Supplementary Table S2. Class II flagellar gene expression in *Escherichia coli*.** The values show β-galactosidase activity (Miller units) stationary phase cultures of *E. coli* ET8000 bearing the empty vector pSB1K3, or pMRB99, expressing *P. putida* KT2440 FleQ from its own promoter grown in LB. Also shown are the number of biological replicates (n) for each fusion and strain, the averages and standard deviations, the fold differences (ET8000/pMRB99 / ET8000/pSB1K3 ratio) in the β-galactosidase activity and the T-test *p*-values are shown. * = *p*<0.05; ** = *p*<0.01; *** = p <0.001; ns = not significant.

| | ET8000/pSB1K3 | | | ET8000/pMRB99 | | | pMRB99 vs. pSB1K3 | | |
|---|---|---|---|---|---|---|---|---|---|
| Promoter | n | β-galactosidase | | n | β-galactosidase | | Fold difference | T-test | |
| | | Average | SD | | Average | SD | | | |
| P*parC* | 4 | 247 | 34 | 4 | 6721 | 1345 | 27 | 0.002369 | ** |
| P*flhF* | 4 | 774 | 59 | 4 | 19539 | 9945 | 25 | 0.032582 | * |
| P*flhA* | 4 | 231 | 52 | 4 | 5450 | 1097 | 24 | 0.002426 | ** |
| P*fliL* | 4 | 309 | 38 | 4 | 2379 | 1016 | 8 | 0.026567 | * |
| P*fliK* | 4 | 381 | 41 | 4 | 1351 | 210 | 4 | 0.002107 | ** |
| P*fliE* | 4 | 361 | 49 | 4 | 11498 | 1109 | 32 | 0.000264 | *** |
| P*fleS* | 4 | 286 | 46 | 4 | 241 | 30 | 1 | 0.16374 | ns |
| P*fliD* | 4 | 481 | 44 | 4 | 3452 | 713 | 7 | 0.003545 | ** |
| P*flgF* | 4 | 165 | 21 | 4 | 8364 | 1130 | 51 | 0.000708 | *** |
| P*flgB* | 4 | 181 | 6 | 4 | 14664 | 3013 | 81 | 0.002388 | ** |
| P*flgA* | 4 | 160 | 32 | 4 | 11014 | 1414 | 69 | 0.000598 | *** |
| P*flgZ* | 4 | 3889 | 430 | 4 | 3407 | 717 | 1 | 0.301567 | ns |

**Supplementary Table S3. Effect of *rpoN* deletion on Class II flagellar gene expression.** The values show β-galactosidase activity (Miller units) from exponential and stationary phase cultures of *P. putida* KT2442 (wild-type) and MRB149 (Δ*rpoN*) grown in LB containing 10 mM glutamine. The number of biological replicates (n) for each fusion and strain, the averages and standard deviations, the fold differences (wild-type/Δ*rpoN* ratio) in the β-galactosidase activity and the T-test *p*-values are shown. * = *p*<0.05; ** = *p*<0.01; *** = p <0.001; ns = not significant.

| Exponential phase | | | | | | | | | |
|---|---|---|---|---|---|---|---|---|---|
| | Wild-type | | | Δ*rpoN* | | | Wild-type vs. Δ*rpoN* | | |
| Promoter | n | β-galactosidase | | n | β-galactosidase | | Fold difference | T-test | |
| | | Average | SD | | Average | SD | | | |
| P*parC* | 4 | 1690 | 410 | 4 | 185 | 100 | 9 | 0.003872 | ** |
| P*flhF* | 4 | 7511 | 1500 | 4 | 3616 | 1393 | 2 | 0.009015 | ** |
| P*flhA* | 4 | 980 | 173 | 4 | 145 | 47 | 7 | 0.001463 | ** |
| P*fliL* | 4 | 5145 | 1747 | 4 | 428 | 105 | 12 | 0.012278 | * |
| P*fliK* | 7 | 497 | 71 | 8 | 304 | 76 | 2 | 0.000214 | *** |
| P*fliE* | 4 | 6816 | 1668 | 4 | 320 | 133 | 21 | 0.004245 | ** |
| P*fleS* | 6 | 291 | 29 | 8 | 296 | 60 | 1 | 0.856722 | ns |
| P*fliD* | 6 | 899 | 148 | 7 | 523 | 61 | 2 | 0.000869 | *** |
| P*flgF* | 4 | 3112 | 921 | 4 | 177 | 60 | 18 | 0.007666 | ** |
| P*flgB* | 4 | 7239 | 334 | 4 | 152 | 62 | 48 | 0.000017 | *** |
| P*flgA* | 4 | 4783 | 1186 | 4 | 1808 | 226 | 3 | 0.013557 | * |
| P*flgZ* | 4 | 6759 | 797 | 4 | 5009 | 761 | 1 | 0.019201 | * |
| **Stationary phase** | | | | | | | | | |
| | Wild-type | | | Δ*rpoN* | | | Wild-type vs. Δ*rpoN* | | |
| Promoter | n | β-galactosidase | | n | β-galactosidase | | Fold difference | T-test | |
| | | Average | SD | | Average | SD | | | |
| P*parC* | 4 | 4185 | 356 | 4 | 548 | 140 | 8 | 0.000054 | *** |
| P*flhF* | 4 | 25924 | 1694 | 4 | 13810 | 1444 | 2 | 0.000042 | *** |
| P*flhA* | 4 | 2054 | 111 | 4 | 320 | 129 | 6 | 0.000001 | *** |
| P*fliL* | 4 | 11477 | 659 | 4 | 1482 | 420 | 8 | 0.000001 | *** |
| P*fliK* | 6 | 1747 | 176 | 6 | 728 | 215 | 2 | 0.000005 | *** |
| P*fliE* | 4 | 17830 | 701 | 4 | 1110 | 268 | 16 | 0.000002 | *** |
| P*fleS* | 6 | 1041 | 132 | 6 | 614 | 124 | 2 | 0.000185 | *** |
| P*fliD* | 6 | 2306 | 417 | 6 | 1146 | 175 | 2 | 0.000485 | *** |
| P*flgF* | 4 | 10376 | 668 | 4 | 464 | 36 | 22 | 0.000081 | *** |
| P*flgB* | 4 | 20202 | 512 | 4 | 519 | 89 | 39 | 0.000003 | *** |
| P*flgA* | 6 | 10114 | 1913 | 6 | 2005 | 426 | 5 | 0.000092 | *** |
| P*flgZ* | 6 | 16987 | 2709 | 6 | 9295 | 588 | 2 | 0.000731 | *** |

**Supplementary Table S4. Bacterial strains, plasmids and oligonucleotides used in this work.** Underlined bases indicate oligonucleotide positions that differ from the corresponding templates. [1]References list for this Table shown as Supplemental References below.

| Bacterial strain | Genotype/phenotype | Reference[1]/source |
|---|---|---|
| ***E. coli*** | | |
| DH5α | ϕ80d*lacZ*ΔM15 Δ(*lacZYA-argF*)U169 *recA*1 *endA*1 *hsdR*17 ($r_k^-$ $m_k^+$) *supE*44 *thi*-1 *gyrA relA*1 | Hanahan, 1983 |
| DH5α λ-pir | DH5α with lysogenic phage λ-pir. Host for R6K replication origin plasmids | Víctor de Lorenzo |
| ET8000 | *rbs gyrA hutCK*$^{C}$ *lacZ*::IS*1* Mucts62 | Buck *et al*., 1986 |
| ***P. putida*** | | |
| KT2440 | mt-2 *hsdR*1 ($r^-$ $m^+$). | Franklin *et al*., 1981 |
| KT2442 | mt-2 *hsdR*1 ($r^-$ $m^+$). Rif$^{r}$. | Franklin *et al*., 1981 |
| MRB34 | KT2442 *lapA*::miniTn5-Km. Cm$^{r}$ Rif$^{r}$ Km$^{r}$. | López-Sánchez *et al*., 2016 |
| MRB34/ miniTn*7*-*nahR*-P*sal*-*yhjH* | MRB34 containing the miniTn*7*-*nahR*-P*sal*-*yhjH* transposon. Cm$^{r}$ Rif$^{r}$ Km$^{r}$ Gm$^{r}$ | This work |
| MRB34/ miniTn*7*-*nahR*-P*sal*-*pleD* | MRB34 containing the miniTn*7*-*nahR*-P*sal*-*pleD* transposon. Cm$^{r}$ Rif$^{r}$ Km$^{r}$ Gm$^{r}$ | This work |
| MRB49 | KT2442 *flhF*::miniTn5-Km. Cm$^{r}$ Rif$^{r}$ Km$^{r}$. | Navarrete *et al*., 2019 |
| MRB52 | KT2442 Δ*fleQ*. Cm$^{r}$ Rif$^{r}$. | Navarrete *et al*., 2019 |
| MRB71 | KT2442 Δ*fleN*. Cm$^{r}$ Rif$^{r}$. | Navarrete *et al*., 2019 |
| MRB92 | KT2442 Δ*fleS*. Cm$^{r}$ Rif$^{r}$. | This work |
| MRB93 | KT2442 Δ*fleR*. Cm$^{r}$ Rif$^{r}$. | This work |
| MRB130 | KT2442 Δflagella. Cm$^{r}$ Rif$^{r}$. | This work |
| MRB130/ miniTn*7*-*nahR*-P*sal*-*fleQ* | MRB130 containing the miniTn*7*-*nahR*-P*sal*-*fleQ* transposon. Cm$^{r}$ Rif$^{r}$ Gm$^{r}$ | This work |
| MRB130/ miniTn*7*-*nahR*-P*sal*-*fliA* | MRB130 containing the miniTn*7*-*nahR*-P*sal*-*fliA* transposon. Cm$^{r}$ Rif$^{r}$ Gm$^{r}$ | This work |
| MRB149 | KT2442 Δ*rpoN*. Cm$^{r}$ Rif$^{r}$. | This work |

| Plasmid | Genotype/phenotype | Reference[1]/source |
|---|---|---|
| pBBR1-MCS4 | Broad host-range cloning vector. Ap$^{r}$, Mob$^{+}$ | Kovach *et al*., 1995 |
| pENTR™/D-TOPO® | Vector for directional TOPO® cloning. Km$^{r}$ | Thermo Fisher Scientific |
| pEX18-Tc | Gene replacement vector with MCS from pUC18. Tc$^{r}$ Sac$^{s}$ Mob$^{+}$ | Hoang *et al*., 1998 |
| pEMG | pJP5603 bearing a lacZα polylinker with two flanking I-SceI sites. *ori*R6K, Km$^{r}$ | Martínez-García and de Lorenzo, 2011 |
| pEMG-flagella | pEMG bearing 1.56 Kb insert with flanking regions of flagellar cluster. Km$^{r}$. | Martínez-García and de Lorenzo, 2011 |
| pFLP2 | FLP recombinase expression plasmid. Ap$^{r}$ | Hoang *et al*., 1998 |
| pMPO284 | pPS854-derived vector containing pUTminiTn*5*Km Km$^{r}$ gene flanked by the FRT sites. Ap$^{r}$ Km$^{r}$ | Jiménez-Fernández *et al*., 2014 |

| | | |
|---|---|---|
| pMRB1 | pBBR1-MCS4-derived broad host-range *gfp*mut3::*lac*Z transcriptional fusion vector. Ap$^r$ | Jiménez-Fernández *et al*., 2014 |
| pMRB2 | pMRB1-derived vector containing the Gateway conversion cassette *attR*2-*ccdB*-Cm$^r$-*attR*1. Ap$^r$ Cm$^r$ | Jiménez-Fernández *et al*., 2014 |
| pMRB3 | pMRB1-derived vector containing the Gateway conversion cassette *attR*2-*ccdB*-Cm$^r$-*attR*1. Ap$^r$ Cm$^r$ | Jiménez-Fernández *et al*., 2014 |
| pMRB96 | pEX18-Tc bearing *fleQ* upstream and downstream flanking regions. Tc$^r$ Sac$^s$ Mob$^+$ | Navarrete *et al*., 2019 |
| pMRB99 | pSB1K3-derived plasmid expressing fleQ from its own promoter. Km$^r$ | Jiménez-Fernández *et al*., 2014 |
| pMRB104 | pEX18-Tc bearing *fleN* upstream and downstream flanking regions. Tc$^r$ Sac$^s$ Mob$^+$ | Navarrete *et al*., 2019 |
| pMRB119 | pMRB3-derived vector containing a *gfp*mut3::*lac*Z transcriptional fusion to the P*fleQ* promoter. Ap$^r$. | Jiménez-Fernández *et al*., 2016 |
| pMRB192 | pEX18-Tc bearing *fleR* upstream and downstream flanking regions. Tc$^r$ Sac$^s$ Mob$^+$ | This work |
| pMRB164 | pMRB172-based delivery plasmid for miniTn*7*-*nahR*-P*sal*-*yhjH*] Ap$^r$ Gm$^r$. | This work |
| pMRB165 | pMRB172-based delivery plasmid for miniTn-*nahR*-P*sal*-*pleD*. Ap$^r$ Gm$^r$. | This work |
| pMRB172 | pUC18Sfi-miniTn*7*BB-Gm-based delivery plasmid for miniTn*7*BB-Gm [*nahR* P*sal*]. Ap$^r$ Gm$^r$. | This work |
| pMRB178 | pMRB172-based delivery plasmid for miniTn*7*-*nahR*-P*sal*-*fleQ*. Ap$^r$ Gm$^r$. | This work |
| pMRB192 | pEX18-Tc bearing *fleR* upstream and downstream flanking regions. Tc$^r$ Sac$^s$ Mob$^+$ | This work |
| pMRB196 | pMRB3-derived plasmid including a polylinker between *Not*I and *Asc*I restriction sites. Ap$^r$ Cm$^r$. | This work |
| pMRB250 | pMRB196-derived vector containing a *gfp*mut3::*lac*Z transcriptional fusion to the P*flgF* promoter. Ap$^r$ | This work |
| pMRB259 | pMRB196-derived vector containing a *gfp*mut3::*lac*Z transcriptional fusion to the P*fliE* promoter. Ap$^r$ | This work |
| pMRB260 | pMRB3-derived vector containing a *gfp*mut3::*lac*Z transcriptional fusion to the P*fliL* promoter. Ap$^r$. | This work |
| pMRB261 | pMRB3-derived vector containing a *gfp*mut3::*lac*Z transcriptional fusion to the P*parC* promoter. Ap$^r$. | This work |
| pMRB262 | pMRB2-derived vector containing a *gfp*mut3::*lac*Z transcriptional fusion to the P*fliK2* promoter. Ap$^r$. | This work |
| pMRB264 | pMRB3-derived vector containing a *gfp*mut3::*lac*Z transcriptional fusion to the P*flhA* promoter. Ap$^r$. | Jiménez-Fernández *et al*., 2016 |
| pMRB265 | pMRB3-derived vector containing a *gfp*mut3::*lac*Z transcriptional fusion to the P*flhF* promoter. Ap$^r$. | This work |
| pMRB266 | pMRB3-derived vector containing a *gfp*mut3::*lac*Z transcriptional fusion to the P*fliK* promoter. Ap$^r$. | Jiménez-Fernández *et al*., 2016 |
| pMRB267 | pMRB3-derived vector containing a *gfp*mut3::*lac*Z transcriptional fusion to the P*hsbA* promoter. Ap$^r$. | Jiménez-Fernández *et al*., 2016 |
| pMRB268 | pMRB3-derived vector containing a *gfp*mut3::*lac*Z transcriptional fusion to the P*fliS* promoter. Ap$^r$. | Jiménez-Fernández *et al*., 2016 |
| pMRB269 | pMRB3-derived vector containing a *gfp*mut3::*lac*Z transcriptional fusion to the P*fliD* promoter. Ap$^r$. | Jiménez-Fernández *et al*., 2016 |
| pMRB270 | pMRB3-derived vector containing a *gfp*mut3::*lac*Z transcriptional fusion to the P*fliC* promoter. Ap$^r$. | Jiménez-Fernández *et al*., 2016 |
| pMRB271 | pMRB3-derived vector containing a *gfp*mut3::*lac*Z transcriptional fusion to the P*motA* promoter. Ap$^r$. | Jiménez-Fernández *et al*., 2016 |
| pMRB272 | pMRB3-derived vector containing a *gfp*mut3::*lac*Z transcriptional fusion to the P*flgB* promoter. Ap$^r$. | Jiménez-Fernández *et al*., 2016 |
| pMRB273 | pMRB3-derived vector containing a *gfp*mut3::*lac*Z transcriptional fusion to the P*cheV* promoter. Ap$^r$. | Jiménez-Fernández *et al*., 2016 |
| pMRB274 | pMRB2-derived vector containing a *gfp*mut3::*lac*Z transcriptional fusion to the P*flgA* promoter. Ap$^r$. | Jiménez-Fernández *et al*., 2016 |
| pMRB275 | pMRB2-derived vector containing a *gfp*mut3::*lac*Z transcriptional fusion to the P*flgM* promoter. Ap$^r$. | Jiménez-Fernández *et al*., 2016 |
| pMRB276 | pMRB3-derived vector containing a *gfp*mut3::*lac*Z transcriptional fusion to the P*cheA* promoter. Ap$^r$. | Jiménez-Fernández *et al*., 2016 |
| pMRB277 | pMRB3-derived vector containing a *gfp*mut3::*lac*Z transcriptional fusion to the P*fliH* promoter. Ap$^r$. | Jiménez-Fernández *et al*., 2016 |
| pMRB278 | pMRB3-derived vector containing a *gfp*mut3::*lac*Z transcriptional fusion to the P*flgZ* promoter. Ap$^r$. | Jiménez-Fernández *et al*., 2016 |
| pMRB291 | pMRB196-derived vector containing a *gfp*mut3::*lac*Z transcriptional fusion to the P*fleS* promoter. Ap$^r$ | This work |
| pMRB286 | pMRB172-based delivery plasmid for miniTn*7*-*nahR*-P*sal*-*fliA*. Ap$^r$ Gm$^r$. | This work |
| pMRB290 | pEMG bearing 1 Kbp insert with upstream and downstream flanking regions of *rpoN*. Km$^r$ | This work |
| pMRB291 | pMRB2-derived vector containing a *gfp*mut3::*lac*Z transcriptional fusion to the P*fleS* promoter. Ap$^r$. | This work |
| pRK2013 | Helper plasmid for triparental mating. ColE1 replicon. Km$^r$ | Figurski and Helinski, 1979 |
| pSB1K3 | High copy-number cloning vector. Km$^r$ | Shetty *et al*., 2008 |
| pSW-I | Plasmid expressing I-SceI from *xylS*-P*m*. *ori*RK2, xylS. Ap$^r$ | Wong and Mekalanos, 2000 |
| pTNS2 | R6K replicon-based helper plasmid expressing the Tn*7* transposase. Ap$^r$ Mob$^+$ | Choi *et al*., 2005 |
| pUC18Sfi-miniTn*7*BB-Gm | pUC18Sfi-based delivery plasmid for the synthetic minitransposon miniTn7BB-Gm. Ap$^r$ Gm$^r$ | Jiménez-Fernández *et al*., 2014 |

**Oligonucleotide Sequence (5' to 3')**

| | |
|---|---|
| cheA-cheB_fwd | GCCACGTGGTGATTCTGTC |
| cheA-cheB_rev | ATGGGCATTTCGTAGTCCAT |
| cheW2-cheW_fwd | GATTTCGGTGCAGGGCTAT |
| cheW2-cheW_rev | ATCAGGCCAAAACGCTGAC |
| cheY-cheZ_fwd | GCTGGAAAACGGTCACTACG |
| cheY-cheZ_rev | GGGTCGATCTGGAAACTGAC |
| cheZ-cheA_fwd | GATCAAGCGCGTCACCAC |
| cheZ-cheA_rev | CTCACCTTTGCGCAGGAC |
| flaG-fliD_fwd | GACCCTGGTGGTCAAGGTTA |
| flag-fliD_rev | TTCATCCGAAGACGTACCCG |
| fleN-down-fwd | GGACGGATCCTATGAACGCCAGCGGCTT |
| fleN-down-rev | TTTTAAGCTTCTTGTCCAATTAGACCTC |
| fleN-fliA_fwd | GACCGCTTCCTTGACGTTG |
| fleN-fliA_rev | CGTCGTATTTGTTGGCCACT |
| fleN-up-fwd | CCTGGAATTCCTGCTGGAAGTGCAACTC |
| fleN-up-rev | TGCAGGATCCCCATGTCTGTTCTTTACC |
| fleQ-fleS_fwd | GAACGTTTGCGTATTCGC |
| fleQ-fleS_rev | CTTCAAGCAGGCTGTAGGA |
| fleR-up-fwd-EcoRI | GGAAGAATTCCGCCACCCGCTCACT |
| fleR-up-rev-BamHI | TTCAGGATCCACATTACACCCCCTCCGCCAA |
| fleR-down-fwd-BamHI | TTCAGGATCCAATGACTGGCAAATCATCTGCA |
| fleR-down-rev-HindIII | GTGGAAGCTTGAGGCCGATCTGGCG |
| flgK-flgL-fwd | CTGGACCTGCAGACCAAGTC |
| flgK-flgL-rev | CGCTGTACTGCTCCAACAAG |
| flgL-P4379-fwd: | CAGCTAAGCCTGTTCGACAA |
| flgL-PP4379-rev | CGTAACTCGCAATGCTTTTG |
| flhA-flhF_fwd | CTTCTGGAACCTAGCATGGC |
| flhA-flhF_rev | AATGCGCGTGTGGGTCTT |
| flhF-fleN_fwd | GAGCCTTGCCATCAGTCATG |
| flhF-fleN-rev | CAGCAACACGTCGACATTGG |
| FleQ-SpeI-rev | GCATACTAGTATTATTATCAATCCTCCGCCTGGT |
| FleQ-XbaI-SD-fwd | ATGCTCTAGAGAAAGAGGAGAAATACTAGATGTGGCGTGAAACCAAGA |
| fliA-cheY-fwd | CAAGGAAATCGGTGAGGTGC |
| fliA-cheY-rev | CCGCTTCAATGATCTGGTCG |
| FliA-fwd-XbaI-SD | ATGCTCTAGAGAAAGAGGAGAAATACTAGATGAACGCCAGCGGCTT |
| FliA-rev-SpeI-Stop | GCATACTAGTTCATCATCAACGCGCCCGCCATT |
| fliC-fwd | GCAGCGTATGCGTGAACTGG |
| fliC-rev | CAACCACATCCTGCAGGGTG |
| fliC-flaG-fwd | CGTTTCGATAGCACTGTCGC |
| fliC-flaG-rev | CGTGGAATCGTCCACGGAAA |
| fliD-fliS-fwd | GGAATCTTGGCAACGCGGAC |
| fliD-fliS-rev | CCTTGCCCAGCATTACACCC |
| fliF-fliG-fwd | GTTCATCCTGGTGCTGGTGT |
| fliF-fliG-rev | ATGTCGACGAACTCGCTCAT |
| fliG-fliH-fwd | GCGCAGAAAGAAATCCTCAC |
| fliG-fliH-rev | CCTCGTTGTAAGCCTCCTGA |

| | |
|---|---|
| fliH-fliI-fwd | ACAATATCCGCATCCACCTC |
| fliH-fliI-rev | CTGCACCGGGTGATAACTGT |
| fliK-fwd | AACTGATTCAGGCGCAGG |
| fliK-rev | GCGTTCTGGTTCATCGGC |
| fliK-fliL-fwd | GTCAACGTGGCTGACCAG |
| fliK-fliL-rev | GCTCGCTCTTGTGCATGA |
| fliL-fliM-fwd | GACACCCTCGCCAGCAGC |
| fliL-fliM-rev | GCGCAGCGGCTTGATCTT |
| fliT-fleQ_fwd | CTGCGAGAGAATCAGGATG |
| fliT-fleQ_rev | AGGACAATGGCTCGACC |
| hptB_fwd | CAAGGTACTCAGTGACCTGC |
| hptB_rev | AAATACGTTGCTGCTCACC |
| hptB-fliK_fwd | TGTCAGCAGCTCGAAGAG |
| hptB-fliK_rev | CAACTTGTTGCCGTCATC |
| motD-parC-fwd | TTTCCCGTAACCTGGAAGTG |
| motD-parC-rev | AGGTCGTAGCAGCTGTGCTC |
| P4331-P4332-fwd | GGTGAGGCCATCCATGAG |
| P4331-P4332-rev | GTCGATCGCCATCTACCC |
| P4334-fwd | CACCGGCAATGGTAGTGGTGGTCT |
| P4334-rev | TCACTGACAACCTGCCAATC |
| P4379-fwd | CGACCTTGATCCGCAATC |
| P4379-rev | CGATATCCGACGACTTCAGG |
| PcheA-fwd | CACCCCTGGAGGATTTCTTCATCG |
| PcheA-rev | GCCTGTCGTATGTGGTCAAG |
| PfleSR-fwd-SpeI | AGCTACTAGTGTGAGCGGGTGGAAGACATC |
| PfleSR-rev-PstI | AGCTCTGCAGGTGCTCTCTCGCGTGGCT |
| PflgF-fwd-SpeI | ATGCACTAGTCAGAGCAACTACCAGGCCAAT |
| PflgF-rev-PstI | ATGCCTGCAGAGTTTCTCCTCTGCGCGTAGCT |
| PflgZ-fwd | CACCTAGGCTAGCCACCAGATTGC |
| PflgZ-rev | GGCAATGTCATCTTCGATCA |
| PfliE-fwd-SpeI | ATGCACTAGTGCCATTCCATTGTCGGCT |
| PfliE-rev-PstI | ATGCCTGCAGGACCCTTCTCTCTCCTGCG |
| PfliL-fwd | CACCCCAGCAGGATCAGCTTGAGT |
| PfliL-rev | TCAAGTCGGCTGATATCCAA |
| PmotA-fwd | CACCCAGCACCTCCAGGTAGAAGG |
| PmotA-rev | CATTGGTGGGGTGATTGG |
| rpoN fwd EcoRI | ACGTGAATTCCGGCATGAGCCTGTCTGG |
| rpoN-overlap-fwd | GGTATTAAGCCCCTGCCATGCCCCTGAGATGTGCCACA |
| rpoN overlap-rev | TGTGGCACATCTCAGGGGCATGGCAGGGGCTTAATACC |
| rpoN-rev-BamHI | ACGTGGATCCCGATGACAGTGGCGACCTTT |

### SUPPLEMENTARY FIGURES

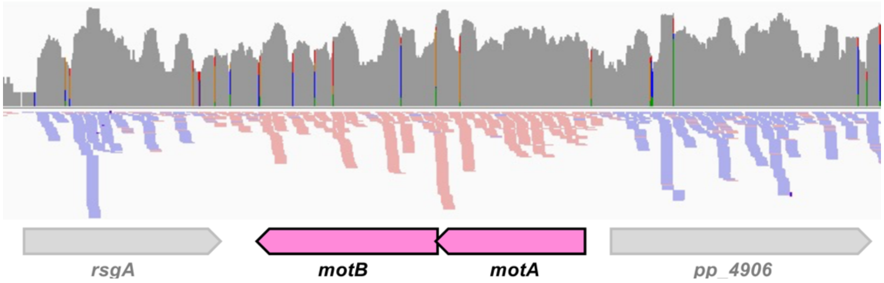


**Supplementary Figure S1. RNA-seq analysis of the *motAB* operon.** Output of the RNA-seq analysis of the genomic region containing the *motAB* operon. Coverage is represented in grey on a logarithmic scale. Top strand and bottom strand reads are shown in blue and pink, respectively. A cartoon of the flagellar genes is aligned with the RNA-seq reads. Genes are color-coded as in **Fig. 1**.

```
PfliK2
Ppu KT2440  TATCTTCAGTAGGGCTTTGAAGCAGCCAACACCGCTCCAGAACCCTTGTGTCTCAAGTCGCCCCGCGCGCTGCCGTTATACTGGCAACCA
Ppu F1      AGGAGCGGGTTCACCCGCGAAGCAGCCAACATCGCTCCAGAACCCTTGTGTCTCAAGTCGCCCCGCGCGCTGCCGTTATACTGGCAACCA
Psy DC3000  CTGTTGCGTCGCGAAATGTGTCAAAACAGACACATACGGAGTAAAAGACGGTTCAAGTCAGGATTGATGCAGCCGTTACATGAAGAGAAC
Psy B728a   GTTTTGCGTCGCGAAATGTGCCAAAACAGACACATACGGAGTAAAACGCGGTTCAAGTCGGGGTTGATACGGCCGTTACATGAAGAGAAG
Psy 1448a   CTGGAGCGTCGCGAAATGTACCAAAACAGACACATACGCAGTAAAAGGCGGTTCAAGTCGGGGGTGATATGGCCGTTACATGAAGAGAAG
Pfl SBW25   GCGCTATGGCTTCAAGAAGAGGGCTGGTCAAAAGATCGCGGGCCTTTACGGTTCAAGTCGGCGGTGCAGCGGCCGTTAAAGAGAAATGCA
Ppr Pf-5    TGCTACGGCTTCGAGAAGAGGGCTGGTCAAGGGATAGCGGGCCTTTACAGGTTCAAGTCGGCAACGGAGCGGCCGTTAAAGAGATGCAGG
                                                                ******             **** ***
                                                 FliA consensus TcAAgNw   —N12-13- gcCGAtaNc
PparC
Ppu KT2440  TGCGCCGCAGCCTGACCGGCTCGGGCAGCGCCAATGCCACACCGGATGCCGCATTGCGCCGGGCTGGCACACAAAGTGCACCGGCAATAC
Ppu F1      TGCGCCGCAGCCTGACAGGCTCGGGCAGCGCCAATGCCACACCGGATGCCGCATTGCGCCGTGCTGGCACACAAAGTGCACCGGCAATAC
Psy DC3000  TTCGCCGCAGCCTGACCGGTTCCGGCACGGCCAATGCGCAGCCCGACGCTGCATTACGCCGTGCTGGCACACAAACTGCACCGGTGCCAG
Psy B728a   TTCGCCGCAGCCTGACCAGCGCCGGATCAGCCAATGTGCAGCCGGATGCTGCATTGCGGCGTGCTGGCACACAAACTGCACCGGTGCCAG
Psy 1448a   TGCGTCGCAGCCTGACCAGTTCAGGGTCGGCCAATGTGCAACCGGATGCTGCACTGAGGCGTGCTGGCACACAAACTGCACCGGTGCCAG
Pfl SBW25   TACGCCGCAGCCTGACCGGCACGGGCACCGCCAATGCAACACCGGATGCGGCCTTGAAGCGGGCTGGCACACAAACTGCACCGGCCCCTG
Ppr Pf-5    TGCGCCGCAGCCTGACCGGTACCGGCACCGCCCATGCAACACCGGATGCGGCACTCAAGCGTGCTGGCACACAAACTGCACCAACCCCGG
                                                                            ******      ****
                                                              σ54 consensus TGGCACG-N4- TTGCw
PcheA
Ppu KT2440  ACCACGATCAATTGCGTGCAGAAAAAGATCGAGAAAAACATCCGACTCGGGGTGAAGGTCCGCAGATTCATGCCGATAAGCGTGAAGACG
Ppu F1      ACCACGATCAATTGCGTGCAGAAAAAGATCGAGAAAAACATCCGACTCGGGGTGAAGGTCCGCAGATTCATGCCGATAAGCGTGAAGACG
Psy DC3000  ACGAAGAATCCATCCTTGCTGAAAAAGATCCTAAAAAACATCTCGACAAGGGTGAAGGTCCGCAGATTCATGCCGATAAACGTGAAGACG
Psy B728a   ACGAACAATCCATCCTCAATGAAAAAGATCCTAAAAAACATCTCGCTCAGGGTGAAGGTCCGCAGATTCATGCCGATAAACGTGAAGACG
Psy 1448a   ACGAAGAATCCATCCTTGCTGAAAAAGATCCTAAAAAACATCTCGCCAAGGGTGAAGGTCCGCAGATTCATGCCGATAAACGTGAAGACG
Pfl SBW25   ACCGCGAAGCGATCCTCTCGGAAAAAGATCCACAAAAACATCTCGCCAAGGGTGAAGGTCCGCAGATTCATGCCGATAAACGTGAAGACG
Ppr Pf-5    ACCGTGAATCGATCCTCGCTGAAAAAGATCCGCAAAAACATCTCGCCAAGGGTGAAGGTCCGCAGATTCATGCCGATAAACGGGAAGACG
                                                                * *****            ********
                                                 FliA consensus TcAAgNw   —N12-13- gcCGAtaNc
PflhF
Ppu KT2440  GCCATCGTGCAAAGCATTGTCGGCGTTGAGTCGGAGCTACCAGTGATTACCCTGGAGCCAAGGTTGGAACAGATTTTGCTGAATAGTCTG
Ppu F1      GCCATCGTGCAAAGCATTGTCGGCGTTGAGTCGGAGCTGCCAGTGATTACCCTGGAGCCAAGGTTGGAACAGATTTTGCTGAATAGTCTG
Psy DC3000  GCAATCGTCCAAAGCATTGTAGGCGTTGAGCCGGAGCTGCCTGTTATCACTCTGGAACCAAGGTTGGAACAGATATTGCTCAATAGTTTG
Psy B728a   GCAATCGTCCAAAGCATTGTAGGCGTTGAGCCGGAGCTGCCTGTTATCACTCTGGAGCCAAGGTTGGAACAGATATTGCTCAATAGTTTG
Psy 1448a   GCAATCGTCCAAAGCATTGTAGGTGTTGAGCCGGAGCTGCCTGTTATCACTCTGGAACCAAGGTTGGAACAGATATTGCTCAATAGTTTG
Pfl SBW25   GCAATCGTGCAAAGCATTGTAGGGCTTGACTCCGAGCTGCCTGTGATCACCTTGGAACCAAGGTTGGAACAAATATTGCTCAATAGTATT
Ppr Pf-5    GCCATCGTCCAAAGCATTGTAGGCACTGAGTCTGAGCTGCCTGTGATCACCTTGGAGCCAAGGTTGGAACAAATTTTGCTCAGCAGTCTG
                                                                            *** **     *****
                                                              σ54 consensus TGGCACG-N4- TTGCw
PflhA
Ppu KT2440  GAGCCACCCGTCAACGCGGTGGTGCTCGCCCGGTTCATCTTTTCGTGTGGGTGTCTGCCAAAGTTGGAAAGCTTCTTGCAAAGCCACGCC
Ppu F1      GAGCCATCAGCCAACGCGATGGAGTTCGCCCGGTTCATCTTTTCGTGTGGGTGTCAGCCAAAGTTGGAAAGCTTCTTGCAAAGCCACGCC
Psy DC3000  CTGATCACCTCGTCTTGCGCAATTTCTACCTCTAAAGCCTTGCCGGCACTGGCTTTGTAAAAGTTGGACAGCTTTTTGCAATAAGGTCGA
Psy B728a   TCAGCGCACCGGATTTACGTGACTTCTGTGCCTAAAGCCTTACCGCCACTGGCTTTGTAAAAGTTGGACAGATTTTTGCAACAGGGTCGA
Psy 1448a   AAGGTATGCCGGTTTCACATTGCTTCTGCCTTTAAAGCCTTACCAGCACTGGCTTTGTAAAAGTTGGACAGCTTTTTGCAACAGGGTCGA
Pfl SBW25   CAGCCCAGCGGGAGCAAGCTCCCTCGCCACAGGTGCGACATGATCTGCGGCATGGGGCAAAAGTTGGAAGGCTTCTTGCAATACCGCCTG
Ppr Pf-5    GCCACTCTGAGGTTTCCGGCATTTTCGGCTCAATGCCGGCGCCATATTTCCTTTCGGTAAAAGTTGGAAGGTTTCTTGCAGTAGCAGCCC
                                                                            ****  *    *****
                                                              σ54 consensus TGGCACG-N4- TTGCw
PfliL
Ppu KT2440  CGCGGGCGAGCCCGCGAAGAAGCCAGCACCGCAAAACCTGCTGGAATCCATCCCCTGCAAAATCTGGCATAACACTTGCTCAAGCCAAGC
Ppu F1      AGCGGGCGTGGCCGCGAAGCAGCCAGCACCGCAAAACCTGCTGGAATCCATCCCCTGCAAAATCTGGCATAACACTTGCTCAAGCCAAGC
Psy DC3000  ATGTCCAGCGCCTGTCTCCGCGCGTAACCGTTTGTGTCAGATACATGCCCCTCTCGTACAACTCTGGCATAACACTTGCTCTTCCTCTGT
Psy B728a   ACGATCCCTGCGCGCCCTGGCGCGTAACCGGTTGTGTCCGATACATCCCCGTTTCGTACAACTCTGGCATAACACTTGCTCTGCCTCTGT
Psy 1448a   ACCATCCCTCCGCGCCCAGCCGCGTAACCGGTTGTGTCCAATACATCTCCATCTCGTACAACTCTGGCATAACACTTGCTCTGCCTCTGT
Pfl SBW25   GGGGGCAAGCCCCCTCCCACAATTTGACTTGCCGCGTCCGTACGTCCCCCACTTCAGACAACTCTGGCATAACACTTGCTCTTGCCTTTC
Ppr Pf-5    TGCACCCCAGGTTTCCCTTCGCAGCGTTGCCCTTCGCTGTTGATCCCTGCCTTTCTGACAAATCTGGCATAACACTTGCTCTTGCCTTGC
                                                                            ***** *    *****
                                                              σ54 consensus TGGCACG-N4- TTGCw
PfliK
Ppu KT2440  CGTCGTTGTGCAGCACTTTTACCGTGGTGAGCAGCAACGTATTTCCGCAGGCTGATACGCGAGTTGGCGCGAGTCTTGCCTGTCTCTGCC
Ppu F1      CGTCGTGGTGCAGCACTTTTACCGCGGTGAGCAGCAACGTATTTCCGCAGGCTGATACGCGAGTTGGCGCGAGTCTTGCCTGTCTCTGCC
Psy DC3000  TGCTGAGCGCCAGCTCTTCATCAGTTGATCGATAATGCCCTGTCGAAGGGCTGTATTTTAAAGTTGGCCCACATATTGCTATGTCTTGTT
Psy B728a   TGTTCGTCATCTGAGCCCTTAGAGTCATACCCTCGATCCCCGCTATGGGGAATCATTAGATAACTGGCCCGTATATTGCTAACTATTTCG
Psy 1448a   ATCACTGATTTCCAGATCCCCGATCCTCCTTTATTCGGCAACCTTTGAAGTGGGTGTGGAAGGTTGGCCCGCATATTGCTCAGTGTTCTG
Pfl SBW25   TGCTGGGGCGGACCCCGATCAGCCGTTACCCACACAACGGATAGGTACCCGTAAAATCAAAACCTGGCCCAACTCTTGCTCTACCTCTAG
Ppr Pf-5    GCGAGCAGGAGCTGCAGCGCTTTCATTGTTGAGAAGCAGCTTTTTATCTGCGGATTGATAAAGCTGGCCCGGCCTTTGCAGTCAAGTCTT
                                                                            **** *     ****
                                                              σ54 consensus TGGCACG-N4- TTGCw
PhsbA
Ppu KT2440  TGCCCGCGAAGAGGCCGGTACAGGCCATCAAAGCCATGAGCCTGCATCCGCTTGCCTCCCCCACCATCGCC TGCTAAACCTTAAAGCAG
Ppu F1      TTACCCGCGAAGAGGCCGTACAGGCCATCAAAGCCACGAGCCTGCATCCGCTTGCCACCCCCACCATCGCC TGCTAAACCTTAGAGCAG
Psy DC3000  CGAACAATTCTGACGTTTCATTTAATCAGTCCCGGAGATCTTCCGGTCTGGTCTACGCTATTAATGTCTCGGTGATGAAACACCTCGAGG
Psy B728a   AACAATTCTGACGTTTGACTCAATCAGACCCGCAGTGATCTCTCTGGCTGGTCTACGCTATCAATGTCTCGGTGATGAAACACCTCGAGG
Psy 1448a   AACAATTCTGACGTTTGACCCAATCAGACCCGCGGTGATTTCTCTGGCTGGTCTACGCTATCAATGTCTCGGTGATGAAACACCTCGAGG
Pfl SBW25   AGCCCGCTCACTACAAATAGTGCATAGCCCACACACATTTGATGTGTTGTGTTGCCCATAGCCGGTTCACC TGCTAAACCTTGTGTCAT
Ppr Pf-5    CGCTTGCCTCGTCAGAGCCCATACTGATGGTGCTCATTTGCCCCGTGCGGGTTGTCCCCCCGGCGAGTGGG TGCTAAACCTTTTAGACG
                                                               *   *                ****  *
                               σ70 consensus with extended -10 TTGACA     -N17-18-  TGNTATAaT
PfliE
Ppu KT2440  CTATCGCGACGCAAGGCCGCTCCCGCAGGAATTGTGTGATCTCTGGGAAATGAGTTGCGAAGGCTGGCACCTTTGTTGCTTTAGGTAGGT
Ppu F1      CTATCGCGACGCAAGGCCGCTCCCACAGGAATTGTGTGATCCCTGGGAAATGAGTTGCGAAGGCTGGCACCTTTGTTGCTTTAGGTAGGT
Psy DC3000  AGGGCGATTTGATGTCGGGCAACACGGCATTCCTGCAGGGTGTGCCAAATTGCCACTATCCAGCTGGCACCGTTGTTGCTAAGTCTTCTG
Psy B728a   GGCGCGAACTCGTGTCAAGCGACATGATGCACTCATCTGGTGTGGCAAATAGCCGCCATCTAGCTGGCATCGTTGTTGCTATACCTTCTG
Psy 1448a   GGAGTGATTTCATGTCGGGCAACATGGTATTCCCATTTGGTGTGGCAAATAGCCGCCATCTAGCTGGCACCGTTGTTGCTATGTCTTCTG
Pfl SBW25   CGCCACATAACTGGTAACAAGGTCGACAACCTTTGTCGCCTTTTCTTACGAGTTGATCCAGAGCTGGCACCCTTGTTGCTACTTCCCTAG
Ppr Pf-5    GTTCGCGACCTGATCTGTGAGAGGCAAGCCCGGGATGGGCTTTTGTCGATAAGGGCCATGGAGCTGGCACCCTTGTTGCTAACACCTCAC
                                                                            ***** *    *****
                                                              σ54 consensus TGGCACG-N4- TTGCw
PfleS
Ppu KT2440  AAAACTCACGGCGAACCCTTCTCGGTCTTAACGCAATCCTACGTCAAACCCAAACCCCCATCTCCGGCACGGCTATTGCTACACCGCTGG
Ppu F1      CGGTTGCCTGTGCCGGCTTCTTCGCGGGCGTGCCCGCGAAGAGGCCGGCACAAATCCCTATCTCCGGCACGGCTATTGCTACATCGCTGG
Psy DC3000  CGGAGGATTGACGCCTGTTTCAAACGCCAAAAATCAAGCCCTTGAATTTTCAAGGGTTTTTTTTCGGCACGAGGATTGCTAAGTCTCTCT
Psy B728a   CAGATGATTGACGCCGGTTTTTGTGCCAAAAACCAAGCCCCTGATTTTCAGGGGTTTTTTTTAGGCACGGGGATTGCTAAGTCTCTCT
Psy 1448a   CAGATGATTGACGCCTGTCTGTAGCGCCAAAAATCAAGCCCCTGAATTTTCAGGGGTTTTTTTTCGGCACGAGGATTGCAAAGTCCCTCT
Pfl SBW25   GACGGCAACAGCCACTCGTCAAAACGAAACTGTATTGGCGAGGCCGGGCACCTTGGTTTTTTTTCGGCACAAGTATTGCTACAGCCCTCG
Ppr Pf-5    GTGATGAACAGGCGGATGATTGACGCCGGTTTTTCAAGTCTTTGAAAACTGGGTGGTTTTTTTTAGGCACGGGTATTGCTAAGTCTCTTG
                                                                             *****     *****
                                                              σ54 consensus TGGCACG-N4- TTGCw
PfleQ
Ppu KT2440  TGGGCAGAGAGCAACGCTTCGGCGCCATAAATTTGACTGCCTTTAAGTTTTTGACTTTACTAGTGGCTGTTTTCGAATTTCAGACGTCCG
Ppu F1      ACCGATGTTGCAAAAAGCCGTGCGCCATTAATTTGACTGATCGGTGTTTTTTGACTTTACTAGTGGCTGTTTTCGAATTTCAGACGTCCG
Psy DC3000  TTTAATCCGTCGACAAGAGTTTCGCCATAAATTTGACTGTGTTCGTTTTTTTGACTTAACTAGTG ATGTTTACTATTTTCTGGCGTCTC
Psy B728a   TTTTATTGTTCGACAAGAATTTCGCCATAAATTTGACTGTGTCCGTTTTTTTGACTTAACTAGTG ATGTTTACTATTTTCTGGCGTCTC
Psy 1448a   TTTTATTGTTCGACAAGAATTTCGCCATAAATTTGACTGTGTTCGTTTTTTTGACTTAACTAGTG ATGTTTACTATTTTCTGGCGTCTC
Pfl SBW25   GTCGCAGTTAATCCTACCGGTGCGTCATAAATTTGACCGTGCCAGCTTTTTTGACTTAACTAGTG CTGTTTTCAGATTTCAGACGTCTA
Ppr Pf-5    TGAGTTAATCCCGAACACTGCGCGCCATAAATTTGACTGTGCACGCTTTTTTGACTTAACTAGTGGCTGTTTTCAGATTTCAGGCGTCTA
```

```
                                           *****                  *  * *
                             σ70 consensus TTGACA      -N17-18-   TATAaT
                                                             *****                        *
                                               σ70 consensus TTGACA      -N17-18-    TATAaT
PfliS
Ppu KT2440 GACGACCCTGAACGCGATGAACAAGGCCAATAACGACGACTGATGGTCGTATTCAAGATCTTTGGGCTAAGCCGATCACTTAGGTATGT
Ppu F1     ATTGACAGCAGTATCAAGAACCCGACTTCGTGTCGGGTTCTTGCTTTTGGGCTAAAGTTTCTGCGATGCGTCGATAGCTTGAATATCA
Psy DC3000 ATTGACAGCAGTATCAAGAACCCGACTTCGTGTCGGGTTCTTGCTTTTGGGCTAAAGTTTCTGCGATGCGTCGATAGCTTGAATATCA
Psy B728a  CAGTGCAGAAGTACCAGGAACCCGACGCTGTGTCGGGTTCTTGCTTTTAGGCTAAAGTTTCCGCGCACTGCGTCGATAGCTTGAGTATCA
Psy 1448a  TTTTACAGAAGTATCAAGAACCCGACTCCGTGTCGGGTTTCTGCTTTTGGGCTAAAGTTTCTGCGCCATGCGTCGATAGCTTGAGTATCA
Pfl SBW25  AAAAGGCCAGACGGTTGTCTGGCCTTTTTTTTCATCTGGAGACGACTCGCGCTAAAGCTTTTTGCGAGCCAGTCGACATATTGGTTACTG
Ppr Pf-5   GCGCCAAAAGCCCGGCAGCGTTTTTCAGCGCTCCGGGCTTTTGTATTTGGCCTAAAGTTTTTTGACGTTGCGTCGATATATTGCTTATAC
                                                               * *** *           * *** **
                                                FliA consensus TcAAgNw  -N12-13- gcCGAtaNc
PfliD
Ppu KT2440 CCTGAGCCTGGCTCGCAGTCTGGCGGAGGGTGATGGTTTTTTACTTGATGACAATGTGTAATGTTGGCATGAGTCTTGACTCGGATTGTG
Ppu F1     GCATTGCGGATTGCGCATAACCTGAGCGATGTAAACAGTATTTTGTTCGACGCCAAGGTTTGATCGGCGGAGAATGTGCTAAAGCAGGAA
Psy DC3000 CTGAAACTGGCCGATAGCCTGAGTGATGCGAACAGCTTGTTGTTCCGCGCCAGAGCGTGATCGCCGGTACGGTACTTGTTGCAGGTTGCC
Psy B728a  CTGAAACTGGCCGATAGCTTGAGTGATGCGAACAGTTTGTTGTTCCGCGCCAAGGCCTGATCGCCGGTACCGCATGTGTTGTCGGGTCAT
Psy 1448a  CTGAAACTGGCCGATAGCCTGAGTGATGCGAACAGCTTGTTGTTCCGCGCCAAGGCCTGATTGTCAGTACGGTATTTGTTGCCGTTTTGC
Pfl SBW25  CACTGAAGCTGGCACAGAACCTTTCCAGCGCAAGCCACCTGTTGTTTGACGACAAGGTCTGAGCTGGCATGAAAATTGTTGCTGTCATGT
Ppr Pf-5   CTTAAACTGGCCAATAGCTTGAATGATGCGAGTAGTCTGTTGTTCAGTGCCAAGGCCTGACAGCTGGCATGAAAATTGTTGCTGGTTACG
                                                                             *         ** *
                                                             σ54 consensus TGGCACG-N4- TTGCw
PfliC
Ppu KT2440 CTTGATTTCTCAGGGCTGGCCCCATGCCAGTATTTTTTTTTTTTGAAAAAAACCCTCAAGCAACCCGCGCACCCGACGATAACCATTACGAAG
Ppu F1     CCTTGATTTCATTGGACTGGCACGGTGCCAGCAATTTTTTTTGAAATTGCGCTAAAGCAAGACGCCCTGACGACGATAACTATTACGAAG
Psy DC3000 CCTTGTTTTATAAGGGGTGGAGGAATGATGGCAAAATATTCAAAAAAAACGCTCAAGCAACCTGCCATCGCGACGATAACTATTACGAAG
Psy B728a  CCTTGTTTTATAAGGGGTGTCACGATGATGGCAAAATATTAAAAAAAAACACTCAAGCAACCTGCCATCGCGACGATAACTATTACGAAG
Psy 1448a  CCTTGTTTTATAAGGGGTACAGGCATGATGGCAAAATATTCAAAAAAAACACTCAAGCAACCTGCCATCGCGACGATAACTATTACGAAG
Pfl SBW25  CCTTGAATTACAAGGGGTGGCACGACGCCAGTAAATATTTTCAAAAAAATCCTCAAGCAACCGGCTACCACGACGATAACTATTACGTAG
Ppr Pf-5   CCTTGTTTTCCAAGGGATGGCCGGGTGATGGCAAATATTTTTGAAAAAGCCCTAAAGCAACTTGCATTACCGACGATAACTATTACGAAG
                                                               * *****           * *******
                                                FliA consensus TcAAgNw  -N12-13- gcCGAtaNc
PflgF
Ppu KT2440 CGACAAACTGACGGTTATTGCCCCTTGCCGCGCTGAAAGCCCCGTAAACCGGGTGTTTCAGACTTGGTTCAATTATTGCTTGGAACCTGC
Ppu F1     GCCGTCGCTGCGCGGGCTTTCTCAACTGCCCGTATTCATGGGCTTTTTCCTGCTTTTGGCAAGTTGGTTCGGAACTTGCTTTAGCCCCTG
Psy DC3000 GAAGGTGATCGTCGCGTGCCGGTTTTTGCTGGTTAATCAGGCATTTGTATCGAGGCTGACAAGTTGGTTCAAAAATTGCTTTAACCCTTT
Psy B728a  TCAGGCCGTCGGCGAATGCCGGTTTTTACTGGCTAATCAGGCATTTGTATCGAGCCTGACAAGTTGGTTCAAAAATTGCTTTAACCCTTT
Psy 1448a  GAAGGGCATCATCGAATGCCGGTTTTTGCTGGTTAATCAGGCATTTGTATCGAGCCCGACAAGTTGGTTCAAAAATTGCTTTAACCCTTT
Pfl SBW25  CTGAAATCTTCATACCTCGCCGCAAACCCCCGCCCCGCCTGCACTTCATGCTCTTTTTAAAAGTTGGTTCGCAAATTGCTTAAGCCTCAT
Ppr Pf-5   GTTTTTTCCTCTGTAATCAGGCAAAGCCTTGCCCCGTCTGGGGATCACCACTGCGTGAAAAAGTTGGTGCGCTTATTGCTTATGGCTTCG
                                                                           ***  *     *****
                                                             σ54 consensus TGGCACG-N4- TTGCw
PflgB
Ppu KT2440 TTTTCTGGCATCAATCTGACAGTACCGCGCCGCAAAACCCCATAAATACGGGCTTTCCAGTGGTTGGCACAGCCCTTGCTATGCCTTGCT
Ppu F1     TTTTCTGGCATCAACCTGACAGTACTGCGCCGCAATCCCCCCTAAACACGGGCTTTTCAGTGGTTGGCACAGCCCTTGCTATACCTTGCT
Psy DC3000 CGCTTTTCTGGCATTGCCGCCTCTGCGCTGTCTGTAACCCTCGCAATCATTGGCTTTTGAAACATGGCACGCCGATTGCTTTGTCAGTAA
Psy B728a  CGCTTTTCTGGCATTGCCGCCTCGCCGCCGCCTGTGGACCCTGCAAACATTGGCTTTTACAACATGGCACGCCGATTGCTTTGTCAGTAA
Psy 1448a  CGCTTTTCTGGCATCGCTGCCTCGCCGCTGGCTGCGGCCCCCGCAATCATTGGCTTTTGCAACATGGCACGCTGATTGCTTTGTCAGTAA
Pfl SBW25  GCTTTTCTGGCATCGTCGCCGGCAACCGCCTCGCAAGCCCTTGATATACGGGCCTCCACCAGATTGGCATGCACCTTGCTATAACCCTGT
Ppr Pf-5   CTTGCCGCTTCCGGGTTACCCTTAAATAAACTCAAGTTATTGAATATAAAGGTTTTTAATAATTTGGCACGGGCCTTGCTGAAGTGTTGG
                                                                           *****      *****
                                                             σ54 consensus TGGCACG-N4- TTGCw
PcheV
Ppu KT2440 TCATAGCGGCCGGCAGTCTGGGTGGTGGACAGATAATCTTCAACGGTGAATTCAAGAAACCCTTGGGTGACACCGATAAGCTGTTCAGGC
Ppu F1     TCATAGCGGCCGGCAGTCTGGGTGGTGGACAGATAATCTTCAACGGTGAATTCAAGAAACCCTTGGGTGACACCGATAAGCTGTTCAGGC
Psy DC3000 TCGTGACGCCCGTCAATCTGGTTGGTTGCCAAATAGTCTTCTACTGTGAACTCAAGAAACCCTTGGGTGACGCCGATAAGCTGTTCAGGC
Psy B728a  TCATGACGCCCGTCAATCTGGTTGGTTGCCAAATAGTCTTCTACTGTGAACTCAAGAAACCCTTGGGTGACGCCGATAAGCTGCTCAGGC
Psy 1448a  TCATGACGCCCGTCAATCTGGTTGGTTGCCAAATAGTCTTCTACTGTGAACTCAAGAAACCCTTGGGTGACGCCGATAAGCTGTTCAGGC
Pfl SBW25  TCATAACGGCCCGGCGTCTGAGTGGTTGCCAGATAATCTTCTACAGTGAACTCAAGAAAGCCTTGAGTGACGCCGATAAGTAGATCAGGC
Ppr Pf-5   TCATAGCGACCTTCCGTTTGACTGGTGGCCAGGTAGTCTTCTACCGTGAATTCAAGAAACCCTTGAGTGACGCCGATAAGAAGATCAGGC
                                                              *******              *******
                                               FliA consensus TcAAgNw   -N12-13-  gcCGAtaNc
PflgA
Ppu KT2440 CATAACACCCGCCATGCCAGACTCCTTAACGTTTCAGCCTTTCGTTGCAACTCAGGTAGCGAGTCGGCACGGGCTTTGCTTTTTTGAGTG
Ppu F1     CATAACACCCGCCATGCCAGACTCCTTAACGTTTCAGCCTTTCGTTGCAACTCAGGTAGCGAGTCGGCACGGGCTTTGCTTTTTTGAGTG
Psy DC3000 ATTACACCTGCCATGCCGAACCCCCTTACCAAAATGACCGCCACCCAGGCACCAGACCAACAAGCGGCACGGGCTTTGCTATTTGTTACG
Psy B728a  CATTACACCTGCCATGCCGAACCCCCTTACAAAATGACCGCCGCCCAGACACCAGATCAACAAGCGGCACGGGCTTTGCTATTTATTACT
Psy 1448a  ATTACACCTGCCATGCCGAACCCCCTTACCAAAATGACCGCCGCCCAGACACCAGATCAACAAGCGGCACGGGCTTTGCTATTTATTACC
Pfl SBW25  CTCCTGCCATGCCTTGACTCCTACGCTAAAACCTTTGGGGGTGCGATGCGCACCCAACACTAAACGGCACGAGCTTTGCTATTTATTGGT
Ppr Pf-5   CCATTACACCAGCCATGCCCAGACTCCCCAAAGCTTTAAGATCCGACGCACGCTCATTGCTAAACGGCACGGCGCTTGCTTTTTAACTGT
                                                                            ******    *****
                                                             σ54 consensus TGGCACG-N4- TTGCw
PflgM
Ppu KT2440 TGTCAGAACACGGGTTCGCGGGCTTGCCGACAGGCGGCAATGCATTTGTGCCTAAAGTTTATATCGGGTTGGCCGAAAACAAGGCAAGCG
Ppu F1     TGTCAGAACACGGGTTCGCGGGCTTGCCGACAGGCGGCAATGCATTTGTGCCTAAAGTTTATATCGGGTTGGCCGAAAACAAGGCAAGCG
Psy DC3000 GGAATGCCCGACGGTGAAAAGCATCTATTGAGAAGCTTTATCACAATCGGCCTAAAGTTTAATTGGGGTTGGCCGAAAACAAGGCAAGCG
Psy B728a  GGAATGCCCGACGGCAAATGTCGTCTATTGGGAAGCTTTCTCACAATCAGCCTAAAGTTTGATGGGGGTTGGCCGAAAACAAGGCAAGCG
Psy 1448a  GGAATATCCGACGGTGAAAAGCGTCTATTGGGAAGCTTTCTCACAATCGGCCTAAAGTTTGATAGGGGTTGGCCGAAAACAAGGCAAGCG
Pfl SBW25  TTCTACACTGTGCACGAGATATTCCGTGCGCAGATTTATTGTCATTCGCGCCTAAAGTTCGTCCGGGTATTGCCGAAAACATGGCAAGCG
Ppr Pf-5   GTGGCCGATAAAGGGTCGCAGATACCTGACCTGGTTTGTCGACAATCGAGCCTAAAGTTTTTCAGGGTATGGCCGAAAACATGGCAAGCG
                                                               * *****           ***** ***
                                                FliA consensus TcAAgNw  -N12-13- gcCGAtaNc
PflgZ
Ppu KT2440 CGCCGCTGGCCAAGCCGCGCGCGCTCAGCCAAGTGTGATTTTTCTATCAAGGCACAGAACATAGTGGCAAAATGCCGTTACTTGCGTGTG
Ppu F1     CGCCGCTGGCCAAGCCGCGCGCGCTCAGCCAAGTGTGATTTTTCTATCAAGGCACAGAACATAGTGGCAAAATGCCGTTACTTGCGTGTG
Psy DC3000 GCTTAGAGCAAAGCCACGCCCCTTGAGTCAGGCGTAACTTCCTGTATCAAGGTTTCGTGCGTAGTGGCAGAATGCCTGTTGTTGCGTTGC
Psy B728a  GCTTAGAGCAAAACCACGCCCTCTGAGCCAGGCGTAACTTCCTGTATCAAGGTTTCGTGCATAGTGGCAGAATGCCTGCTGTTGCGTTGC
Psy 1448a  GCTTAGAGCAAAACCACGCCCTCTGAGTCAGGCGTAACTTCCTGTATCAAGGTTTCGTGCGTAGTGGCAGAATGCCTGTTGTTGCGTTGC
Pfl SBW25  CCAAGCACTCGCAGCCAAGCCTGACGCCGGTTTACAGCGCGGCTATCCAAGACGCGCCACATACTGGCAGAATGCCTGCTCTTGCGTGTA
Ppr Pf-5   CACGCCCACTCAGTCAGGCTTAAACCCAAAAAATGAGCGCGCGCTACCAAGCCGTGAAACATGCTGGCAAAATGCTGGCTCCTGCGTGTA
                                                                           *****
                                                             σ54 consensus TGGCACG-N4- TTGCw
PmotA
Ppu KT2440 TTTCAATTGGCATTTTGACATCAGTCCGAACTGGCTTTTCGCATTGAGTGGCTATAGTCTGGCGCAGTTCTGCCGATAAGTAGAAGAAGA
Ppu F1     TTTCAATTGGCATTTTGACATCAGTCCGAACTGGCTTTTCGCATTGAGTGGCTATAGTCTGGCGCAGTTCTGCCGATAAGTAGAAGAAGA
Psy DC3000 CCAGTTTCCCCTTTTTATCGGACCGGGAAAATGGTTTTTCACAAGAACCGGCTATAGTCTGGCCCAGTTAGGCCGATAAGGAGAAGAAGA
Psy B728a  CATTTTCCTCCTTTTATGGAGGCGCATAAATGGGTTTTTCAAAAGAACTGGCTATAGTCTGGCCCAGTTAGGCCGATAAGGAGAAGAAGA
Psy 1448a  CATTTTCCTCCTTTTATGGAGGCGGATAAATGGGTTTTTCAAAAGAACCGGCTATAGTCTGGCCCAGTTAGGCCGATAAGGAGAAGAAGA
Pfl SBW25  TTGCGGTCAACACACCCGGAGCGGGCCGAACTGGCTTTTCGCCTTAACGGGCTATAGTCTGGCGCAGTTTTGCCGATAAGAGGAACAAGA
Ppr Pf-5   CTGCTGTGAACACGCCCAGAGCGGGCCAAACTGGCTTTTCGCCTTAACTGGCTATAGTCTGGCGCAGTTTTGCCGATAAGTAGAAGAAGA
                                                               *  ***            ********
                                                FliA consensus TcAAgNw  -N12-13- gcCGAtaNc
```

**Supplementary Figure S2. Identification of putative flagellar promoters.** Sequences from the upstream regions of selected flagellar genes from *P. putida* KT2440 and F1, *P. syringae* DC3000 and B728a, *P.*

*savastanoi* 1448a, *P. fluorescens* SBW25 and *P. protegens* Pf-5 with the putative promoters aligned along with the established consensus for $\sigma^{54}$, FliA and $\sigma^{70}$-dependent promoters. No gaps are introduced, except when required for alignment of the promoter motifs. Promoter sequences are indicated in red and coding sequences of the upstream genes are indicated in blue.

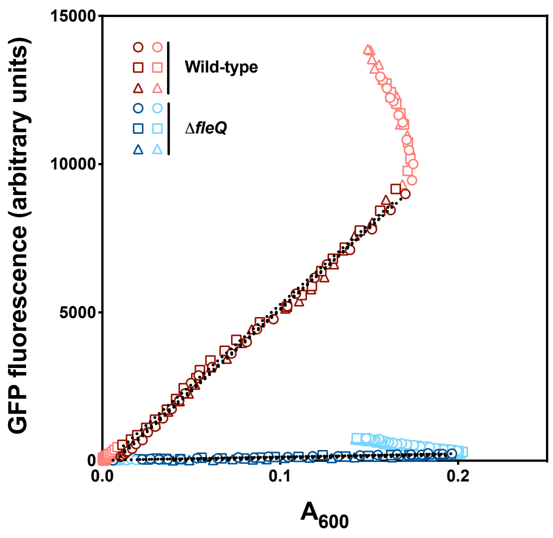


**Supplementary Figure S3. An example of the GFP fluorescence vs. absorbance plot.** Fluorescence data collected along the growth curves of the wild-type (red) and Δ*fleQ* (blue) strains bearing the P*flgB-gfp-lacZ* fusion plasmid pMRB272 were plotted against the $A_{600}$ data. Circles, squares and triangles denote three independent biological replicates, with the darker colors denoting the data used for the linear regression and the lighter colors denoting those not used. Dotted lines denote the linear fit obtained for the selected data of each replicate.

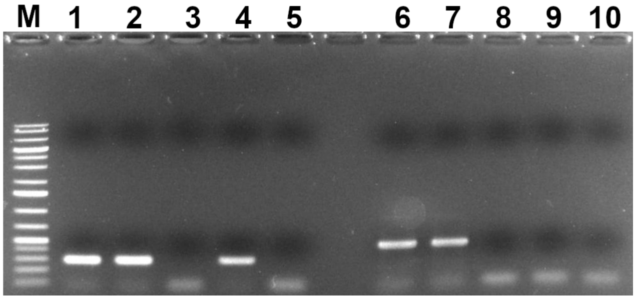


**Supplementary Figure S4. RT-PCR of *fleQ* expressed from its natural genomic context and ectopically.** Ethidium bromide-stained 1% agarose electrophoresis gel showing the amplification reaction products obtained with *fleQ*-specific (lanes 1-5) and *fliT*-specific (lanes 6-10) oligonucleotides. The template DNA was KT2442 genomic DNA (lanes 1 and 6), KT2442 cDNA (lanes 2 and 7), MRB130(Δflagella) cDNA (lanes 3 and 8), or MRB130/miniTn*7*-*nahR*-P*sal*-*fleQ* cDNA (lanes 4 and 9). Lanes 5 and 10 contain reactions performed with no template DNA. The picture shows one representative result of three biological replicates.

**A**

| | |
|---|---|
| ***PparC*** | TGGCAC-$N_5$-GTGCA-$N_{36}$-GTCAATTCCCCGTC |
| ***PflhF*** | GTCAAGATACCGCC-$N_{31}$-GTCGCGCCATCGTG-$N_9$-GTCGGCGTTGAGTC-$N_{32}$-TGGAAC-$N_5$-TTGCT |
| ***PflhA*** | TGGAAA-$N_5$-TTGCA-$N_{26}$-GTCAAAGTTTTGCA |
| ***PfliL*** | TGGCAT-$N_5$-TTGCT-$N_{36}$-GACGGATTATTGGC |
| ***PfliK*** | TGGCGC-$N_5$-TTGCC-$N_{87}$-GCCAAAACCTCGCG |
| ***PfliE*** | TGGCAC-$N_5$-TTGCT-$N_{26}$-GTCAAAAAAATGCG |
| ***PfleS*** | CGGCAC-$N_5$-TTGCT-$N_{15}$-CACCGTTTTATGAC |
| ***PfliD*** | TGGCAT-$N_5$-TTGAC-$N_{25}$-GTCAAGCAATTGGC |
| ***PflgF*** | GGCAAACATCCGCC-$N_{63}$-TGGTTC-$N_5$-TTGCT-$N_{23}$-GGCAGCTACGCGCA |
| ***PflgB*** | GCCGCTTTTGTCAC-$N_{18}$-GCCGCTTTTCTGGC-$N_{55}$-TGGCAC-$N_5$-TTGCT |
| ***PflgA*** | CGGCAC-$N_5$-TTGCT-$N_{26}$-GACATTTTCCCGAC |
| ***PflgZ*** | TGGCAA-$N_5$-CGTTA-$N_{18}$-GCCTGGAGAACAAT |

**Putative $\sigma^{54}$-dependent promoter**
**Putative FleQ binding site**

**B**

| | | |
|---|---|---|
| ***PparC*** | GACGGGGAATTGAC | (-) |
| ***PflhF*** | GTCAAGATACCGCC | (+) |
| | GTCGCGCCATCGTG | (+) |
| | GACTCAACGCCGAC | (-) |
| ***PflhA*** | TGCAAAACTTTGAC | (-) |
| ***PfliL*** | GCCAATAATCCGTC | (-) |
| ***PfliK*** | GCCAAAACCTCGCG | (+) |
| ***PfliE*** | GTCAAAAAAATGCG | (+) |
| ***PfleS*** | GTCATAAAACGGTG | (-) |
| ***PfliD*** | GTCAAGCAATTGGC | (+) |
| ***PflgF*** | GGCAAACATCCGCC | (+) |
| | GGCAGCTACGCGCA | (+) |
| ***PflgB*** | GTGACAAAAGCGGC | (-) |
| | GCCAGAAAACGGCC | (-) |
| ***PflgA*** | GTCGGGAAAATGTC | (-) |
| ***PflgZ*** | GCCTGGAGAACAAT | (+) |

**C**

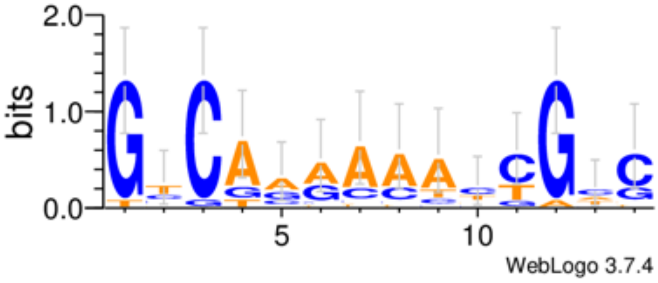


**Supplementary Figure S5. Putative FleQ binding sites at the Class II promoter regions. A.** Cartoon of each Class II promoter region showing the putative s54 promoter in red and the putative FleQ binding sites in green, along with the distances between the different *cis*-acting elements. **B. and C.** The putative FleQ binding sites, aligned, and the logo derived from the sequences of these sites using WebLogo 3.7.4. (Crooks *et al*., 2004) (+) and (-) indicate the orientation of the sequences as shown in **B.** relative to the promoter (coding and non-coding strand, respectively).

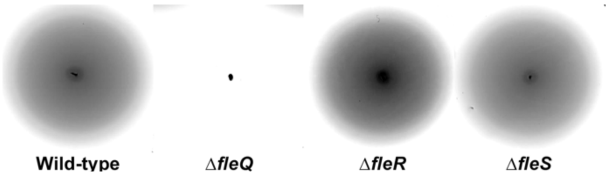


**Supplementary Figure S6. Swimming assay of the Δ*fleR* and Δ*fleS* mutants.** Swimming halos obtained in tryptone-soft agar plates with the wild-type strain KT2442 and its Δ*fleQ*, Δ*fleR* and Δ*fleS* derivatives. The images show representative halos from identical plates grown in parallel in the same conditions in triplicate. Cropping was performed maintaining the scale of the respective images.
